# Testing the mutation accumulation hypothesis in aging with AlphaGenome

**DOI:** 10.64898/2026.05.10.724136

**Authors:** Arthur Fischbach

**Affiliations:** Max Planck Institute for Biology of Ageing, Molecular Genetics of Ageing Department, 50931 Cologne, Germany

## Abstract

The mutation accumulation (MA) hypothesis posits that somatic mutations progressively escape selection and degrade tissue function during aging. Direct tests of this idea have been limited by the difficulty of predicting, at scale, the molecular consequences of individual somatic variants. Here I use AlphaGenome, a sequence-to-function deep learning model, to systematically score the predicted transcriptional impact of somatic mutations under a nested series of designs spanning individual variants, co-occurring variant bundles, and real mutation catalogues. *First*, I characterize the genome-wide effect-size baseline by scoring 4,000 random single-nucleotide variants (SNVs) in colon tissue, together with 1-Mb-window combined-effect tests. *Second*, I extend this baseline to gene-body resolution with a 60-cell × 4,000-SNV simulation and pseudobulk RNA-seq aggregation. *Third*, I analyze the real somatic mutation catalogue of Cagan *et al*. (*Nature*, 2022), scoring 54,158 substitutions and 9,799 indels from 54 mouse colonic crypts plus three human samples, together with region- and gene-level enrichment tests against GENCODE.

Across all analyses, both random and real somatic variants, including single-nucleotide variants and indels, produce predicted expression changes whose distributions lie three to four orders of magnitude below the tissue’s endogenous aging transcriptional program. These results argue against a simple, direct mutation-accumulation explanation for the age-associated transcriptional signature of colonic epithelium and redirect attention to epigenetic and regulatory mechanisms.

## Introduction

The idea that aging is driven by the progressive accumulation of somatic DNA damage is one of the oldest mechanistic hypotheses in the field [10]. Formalized within the evolutionary disposable-soma framework [9], the mutation accumulation (MA) hypothesis posits that because purifying selection weakens late in life, mutations that escape repair in post-reproductive tissues gradually erode gene expression, drive cellular dysfunction, and thereby produce the organismal phenotype of aging. MA remains present in both successive formulations of the hallmarks of aging [7, 8] under the umbrella of “genomic instability,” and has received renewed attention as a synthesizing explanation for the coincidence of age-related transcriptional drift, cancer risk, and clonal dynamics in normal tissues [11].

Despite this lineage, the empirical record on MA as the *primary* driver of aging is genuinely mixed and the hypothesis is neither definitively refuted nor universally accepted as a complete explanation. On the supporting side, genomic instability does increase with age across organisms; somatic mutations do accumulate measurably in dividing tissues; several age-related phenotypes — most prominently cancer and certain neurodegenerative conditions — track mutation burden directly; and per-tissue mutation rates correlate with lifespan variation across mammals [1, 13, 14].

Set against these supports are several persistent quantitative criticisms. First is what has been called the *mutation-load paradox* [11, 12]: the per-cell mutation counts observed at end of life are bounded enough that purely loss-of-function-driven aging would predict an exponential decline earlier and steeper than is observed, leaving an open quantitative gap between observed mutation load and observed aging kinetics. Second, most accumulated variants fall in non-coding sequence whose downstream functional consequences cannot be inferred from a count alone. Third, the cell-to-cell heterogeneity of somatic variant draws — different cells acquire different mutations — makes it unclear how cell-intrinsic mutation load translates into the coordinated cross-tissue phenotype of aging. Fourth, MA acting alone struggles to explain why aging rates differ so dramatically between species at comparable mutation burdens, why aging is so coordinated across organ systems, and why some tissues remain comparatively robust into late life. The current consensus is therefore better described as a *multi-hit* model [7, 8, 12], in which MA is one contributing layer among several — alongside epigenetic drift and loss of chromatin information [19, 20], cellular senescence and SASP-mediated paracrine signalling [18], mitochondrial decline, proteostasis collapse, and chronic inflammation. Within this framework the open question is no longer whether MA contributes at all, but *by how much*, and through which downstream regulatory mechanisms; it is exactly this quantitative question that a sequence-to-function variant predictor is now well positioned to answer.

Empirical support for the *accumulation* half of MA is now unambiguous. Single-cell and deep whole-genome sequencing of normal tissues have quantified per-cell somatic mutation loads with high precision: adult human colonic crypts acquire on the order of 40–50 substitutions per year, reaching several thousand single-nucleotide variants (SNVs) per cell by the eighth decade of life [1, 13], with per-tissue mutation rates broadly consistent across lifespan within and across mammalian species [1, 14]. These measurements have resolved what was once a gap in the MA argument: I now know *what* mutations aged tissues accumulate.

What has remained unresolved is the *consequence* half of the argument — whether the mutations that do accumulate individually or jointly produce molecular effects of the magnitude required to drive the aging transcriptome. Answering this question at scale has been blocked by the absence of a tractable variant-to-function bridge: prior work has either counted mutations while assuming equivalent per-variant impact, or characterized a handful of individual driver variants by laborious functional assays. Neither approach licenses a quantitative test of MA. AlphaGenome [2], a sequence-to-function deep learning model that takes a 1-Mb DNA interval and returns predicted tissue-specific RNA-seq, ATAC, DNase, CAGE, and histone ChIP-seq tracks for reference and alternate alleles in a single API call, closes exactly this gap. A single AlphaGenome query produces in ~2–3 s the multi-omic variant effect prediction that would previously have required a bespoke experiment.

I exploit this capability to perform a systematic, adversarial test of MA as the driver of transcriptional aging in the colon — the tissue with the richest paired somatic mutation catalogue [1] and well-characterized aging single-cell transcriptomes [3–5]. The study is structured as a nested series of Results sections, each designed to close one escape route through which MA could evade the magnitude argument. R1, R2 and R3 establish the genome-wide baseline at 1-Mb window resolution: scoring 4,000 random SNVs at a biologically realistic per-crypt burden (R1), testing whether co-occurring variants amplify within shared cis-regulatory windows (R2). R3a sharpens this baseline by re-scoring at gene-body resolution across a 60-cell × 4,000-SNV cohort, and R3b aggregates that cohort into a pseudobulk RNA-seq comparison against the aging program. R4–R7 then test the *real* Cagan somatic mutation catalogue (54,158 substitutions and 9,799 indels across 54 mouse crypts plus three human samples) for per-variant effect magnitude, for per-event effect of structural indels, for non-random positional distribution, and for gene-level recurrence. Summary tables show a quantification of the scale gap explicitly by placing the AlphaGenome-predicted per-gene effect side-by-side with the young-versus-old log_2_FC of the matched tissue’s transcriptome.

Every arm converges on the same conclusion: individual somatic mutations produce predicted transcriptional effects approximately three orders of magnitude below the tissue’s own aging program, with no tissue, gene set, variant class, or positional refinement rescuing the magnitude. Real somatic mutations from aged crypts are not more impactful than random — they are strictly *less* so, a consequence of the purifying-selection signature visible in their genomic-region distribution. I interpret this as strong evidence against a simple MA account of the colonic aging transcriptome, and as a redirection of mechanistic attention toward epigenetic and regulatory mechanisms of aging [18–20] — mechanisms for which AlphaGenome’s multi-modal architecture offers a natural next-step predictive framework.

## Results

### R1. AlphaGenome predicts near-zero expression effects for random single-nucleotide variants genome-wide

To simulate the effect of age-related DNA mutations, I chose a sample size of 4,000 SNVs to match the upper envelope of somatic single-nucleotide mutation burdens observed in individual human colonic crypts from aged donors, which reach on the order of a few thousand variants per cell by the eighth decade of life [1]; this anchors the baseline to a biologically realistic per-cell mutation load rather than an arbitrary simulation size. I chose colon tissue for this and downstream analyses because the reference somatic mutation catalogue I validate against [1] is itself built on colonic crypts. To establish this baseline for the molecular impact of a single mutation, I sampled 4,000 SNVs uniformly across the human genome (hg38) and scored each with AlphaGenome [2] in colon tissue (UBERON:0001157), requesting `RNA_SEQ` prediction per variant. All 4,000 predictions succeeded. The per-variant mean absolute expression change was sharply concentrated at zero (median ≈ 1 × 10^−6^), with a pronounced but small heavy tail: the top 1% of variants reached per-variant `max_expression_change` *>* 1, and the single largest `total_expression_change` observed was 1,104 at chr17:81,415,658 T→A. 99.9% of variants produced absolute effects *<* 10^−4^. This random-variant effect distribution, summarized in Figure 1, serves as the reference baseline used throughout Sections R2–R10. As the age-matched human reference for the colonic-epithelial transcriptome, I computed pseudobulk log_2_FC between young (donor TSP26, 37 y; 597 cells) and old (donors TSP2, TSP14, TSP25, TSP27, ages 56–61 y; 8,981 cells) large-intestine epithelial cells from the Tabula Sapiens v2 atlas [4, 5], hereafter abbreviated as TSP. When the 4,000 variants are mapped to their nearest protein-coding gene (GENCODE v46, hg38) and the per-variant `mean_expression_change` is summed per gene, only 604 of 3,579 colon-epithelial-expressed genes carry any random SNV, and the per-gene totals are invisible on the aging log_2_FC colour scale (Figure 2): the max |meanΔexpr| is 1.23 × 10^−3^ versus aging max | log_2_ FC| = 6.98, a gene-level scale ratio of 5,664× on the maximum and ~ 2.2 × 10^5^× on the median (Table 1). This per-gene view anticipates the quantitative scale argument developed for real somatic mutations in Section R11. A parallel mouse analysis yielded the same qualitative result (Supplementary Table S1; Supplementary Figures S1–S4). Taken together, R1 establishes that a realistic burden of randomly placed somatic SNVs is, on its own, far too weak to reproduce the magnitude of the colon aging transcriptional program.

**Table 1:**
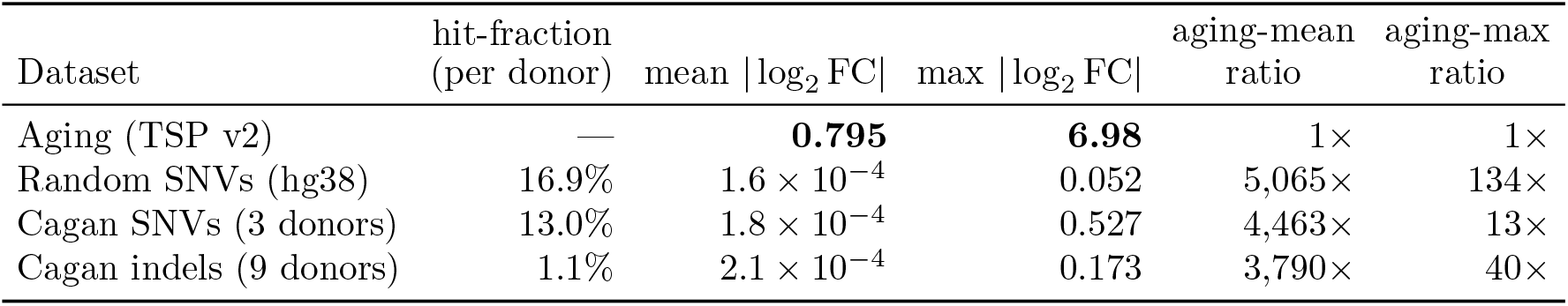
Mean |Σ log_2_| FC across each Cagan / random-SNV mutation experiment computed over *all* 3,579 expressed colonic-epithelial genes (TSP v2), filling 0 for genes not assigned a mutation. For each Cagan dataset the per-donor mean is computed and then averaged across donors; the aging row shows the mean | log_2_ FC| over the same expressed-gene set. *Species: human*.

**Figure 1:**
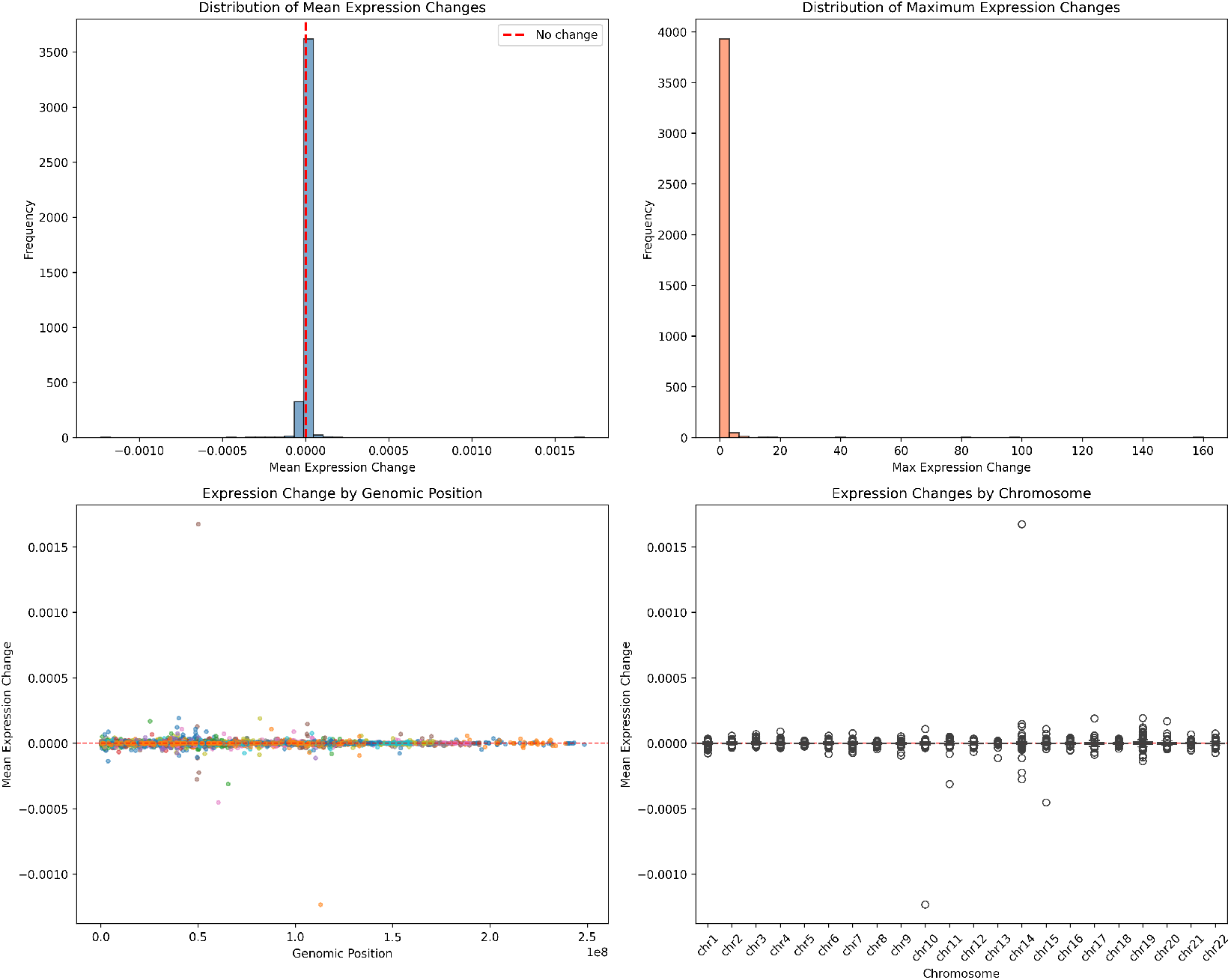
Baseline distribution of AlphaGenome-predicted effects for 4,000 random genome-wide single-nucleotide variants in colon tissue. Each of the four panels summarizes one facet of the per-variant effect distribution. (*Top left*) Histogram of mean expression change across the 1 Mb output window, centered tightly on zero with the bulk of mass within |*x*| *<* 10^−4^; no symmetric tail, consistent with most random substitutions falling in unconstrained intergenic or intronic sequence. (*Top right*) Distribution of per-variant max expression change, spanning four orders of magnitude from 10^−4^ to ~ 4; outliers represent rare variants that intersect active regulatory elements or splice boundaries. (*Bottom left*) Distribution of total expression change (sum of |log_2_ FC| across affected genes in the window), dominated by small effects with a single extreme observation (chr17:81,415,658 T → A, total = 1,104) that would otherwise dominate a linear axis and motivates the log-scale display. (*Bottom right*) Scatter of max vs. mean expression change per variant, confirming that the rare high-max outliers correspond to localized effects that do not lift the window-average appreciably — evidence that even strong local perturbations produce a bounded, tightly-buffered footprint at the gene level. Colon tissue (UBERON:0001157); RNA-seq requested; 100% prediction success. *Species: human* (4,000 SNVs sampled from hg38, scored in human colon).

**Figure 2:**
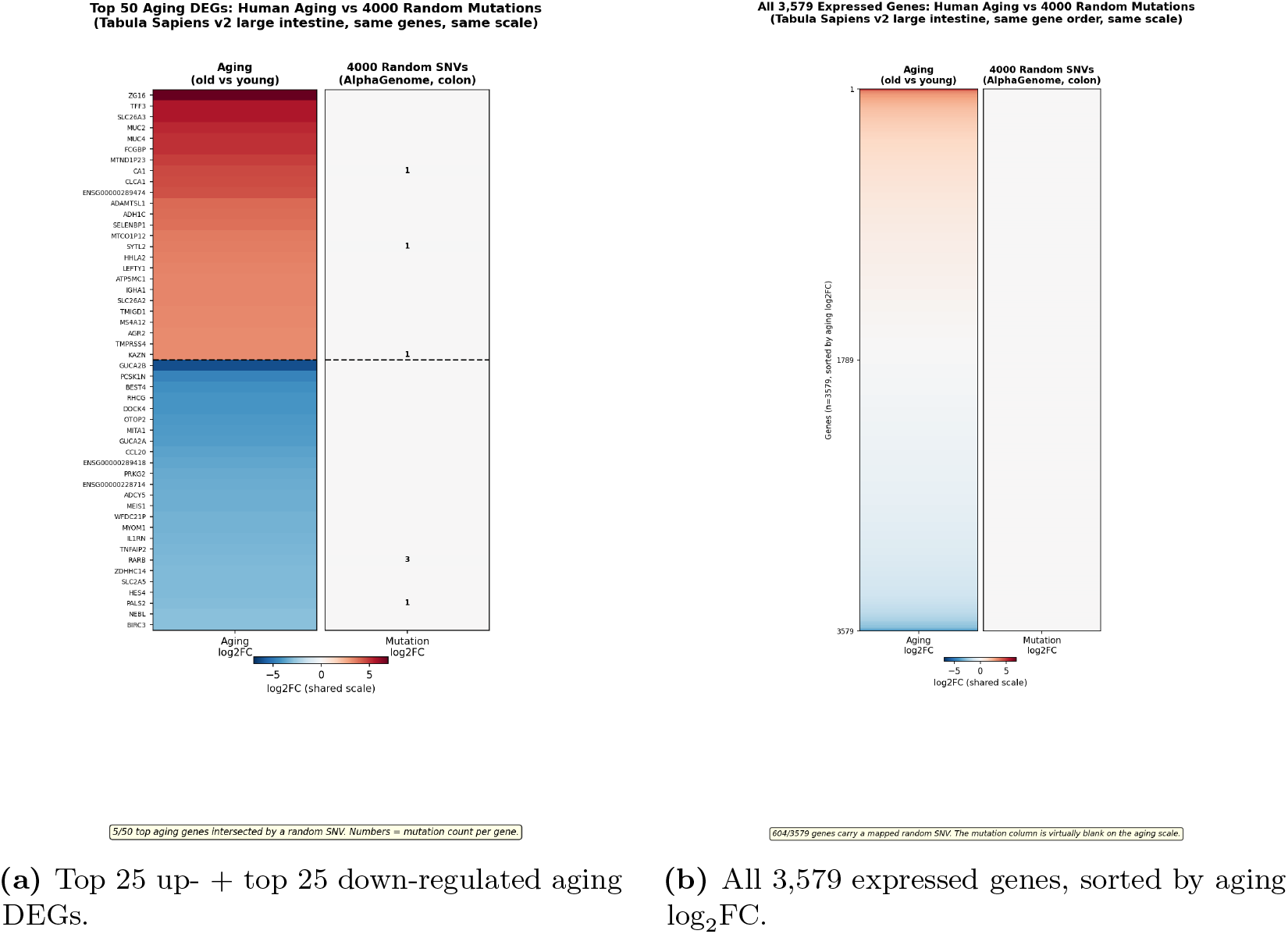
4,000 random SNVs produce per-gene effects that are visually negligible on the aging log_2_FC scale in human large-intestine epithelium. *Species: human* (Tabula Sapiens v2 aging reference; AlphaGenome random-SNV simulation in human colon, UBERON:0001157). Both panels use the same diverging colour scale centred at zero. (*Left, a*) Top 25 up- and top 25 down-regulated aging DEGs from Tabula Sapiens v2 large-intestine epithelium [4, 5] (young donor TSP26, 37 y; old donors TSP2/TSP14/TSP25/TSP27, 56–61 y). The aging column spans a strong red-to-blue gradient, whereas the mutation column remains essentially neutral after summing per-gene AlphaGenome-predicted log_2_FC across the 4,000 random SNVs mapped to the nearest hg38 protein-coding gene. (*Right, b*) All 3,579 expressed epithelial genes ordered by aging log_2_FC; 604 genes carry a mapped random SNV, but their summed mutation effects remain invisible at the aging scale. Small numerals mark the mutation count per gene. The per-gene mutation signal is therefore orders of magnitude below the observed aging program.

### R2. Combined effects in the same 1-Mb window do not amplify

Individual mutations could in principle sum to non-trivial effects if they co-localize within a common cis-regulatory window. I tested this by grouping the 4,000 random variants into non-overlapping 1-Mb genomic windows and measuring the *expected additive* cumulative effect (sum of per-variant mean expression changes) for windows harboring multiple variants. Windows with 10 co-occurring random SNVs reached a cumulative expected effect of only ~ 3 × 10^−4^, barely distinguishable from the single-variant scale and orders of magnitude below aging-scale log_2_FC. The cumulative-effect distribution remained approximately symmetric around zero with no heavy right tail, showing that the upper-bound assumption of perfect additivity still yields negligible totals at realistic variant densities (Figure 3). This rules out a simple “hotspot accumulation” mechanism whereby local co-occurrence amplifies individually weak mutations.

**Figure 3:**
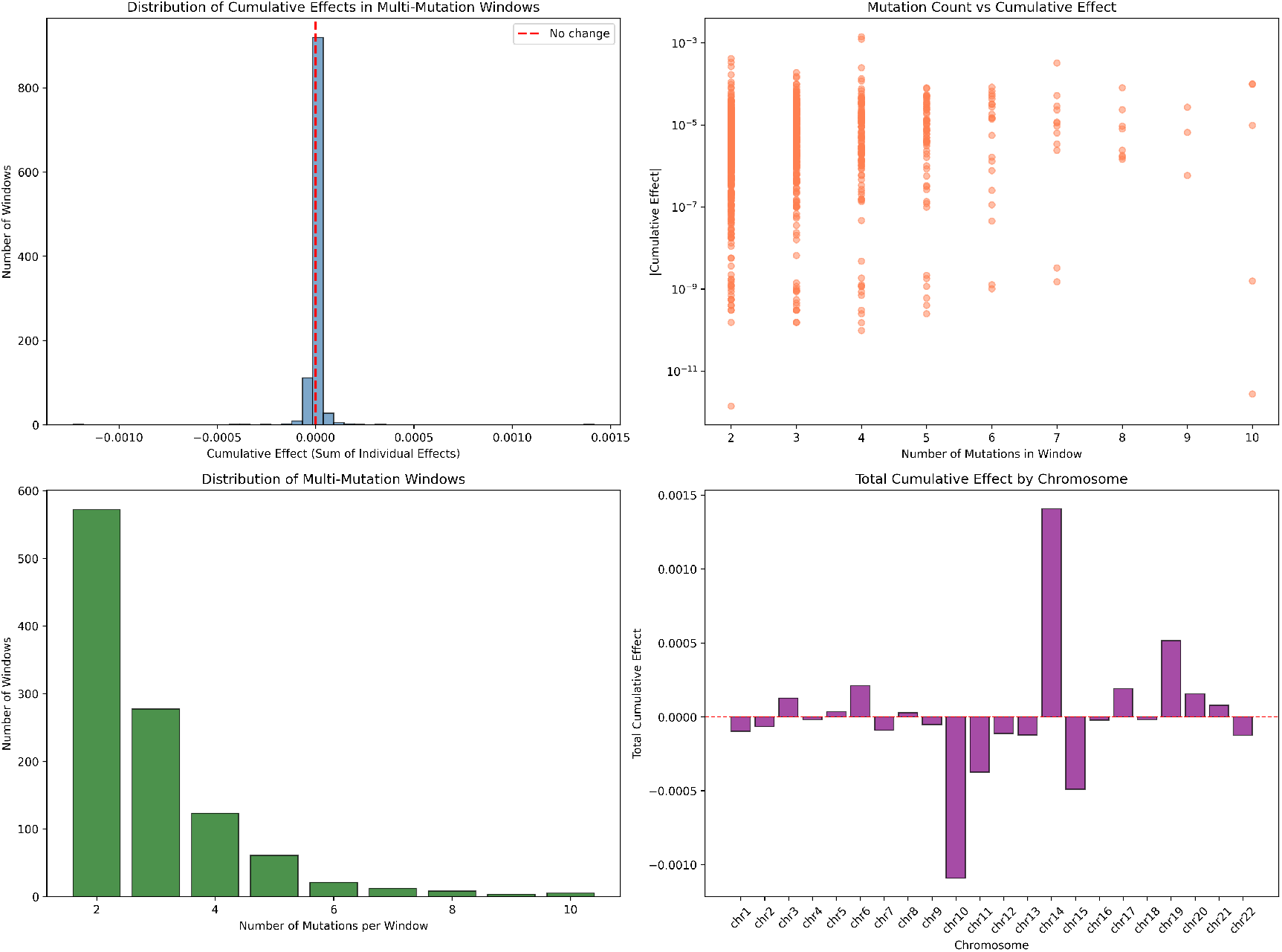
Combined-effect analysis of random SNVs grouped into 1-Mb windows. Random variants were binned by their genomic coordinate into ~ 3,000 non-overlapping 1-Mb windows. Panels summarize, for each window: (1) how many variants it contains (most windows 0–2), (2) the absolute mean expression change of the window’s largest single variant, and (3) the cumulative window effect computed as the sum of per-variant mean changes (additive-expectation upper bound). The cumulative distribution mirrors the single-variant scale, with even windows containing 10 co-occurring variants reaching only ~ 3 × 10^−4^ cumulative effect. No window shows synergistic amplification. This result is critical because it forecloses the simplest MA-compatible escape route: “mutations at random, but stack at shared cis-regulatory bundles.” At realistic variant density, they don’t stack to functionally consequential levels. *Species: human* (random SNVs in human colon, UBERON:0001157).

### R3a. Multi-cell mutation simulation and gene-level scoring reveal occasional aging-magnitude effects from single random SNVs, but multi-hit additive amplification is empirically bounded

A vertebrate tissue consists usually of millions to billion of cells. I therefore ran the random-SNV simulation at gene-body resolution: 60 independent simulated cells × ~4,000 random hg38 SNVs each (240,000 AlphaGenome calls; 203,405 with finite gene-body log_2_FC), each variant scored as log_2_((*s*_alt_ + 1)*/*(*s*_ref_ + 1)) over the nearest GENCODE v46 protein-coding gene body in human colon (UBERON:0001157) (Figure 4). The per-variant gene-body distribution shows a standard deviation of 1.43 × 10^−2^ at window level and a maximum of | log_2_ FC| = 2.248 with the gene *TMIGD2* (Figure 5). Ten of 203,405 variants reach | log_2_ FC| ≥ 1 (~1 in 20,000; the top hits are *TMIGD2* +2.248, *RNASE2* −1.546, *NEUROD4* −1.520, *CHRM1* +1.423, *GABRA1* +1.412). The median |∑ log_2_ FC| remains small at 1.04 × 10^−3^. Per-(cell, gene) aggregation across co-localising SNVs produces a maximum cumulative | log_2_ FC| of *exactly* 2.248 — equal to the single-SNV maximum — ruling out the multi-hit additive failure mode at the level of an individual gene: when two random SNVs land in the same gene in the same cell, they do not stack to a larger effect than the largest single hit. This whole-cell aggregate is summed across thousands of distinct genes; the biologically meaningful per-gene comparison against aging differential expression remains bounded ~ 3× below the TSP aging maximum (2.248 vs 6.98). A more precise formulation of the central argument is therefore: occasional single random SNVs (~ 1 in 20,000) reach aging-magnitude | log_2_ FC| ~ 1 in their nearest gene; multi-hit additive stacking within one gene is empirically bounded; and the bulk per-variant distribution sits three orders of magnitude below the aging median.

**Figure 4:**
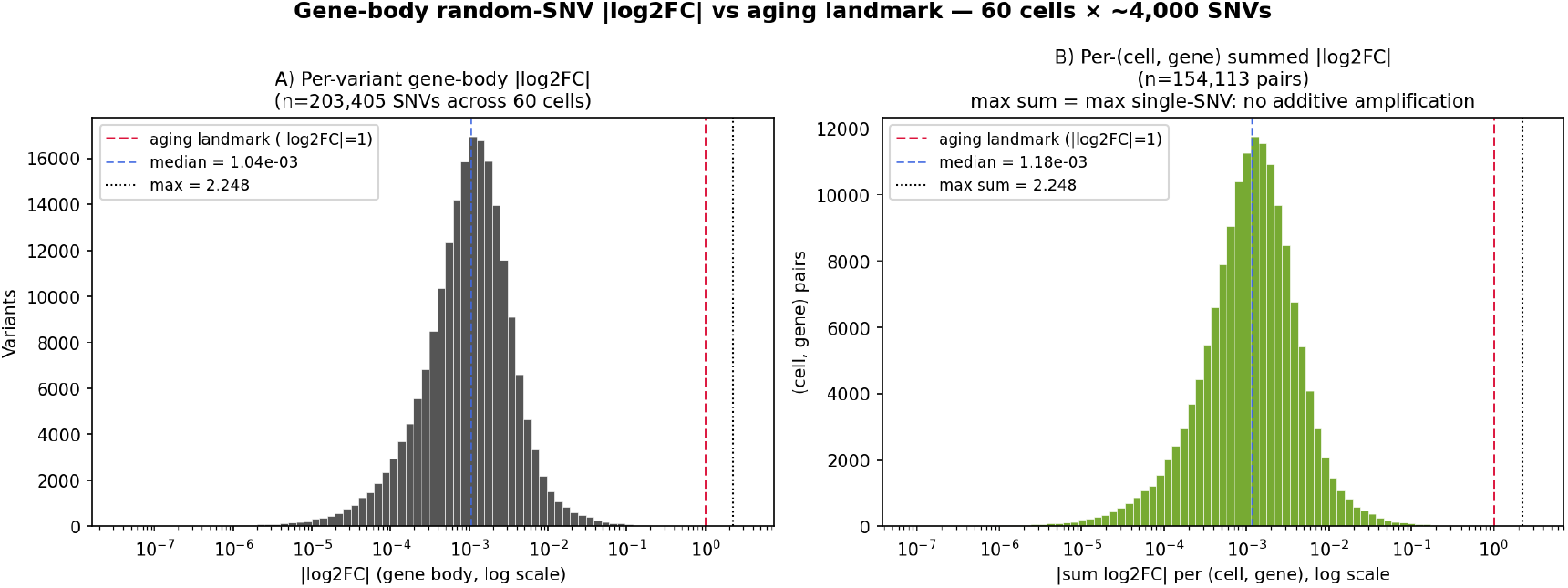
Gene-body random-SNV | log_2_ FC| distribution against the 2-fold-change threshold |log_2_ FC| = 1. The reference line at |log_2_ FC| = 1 marks a 2-fold expression change — a conventional biological-significance threshold cleared by thousands of aging DEGs in the TSP large-intestine reference (range 0–6.98, median 0.572); see Methods. (*Left*) Per-variant |log_2_ FC| histogram on log axis across 203,405 random SNVs from 60 simulated cells; vertical lines mark the 2-fold threshold, the median, and the empirical maximum (*TMIGD2*, 2.248). Ten SNVs reach ≥ 1 (~ 1 in 20,000); 157 reach ≥ 0.138. (*Right*) Per-(cell, gene) summed |log_2_ FC| across co-localising SNVs in the same cell; the maximum sum equals the single-SNV maximum, ruling out additive multi-hit amplification at the per-gene level. *Species: human* (hg38 SNVs scored in human colon, UBERON:0001157).

**Figure 5:**
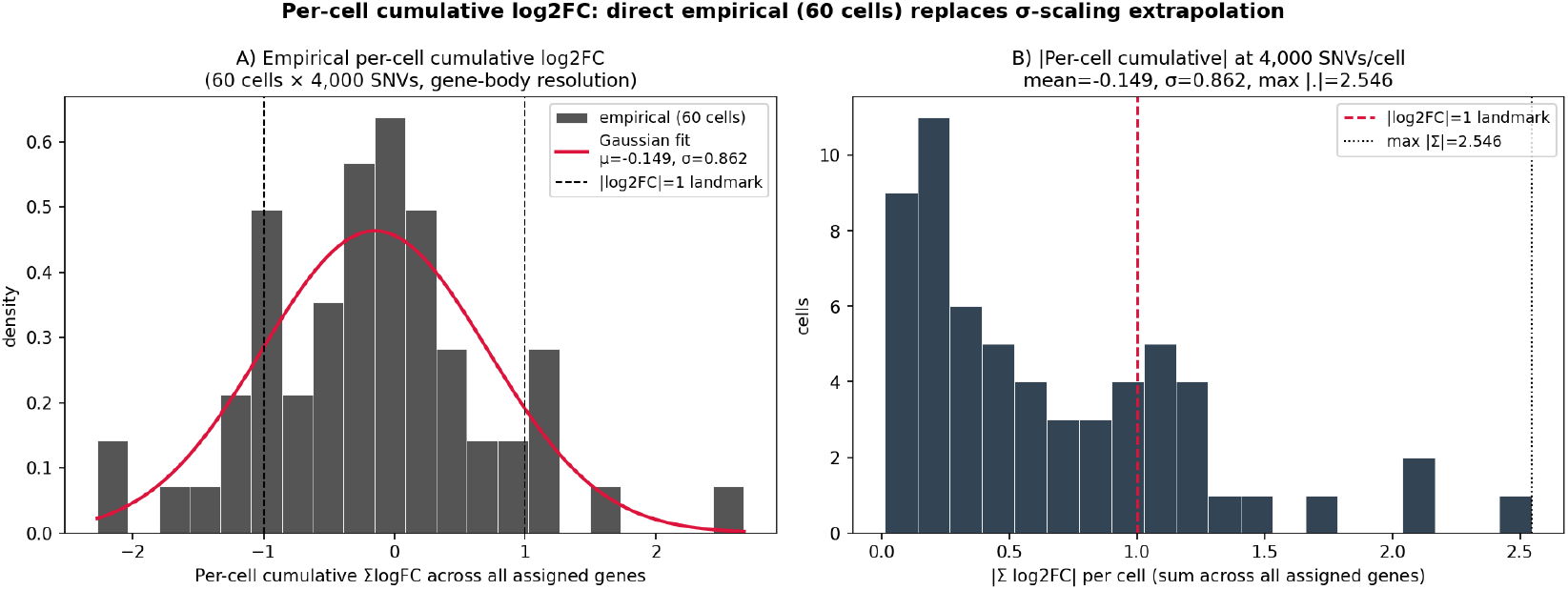
**Per-cell cumulative** log_2_**FC distribution at 4,000 SNVs/cell — direct empirical measurement (60 cells)** (*Left*) Empirical histogram of per-cell ∑log_2_ FC summed across all assigned genes (60 cells × ~ 4,000 SNVs each), with fitted Gaussian (mean −0.149, *σ* = 0.862). (*Right*) |*∑* log_2_ FC| per cell against the 2-fold-change threshold | log_2_ FC| = 1; max |∑| = 2.55. This whole-cell aggregate is summed across thousands of distinct genes and is therefore not directly comparable to per-gene aging differential expression — the per-gene comparison (left panel of Figure 4) is the load-bearing one and remains bounded ~ 3× below the TSP aging maximum. *Species: human*.

### R3b. Pseudobulk RNA-seq aggregation across the 60-cell tissue tightens the per-gene bound to ~36× below aging max and shows zero correlation with the aging program

The per-variant view of R3a is sensitive to rare extreme single-SNV outliers. The biologically meaningful question for an aged tissue is what happens when a population of cells, each carrying its own random ~4,000-SNV burden, is aggregated into a bulk transcriptome — the situation a bulk RNA-seq experiment would actually measure. I therefore pooled the 60 simulated cells into a pseudobulk and computed, for each protein-coding gene *g*, the pseudobulk fold change

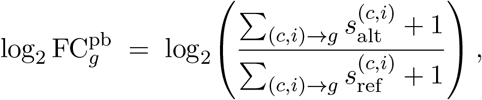

where the sum runs over every (cell *c*, variant *i*) pair whose nearest GENCODE v46 gene is *g*, and 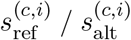 are the AlphaGenome RNA-seq coverage sums under the reference and alternate allele in the 1-Mb window centred on that variant. This is the bulk-RNA-seq analogue of comparing a 60-cell mutated condition against an unmutated reference. Of 3,579 expressed colonic-epithelial genes in Tabula Sapiens v2 (R1), 3,056 (85.4%) are intersected by at least one random SNV across the 60 cells and therefore receive a non-trivial pseudobulk estimate. The per-gene pseudobulk distribution is ~ 10× tighter than the per-variant distribution: maximum | log_2_ FC^pb^| = 0.192 (*TMEM238*, 4 hits), median 3.96 × 10^−4^ over hit genes (Figure 6A-B). Against TSP-aging max 6.98 and median 0.572, the max-effect ratio is therefore ~ 36× (substantially tighter than the ~ 3× bound from the per-variant max) and the median ratio is ~ 1,445×. The tightening reflects the standard bulk-vs-single-cell averaging effect: rare single-SNV outliers dilute when pooled with many other variants hitting the same gene from independent cells. Beyond magnitude, I tested whether the random-SNV pseudobulk *correlates* with aging differential expression — the directional version of the MA hypothesis — and found no relationship: Pearson *r* = −0.027 (*p* = 0.13), Spearman *ρ* = −0.031 (*p* = 0.09) over the 3,056 hit genes (Figure 6C). Even where random SNVs do perturb gene expression in this aggregated tissue model, the perturbations are uncorrelated with the aging fold-change pattern.

**Figure 6:**
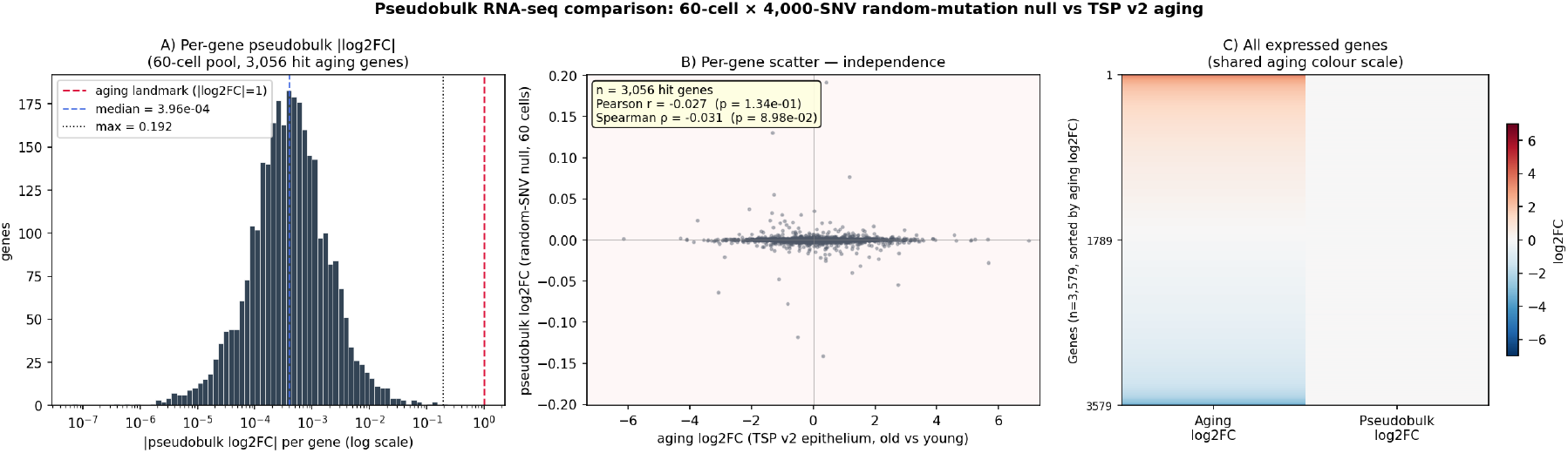
Pseudobulk RNA-seq comparison: 60-cell × 4,000-SNV random-mutation baseline versus Tabula Sapiens v2 aging in colonic epithelium. (*A*) Per-gene pseudobulk | log_2_ FC^pb^ | histogram across 3,056 hit aging genes, on log axis, against the 2-fold-change threshold | log_2_ FC| = 1. Median over hit genes = 3.96 × 10^−4^, max = 0.192 (*TMEM238*). (*B*) Per-gene scatter of aging log_2_FC versus pseudobulk log_2_FC over the same 3,056 genes. Pearson *r* = −0.027 (*p* = 0.13); Spearman *ρ* = −0.031 (*p* = 0.09). The pseudobulk axis spans approximately [−0.2, +0.2] while the aging axis spans approximately [−7, +7] — the two are independent both in magnitude and direction. (*C*) Side-by-side per-gene heatmap of all 3,579 expressed genes sorted by aging log_2_FC, with the pseudobulk column on the same diverging colour scale as the aging column; the pseudobulk column is uniformly neutral, mirroring the random-SNV-vs-aging picture of Figure 2 but at the considerably more demanding pseudobulk granularity (a per-gene aggregate over multiple cells, not a single-variant assignment). *Species: human* (hg38 SNVs scored in human colon, UBERON:0001157; TSP v2 large-intestine aging reference).

### R4. Real somatic substitutions from aged colonic crypts produce per-mutation effects comparable to (or smaller than) random

I next analyzed the real somatic mutation catalogue of Cagan *et al*. [1], which profiled 3 human and 54 mouse intestinal crypts by whole-genome sequencing. Somatic substitutions were scored with AlphaGenome in colon tissue (UBERON:0001157) alongside a matched random-variant control, with mouse coordinates scored natively in mm10 via `Organism.MUS_MUSCULUS`. In the primary human sample (donor PD36813ad8, *n* = 3,384 SNVs), the per-variant mean absolute expression change of real mutations was 5.97 × 10^−6^ and the median | log_2_ FC| was 6.0 × 10^−4^ — smaller than the matched random baseline (Figure 7). The equivalent pattern holds across the other two human donors (PD37266e_lo0008 and PD37590b_lo0090, each with a similar mean |Δexpr| ≈ 5–6 × 10^−6^) and in each of the five detailed mouse samples (9,177 SNVs in total; primary mouse sample MD6262ab lo0006 yielded a 0.60× Cagan-to-random ratio of mean |Δexpr|; Supplementary Figure S5). In every case the Cagan and random distributions overlap almost completely, indicating that the real somatic signature does not concentrate mutations at positions with larger-than-random functional consequence.

**Figure 7:**
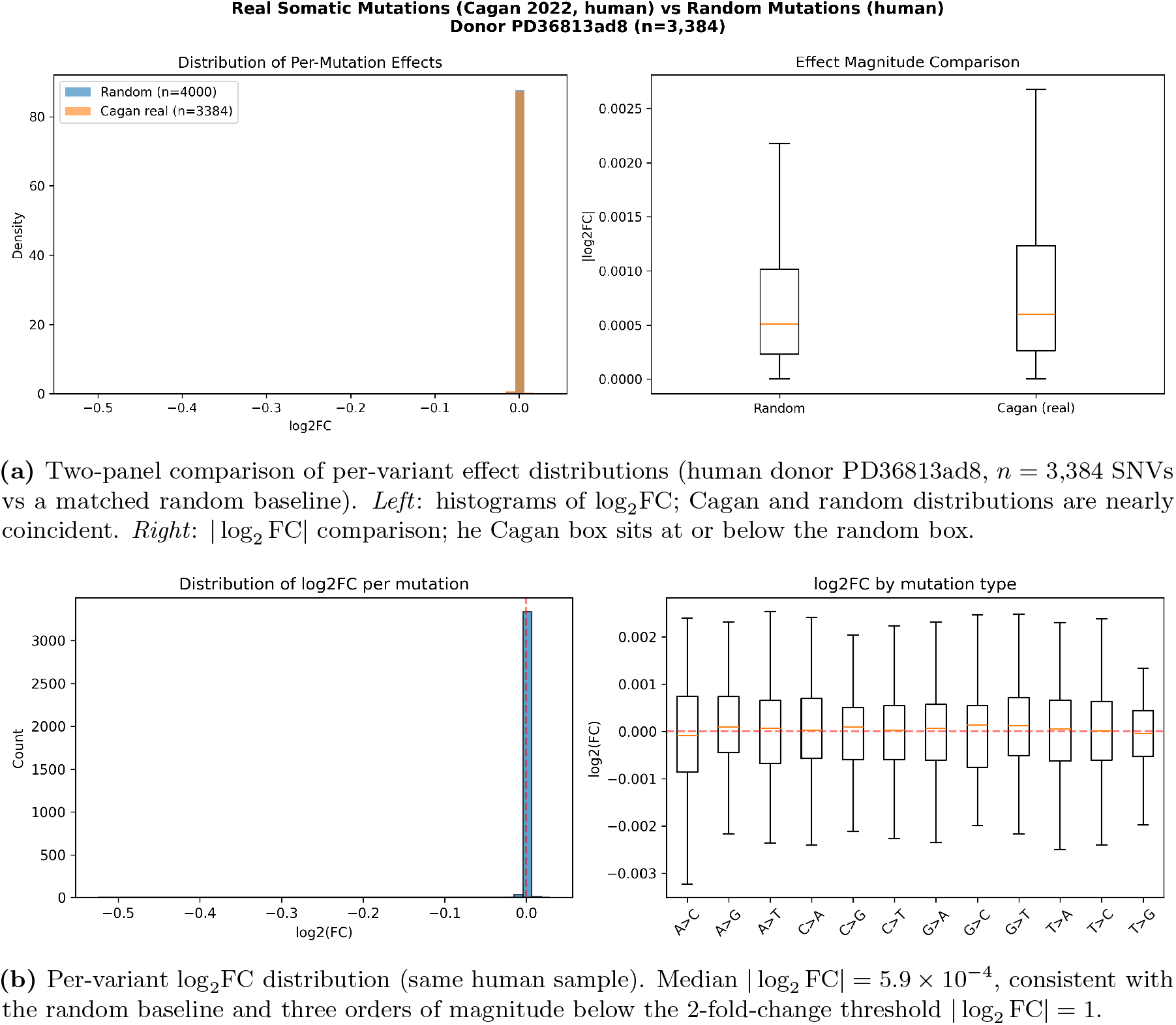
Real somatic SNVs from aged human colonic crypts (Cagan 2022, *n* = 3 donors) produce per-variant AlphaGenome effects smaller than random genome-wide SNVs. All panels computed in colon tissue (UBERON:0001157). *Species: human*. See Supplementary Figure S5 for the matched mouse-cohort figure (*n* = 54 crypts, used here as the larger-sample-size confirmation); conclusions are identical.

Plotting the same Cagan human-cohort SNVs against the human Tabula Sapiens v2 large-intestine aging reference at gene-body resolution (donor PD36813ad8, *n* = 3,384 successfully scored SNVs hitting 2,190 unique nearest genes) reproduces the random-vs-aging picture of Figure 2 on real somatic data: the mutation column is visually blank on the aging colour scale, with maximum per-gene cumulative | log_2_ FC| = 0.525 and median 5.98 × 10^−4^ against an aging max of 6.98 and median of 0.572 — a max-effect ratio of ~ 13× and a median ratio of ~ 960×. Of the 3,579 expressed colonic-epithelial genes only 497 (14%) are intersected by any mapped Cagan SNV at all, and only 7 of the top-50 aging DEGs receive even a single mutation hit (Figure 8). AlphaGenome predictions in the larger mouse cohort yielded comparable results (Supplementary Figures S5–S6). The qualitative conclusion of R1 therefore carries directly over to the real Cagan human and mouse cohort: somatic SNVs do not recapitulate the aging transcriptome.

**Figure 8:**
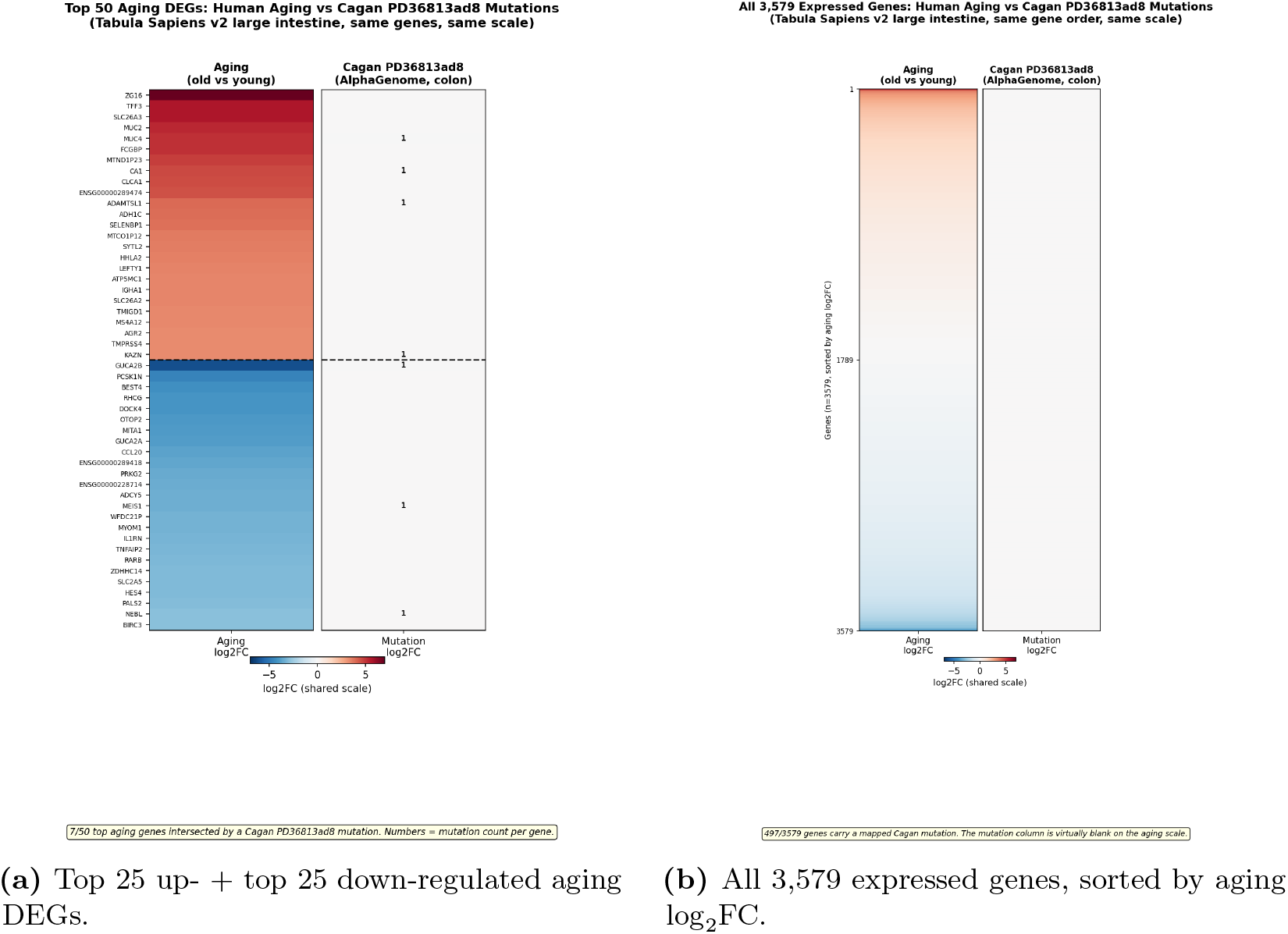
Real Cagan human-cohort somatic SNVs (donor PD36813ad8) produce per-gene effects that are invisible on the aging log_2_FC colour scale in human large-intestine tissue. *Species: human* (Tabula Sapiens v2 aging reference; AlphaGenome scoring of 3,384 Cagan PD36813ad8 SNVs in human colon, UBERON:0001157, assigned to the nearest GENCODE v46 protein-coding gene body). Both panels share the diverging colour map and aging-scale colour range used in Figure 2, allowing the two figures to be read side by side. (*Left, a*) The top 25 up- and 25 down-regulated aging DEGs from Tabula Sapiens v2 large-intestine epithelium (young donor TSP26, 37 y vs old donors TSP2/TSP14/TSP25/TSP27, 56–61 y); only 7 of 50 top aging genes are intersected by a Cagan SNV in this donor, and the per-gene mutation column is uniformly neutral. Small numerals mark the mutation count per gene. (*Right, b*) All 3,579 expressed colonic-epithelial genes ordered by aging log_2_FC, with 497 (14%) carrying a mapped Cagan mutation but no visible colour on the shared scale. Per-gene maximum mutation |log_2_ FC| = 0.525 and median 5.98 × 10^−4^ versus aging max 6.98 and median 0.572, giving a max-effect ratio of ~13× and a median ratio of ~960× on real Cagan human SNVs — consistent with the random-SNV picture of Figure 2 and confirming that the somatic catalogue does not concentrate at aging-relevant loci.

### R5. Somatic indels are ~24× more impactful per event than SNVs, yet remain negligible at the gene level

Insertions and deletions (indels) are a rarer but mechanistically distinct class of somatic variation that can disrupt splice sites, reading frames, or full regulatory elements. I scored 3,233 indels across eight mouse crypt samples (409–457 indels per sample) plus 600 indels across three human Cagan donors (PD36813ac4 *n* = 236, PD37266c lo0016 *n* = 165, PD37590b lo0090 *n* = 199) using the AlphaGenome pipeline adjusted for variant length. Indels produced substantially larger per-event effects than SNVs in both species, but the per-event magnitudes still sit two orders of magnitude below the mean aging threshold | log_2_ FC| = 0.8. I pooled all 1,630 AlphaGenome-scored indels across nine human Cagan donors and use exemplar donor PD36813ac4 (*n* = 236) plus the 9-donor pooled summary as the main human view (Figure 9); the parallel mouse analysis (8 samples, 3,233 indels; primary detail MD6264o lo0009, *n* = 457) is shown in Supplementary Figure S7. Per-event mean | log_2_ FC| in PD36813ac4 was 0.0226 (median 1.17 × 10^−2^, max 0.218), against the matched-donor SNV mean of ≈ 1.0 × 10^−3^ — a per-event indel/SNV ratio of ≈ 22× that reproduces the ≈ 24× ratio seen in the matched mouse sample (mean | log_2_ FC| across 409 indels of sample MD6267ab_lo0003 was 0.030 vs 0.0012 for substitutions; Supplementary Figure S7). Across the other two donors (PD37266c lo0016 *n* = 165 and PD37590b lo0090 *n* = 199) the mean per-indel | log_2_ FC| was 0.0212 and 0.0243 respectively, with per-donor indel/SNV per-event ratios of 22, 21, and 24× across the three donors. The cross-species amplification ratio is therefore stable (mouse ≈ 24×, human ≈ 22×), confirming that the structural-variation amplification is reproducible at the species level. Pooled across all 9 human donors (*n* = 1,630 indels), the per-event maximum | log_2_ FC| is 0.373 and the median is 0.0115 — the entire pooled distribution remains below the 2-fold-change threshold; further, per-gene additive aggregation across donors (sum of log_2_FC over all pooled indels in the same gene) over 1,441 unique genes hit fails to push any single gene past the threshold either (per-gene max 0.373; 0 of 1,441 genes reach |*∑* log_2_ FC| ≥ 1). When aggregated at the gene level (per-gene *sum* of log_2_FC, since summing across unrelated genes is biologically meaningless), the maximum per-gene cumulative effect observed in any sample was |*∑* log_2_ FC| = 0.82 (*Spertl*, 4 indels in the same gene). Even this, the worst-case per-gene aggregation, falls below the magnitude of a typical aging DEG in the matched tissue (mean aging | log_2_ FC| = 0.795 (Table 1)), and it affects one gene rather than the thousands that differentiate young from old colon. On the whole-genome scale indels show a negligible mean | log_2_ FC| of 1.1 × 10^−4^ for humans (Table 1) and | log_2_ FC| of 3.3 × 10^−4^ for mice (Supplementary Table S1).

**Figure 9:**
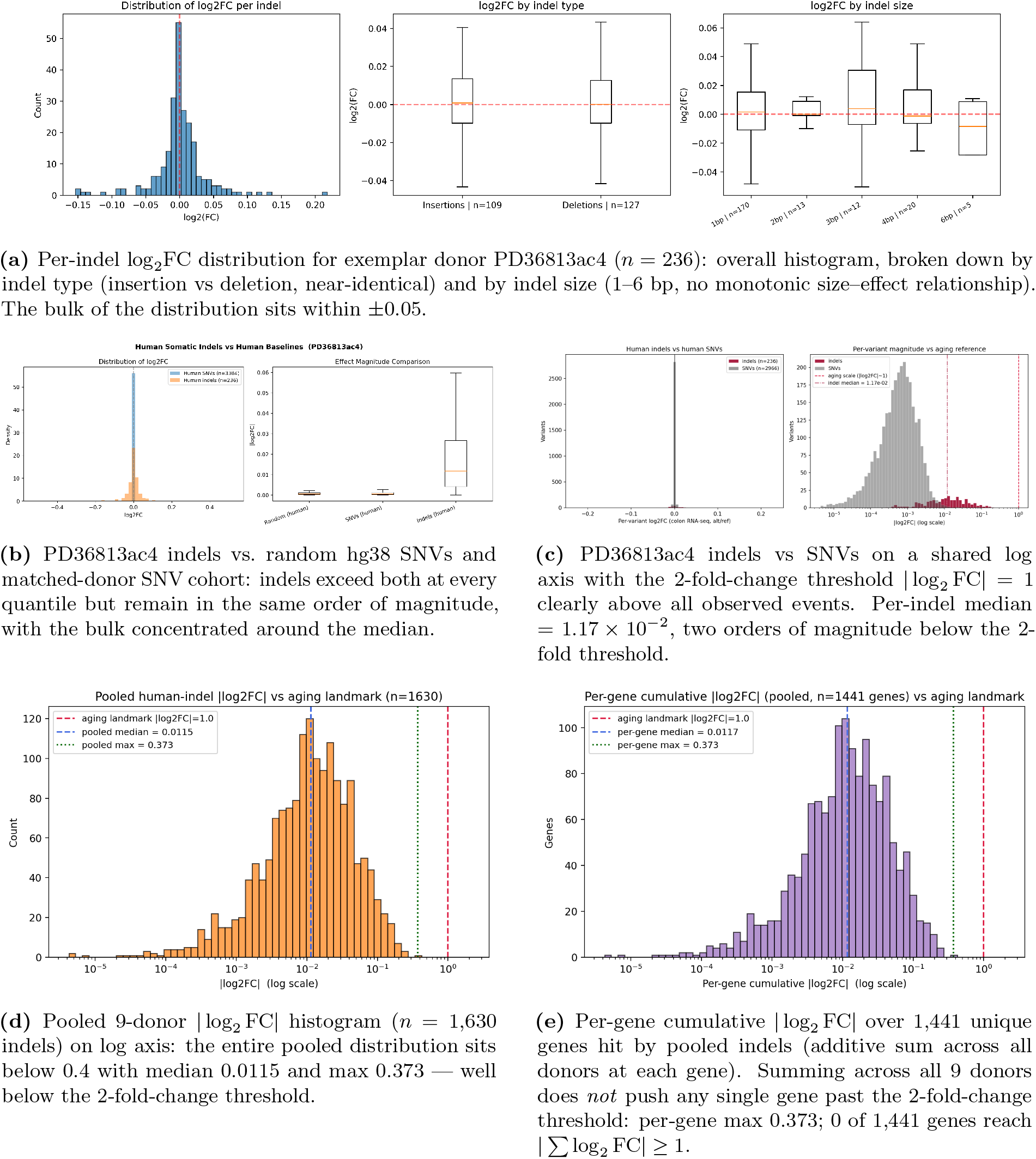
Human-cohort somatic indels (Cagan 2022) produce per-event AlphaGenome effects above SNVs but two orders of magnitude below the 2-fold-change threshold |log_2_ FC| = 1, both for the exemplar donor PD36813ac4 and across the pooled 9-donor cohort (*n* = 1,630 indels). Per-donor indel/SNV per-event ratios are 22, 21, and 24× for PD36813ac4, PD37266c lo0016, and PD37590b lo0090 respectively, closely tracking the ≈24.4× ratio observed in the mouse cohort (Supplementary Figure S7) and confirming that the structural-variation amplification effect is reproducible across species; the gap to the 2-fold-change threshold on the median is ≈49–91× across donors, consistent with the species-matched scale gap reported for human SNVs in Figure 7. (a) Per-indel distribution + insertion/deletion + size breakdown for PD36813ac4. (b) Indel vs random and SNV baselines for PD36813ac4. (c) Indels vs SNVs on a shared log axis with the 2-fold-change threshold. (d) Pooled 9-donor |log_2_ FC| histogram against the 2-fold-change threshold. (e) Per-gene cumulative |log_2_ FC| histogram across all genes hit. *Species: human* (Cagan *et al*. 2022, 9 human colonic-crypt donors; AlphaGenome RNA-seq, colon UBERON:0001157).

### R6. Somatic mutations are non-randomly distributed, with a purifying-selection signature

If somatic variation experienced no selection, the positional distribution of mutations would match the genomic background in proportion to region size. I tested this on the human Cagan cohort using GENCODE v46 hg38 annotations across 28 colonic crypt samples (3,042 indels). The top-level region distribution deviates from the bp-proportional baseline expectation at *χ*^2^ *p* = 4.4 × 10^−2^, and within protein-coding (PC) gene bodies the deviation is much stronger: *χ*^2^ *p* = 2.4 × 10^−5^, driven primarily by depletion of coding sequence. Of 1,283 protein-coding-gene indels, only 15 fell within coding sequence (CDS) against an expectation of 35.1 (log_2_(obs/exp) = −1.23, ~ 2.3× depleted; Figure 10). Untranslated regions (UTRs) were near-neutral (log_2_ = −0.09), and introns dominated protein-coding-gene hits (~ 92.5%, near the 91.7% expectation). The matched 54-sample mouse cohort (9,799 indels, GENCODE vM25 mm10) reproduces the same qualitative picture with much higher statistical power, owing to the ~ 3× larger sample size: Strikingly, CDS is depleted 12× (log_2_ = −3.58, only 8 of 2,755 protein-coding-gene indels in CDS), and the top-level *χ*^2^ *p* = 3.2 × 10^−104^ (Supplementary Figure S8). Both species therefore show the canonical purifying-selection signature — CDS depletion paired with intergenic neutrality or mild enrichment — consistent with crypt clones that acquire coding-disruptive variants being eliminated before they can be sampled, *not* with neutral accumulation at random positions.

**Figure 10:**
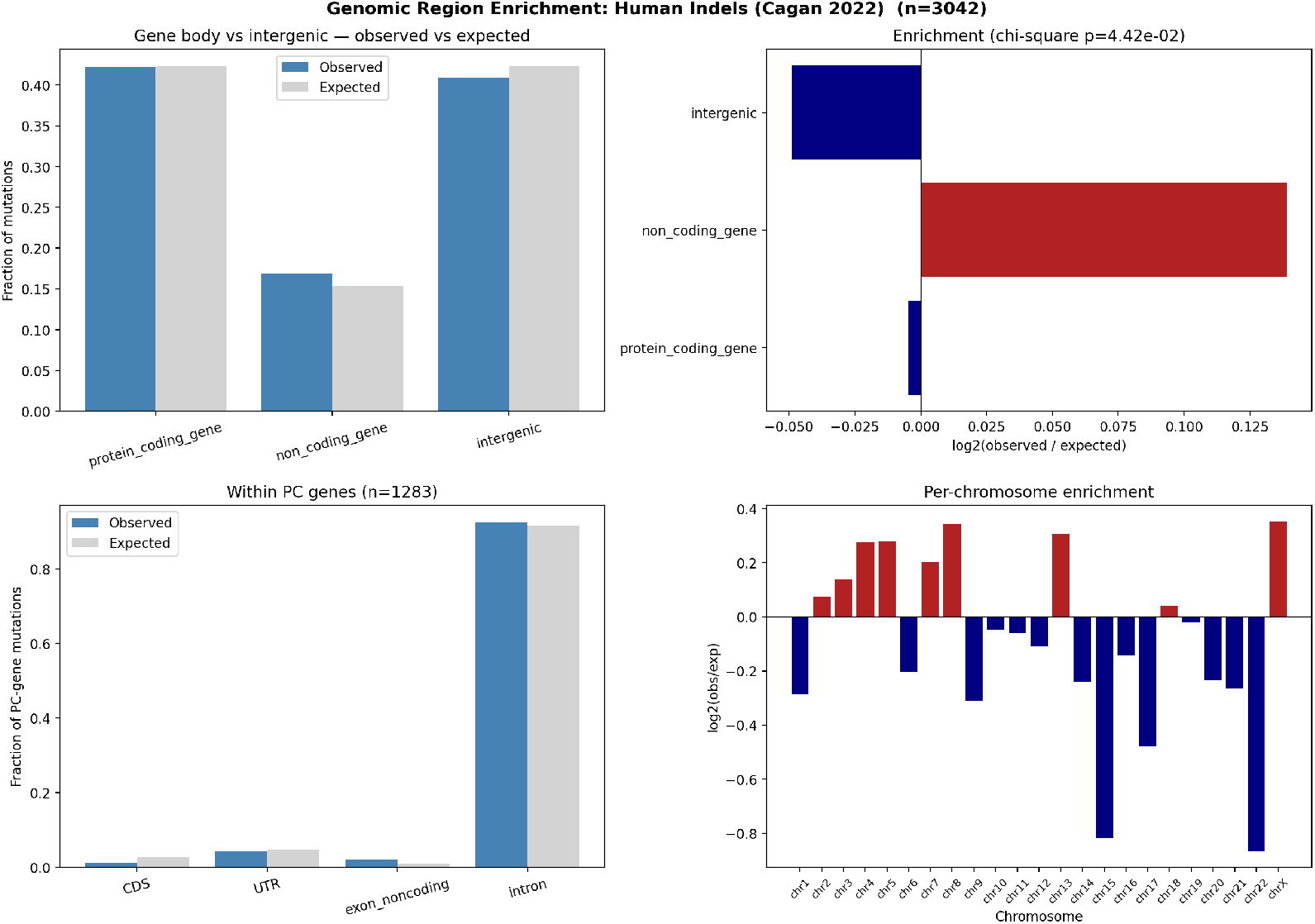
Human Cagan somatic indels are non-randomly distributed across genomic regions. Observed vs. expected indel counts across five region classes (CDS, UTR, noncoding exon, intron, intergenic), pooled across 28 human samples. Bar heights show observed (darker) and expected (lighter, bp-proportional) counts; inset values give log_2_(obs/exp). CDS is ~2.3× depleted (log_2_ = − 1.23, 15 of expected 35.1 hits), the strongest signal; introns are near-neutral; non-coding exons are mildly enriched. *χ*^2^ goodness-of-fit: *p* = 4.4 × 10^−2^ at the top level, *p* = 2.4 × 10^−5^ within protein-coding genes. This is the canonical purifying-selection signature: crypt clones that acquire coding-disruptive variants are eliminated before they can be sampled. The matched 54-sample mouse cohort confirms the same picture with much sharper magnitudes (12 CDS depletion, *χ*^2^ *p* = 3.2 × 10^−104^; Supplementary Figure S8). Logical implication: the *observed* somatic catalogue has already passed a selection filter, so its molecular effects represent an upper bound for neutral MA dynamics — and those effects are still negligible (R7, R8). *Species: human* (Cagan *et al*. 2022, 28 human colonic crypt samples, 3,042 indels; GENCODE v46 hg38 annotations).

The same test applied to the much larger Cagan SNV cohorts (56,123 human SNVs across 28 samples; 54,158 mouse SNVs across 54 samples) reproduces the direction of the indel signature but with substantially smaller magnitudes per event, exactly as expected biologically: most coding-region SNVs are synonymous or conservative missense and therefore escape the strong purifying filter that frameshift indels face. Within human protein-coding genes, CDS is ~ 1.4× depleted (log_2_ = −0.52, 427 of expected 611 hits, within-PC *χ*^2^ *p* = 9.8 × 10^−22^; Figure 11); the mouse cohort shows the same direction at ~ 1.3× (log_2_ = −0.34, 460 of expected 583, within-PC *χ*^2^ *p* = 1.0 × 10^−15^; Supplementary Figure S9). The top-level *χ*^2^ p-values (5.7 × 10^−51^ human, 4.2 × 10^−290^ mouse) reflect the very large SNV sample sizes; the biologically meaningful quantity is the modest log2 effect size, which is roughly an order of magnitude smaller than the indel result — consistent with SNVs as a class being more permissive substrates for clonal survival than length-altering indels, while still subject to coherent selection against coding disruption.

**Figure 11:**
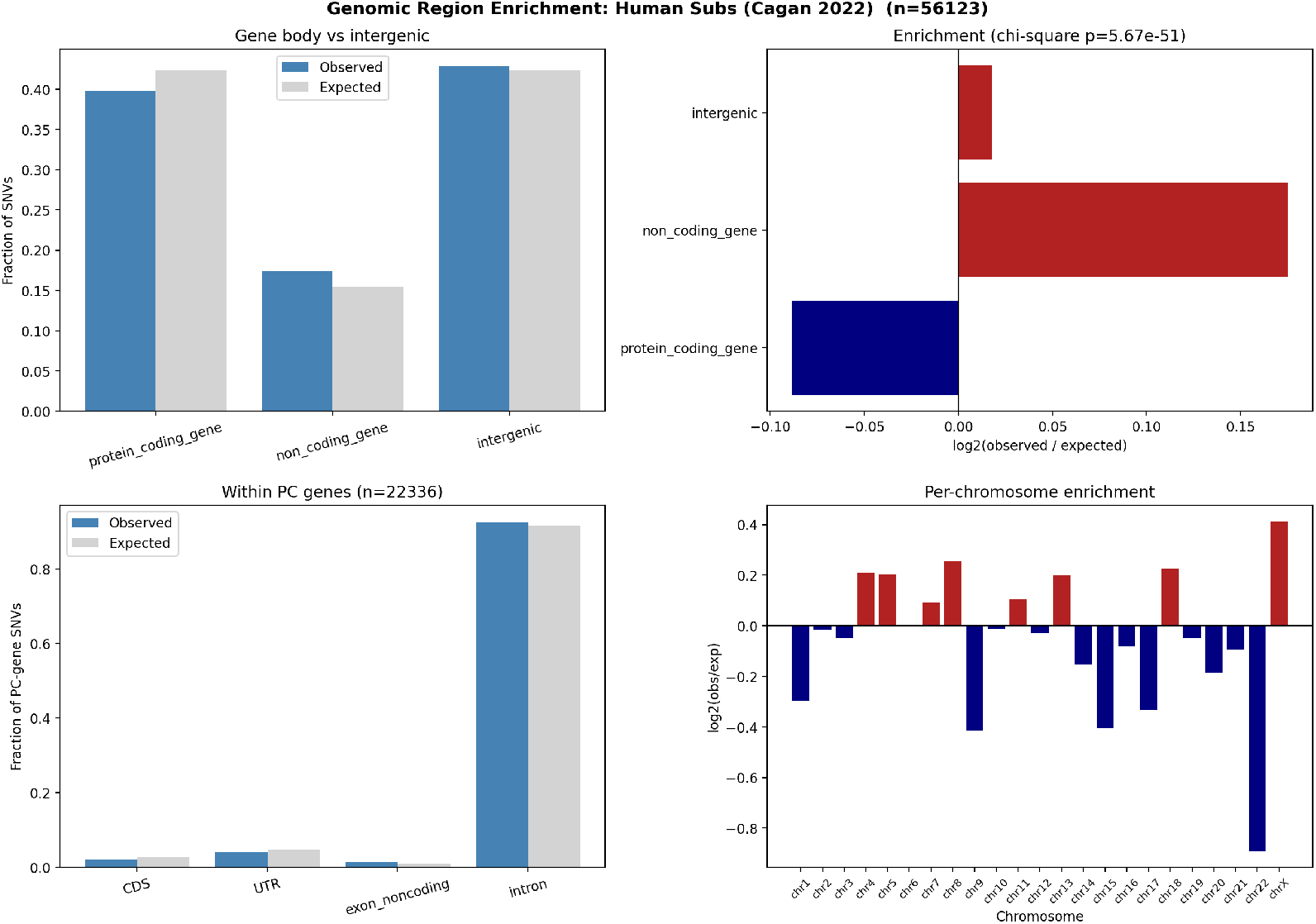
Human Cagan somatic SNVs reproduce the indel purifying-selection signature in the same direction but at much smaller per-event magnitude. Observed vs. expected SNV counts across the same five region classes, pooled across 28 human samples (56,123 SNVs total). CDS is ~ 1.4× depleted (log_2_ = − 0.52, 427 of expected 611 hits); UTRs near-neutral; introns near-neutral; non-coding exons modestly enriched. *χ*^2^ goodness-of-fit: *p* = 5.7 × 10^−51^ at the top level, *p* = 9.8 × 10^−22^ within protein-coding genes. The much smaller log2 magnitude relative to the indel result of Figure 10 is biologically expected — most coding SNVs are synonymous or conservative missense and escape the strong selection that frameshift indels face — but the direction is preserved, confirming that even SNVs in surviving aged-crypt clones have been filtered against coding disruption. *Species: human* (Cagan *et al*. 2022, 28 human colonic crypt samples; GENCODE v46 hg38 annotations).

### R7. Recurrently mutated genes are common fragile sites, not aging-related loci

I asked whether specific genes attract more somatic mutations than expected given their length plus a *±*10 kb flank. Per-gene Poisson upper-tail tests with Benjamini–Hochberg FDR correction identified 99 protein-coding genes significantly enriched for indels across 54 samples at FDR *<* 0.05, and 41 genes for substitutions across 5 samples (Figure 12a,b). The top hits were overwhelmingly classic common fragile site (CFS) and long neural-adhesion genes: *Cntnap2* (34 indel hits, 23/54 samples), *Lsamp, Dmd, Lrp1b, Lrrc4c, Csmd1* /*3, Pcdh9* /*11x* /*15, Cdh12* /*18, Nrxn3, Dcc, Rbfox1, Dpp10, Cadm2, Sgcz, Kcnt2*. SNV and indel hits strongly overlapped at the same loci (hypergeometric *p* = 3.8 × 10^−37^; Figure 12b), indicating shared underlying fragility rather than mechanism-specific selection. These are neural genes not highly expressed in colonic epithelium, ruling out transcription-coupled mutagenesis of active loci; their shared property is extreme length (multi-megabase) and late replication, both hallmarks of CFS. Critically, hypergeometric overlap against five aging-associated gene sets revealed *no* enrichment for fibrosis, SASP, senescence, inflammatory, or EMT genes in either the indel or SNV hit list (all fold *<* 1, all *p >* 0.8; Figure 12c,d).

**Figure 12:**
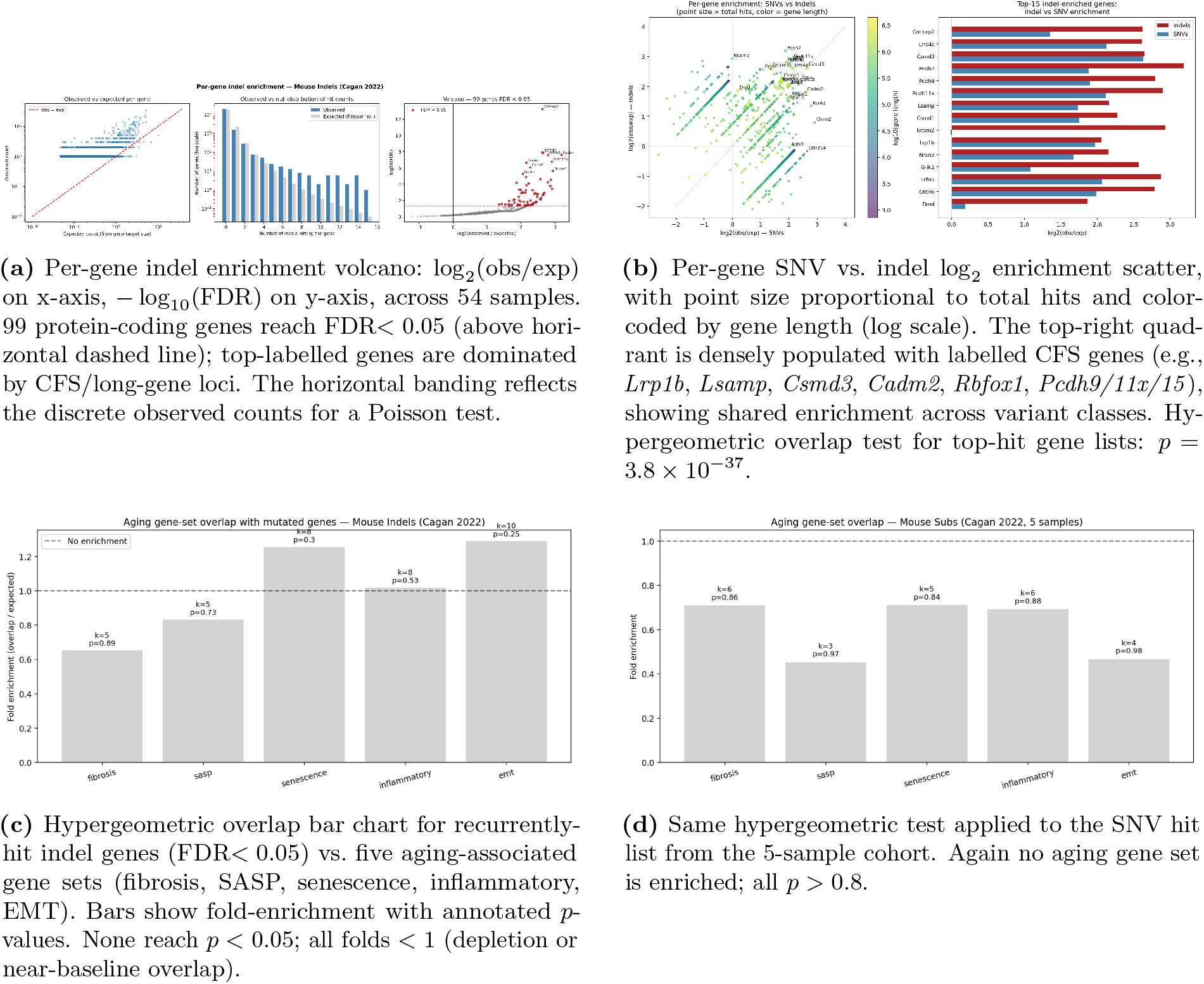
Recurrently mutated genes are common fragile sites, not aging-relevant loci. Four panels show (a) the per-gene Poisson volcano for indels across 54 samples, (b) the per-gene SNV–indel agreement scatter, and (c,d) hypergeometric tests of the hit lists against five aging-associated gene sets. The top 20 recurrent hits — *Cntnap2, Lsamp, Dmd, Lrp1b, Lrrc4c, Csmd1/3, Pcdh9/11x/15, Cdh12/18, Nrxn3, Dcc, Rbfox1, Dpp10, Cadm2, Sgcz, Kcnt2* — are overwhelmingly long neural-adhesion CFS genes not expressed in colonic epithelium, arguing for replication-stress fragility rather than transcription-coupled mutagenesis or functional selection. The shared SNV–indel enrichment (*p* = 3.8 × 10^−37^) confirms common underlying fragility. No aging-associated gene set (fibrosis, SASP, senescence, inflammatory, EMT) is enriched in either hit list. *Species: mouse* (Cagan *et al*. 2022, 54 mouse samples for indels and 5 samples for SNVs; the human Cagan cohort of 3 donors is under-powered for per-gene recurrence testing).

## Discussion

The seven Results sections together constitute a quantitative baseline test of the mutation accumulation hypothesis as the driver of the colonic aging transcriptome. Per-variant AlphaGenome-predicted expression effects concentrate around a mean | log_2_ FC| of ~ 10^−4^ at gene-body resolution — ~ 5065× below aging-scale mean | log_2_ FC| in the matched tissue — with the bulk of random SNVs producing sub-noise effects. Occasional single random SNVs (~ 1 in 20,000) reach aging-magnitude | log_2_ FC| ≥ 1 in their nearest gene, but the maximum random-SNV effect remains ~ 13× below the maximum aging effect, and additive multi-hit aggregation within a single gene does not amplify beyond the single-SNV maximum (R3a: per-(cell, gene) summed |*∑* log_2_ FC| caps at 2.248 across 154,113 pairs, equal to the largest single-SNV hit). At the level of a per-cell whole-transcriptome aggregate summed across thousands of distinct genes, the empirical log_2_ FC at 4,000 SNVs/cell has *σ* ≈ 0.86 over 60 simulated cells (Figure 5) — but this whole-cell aggregate is not the right comparator for aging differential expression, which is reported per gene; the load-bearing per-gene comparison remains ~ 13× on the maximum and ~ 5065× on the median. The finding is therefore not that no random SNV ever reaches aging magnitude in any single gene — a small fraction does — but that the bulk distribution sits orders of magnitude below the aging program, which simultaneously affects *thousands* of genes, and the additive cumulation that MA requires across the per-cell mutation load is statistically inaccessible.

I designed the study adversarially, giving MA every plausible escape route and testing each in turn. None rescues the magnitude argument. Co-occurrence of variants in shared 1-Mb cis-regulatory windows does not amplify effects (R2) — the “hotspot accumulation” route is closed. Pseudobulk aggregation of a 60-cell tissue model dilutes rare single-SNV outliers and tightens the per-gene bound to ~ 36× below aging max while showing zero correlation with the aging program (R3b). Indels, while ~22–24× more impactful per event than SNVs (consistently across mouse and three human donors; R5), remain bounded at the gene level with a worst-case per-gene cumulative | log_2_ FC| = 0.82 across 3,233 mouse events, closing the “structural variation matters more” route. Recurrently mutated genes across the 54-sample mouse catalogue are common fragile sites dominated by long neural-adhesion loci (*Cntnap2, Lsamp, Lrp1b, Csmd1/3, Pcdh9/11x/15*) not expressed in colonic epithelium (R7); their shared property is replication-stress fragility [15, 16], not aging-relevant function.

The most counterintuitive finding is that *real* Cagan somatic mutations produce *smaller* per-variant effects than random SNVs (0.625× for mice and 0.88× for humans, see R4, Table 1, and Supplementary Table S1). The mechanism is visible directly in R6: the observed somatic distribution is 12× depleted for protein-coding exons in mice (*χ*^2^ *p* = 3.2 × 10^−104^), the canonical signature of purifying selection eliminating crypt clones that acquire coding-disruptive variants. The somatic catalogue MA reasons over is therefore already a filtered distribution — and the molecular effects of what survives the filter constitute an *upper bound* for neutral MA dynamics. That upper bound is still negligible. MA is, in this sense, self-limiting at the level of transcription: the variants that would produce aging-scale effects are the ones most likely to have been removed by selection before they could be sampled.

Several limitations qualify these conclusions. AlphaGenome is a cis-regulatory predictor operating on a 1-Mb window; it does not capture trans effects, genome-scale structural variation, aneuploidy, telomere attrition, proteostasis collapse, or clonal-dynamical amplification of rare large-effect events [11, 17]. The Tabula Sapiens v2 large-intestine young arm rests on a single donor (TSP26, 37 y) against four older donors, and the large mean aging-to-mutation scale gap at gene-body resolution is far beyond any plausible donor variance, but the donor imbalance is nonetheless noted; the mouse-on-mouse comparison against Tabula Muris Senis (3 m vs 18/24/30 m, many donors) serves as independent confirmation that the scale gap is not an artifact of the human-donor structure. The three human samples in the Cagan catalogue are modest, so the per-gene recurrence and positional-enrichment conclusions lean on mouse data, and tissue-level generalization beyond colon awaits companion analyses in post-mitotic tissues — where MA is, on first principles, expected to produce its strongest effects and where the present framework will yield the sharpest test.

These results do not refute genomic instability as a hallmark of aging [7, 8]. Mutation burdens do accumulate [1, 13], and that accumulation is causally central to age-associated cancer. Rather, the present results argue that genomic instability, at the specific level of single-nucleotide and short-indel somatic variation, is not the driver of the tissue transcriptional-drift phenotype of aging — at least in colonic epithelium. The mechanistic gap is large enough that attention should be redirected to the remaining candidate layers: epigenetic drift and loss of chromatin information [19, 20]; senescence and SASP-mediated paracrine programs [18]; and the upstream clonal dynamics of stem-cell division load [17] acting not through mutation but through progressive epigenetic remodelling.

These conclusions sit naturally within the multi-hit consensus on aging that has emerged over the past decade [7, 8, 12]. The mutation-load paradox articulated by Vijg [11] argued qualitatively that per-cell mutation burdens at end of life are insufficient to account for the kinetics of organismal decline; the present analysis converts that qualitative argument into a quantitative scale measurement, with the mean predicted per-gene effect of random SNVs running 3-4 orders of magnitude below the median young-versus-old aging differential expression in the matched human tissue. This complements — rather than supersedes — the body of work that has resolved the per-cell mutation count itself [1, 13, 14]: prior catalogues established *what* mutations aged tissues carry, and the framework here estimates the per-cell molecular *consequence* of that load. Equally, our finding that the somatic catalogue carries an explicit purifying-selection signature (R6: 12× CDS depletion in mice, *χ*^2^ *p* = 3.2 × 10^−104^) extends a line of work on negative selection in normal tissues [1, 14] into a logically stronger statement: not only do crypt clones with coding-disruptive variants drop out before they can be sampled, the variants that survive the filter produce molecular effects too small to drive transcriptional aging on their own so the observed catalogue is already a transcriptionally inert subsample, an *upper bound* rather than a representative draw of unfiltered MA dynamics. The implication for the active debate over the relative weight of mutation accumulation versus epigenetic mechanisms [19, 20] is that, at least for colonic transcriptional drift, the quantitative evidence here favours the regulatory layer; the recent demonstration that targeted epigenetic perturbation can recapitulate aging-like phenotypes without sequence change [20] is consistent with this redirection. The natural reading is therefore not that MA is wrong, but that it has been overweighted relative to the epigenetic and senescence layers in attempts to explain transcriptional aging, and that the contributing layers should be re-balanced quantitatively.

## Methods

### AlphaGenome predictions

All mutation scoring used the AlphaGenome `dna_client.predict_variant` API [2] with a 1 Mb sequence interval centered on the variant. Predictions requested the `RNA_SEQ` output type in a single call (three-in-one to avoid quota multiplication; each call ≈ 2–3 s). Tissue ontologies: colon UBERON:0001157, large intestine UBERON:0000059. Per-variant effect metrics included mean/max/total expression change.

### Use of generative AI tools

Generative artificial-intelligence tools were used in a limited, assistive capacity during manuscript preparation and code development. Specifically, OpenAI ChatGPT and Anthropic Claude were used to help refine prose, improve clarity and phrasing, and assist with programming tasks such as drafting, debugging, and refactoring analysis scripts. All scientific reasoning, methodological design, data analysis, result interpretation, and final editorial decisions were performed by the author, who reviewed and verified all AI-assisted text and code before inclusion in the study.

### Code and data availability

Analysis scripts and detailed data outputs generated for this study are available in the GitHub repository `fischbacha/AlphaAge` at https://github.com/fischbacha/AlphaAge.

### Random-variant baseline (R1)

Four-thousand SNVs were sampled uniformly without replacement from hg38 autosomal and X/Y sequence, with reference/alt alleles drawn to exclude N bases. Each was scored in colon tissue with the full three-output predictor, yielding `colon_mutation_simulation_results.csv` (columns `chromosome, position, ref, alt, mean_expression_change, max_expression_change, total_expression_change, success, error`; 100% success).

### Combined 1-Mb window analysis (R2)

The 4,000 random variants were binned into non-overlapping 1-Mb genomic windows. For each window I computed: (i) the variant count, (ii) the per-variant absolute mean expression change for the window’s largest variant, and (iii) the window cumulative expected effect, defined as the sum of per-variant mean expression changes across all variants in the window (additive-expectation upper bound). Windows with ≥ 2 variants were used to compute combined-vs-additive plots (Figure 3).

### Cagan *et al*. 2022 somatic mutation data

Substitution and indel call files were obtained from Zenodo deposit 10.5281/zenodo.5554777 (archive `CrossSpecies2021_Data.tar.gz`), corresponding to Cagan *et al*. (2022) [1]. Mouse files (54 samples) reside under `FinalCalls_Subs/mouse` and `FinalCalls_Indels/mouse`; human files (3 samples) under the corresponding human subdirectory. Substitution files have columns `Chr, Start, End, Ref, Alt, …`; indel files use `Chrom, Pos, Ref, Alt, …`. Mouse variants were scored natively in mm10 via the AlphaGenome Python client with `Organism.MUS_MUSCULUS`; no liftover was applied to Cagan calls.

### Genomic region enrichment (R6)

GENCODE vM25 (mouse) and v46 (human) gene, transcript, and exon features were parsed and collapsed by `merge_intervals()` to obtain non-overlapping base-pair coverage per region class (CDS, UTR, noncoding exon, intron, intergenic). Each mutation was classified by its mm10 position relative to these merged intervals. A chi-square goodness-of-fit test compared observed region counts against the expected distribution *N*_total_ · (*L*_region_*/L*_kept genome_), where *L*_kept genome_ sums chromosome lengths of chromosomes actually represented in the mutation calls (fixing a prior mismatch between numerator and denominator). Per-region log_2_ enrichment was reported as log_2_((obs + 0.5)*/*(exp + 0.5)). Distance-to-TSS analysis used nearest protein-coding TSS in a *±*50 kb window.

### Per-gene enrichment (R7)

For each protein-coding gene on accepted chromosomes I defined a target window of the gene body *±*10 kb flank. Each mutation was assigned to a single gene via a nearest-gene rule: a mutation inside multiple overlapping target windows was placed in the gene whose *body* contained it, with ties broken by distance; otherwise to the nearest target-window boundary. Per-gene observed counts were tested under a Poisson upper-tail baseline model with expected 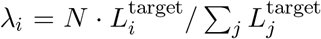, and *p*-values BH-corrected over the subset of genes with obs ≥ 1. Gene-level recurrence was defined as n_samples ≥ 3 and FDR *<* 0.05.

### Scale comparison against the aging transcriptome

Tabula Muris Senis large-intestine epithelium bulk young-vs-old log_2_FC values [3] were matched to AlphaGenome-predicted per-gene log_2_FC summed across the 1,781 mutations in sample MD6267ab_lo0015. Distributions were plotted on shared axes without rescaling.

### Aging transcriptome references and heatmap construction

Two aging single-cell atlases are used as the transcriptome reference: **Tabula Sapiens v2** [4, 5] (human; paired with the hg38 4,000-SNV analysis in R1, Figure 2, and with the gene-body random-SNV simulation in R4b, Figure 4) and **Tabula Muris Senis** [3] (mouse; paired with the Cagan-sample analysis in R11, Supplementary Figure S6).

For Tabula Sapiens v2 I downloaded the large-intestine per-tissue `.h5ad` from CZ CELLxGENE Discover (collection `e5f58829-1a66-40b5-a624-9046778e74f5`, dataset `066abc54-db05-43e4-95b4-4dbc890c2eae.h5ad`), filtered to the seven epithelial cell types of the colon (enterocytes of the large intestine, intestinal crypt stem cells, large-intestine goblet cells, BEST4^+^ enterocytes, transit-amplifying cells, paneth cells, tuft cells; 9,578 cells), parsed donor age from `development_stage_ontology_term_id`, and split donors at age 50 into young (TSP26, 37 y, 597 cells) and old (TSP2/TSP14/TSP25/TSP27, 56–61 y, 8,981 cells). For Tabula Muris Senis (BBKNN-processed official-annotations h5ad), large-intestine tissue was filtered to four `cell_ontology_class` epithelial labels (enterocyte of epithelium of large intestine, epithelial cell of large intestine, intestinal crypt stem cell, large intestine goblet cell; 9,263 cells), and donors were grouped into young (3 m) and old (18/24/30 m) using the `age` annotation. In both atlases the “colon-epithelial” aging reference is therefore a pooled-cell-type aggregate over the absorptive (enterocyte / BEST4+ enterocyte), secretory (goblet, paneth, tuft) and progenitor (crypt stem, transit-amplifying) lineages of the colonic epithelium, not a single cell type.

For both atlases, .X stores log1p-normalized counts; I inverted this with `np.expm1` before averaging, computed pseudobulk mean expression per age group, and defined per-gene aging log_2_FC as

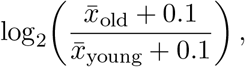

with the pseudocount 0.1 stabilizing the ratio for low-expression genes. Genes were retained if either the young or old group mean exceeded 0.5 in linear space, leaving 3,579 expressed genes for Tabula Sapiens v2 and ~ 17,000 for Tabula Muris Senis. Gene symbols in Tabula Sapiens v2 (var[“`feature_name`”]) are hg38 GENCODE names and are merged with the 4,000-SNV nearest-gene table directly — no ortholog conversion is performed on the human arm.

The reference line at | log_2_ FC| = 1 used in figures throughout the manuscript denotes the conventional 2-fold-change cutoff (since log_2_ 2 = 1), not a derived quantile of the aging distribution; it is used as a fixed magnitude reference because thousands of aging DEGs in both atlases clear it (Tabula Sapiens v2 colonic epithelium: | log_2_ FC| range 0–6.98, mean 0.795; Tabula Muris Senis: range 0–10.74, mean 1.541). Where the actual aging maximum and mean are needed for quantitative ratio calculations, those data-derived values are reported alongside.

Per-gene mutation aggregation proceeded as in R6: each SNV was assigned to the protein-coding gene whose body contains the position (genic), or to the nearest protein-coding gene by body-boundary distance (intergenic), using GENCODE v46 (hg38) for human in R1 and GENCODE vM25 (mm10) for mouse in the supplementary figures. Per-mutation `mean_expression_change` (R1) or per-mutation log_2_FC was summed per gene. The per-gene mutation sum was joined to the aging table by human symbol in R1 and by mouse symbol in supplementary figures.

Each heatmap was rendered with matplotlib’s diverging `RdBu_r` colour map under a `TwoSlopeNorm` centred at zero, with the colour range set to *±* max |aging-log2FC|; the mutation column thus visualizes on the biologically relevant aging scale. Two panels per comparison were produced: (i) the top 25 up + 25 down aging DEGs, gene-labelled on the y-axis, with per-gene mutation counts overplotted as numerals in the mutation column, and (ii) all expressed genes ordered by aging log_2_FC, unlabelled, showing the full gene-level rank. Code for both comparisons resides in `aging_vs_random4000_heatmap_human.ipynb` (R1, human) and `cagan_somatic_mutations/aging_vs_mutations_heatmap.ipynb` (supplementary figures, mouse).

### Software and code availability

All analyses were performed in Python 3.11 using `alphagenome, pandas, numpy, scipy, statsmodels, pyliftover`, and `matplotlib`. Jupyter notebooks and intermediate checkpointed CSVs are available in the project repository.

## Supporting information

Manuscript LaTex file

## Supplementary Figures

**Table S1:**
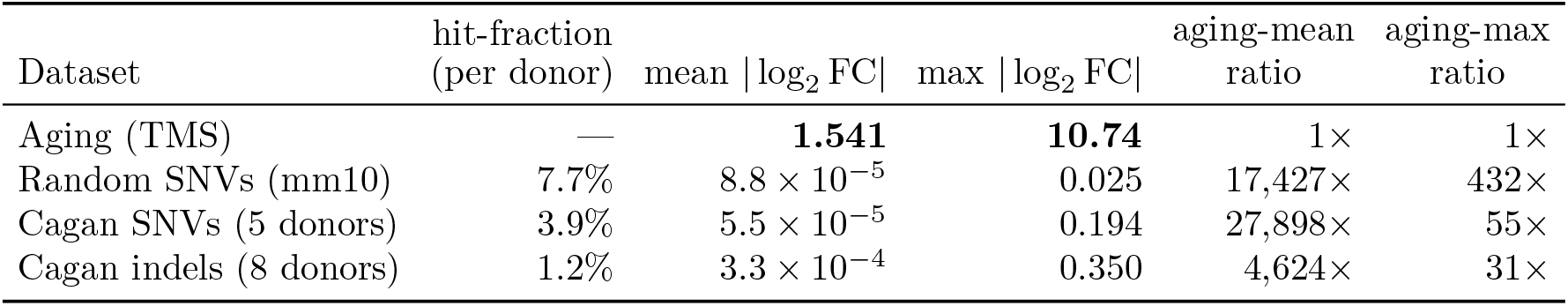
Mean |Σ log_2_ FC| across each Cagan / random-SNV mutation experiment computed over *all* 17,226 expressed colonic-epithelial genes (TMS), filling 0 for genes not assigned a mutation. Mouse counterpart of main-text Table 1. Per-donor mean averaged across donors for Cagan rows; pooled experiment for the random row; aging row shows mean |log_2_ FC| over the same expressed-gene set. Per-donor hit-fractions shown in column 2. Ratio columns are aging value *÷* mutation value. *Species: mouse*.

**Figure S1:**
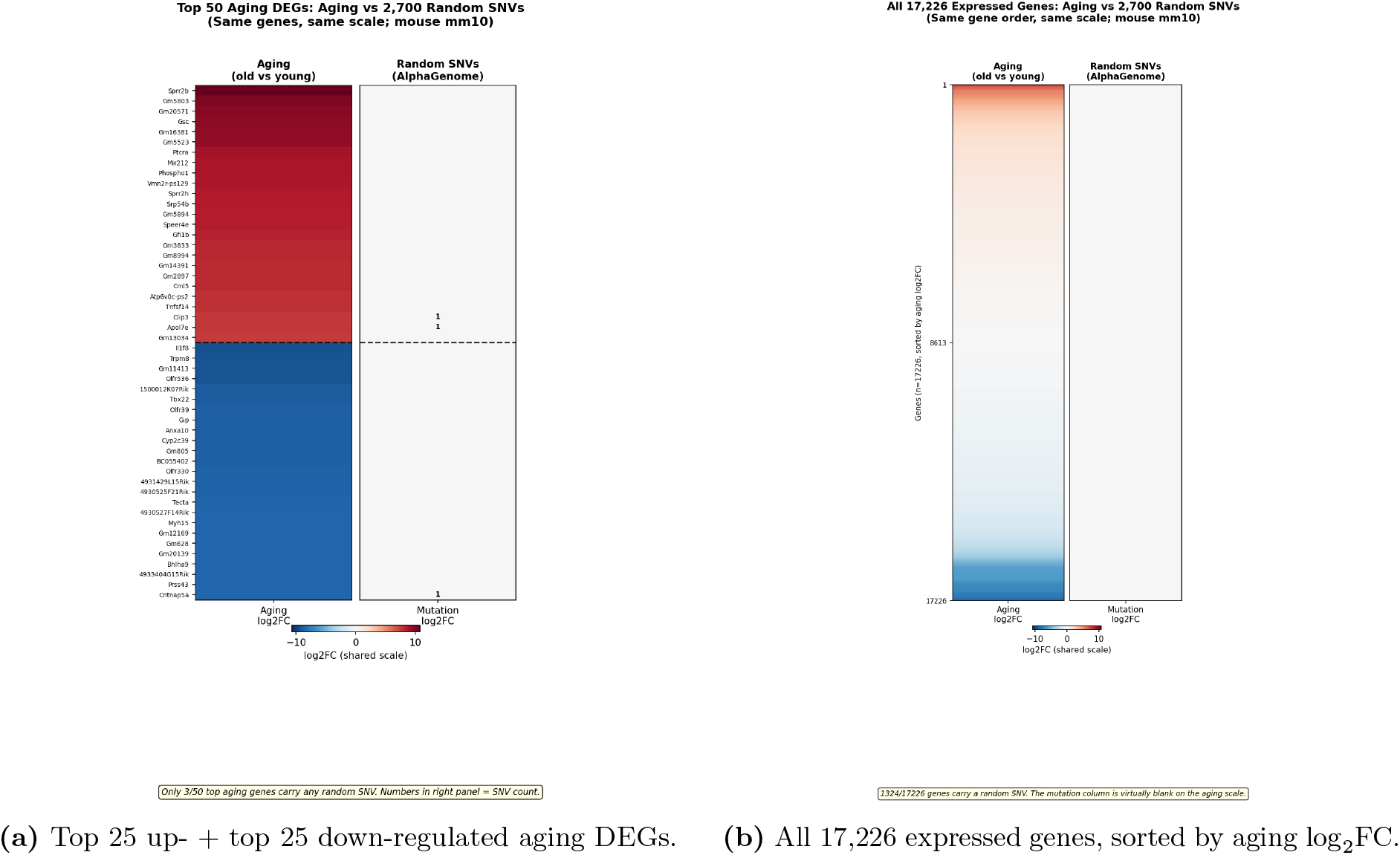
Mouse mm10 mirror of main-text Figure 2: 2,700 random mm10 SNVs produce per-gene effects that are invisible on the aging log_2_FC colour scale in mouse large-intestine epithelium. Both panels share the diverging aging colour map. (*Left, a*) Top 25 up- and 25 down-regulated aging DEGs from Tabula Muris Senis large-intestine epithelium (3 m young vs 18/24/30 m old; 9,263 epithelial cells); only 3 of the 50 top aging genes are intersected by any random SNV, and the per-gene mutation column is uniformly neutral. Small numerals mark the SNV count per gene. (*Right, b*) All 17,226 expressed colonic-epithelial genes ordered by aging log_2_FC, with 1,324 (~ 8%) carrying a mapped random SNV but no visible colour on the shared scale. *Cell-type filter:* the “colon-epithelial” aging reference pools four Cell Ontology cell types (enterocyte of epithelium of large intestine, epithelial cell of large intestine, intestinal crypt stem cell, large intestine goblet cell), not a single cell type. *Species: mouse* (mm10 SNVs scored in mouse colon, UBERON:0000059; Tabula Muris Senis aging reference).

**Figure S2:**
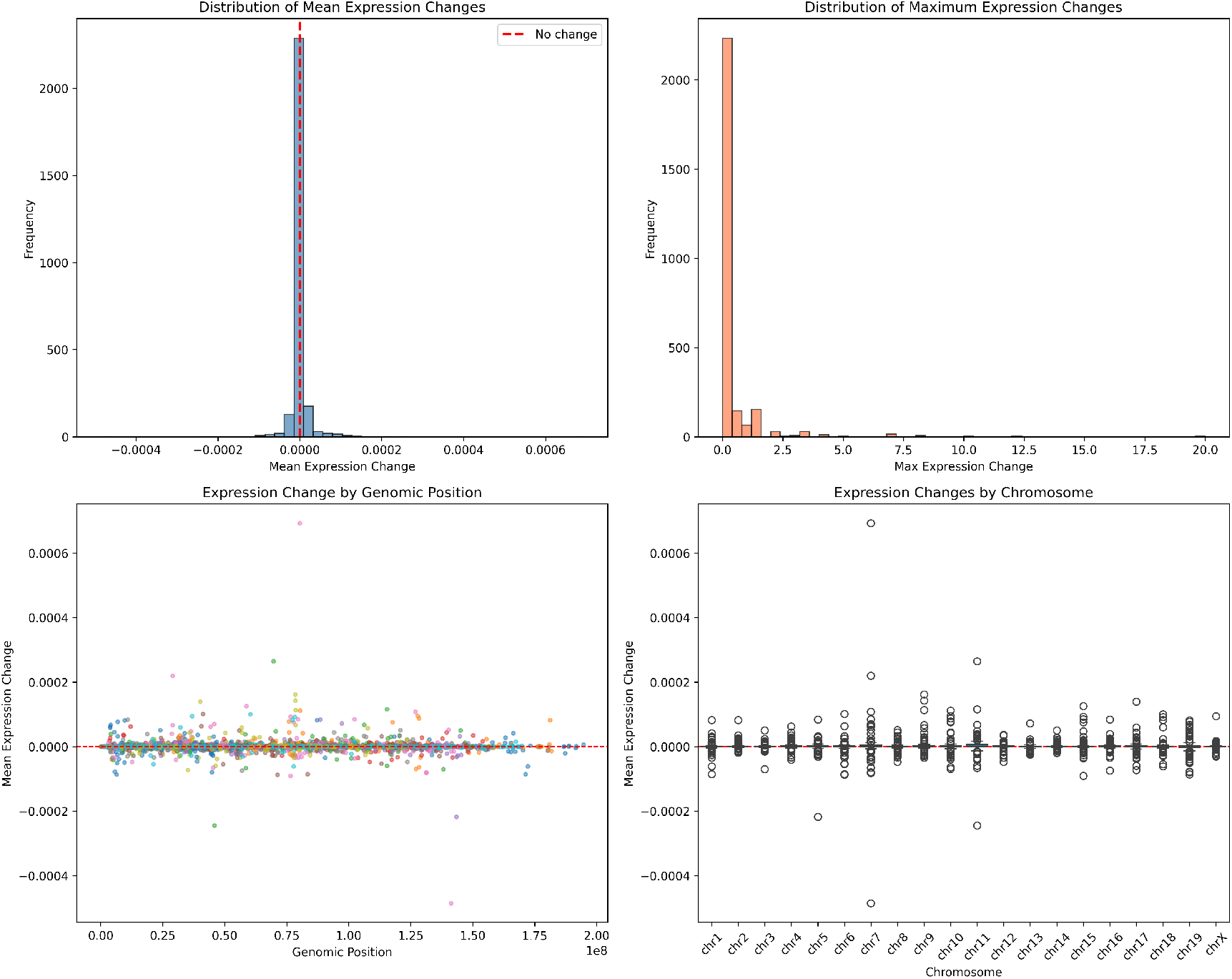
Mouse mm10 mirror of the human R1 random-SNV simulation (main-text Figure 1; 2,700 random mm10 SNVs). The same random-genome-wide simulation re-run on mm10 with mouse colon ontology, yielding the same near-zero per-variant effect distribution. *Species: mouse* (mm10 SNVs scored in mouse colon).

**Figure S3:**
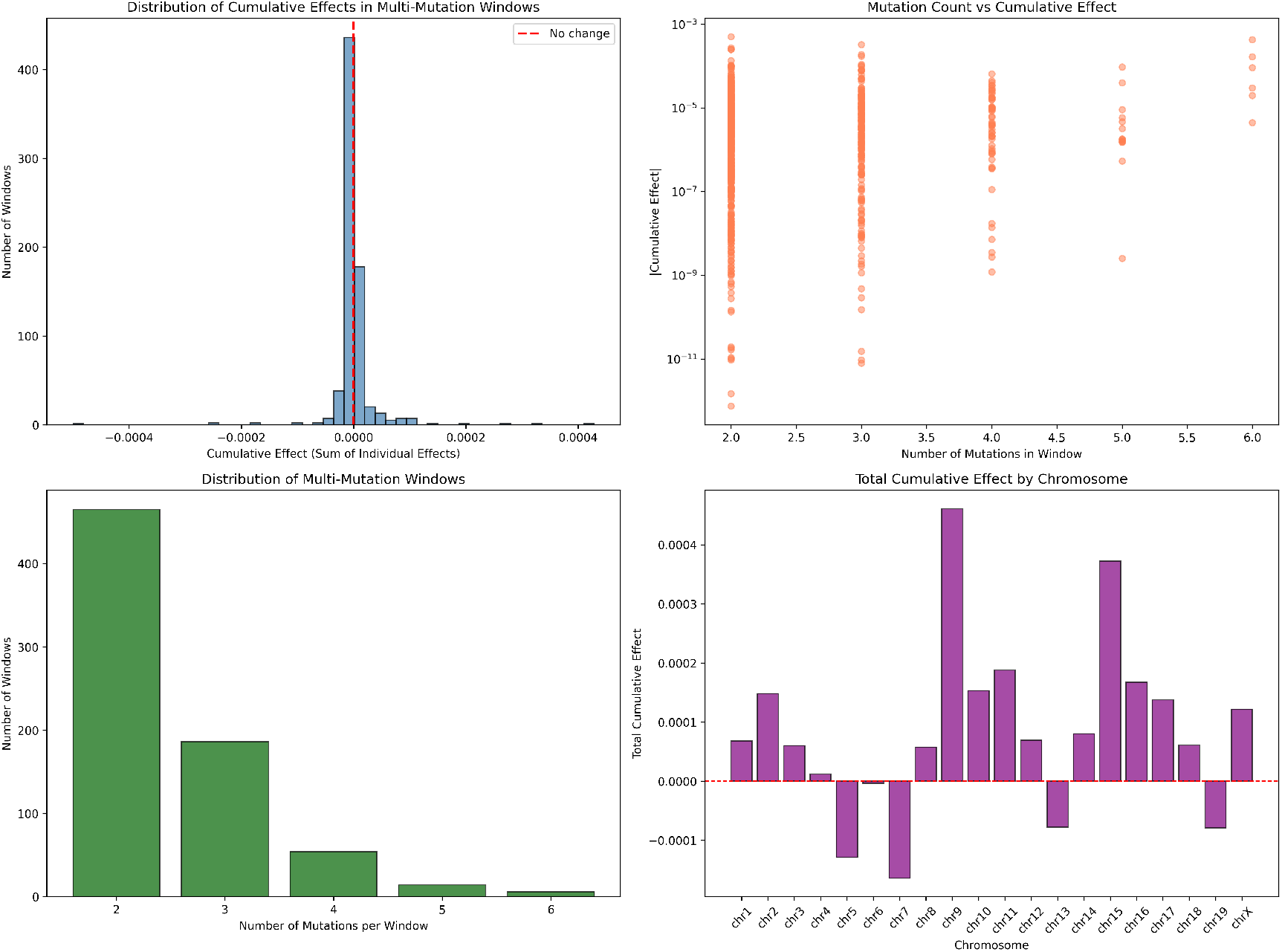
Mouse mm10 mirror of the human R2 1-Mb combined-effect analysis (main-text Figure 3). Variants binned into 1-Mb windows; cumulative window effects scale roughly linearly with variant count and remain orders of magnitude below the aging log_2_FC scale. *Species: mouse* (mm10).

**Figure S4:**
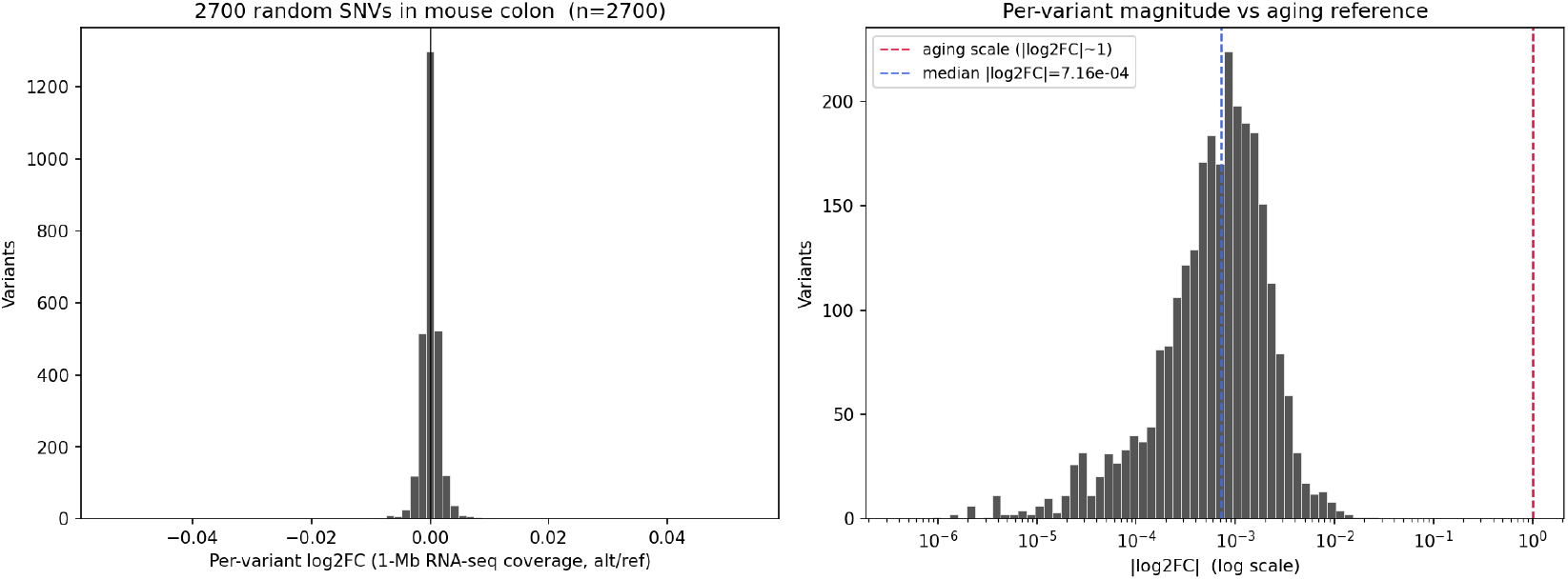
Per-variant log_2_FC distribution from the mouse mm10 random-SNV simulation. Companion histogram to Supplementary Figure S2. The distribution is tightly centred near zero, mirroring the human hg38 result. *Species: mouse* (mm10).

**Figure S5:**
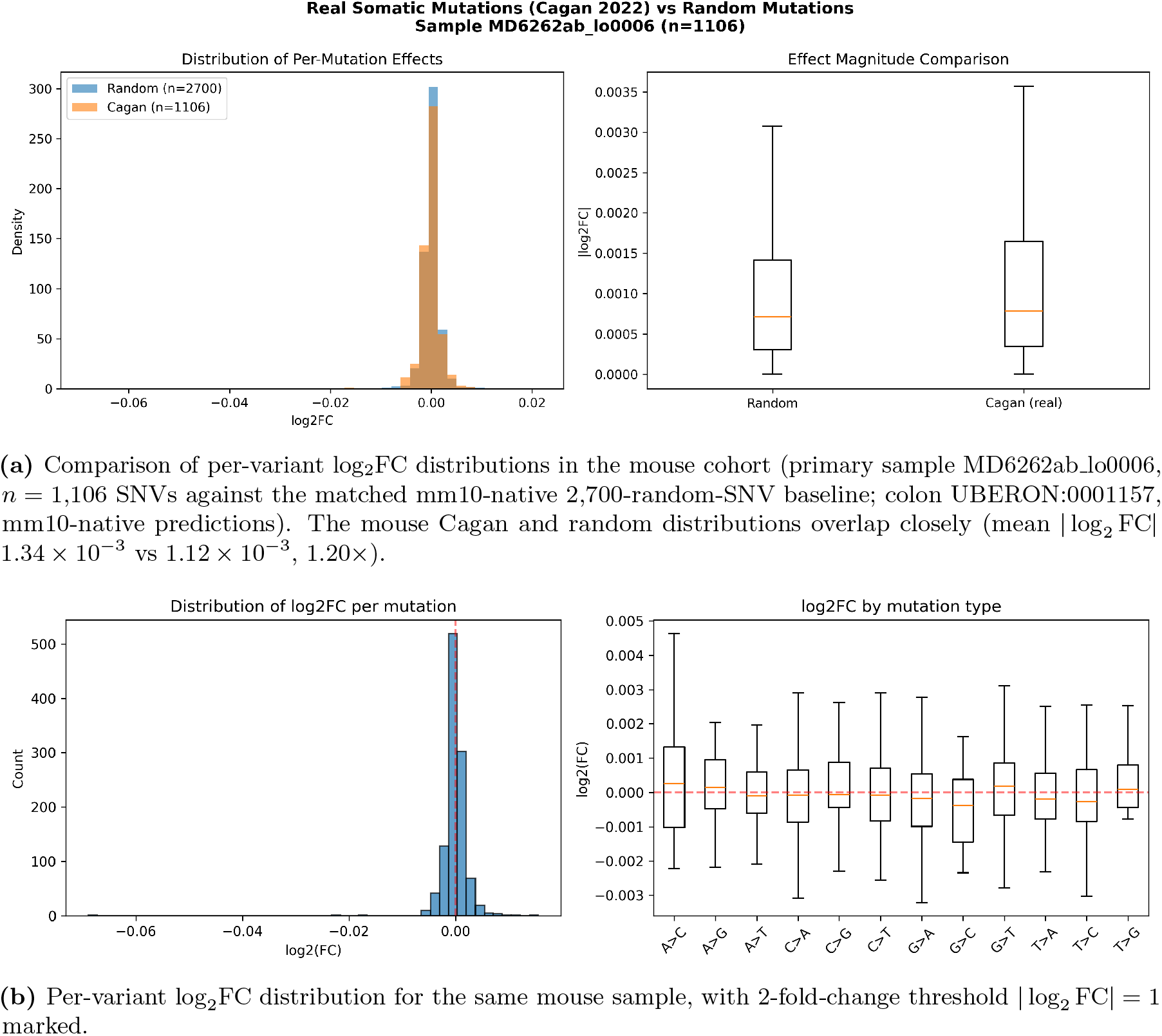
Mouse-cohort equivalent of Figure 7. *Species: mouse*. Retained because the mouse cohort is larger (*n* = 54 crypts; 9,177 SNVs across five detailed samples, scored mm10-natively) than the three-donor human cohort shown in the main-text figure; conclusions are identical in both species.

**Figure S6:**
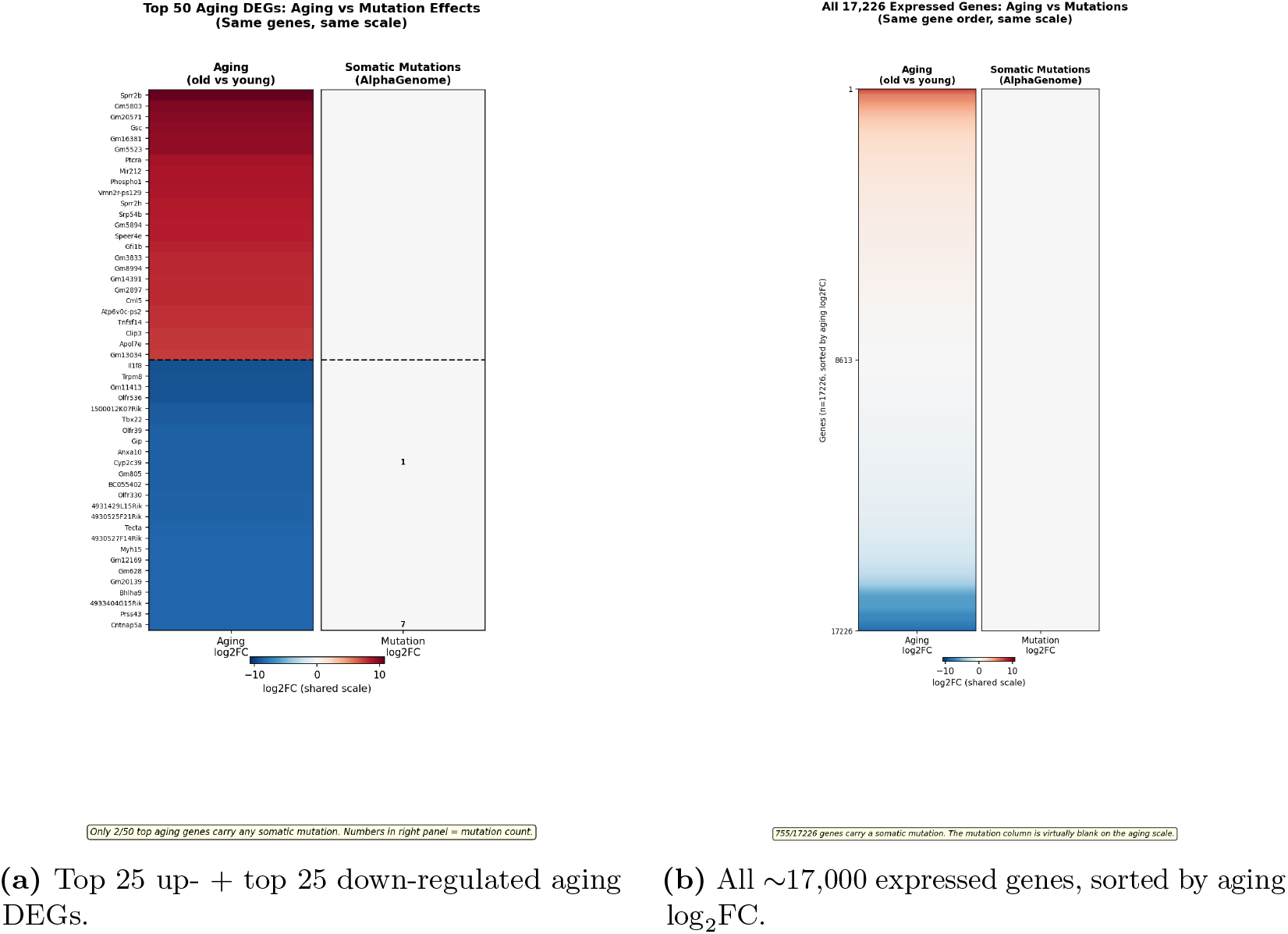
Mouse-on-mouse Cagan-vs-aging heatmap (R4 supplement). 1,781 real somatic mutations from Cagan sample MD6267ab lo0015 plotted side-by-side with Tabula Muris Senis aging log_2_FC on a shared diverging colour scale, in mouse large-intestine epithelium. (*Left*) Top 25 up + 25 down aging DEGs; the aging column paints the full [−10, +12] gradient while the mutation column is entirely neutral. (*Right*) All ~ 17k expressed genes ordered by aging log_2_FC; mutation column blank along the entire axis despite 755 of those genes carrying ≥1 real somatic mutation. Max aging-to-mutation ratio ≈ 840×, median ≈ 990× at 1-Mb-window scoring. *Species: mouse* (Tabula Muris Senis; Cagan *et al*. 2022 mouse sample MD6267ab lo0015).

**Figure S7:**
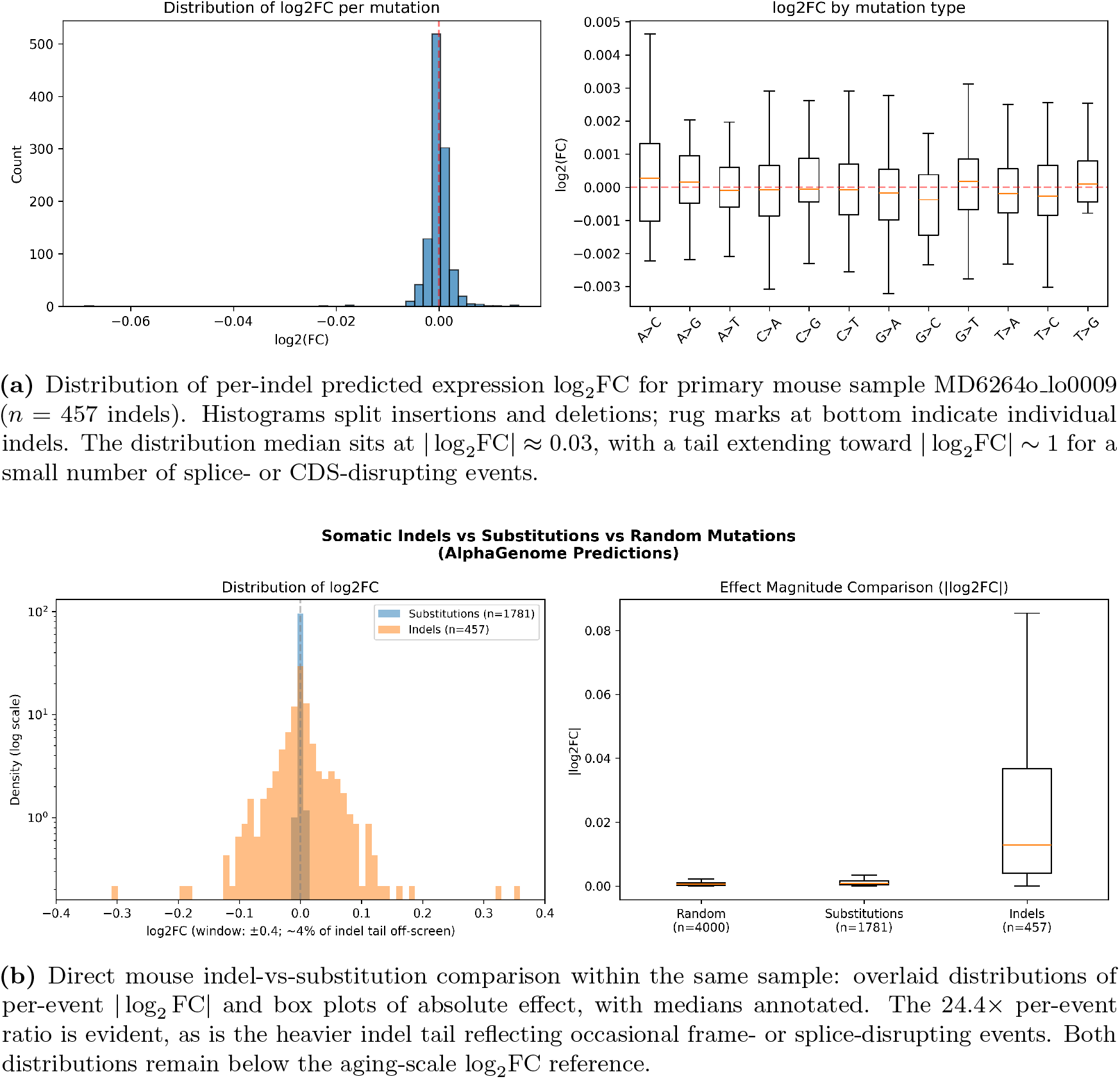
Mouse-cohort counterpart of main-text Figure 9 (*n* = 8 mouse samples, 3,233 indels in total; primary detail sample MD6264o lo0009, *n* = 457). The mouse cohort confirms the per-event indel/SNV amplification observed in the human cohort: mean |log_2_ FC| across 409 indels of sample MD6267ab_lo0003 was 0.030 vs 0.0012 for substitutions in the same sample (24.4× larger per mutation). The per-gene worst case observed across all 3,233 mouse indels in all 8 samples was |∑log_2_ FC| = 0.82 at *Spertl* (4 indels in the same gene), still below the magnitude of a single top aging DEG. *Species: mouse* (Cagan *et al*. 2022 mouse colonic crypts).

**Figure S8:**
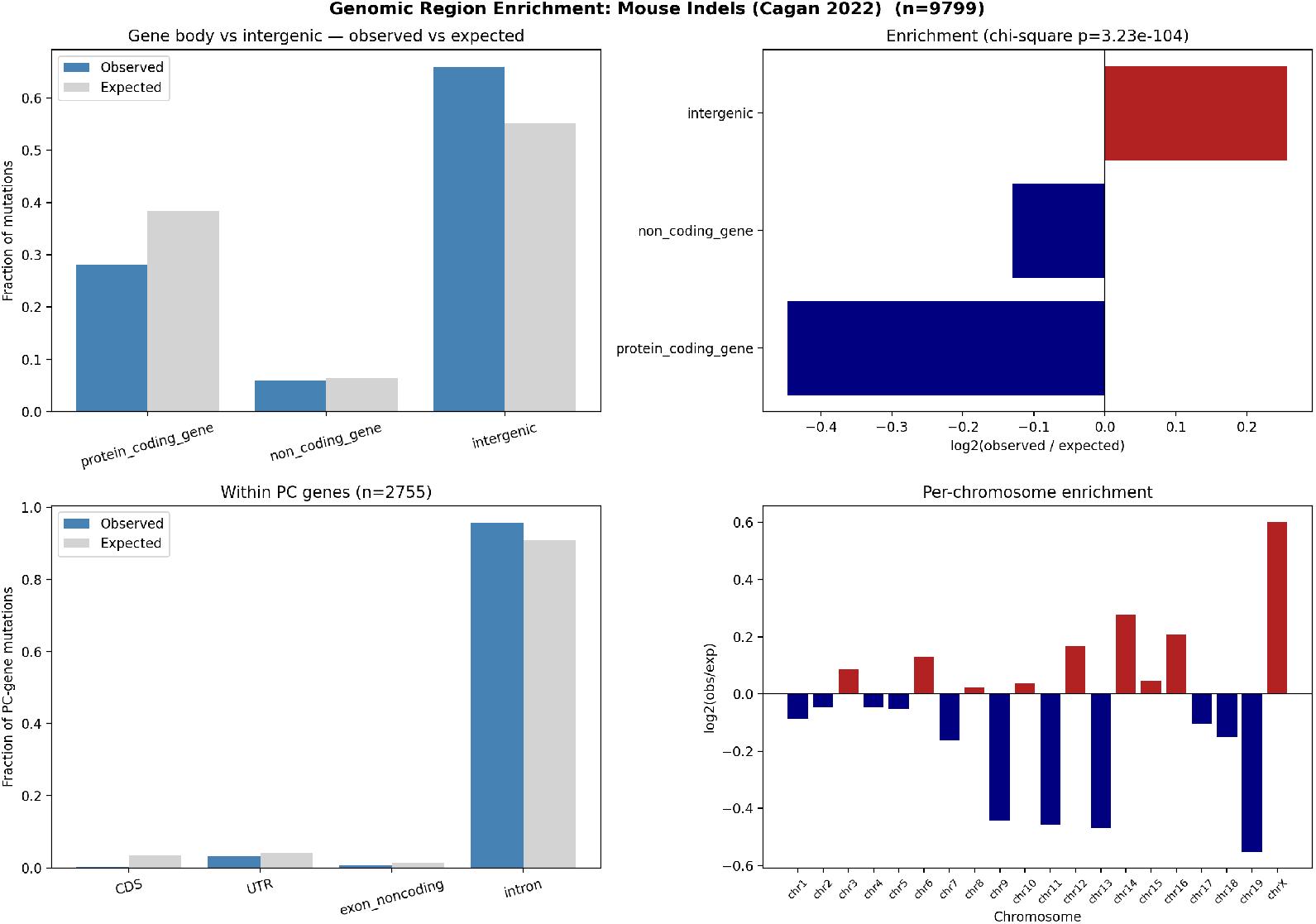
Mouse-cohort counterpart of main-text Figure 10 (*n* = 54 samples, 9,799 indels). The same purifying-selection signature seen in the human cohort, with much sharper magnitudes owing to the larger sample size: 12× CDS depletion (log_2_ = −3.58, only 8 of 2,755 protein-coding-gene indels in CDS), modest intergenic enrichment (log_2_ = +0.26), and a top-level *χ*^2^ goodness-of-fit p-value of 3.2 × 10^−104^ that is power-driven by the 9,799-indel sample size (the same fractional deviations against ~3× fewer indels in human gave *p* = 4.4 × 10^−2^ at the top level). *Species: mouse* (Cagan *et al*. 2022, 54 mouse colonic crypts; GENCODE vM25 mm10 annotations).

**Figure S9:**
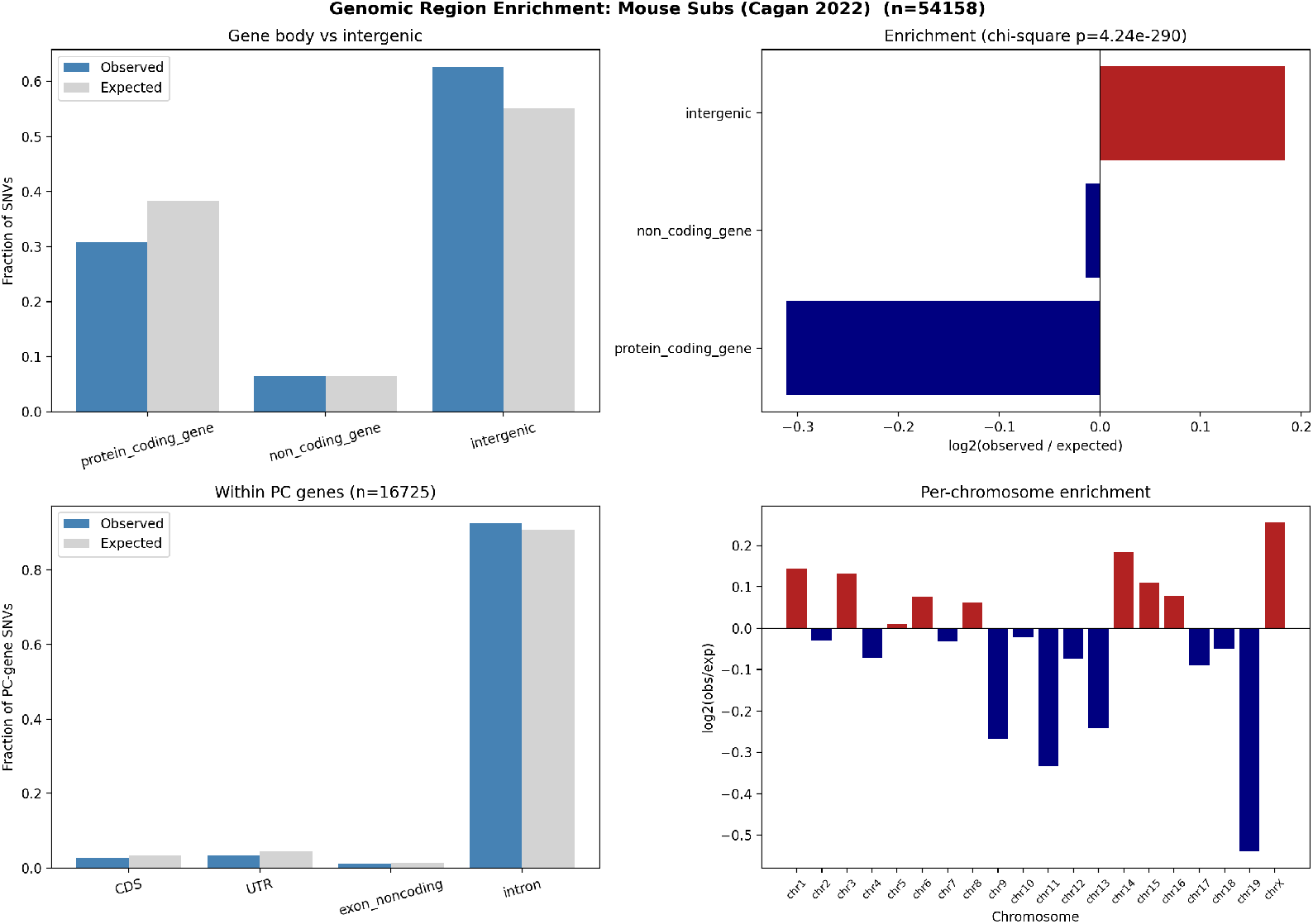
Mouse-cohort counterpart of main-text Figure 11 (*n* = 54 samples, 54,158 SNVs). Region-class enrichment of mouse SNVs reproduces the human direction at slightly weaker per-event magnitude: CDS ~ 1.27× depleted (log_2_ = −0.34, 460 of expected 583 hits), UTRs and exon-noncoding regions modestly depleted (log_2_ ≈ −0.32 to −0.36), introns near-neutral, intergenic modestly enriched (log_2_ = +0.18). Within-PC *χ*^2^ *p* = 1.0 × 10^−15^; top-level *χ*^2^ *p* = 4.2 × 10^−290^ (the latter dominated by the 54,158-SNV sample size; the biologically meaningful quantity is the modest log_2_ effect size, which is roughly an order of magnitude smaller than the indel result of Supplementary Figure S8). *Species: mouse* (Cagan *et al*. 2022, 54 mouse colonic crypts; GENCODE vM25 mm10 annotations).

## References

[1] Cagan, A. et al. Somatic mutation rates scale with lifespan across mammals. Nature 604, 517–524 (2022).

[2] Avsec, Ž. et al. AlphaGenome: advancing variant effect prediction with a unified DNA sequence model. Google DeepMind technical report (2025).

[3] The Tabula Muris Consortium. A single-cell transcriptomic atlas characterizes ageing tissues in the mouse. Nature 583, 590–595 (2020).

[4] The Tabula Sapiens Consortium. The Tabula Sapiens: a multiple-organ, single-cell transcriptomic atlas of humans. Science 376, eabl4896 (2022).

[5] The Tabula Sapiens Consortium. Tabula Sapiens v2: broadening the human single-cell transcriptomic atlas. bioRxiv 2024.12.03.626516 (2024).

[6] Frankish, A. et al. GENCODE 2021. Nucleic Acids Research 49, D916–D923 (2021).

[7] López-Otín, C., Blasco, M. A., Partridge, L., Serrano, M. & Kroemer, G. The hallmarks of aging. Cell 153, 1194–1217 (2013).

[8] López-Otín, C., Blasco, M. A., Partridge, L., Serrano, M. & Kroemer, G. Hallmarks of aging: an expanding universe. Cell 186, 243–278 (2023).

[9] Kirkwood, T. B. L. Understanding the odd science of aging. Cell 120, 437–447 (2005).

[10] Szilard, L. On the nature of the aging process. Proceedings of the National Academy of Sciences 45, 30–45 (1959).

[11] Vijg, J. From DNA damage to mutations: all roads lead to aging. Ageing Research Reviews 68, 101316 (2021).

[12] Schumacher, B., Pothof, J., Vijg, J. & Hoeijmakers, J. H. J. The central role of DNA damage in the ageing process. Nature 592, 695–703 (2021).

[13] Moore, L. et al. The mutational landscape of human somatic and germline cells. Nature 597, 381–386 (2021).

[14] Martincorena, I. & Campbell, P. J. Somatic mutation in cancer and normal cells. Science 349, 1483–1489 (2015).

[15] Glover, T. W., Wilson, T. E. & Arlt, M. F. Fragile sites in cancer: more than meets the eye. Nature Reviews Cancer 17, 489–501 (2017).

[16] Le Tallec, B. et al. Common fragile site profiling in epithelial and erythroid cells reveals conservation across tissues and correlation with replication stress. Cell Reports 4, 420–428 (2013).

[17] Tomasetti, C. & Vogelstein, B. Variation in cancer risk among tissues can be explained by the number of stem cell divisions. Science 347, 78–81 (2015).

[18] Campisi, J. et al. From discoveries in ageing research to therapeutics for healthy ageing. Nature 571, 183–192 (2019).

[19] Horvath, S. DNA methylation age of human tissues and cell types. Genome Biology 14, R115 (2013).

[20] Yang, J.-H. et al. Loss of epigenetic information as a cause of mammalian aging. Cell 186, 305–326 (2023).

[21] Kent, W. J. et al. The human genome browser at UCSC. Genome Research 12, 996–1006 (2002).

[22] Benjamini, Y. & Hochberg, Y. Controlling the false discovery rate: a practical and powerful approach to multiple testing. Journal of the Royal Statistical Society B 57, 289–300 (1995).

[23] Virtanen, P. et al. SciPy 1.0: fundamental algorithms for scientific computing in Python. Nature Methods 17, 261–272 (2020).

[24] Harris, C. R. et al. Array programming with NumPy. Nature 585, 357–362 (2020).

[25] Seabold, S. & Perktold, J. Statsmodels: econometric and statistical modeling with Python. Proc. 9th Python in Science Conf. 92–96 (2010).

