## Supplementary material for "Testing the mutation accumulation hypothesis in aging with AlphaGenome": Manuscript LaTex file: paper.pdf

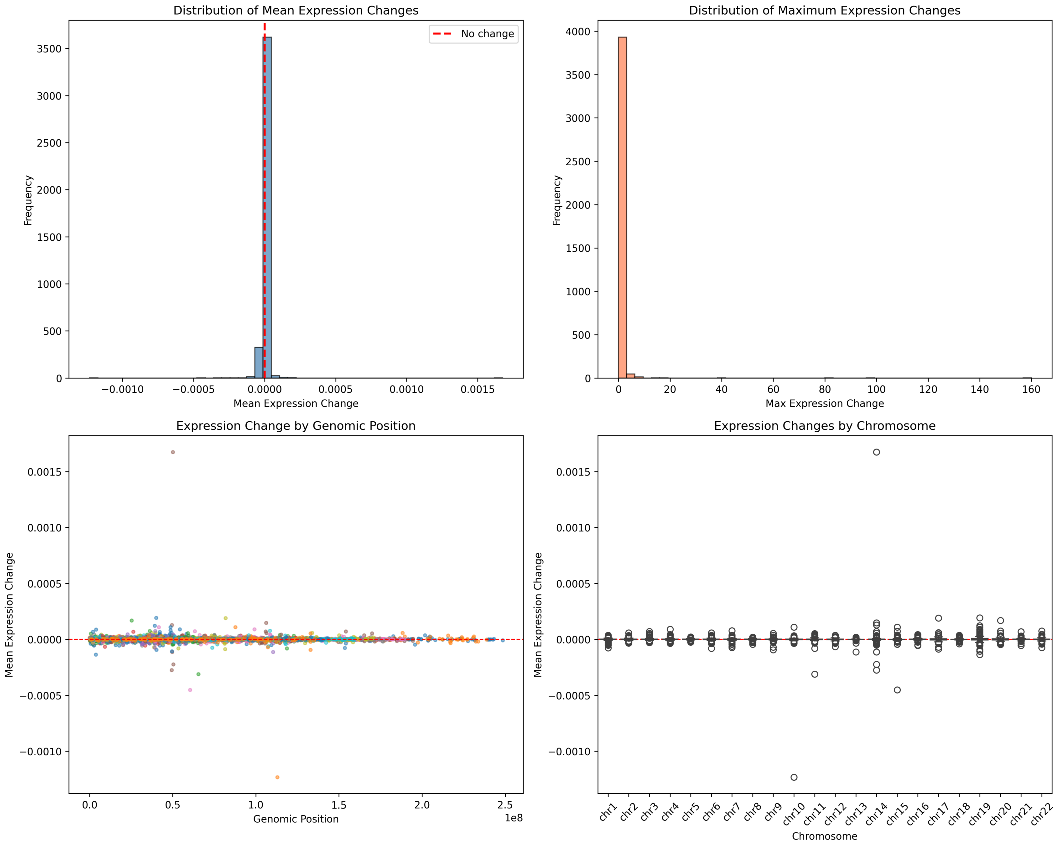


**Figure 1: Baseline distribution of AlphaGenome-predicted effects for 4,000 random genome-wide single-nucleotide variants in colon tissue.** Each of the four panels summarizes one facet of the per-variant effect distribution. (*Top left*) Histogram of mean expression change across the 1 Mb output window, centered tightly on zero with the bulk of mass within $|x| < 10^{-4}$; no symmetric tail, consistent with most random substitutions falling in unconstrained intergenic or intronic sequence. (*Top right*) Distribution of per-variant max expression change, spanning four orders of magnitude from $10^{-4}$ to $\sim 4$; outliers represent rare variants that intersect active regulatory elements or splice boundaries. (*Bottom left*) Distribution of total expression change (sum of $|\log_2 \mathrm{FC}|$ across affected genes in the window), dominated by small effects with a single extreme observation (chr17:81,415,658 T→A, total = 1,104) that would otherwise dominate a linear axis and motivates the log-scale display. (*Bottom right*) Scatter of max vs. mean expression change per variant, confirming that the rare high-max outliers correspond to localized effects that do not lift the window-average appreciably — evidence that even strong local perturbations produce a bounded, tightly-buffered footprint at the gene level. Colon tissue (UBERON:0001157); RNA-seq requested; 100% prediction success. *Species: human* (4,000 SNVs sampled from hg38, scored in human colon).

**Table 1: Mean $|\Sigma \log_2 \mathrm{FC}|$ across each Cagan / random-SNV mutation experiment computed over *all* 3,579 expressed colonic-epithelial genes (TSP v2), filling 0 for genes not assigned a mutation.** For each Cagan dataset the per-donor mean is computed and then averaged across donors; the aging row shows the mean $|\log_2 \mathrm{FC}|$ over the same expressed-gene set. *Species: human.*

| Dataset | hit-fraction (per donor) | mean $\lvert\log_2 \mathrm{FC}\rvert$ | max $\lvert\log_2 \mathrm{FC}\rvert$ | aging-mean ratio | aging-max ratio |
|---|---|---|---|---|---|
| Aging (TSP v2) | — | **0.795** | **6.98** | 1× | 1× |
| Random SNVs (hg38) | 16.9% | $1.6 \times 10^{-4}$ | 0.052 | 5,065× | 134× |
| Cagan SNVs (3 donors) | 13.0% | $1.8 \times 10^{-4}$ | 0.527 | 4,463× | 13× |
| Cagan indels (9 donors) | 1.1% | $2.1 \times 10^{-4}$ | 0.173 | 3,790× | 40× |

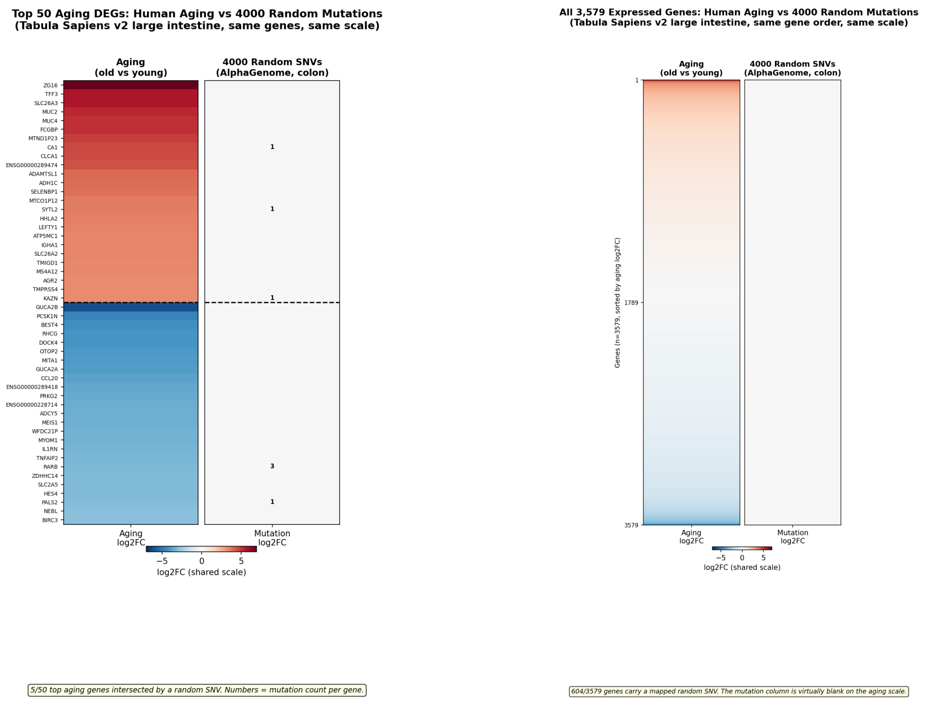


**(a)** Top 25 up- + top 25 down-regulated aging DEGs.

**(b)** All 3,579 expressed genes, sorted by aging $\log_2$FC.

**Figure 2: 4,000 random SNVs produce per-gene effects that are visually negligible on the aging $\log_2$FC scale in human large-intestine epithelium.** *Species: human* (Tabula Sapiens v2 aging reference; AlphaGenome random-SNV simulation in human colon, UBERON:0001157). Both panels use the same diverging colour scale centred at zero. (*Left, a*) Top 25 up- and top 25 down-regulated aging DEGs from Tabula Sapiens v2 large-intestine epithelium [4, 5] (young donor TSP26, 37 y; old donors TSP2/TSP14/TSP25/TSP27, 56–61 y). The aging column spans a strong red-to-blue gradient, whereas the mutation column remains essentially neutral after summing per-gene AlphaGenome-predicted $\log_2$FC across the 4,000 random SNVs mapped to the nearest hg38 protein-coding gene. (*Right, b*) All 3,579 expressed epithelial genes ordered by aging $\log_2$FC; 604 genes carry a mapped random SNV, but their summed mutation effects remain invisible at the aging scale. Small numerals mark the mutation count per gene. The per-gene mutation signal is therefore orders of magnitude below the observed aging program.

#### R2. Combined effects in the same 1-Mb window do not amplify

Individual mutations could in principle sum to non-trivial effects if they co-localize within a common cis-regulatory window. I tested this by grouping the 4,000 random variants into non-overlapping 1-Mb genomic windows and measuring the *expected additive* cumulative effect (sum of per-variant mean expression changes) for windows harboring multiple variants. Windows with 10 co-occurring random SNVs reached a cumulative expected effect of only $\sim 3 \times 10^{-4}$, barely distinguishable from the single-variant scale and orders of magnitude below aging-scale $\log_2$FC. The cumulative-effect distribution remained approximately symmetric around zero with no heavy right tail, showing that the upper-bound assumption of perfect additivity still yields negligible totals at realistic variant densities (Figure 3). This rules out a simple "hotspot accumulation" mechanism whereby local co-occurrence amplifies individually weak mutations.

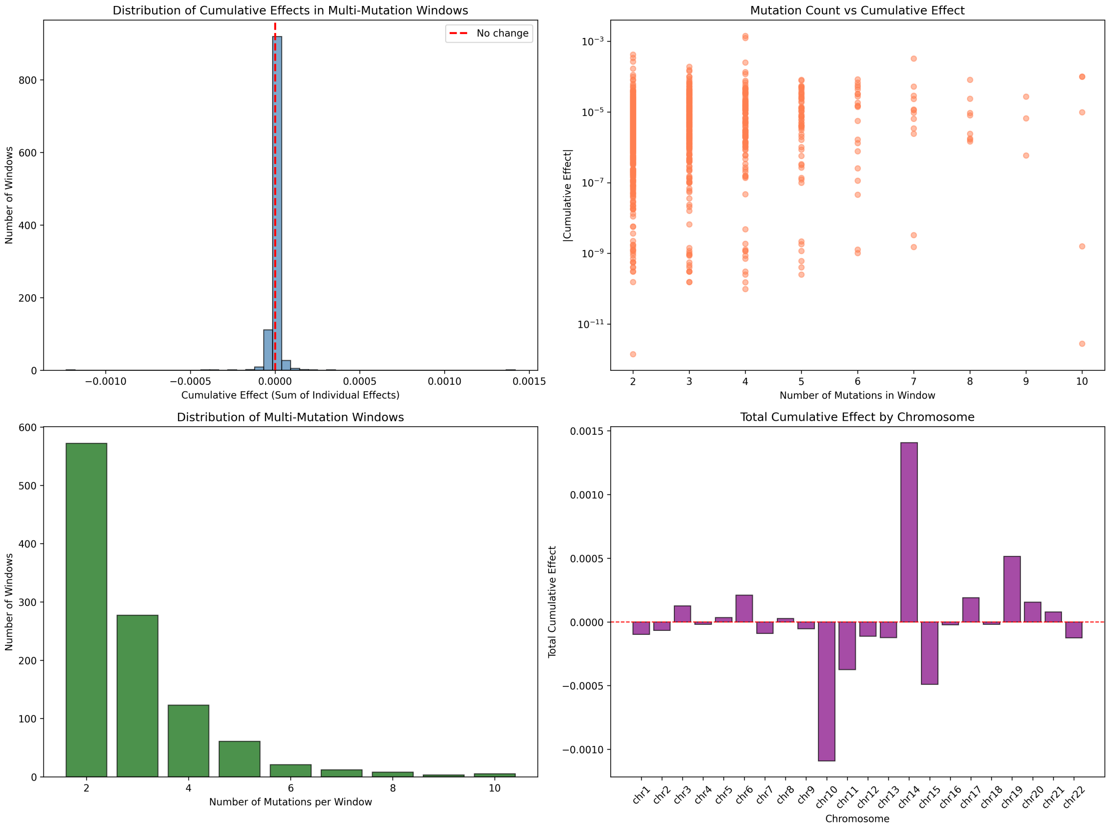


**Figure 3: Combined-effect analysis of random SNVs grouped into 1-Mb windows.** Random variants were binned by their genomic coordinate into $\sim$ 3,000 non-overlapping 1-Mb windows. Panels summarize, for each window: (1) how many variants it contains (most windows 0–2), (2) the absolute mean expression change of the window's largest single variant, and (3) the cumulative window effect computed as the sum of per-variant mean changes (additive-expectation upper bound). The cumulative distribution mirrors the single-variant scale, with even windows containing 10 co-occurring variants reaching only $\sim 3 \times 10^{-4}$ cumulative effect. No window shows synergistic amplification. This result is critical because it forecloses the simplest MA-compatible escape route: "mutations at random, but stack at shared cis-regulatory bundles." At realistic variant density, they don't stack to functionally consequential levels. *Species: human* (random SNVs in human colon, UBERON:0001157).

#### R3a. Multi-cell mutation simulation and gene-level scoring reveal occasional aging-magnitude effects from single random SNVs, but multi-hit additive amplification is empirically bounded

A vertebrate tissue consists usually of millions to billion of cells. I therefore ran the random-SNV simulation at gene-body resolution: 60 independent simulated cells $\times$ ~4,000 random hg38 SNVs each (240,000 AlphaGenome calls; 203,405 with finite gene-body $\log_2$FC), each variant scored as $\log_2((s_{\text{alt}}+1)/(s_{\text{ref}}+1))$ over the nearest GENCODE v46 protein-coding gene body in human colon (UBERON:0001157) (Figure 4). The per-variant gene-body distribution shows a standard deviation of $1.43 \times 10^{-2}$ at window level and a maximum of $|\log_2 \text{FC}| = 2.248$ with the gene *TMIGD2* (Figure 5). Ten of 203,405 variants reach $|\log_2 \text{FC}| \geq 1$ (~1 in 20,000; the top hits are *TMIGD2* +2.248, *RNASE2* −1.546, *NEUROD4* −1.520, *CHRM1* +1.423, *GABRA1* +1.412). The median $|\log_2 \text{FC}|$ remains small at $1.04 \times 10^{-3}$. Per-(cell, gene) aggregation across co-localising SNVs produces a maximum cumulative $|\sum \log_2 \text{FC}|$ of *exactly* 2.248 — equal to the single-SNV maximum — ruling out the multi-hit additive failure mode at the level of an individual gene: when two random SNVs land in the same gene in the same cell, they do not stack to a larger effect than the largest single hit. This whole-cell aggregate is summed across thousands of distinct genes; the biologically meaningful per-gene comparison against aging differential expression remains bounded $\sim 3\times$ below the TSP aging maximum (2.248 vs 6.98). A more precise formulation of the central argument is therefore: occasional single random SNVs ($\sim$ 1 in 20,000) reach aging-magnitude $|\log_2 \mathrm{FC}| \sim 1$ in their nearest gene; multi-hit additive stacking within one gene is empirically bounded; and the bulk per-variant distribution sits three orders of magnitude below the aging median.

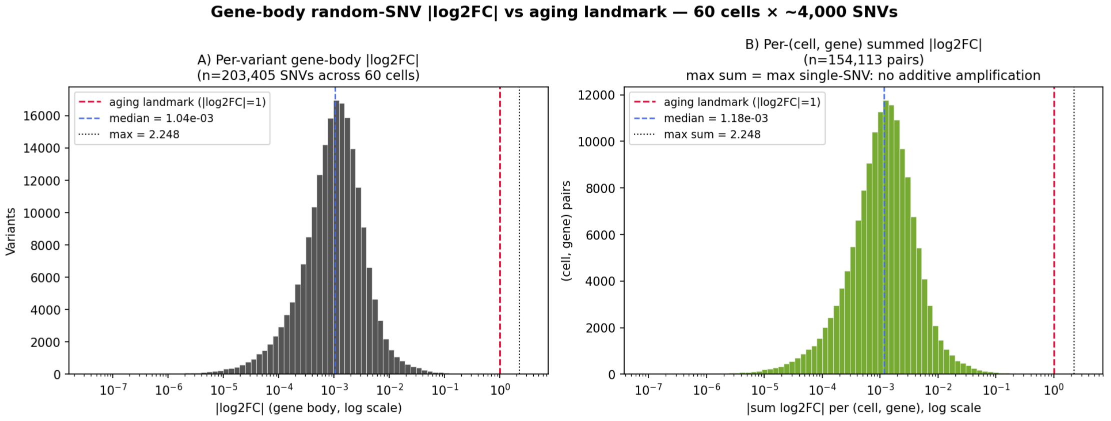


**Figure 4: Gene-body random-SNV $|\log_2 \mathbf{FC}|$ distribution against the 2-fold-change threshold $|\log_2 \mathbf{FC}| = 1$.** The reference line at $|\log_2 \mathrm{FC}| = 1$ marks a 2-fold expression change — a conventional biological-significance threshold cleared by thousands of aging DEGs in the TSP large-intestine reference (range 0–6.98, median 0.572); see Methods. (*Left*) Per-variant $|\log_2 \mathrm{FC}|$ histogram on log axis across 203,405 random SNVs from 60 simulated cells; vertical lines mark the 2-fold threshold, the median, and the empirical maximum (*TMIGD2*, 2.248). Ten SNVs reach $\geq 1$ ($\sim$ 1 in 20,000); 157 reach $\geq 0.138$. (*Right*) Per-(cell, gene) summed $|\log_2 \mathrm{FC}|$ across co-localising SNVs in the same cell; the maximum sum equals the single-SNV maximum, ruling out additive multi-hit amplification at the per-gene level. *Species: human* (hg38 SNVs scored in human colon, UBERON:0001157).

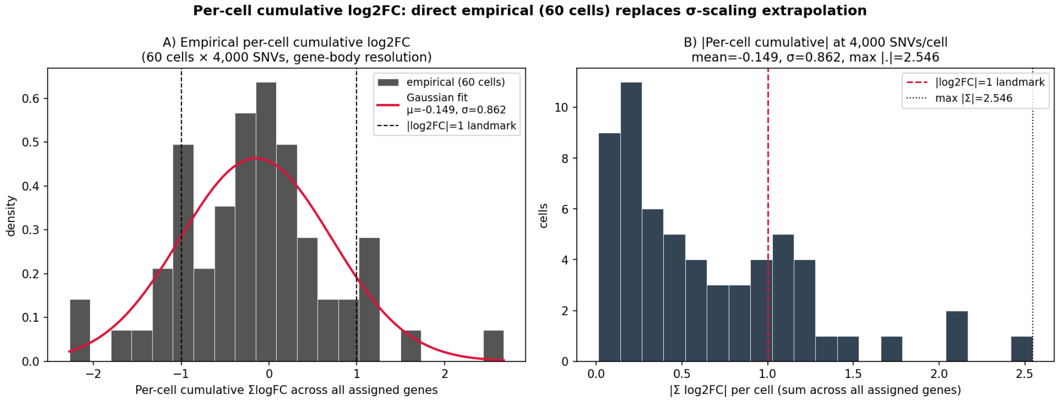


**Figure 5: Per-cell cumulative $\log_2$FC distribution at 4,000 SNVs/cell — direct empirical measurement (60 cells)** (*Left*) Empirical histogram of per-cell $\sum \log_2 \mathrm{FC}$ summed across all assigned genes (60 cells $\times \sim$ 4,000 SNVs each), with fitted Gaussian (mean $-0.149$, $\sigma = 0.862$). (*Right*) $|\sum \log_2 \mathrm{FC}|$ per cell against the 2-fold-change threshold $|\log_2 \mathrm{FC}| = 1$; max $|\sum| = 2.55$. This whole-cell aggregate is summed across thousands of distinct genes and is therefore not directly comparable to per-gene aging differential expression — the per-gene comparison (left panel of Figure 4) is the load-bearing one and remains bounded $\sim 3\times$ below the TSP aging maximum. *Species: human.*

#### R3b. Pseudobulk RNA-seq aggregation across the 60-cell tissue tightens the per-gene bound to $\sim 36\times$ below aging max and shows zero correlation with the aging program

The per-variant view of R3a is sensitive to rare extreme single-SNV outliers. The biologically meaningful question for an aged tissue is what happens when a population of cells, each carrying its own random ∼4,000-SNV burden, is aggregated into a bulk transcriptome — the situation a bulk RNA-seq experiment would actually measure. I therefore pooled the 60 simulated cells into a pseudobulk and computed, for each protein-coding gene $g$, the pseudobulk fold change

$$\log_2 \mathrm{FC}_g^{\mathrm{pb}} \;=\; \log_2\left(\frac{\sum_{(c,i)\to g} s_{\mathrm{alt}}^{(c,i)} + 1}{\sum_{(c,i)\to g} s_{\mathrm{ref}}^{(c,i)} + 1}\right),$$

where the sum runs over every (cell $c$, variant $i$) pair whose nearest GENCODE v46 gene is $g$, and $s_{\mathrm{ref}}^{(c,i)}$ / $s_{\mathrm{alt}}^{(c,i)}$ are the AlphaGenome RNA-seq coverage sums under the reference and alternate allele in the 1-Mb window centred on that variant. This is the bulk-RNA-seq analogue of comparing a 60-cell mutated condition against an unmutated reference. Of 3,579 expressed colonic-epithelial genes in Tabula Sapiens v2 (R1), 3,056 (85.4%) are intersected by at least one random SNV across the 60 cells and therefore receive a non-trivial pseudobulk estimate. The per-gene pseudobulk distribution is $\sim 10\times$ tighter than the per-variant distribution: maximum $|\log_2 \mathrm{FC}^{\mathrm{pb}}| = 0.192$ (*TMEM238*, 4 hits), median $3.96 \times 10^{-4}$ over hit genes (Figure 6A-B). Against TSP-aging max 6.98 and median 0.572, the max-effect ratio is therefore $\sim 36\times$ (substantially tighter than the $\sim 3\times$ bound from the per-variant max) and the median ratio is $\sim 1{,}445\times$. The tightening reflects the standard bulk-vs-single-cell averaging effect: rare single-SNV outliers dilute when pooled with many other variants hitting the same gene from independent cells. Beyond magnitude, I tested whether the random-SNV pseudobulk *correlates* with aging differential expression — the directional version of the MA hypothesis — and found no relationship: Pearson $r = -0.027$ ($p = 0.13$), Spearman $\rho = -0.031$ ($p = 0.09$) over the 3,056 hit genes (Figure 6C). Even where random SNVs do perturb gene expression in this aggregated tissue model, the perturbations are uncorrelated with the aging fold-change pattern.

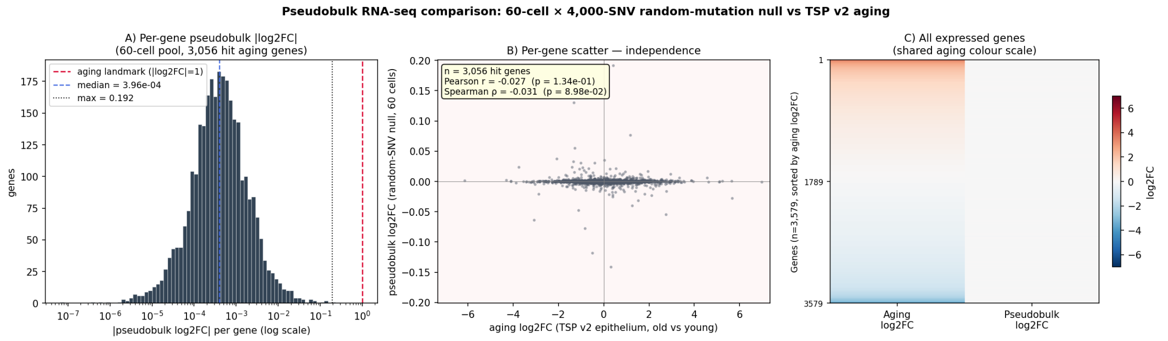


**Figure 6: Pseudobulk RNA-seq comparison: 60-cell × 4,000-SNV random-mutation baseline versus Tabula Sapiens v2 aging in colonic epithelium.** (*A*) Per-gene pseudobulk $|\log_2 \mathrm{FC}^{\mathrm{pb}}|$ histogram across 3,056 hit aging genes, on log axis, against the 2-fold-change threshold $|\log_2 \mathrm{FC}| = 1$. Median over hit genes $= 3.96 \times 10^{-4}$, max $= 0.192$ (*TMEM238*). (*B*) Per-gene scatter of aging $\log_2$FC versus pseudobulk $\log_2$FC over the same 3,056 genes. Pearson $r = -0.027$ ($p = 0.13$); Spearman $\rho = -0.031$ ($p = 0.09$). The pseudobulk axis spans approximately $[-0.2, +0.2]$ while the aging axis spans approximately $[-7, +7]$ — the two are independent both in magnitude and direction. (*C*) Side-by-side per-gene heatmap of all 3,579 expressed genes sorted by aging $\log_2$FC, with the pseudobulk column on the same diverging colour scale as the aging column; the pseudobulk column is uniformly neutral, mirroring the random-SNV-vs-aging picture of Figure 2 but at the considerably more demanding pseudobulk granularity (a per-gene aggregate over multiple cells, not a single-variant assignment). *Species: human* (hg38 SNVs scored in human colon, UBERON:0001157; TSP v2 large-intestine aging reference).

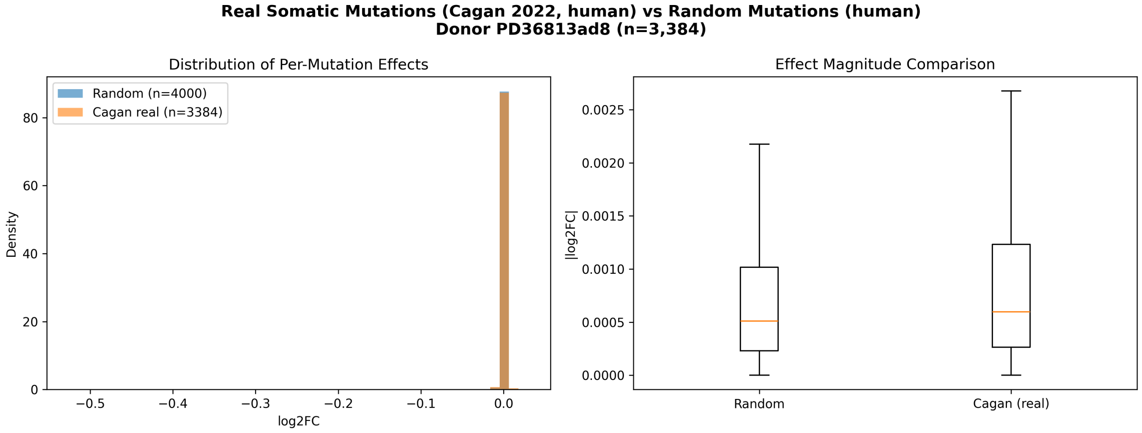


**(a)** Two-panel comparison of per-variant effect distributions (human donor PD36813ad8, $n = 3{,}384$ SNVs vs a matched random baseline). *Left*: histograms of $\log_2$FC; Cagan and random distributions are nearly coincident. *Right*: $|\log_2 \mathrm{FC}|$ comparison; he Cagan box sits at or below the random box.

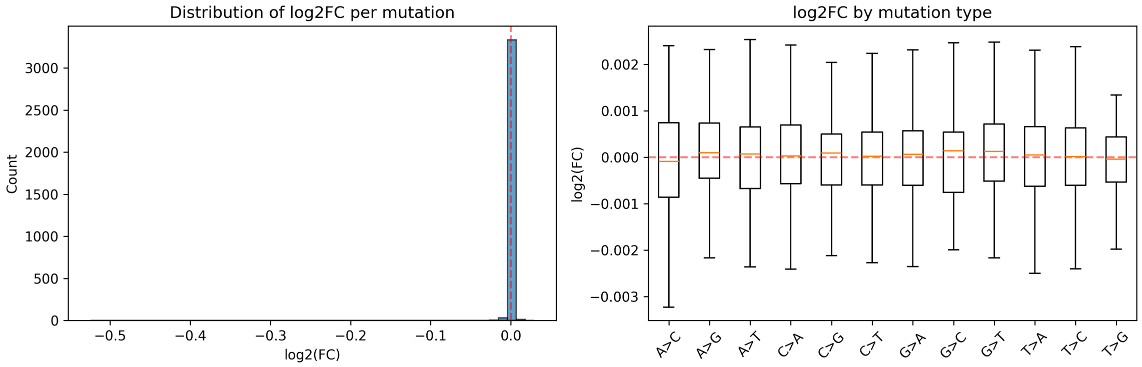


**(b)** Per-variant $\log_2$FC distribution (same human sample). Median $|\log_2 \mathrm{FC}| = 5.9 \times 10^{-4}$, consistent with the random baseline and three orders of magnitude below the 2-fold-change threshold $|\log_2 \mathrm{FC}| = 1$.

**Figure 7: Real somatic SNVs from aged human colonic crypts (Cagan 2022, $n = 3$ donors) produce per-variant AlphaGenome effects smaller than random genome-wide SNVs.** All panels computed in colon tissue (UBERON:0001157). *Species: human.* See Supplementary Figure S5 for the matched mouse-cohort figure ($n = 54$ crypts, used here as the larger-sample-size confirmation); conclusions are identical.

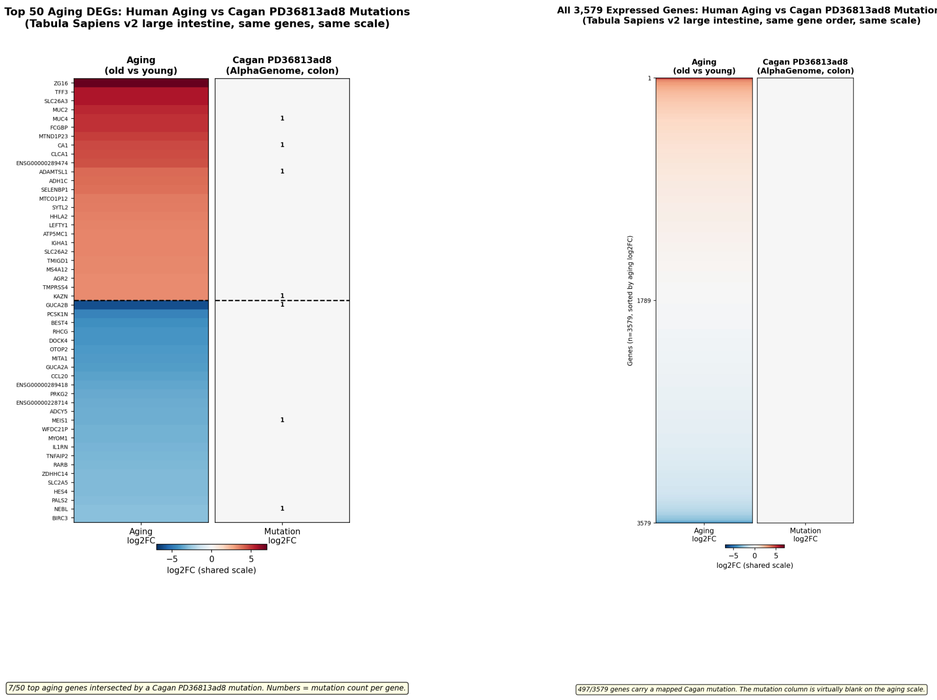


**(a)** Top 25 up- + top 25 down-regulated aging DEGs.

**(b)** All 3,579 expressed genes, sorted by aging $\log_2$FC.

**Figure 8: Real Cagan human-cohort somatic SNVs (donor PD36813ad8) produce per-gene effects that are invisible on the aging $\log_2$FC colour scale in human large-intestine tissue.** *Species: human* (Tabula Sapiens v2 aging reference; AlphaGenome scoring of 3,384 Cagan PD36813ad8 SNVs in human colon, UBERON:0001157, assigned to the nearest GENCODE v46 protein-coding gene body). Both panels share the diverging colour map and aging-scale colour range used in Figure 2, allowing the two figures to be read side by side. (*Left, a*) The top 25 up- and 25 down-regulated aging DEGs from Tabula Sapiens v2 large-intestine epithelium (young donor TSP26, 37 y vs old donors TSP2/TSP14/TSP25/TSP27, 56–61 y); only 7 of 50 top aging genes are intersected by a Cagan SNV in this donor, and the per-gene mutation column is uniformly neutral. Small numerals mark the mutation count per gene. (*Right, b*) All 3,579 expressed colonic-epithelial genes ordered by aging $\log_2$FC, with 497 (14%) carrying a mapped Cagan mutation but no visible colour on the shared scale. Per-gene maximum mutation $|\log_2 \text{FC}| = 0.525$ and median $5.98 \times 10^{-4}$ versus aging max 6.98 and median 0.572, giving a max-effect ratio of $\sim 13\times$ and a median ratio of $\sim 960\times$ on real Cagan human SNVs — consistent with the random-SNV picture of Figure 2 and confirming that the somatic catalogue does not concentrate at aging-relevant loci.

#### R5. Somatic indels are ∼24× more impactful per event than SNVs, yet remain negligible at the gene level

Insertions and deletions (indels) are a rarer but mechanistically distinct class of somatic variation that can disrupt splice sites, reading frames, or full regulatory elements. I scored 3,233 indels across eight mouse crypt samples (409–457 indels per sample) plus 600 indels across three human Cagan donors (PD36813ac4 $n = 236$, PD37266c_lo0016 $n = 165$, PD37590b_lo0090 $n = 199$) using the AlphaGenome pipeline adjusted for variant length. Indels produced substantially larger per-event effects than SNVs in both species, but the per-event magnitudes still sit two orders of magnitude below the mean aging threshold $|\log_2 \text{FC}| = 0.8$. I pooled all 1,630 AlphaGenome-scored indels across nine human Cagan donors and use exemplar donor PD36813ac4 ($n = 236$) plus the 9-donor pooled summary as the main human view (Figure 9); the parallel mouse analysis (8 samples, 3,233 indels; primary detail MD6264o_lo0009, $n = 457$) is shown in Supplementary Figure S7. Per-event mean $|\log_2 \text{FC}|$ in PD36813ac4 was 0.0226 (median $1.17 \times 10^{-2}$, max 0.218), against the matched-donor SNV mean of $\approx 1.0 \times 10^{-3}$ — a per-event indel/SNV ratio of $\approx 22\times$ that reproduces the $\approx 24\times$ ratio seen in the matched mouse sample (mean $|\log_2 \mathrm{FC}|$ across 409 indels of sample MD6267ab_lo0003 was 0.030 vs 0.0012 for substitutions; Supplementary Figure S7). Across the other two donors (PD37266c_lo0016 $n = 165$ and PD37590b_lo0090 $n = 199$) the mean per-indel $|\log_2 \mathrm{FC}|$ was 0.0212 and 0.0243 respectively, with per-donor indel/SNV per-event ratios of 22, 21, and $24\times$ across the three donors. The cross-species amplification ratio is therefore stable (mouse $\approx 24\times$, human $\approx 22\times$), confirming that the structural-variation amplification is reproducible at the species level. Pooled across all 9 human donors ($n = 1{,}630$ indels), the per-event maximum $|\log_2 \mathrm{FC}|$ is 0.373 and the median is 0.0115 — the entire pooled distribution remains below the 2-fold-change threshold; further, per-gene additive aggregation across donors (sum of $\log_2$FC over all pooled indels in the same gene) over 1,441 unique genes hit fails to push any single gene past the threshold either (per-gene max 0.373; 0 of 1,441 genes reach $|\sum \log_2 \mathrm{FC}| \geq 1$). When aggregated at the gene level (per-gene *sum* of $\log_2$FC, since summing across unrelated genes is biologically meaningless), the maximum per-gene cumulative effect observed in any sample was $|\sum \log_2 \mathrm{FC}| = 0.82$ (*Spertl*, 4 indels in the same gene). Even this, the worst-case per-gene aggregation, falls below the magnitude of a typical aging DEG in the matched tissue (mean aging $|\log_2 \mathrm{FC}| = 0.795$ (Table 1)), and it affects one gene rather than the thousands that differentiate young from old colon. On the whole-genome scale indels show a negligible mean $|\log_2 \mathrm{FC}|$ of $1.1 \times 10^{-4}$ for humans (Table 1) and $|\log_2 \mathrm{FC}|$ of $3.3 \times 10^{-4}$ for mice (Supplementary Table S1).

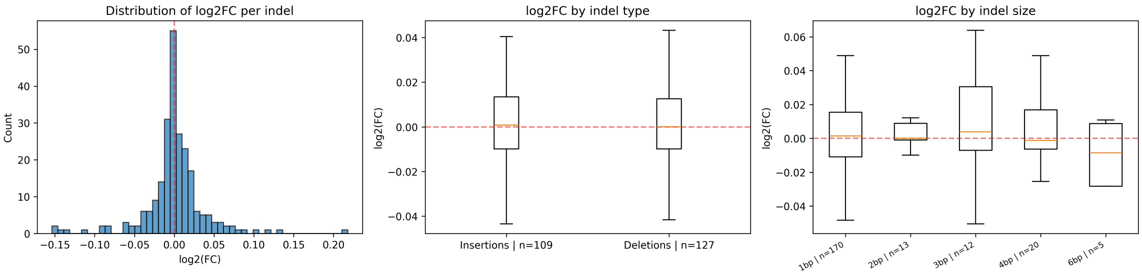


**(a)** Per-indel $\log_2$FC distribution for exemplar donor PD36813ac4 ($n = 236$): overall histogram, broken down by indel type (insertion vs deletion, near-identical) and by indel size (1–6 bp, no monotonic size–effect relationship). The bulk of the distribution sits within $\pm 0.05$.

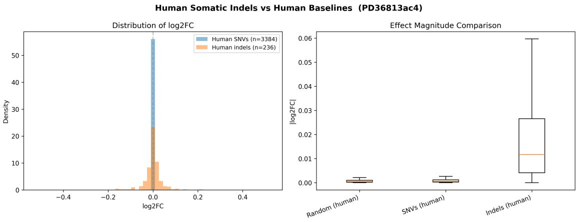


**(b)** PD36813ac4 indels vs. random hg38 SNVs and matched-donor SNV cohort: indels exceed both at every quantile but remain in the same order of magnitude, with the bulk concentrated around the median.

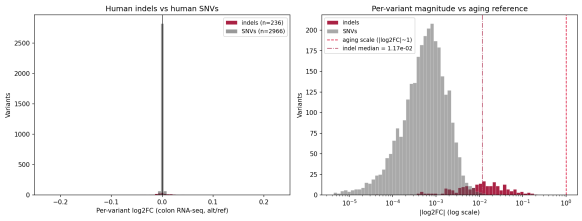


**(c)** PD36813ac4 indels vs SNVs on a shared log axis with the 2-fold-change threshold $|\log_2 \mathrm{FC}| = 1$ clearly above all observed events. Per-indel median $= 1.17 \times 10^{-2}$, two orders of magnitude below the 2-fold threshold.

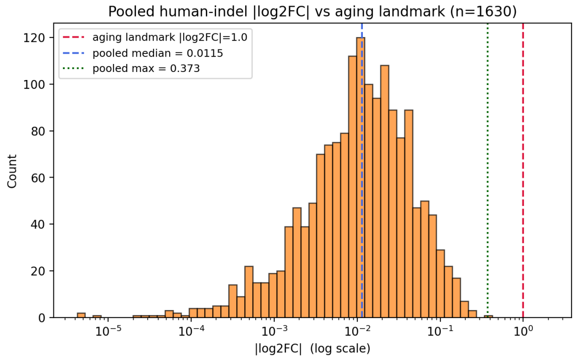


**(d)** Pooled 9-donor $|\log_2 \mathrm{FC}|$ histogram ($n = 1{,}630$ indels) on log axis: the entire pooled distribution sits below 0.4 with median 0.0115 and max 0.373 — well below the 2-fold-change threshold.

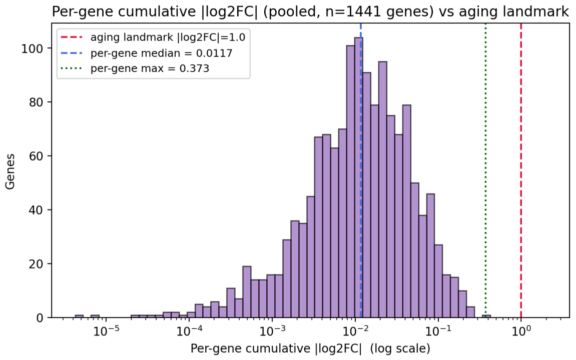


**(e)** Per-gene cumulative $|\log_2 \mathrm{FC}|$ over 1,441 unique genes hit by pooled indels (additive sum across all donors at each gene). Summing across all 9 donors does *not* push any single gene past the 2-fold-change threshold: per-gene max 0.373; 0 of 1,441 genes reach $|\sum \log_2 \mathrm{FC}| \geq 1$.

The same test applied to the much larger Cagan SNV cohorts (56,123 human SNVs across 28 samples; 54,158 mouse SNVs across 54 samples) reproduces the direction of the indel signature but with substantially smaller magnitudes per event, exactly as expected biologically: most coding-region SNVs are synonymous or conservative missense and therefore escape the strong purifying filter that frameshift indels face. Within human protein-coding genes, CDS is $\sim 1.4\times$ depleted ($\log_2 = -0.52$, 427 of expected 611 hits, within-PC $\chi^2\, p = 9.8 \times 10^{-22}$; Figure 11); the mouse cohort shows the same direction at $\sim 1.3\times$ ($\log_2 = -0.34$, 460 of expected 583, within-PC $\chi^2\, p = 1.0 \times 10^{-15}$; Supplementary Figure S9). The top-level $\chi^2$ p-values ($5.7 \times 10^{-51}$ human, $4.2 \times 10^{-290}$ mouse) reflect the very large SNV sample sizes; the biologically meaningful quantity is the modest log2 effect size, which is roughly an order of magnitude smaller than the indel result — consistent with SNVs as a class being more permissive substrates for clonal survival than length-altering indels, while still subject to coherent selection against coding disruption.

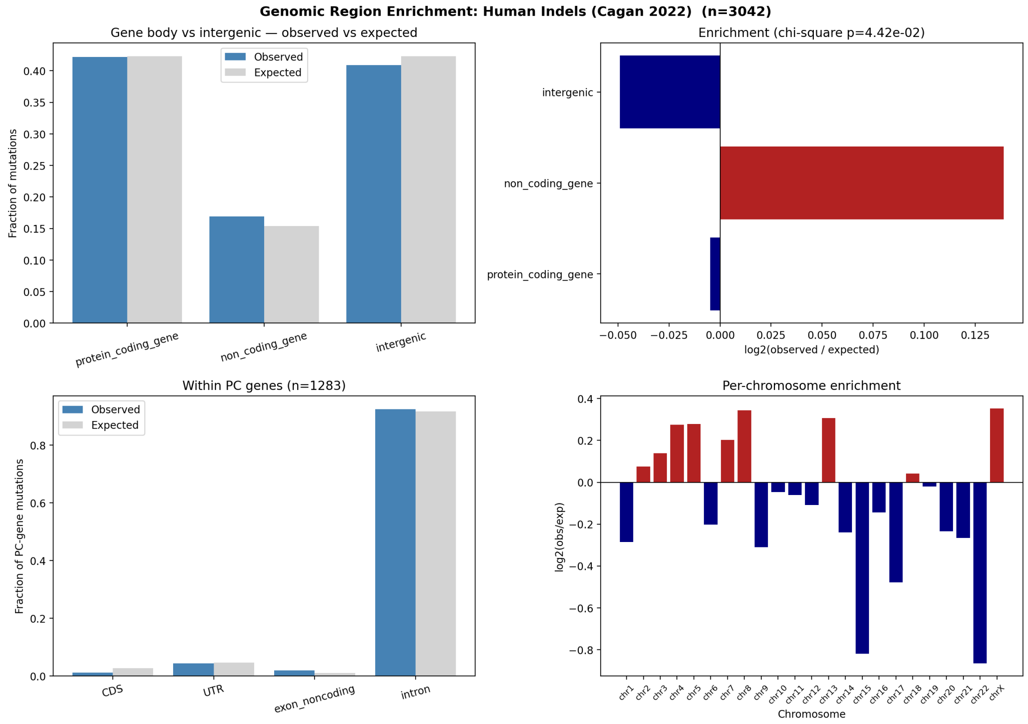


**Figure 10: Human Cagan somatic indels are non-randomly distributed across genomic regions.** Observed vs. expected indel counts across five region classes (CDS, UTR, noncoding exon, intron, intergenic), pooled across 28 human samples. Bar heights show observed (darker) and expected (lighter, bp-proportional) counts; inset values give $\log_2$(obs/exp). CDS is $\sim 2.3\times$ depleted ($\log_2 = -1.23$, 15 of expected 35.1 hits), the strongest signal; introns are near-neutral; non-coding exons are mildly enriched. $\chi^2$ goodness-of-fit: $p = 4.4 \times 10^{-2}$ at the top level, $p = 2.4 \times 10^{-5}$ within protein-coding genes. This is the canonical purifying-selection signature: crypt clones that acquire coding-disruptive variants are eliminated before they can be sampled. The matched 54-sample mouse cohort confirms the same picture with much sharper magnitudes ($12\times$ CDS depletion, $\chi^2\, p = 3.2 \times 10^{-104}$; Supplementary Figure S8). Logical implication: the *observed* somatic catalogue has already passed a selection filter, so its molecular effects represent an upper bound for neutral MA dynamics — and those effects are still negligible (R7, R8). *Species: human* (Cagan *et al.* 2022, 28 human colonic crypt samples, 3,042 indels; GENCODE v46 hg38 annotations).

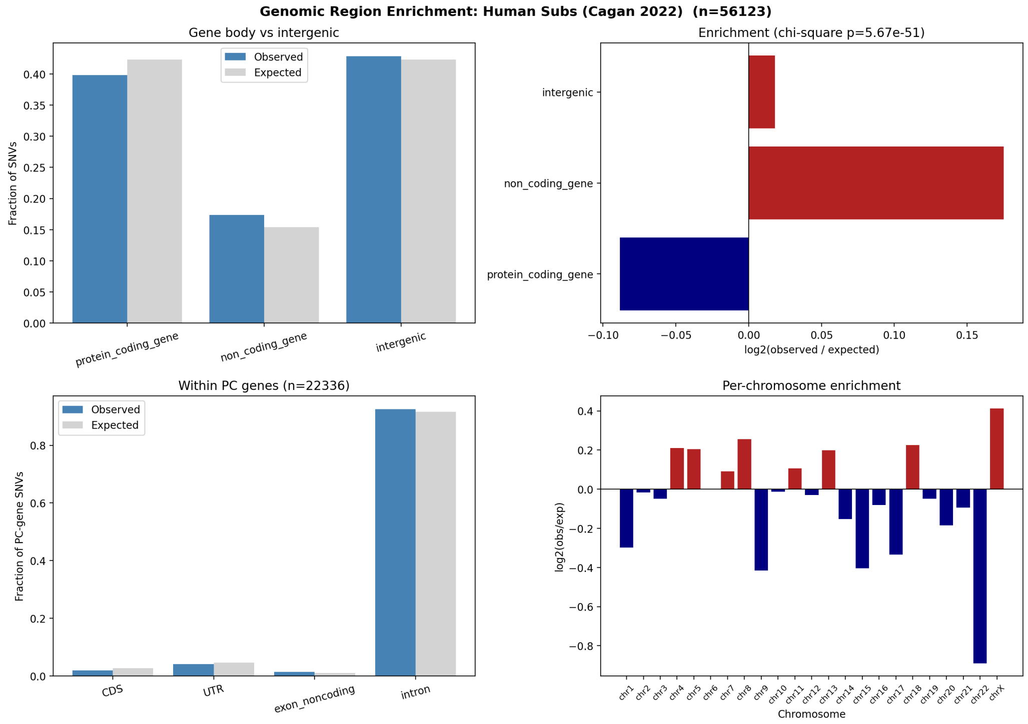


**Figure 11: Human Cagan somatic SNVs reproduce the indel purifying-selection signature in the same direction but at much smaller per-event magnitude.** Observed vs. expected SNV counts across the same five region classes, pooled across 28 human samples (56,123 SNVs total). CDS is $\sim 1.4\times$ depleted ($\log_2 = -0.52$, 427 of expected 611 hits); UTRs near-neutral; introns near-neutral; non-coding exons modestly enriched. $\chi^2$ goodness-of-fit: $p = 5.7 \times 10^{-51}$ at the top level, $p = 9.8 \times 10^{-22}$ within protein-coding genes. The much smaller log2 magnitude relative to the indel result of Figure 10 is biologically expected — most coding SNVs are synonymous or conservative missense and escape the strong selection that frameshift indels face — but the direction is preserved, confirming that even SNVs in surviving aged-crypt clones have been filtered against coding disruption. *Species: human* (Cagan *et al.* 2022, 28 human colonic crypt samples; GENCODE v46 hg38 annotations).

#### R7. Recurrently mutated genes are common fragile sites, not aging-related loci

I asked whether specific genes attract more somatic mutations than expected given their length plus a ±10 kb flank. Per-gene Poisson upper-tail tests with Benjamini–Hochberg FDR correction identified 99 protein-coding genes significantly enriched for indels across 54 samples at FDR $< 0.05$, and 41 genes for substitutions across 5 samples (Figure 12a,b). The top hits were overwhelmingly classic common fragile site (CFS) and long neural-adhesion genes: *Cntnap2* (34 indel hits, 23/54 samples), *Lsamp*, *Dmd*, *Lrp1b*, *Lrrc4c*, *Csmd1/3*, *Pcdh9/11x/15*, *Cdh12/18*, *Nrxn3*, *Dcc*, *Rbfox1*, *Dpp10*, *Cadm2*, *Sgcz*, *Kcnt2*. SNV and indel hits strongly overlapped at the same loci (hypergeometric $p = 3.8 \times 10^{-37}$; Figure 12b), indicating shared underlying fragility rather than mechanism-specific selection. These are neural genes not highly expressed in colonic epithelium, ruling out transcription-coupled mutagenesis of active loci; their shared property is extreme length (multi-megabase) and late replication, both hallmarks of CFS. Critically, hypergeometric overlap against five aging-associated gene sets revealed *no* enrichment for fibrosis, SASP, senescence, inflammatory, or EMT genes in either the indel or SNV hit list (all fold $< 1$, all $p > 0.8$; Figure 12c,d).

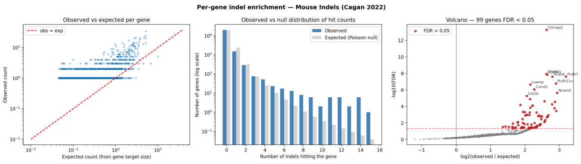


**(a)** Per-gene indel enrichment volcano: $\log_2$(obs/exp) on x-axis, $-\log_{10}$(FDR) on y-axis, across 54 samples. 99 protein-coding genes reach FDR$<0.05$ (above horizontal dashed line); top-labelled genes are dominated by CFS/long-gene loci. The horizontal banding reflects the discrete observed counts for a Poisson test.

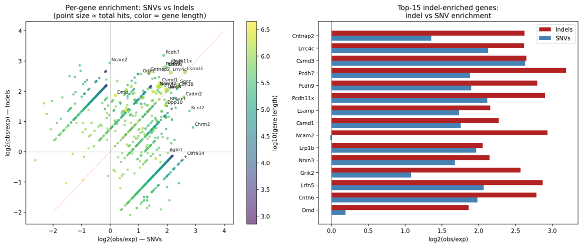


**(b)** Per-gene SNV vs. indel $\log_2$ enrichment scatter, with point size proportional to total hits and color-coded by gene length (log scale). The top-right quadrant is densely populated with labelled CFS genes (e.g., *Lrp1b*, *Lsamp*, *Csmd3*, *Cadm2*, *Rbfox1*, *Pcdh9/11x/15*), showing shared enrichment across variant classes. Hypergeometric overlap test for top-hit gene lists: $p = 3.8 \times 10^{-37}$.

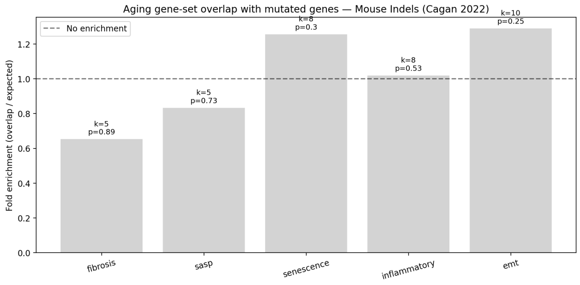


**(c)** Hypergeometric overlap bar chart for recurrently-hit indel genes (FDR$<0.05$) vs. five aging-associated gene sets (fibrosis, SASP, senescence, inflammatory, EMT). Bars show fold-enrichment with annotated $p$-values. None reach $p < 0.05$; all folds $< 1$ (depletion or near-baseline overlap).

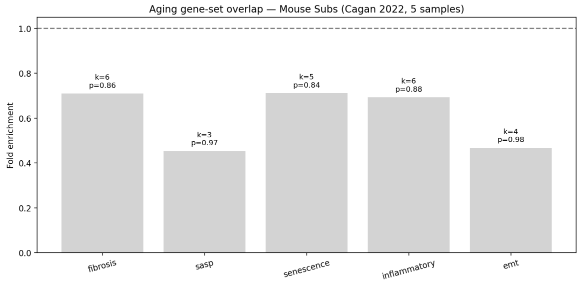


**(d)** Same hypergeometric test applied to the SNV hit list from the 5-sample cohort. Again no aging gene set is enriched; all $p > 0.8$.

**Figure 12: Recurrently mutated genes are common fragile sites, not aging-relevant loci.** Four panels show (a) the per-gene Poisson volcano for indels across 54 samples, (b) the per-gene SNV–indel agreement scatter, and (c,d) hypergeometric tests of the hit lists against five aging-associated gene sets. The top 20 recurrent hits — *Cntnap2*, *Lsamp*, *Dmd*, *Lrp1b*, *Lrrc4c*, *Csmd1/3*, *Pcdh9/11x/15*, *Cdh12/18*, *Nrxn3*, *Dcc*, *Rbfox1*, *Dpp10*, *Cadm2*, *Sgcz*, *Kcnt2* — are overwhelmingly long neural-adhesion CFS genes not expressed in colonic epithelium, arguing for replication-stress fragility rather than transcription-coupled mutagenesis or functional selection. The shared SNV–indel enrichment ($p = 3.8 \times 10^{-37}$) confirms common underlying fragility. No aging-associated gene set (fibrosis, SASP, senescence, inflammatory, EMT) is enriched in either hit list. *Species: mouse* (Cagan *et al.* 2022, 54 mouse samples for indels and 5 samples for SNVs; the human Cagan cohort of 3 donors is under-powered for per-gene recurrence testing).

### Discussion

The seven Results sections together constitute a quantitative baseline test of the mutation accumulation hypothesis as the driver of the colonic aging transcriptome. Per-variant AlphaGenome-predicted expression effects concentrate around a mean $|\log_2 \mathrm{FC}|$ of $\sim 10^{-4}$ at gene-body resolution — $\sim 5065\times$ below aging-scale mean $|\log_2 \mathrm{FC}|$ in the matched tissue — with the bulk of random SNVs producing sub-noise effects. Occasional single random SNVs ($\sim$ 1 in 20,000) reach aging-magnitude $|\log_2 \mathrm{FC}| \geq 1$ in their nearest gene, but the maximum random-SNV effect remains $\sim 13\times$ below the maximum aging effect, and additive multi-hit aggregation within a single gene does not amplify beyond the single-SNV maximum (R3a: per-(cell, gene) summed $|\log_2 \mathrm{FC}|$ caps at 2.248 across 154,113 pairs, equal to the largest single-SNV hit). At the level of a per-cell whole-transcriptome aggregate summed across thousands of distinct genes, the empirical $\sum \log_2 \mathrm{FC}$ at 4,000 SNVs/cell has $\sigma \approx 0.86$ over 60 simulated cells (Figure 5) — but this whole-cell aggregate is not the right comparator for aging differential expression, which is reported per gene; the load-bearing per-gene comparison remains $\sim 13\times$ on the maximum and $\sim 5065\times$ on the median. The finding is therefore not that no random SNV ever reaches aging magnitude in any single gene — a small fraction does — but that the bulk distribution sits orders of magnitude below the aging program, which simultaneously affects *thousands* of genes, and the additive cumulation that MA requires across the per-cell mutation load is statistically inaccessible.

##### Per-gene enrichment (R7)

For each protein-coding gene on accepted chromosomes I defined a target window of the gene body $\pm 10$ kb flank. Each mutation was assigned to a single gene via a nearest-gene rule: a mutation inside multiple overlapping target windows was placed in the gene whose *body* contained it, with ties broken by distance; otherwise to the nearest target-window boundary. Per-gene observed counts were tested under a Poisson upper-tail baseline model with expected $\lambda_i = N \cdot L_i^{\text{target}} / \sum_j L_j^{\text{target}}$, and $p$-values BH-corrected over the subset of genes with obs $\geq 1$. Gene-level recurrence was defined as `n_samples` $\geq 3$ and FDR $< 0.05$.

For both atlases, `.X` stores log1p-normalized counts; I inverted this with `np.expm1` before averaging, computed pseudobulk mean expression per age group, and defined per-gene aging $\log_2$FC as

$$\log_2\left(\frac{\bar{x}_{\text{old}} + 0.1}{\bar{x}_{\text{young}} + 0.1}\right),$$

#### Supplementary Figures

**Table S1: Mean $|\Sigma \log_2 \mathrm{FC}|$ across each Cagan / random-SNV mutation experiment computed over *all* 17,226 expressed colonic-epithelial genes (TMS), filling 0 for genes not assigned a mutation.** Mouse counterpart of main-text Table 1. Per-donor mean averaged across donors for Cagan rows; pooled experiment for the random row; aging row shows mean $|\log_2 \mathrm{FC}|$ over the same expressed-gene set. Per-donor hit-fractions shown in column 2. Ratio columns are aging value ÷ mutation value. *Species: mouse.*

| Dataset | hit-fraction (per donor) | mean $\lvert\log_2 \mathrm{FC}\rvert$ | max $\lvert\log_2 \mathrm{FC}\rvert$ | aging-mean ratio | aging-max ratio |
|---|---|---|---|---|---|
| Aging (TMS) | — | **1.541** | **10.74** | 1× | 1× |
| Random SNVs (mm10) | 7.7% | $8.8 \times 10^{-5}$ | 0.025 | 17,427× | 432× |
| Cagan SNVs (5 donors) | 3.9% | $5.5 \times 10^{-5}$ | 0.194 | 27,898× | 55× |
| Cagan indels (8 donors) | 1.2% | $3.3 \times 10^{-4}$ | 0.350 | 4,624× | 31× |

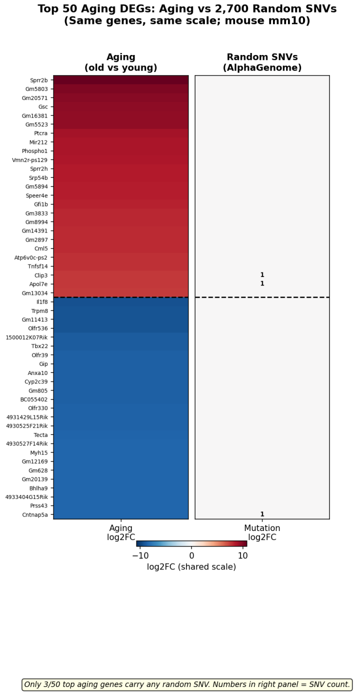


**(a)** Top 25 up- + top 25 down-regulated aging DEGs.

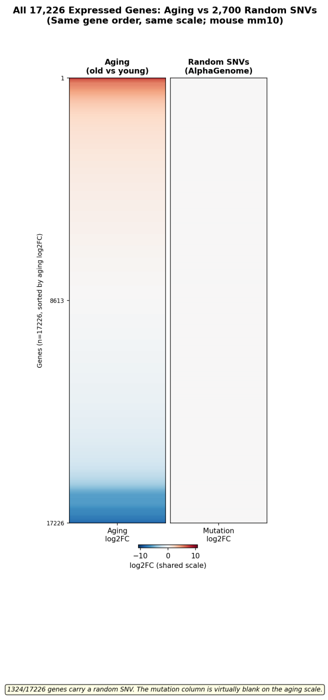


**(b)** All 17,226 expressed genes, sorted by aging $\log_2$FC.

**Figure S1: Mouse mm10 mirror of main-text Figure 2: 2,700 random mm10 SNVs produce per-gene effects that are invisible on the aging $\log_2$FC colour scale in mouse large-intestine epithelium.** Both panels share the diverging aging colour map. (*Left, a*) Top 25 up- and 25 down-regulated aging DEGs from Tabula Muris Senis large-intestine epithelium (3 m young vs 18/24/30 m old; 9,263 epithelial cells); only 3 of the 50 top aging genes are intersected by any random SNV, and the per-gene mutation column is uniformly neutral. Small numerals mark the SNV count per gene. (*Right, b*) All 17,226 expressed colonic-epithelial genes ordered by aging $\log_2$FC, with 1,324 ($\sim$ 8%) carrying a mapped random SNV but no visible colour on the shared scale. *Cell-type filter:* the "colon-epithelial" aging reference pools four Cell Ontology cell types (enterocyte of epithelium of large intestine, epithelial cell of large intestine, intestinal crypt stem cell, large intestine goblet cell), not a single cell type. *Species: mouse* (mm10 SNVs scored in mouse colon, UBERON:0000059; Tabula Muris Senis aging reference).

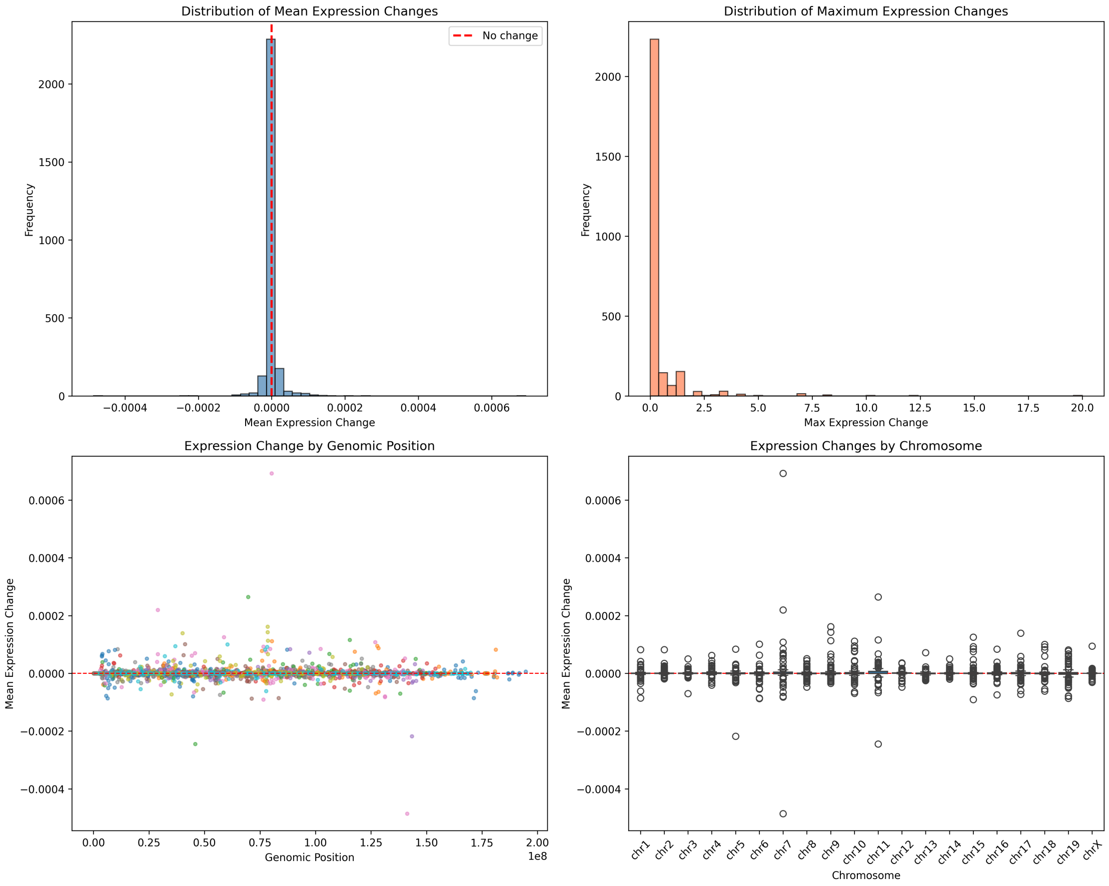


**Figure S2: Mouse mm10 mirror of the human R1 random-SNV simulation (main-text Figure 1; 2,700 random mm10 SNVs).** The same random-genome-wide simulation re-run on mm10 with mouse colon ontology, yielding the same near-zero per-variant effect distribution. *Species: mouse* (mm10 SNVs scored in mouse colon).

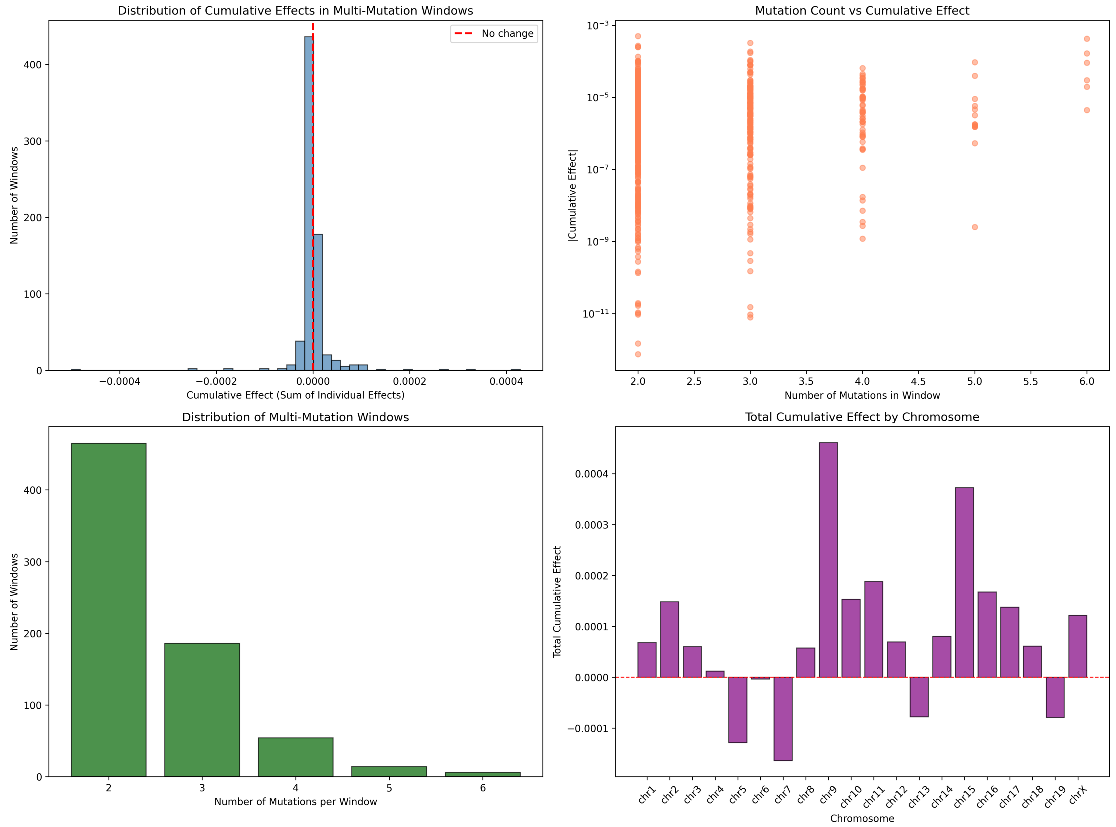


**Figure S3: Mouse mm10 mirror of the human R2 1-Mb combined-effect analysis (main-text Figure 3).** Variants binned into 1-Mb windows; cumulative window effects scale roughly linearly with variant count and remain orders of magnitude below the aging $\log_2$FC scale. *Species: mouse* (mm10).

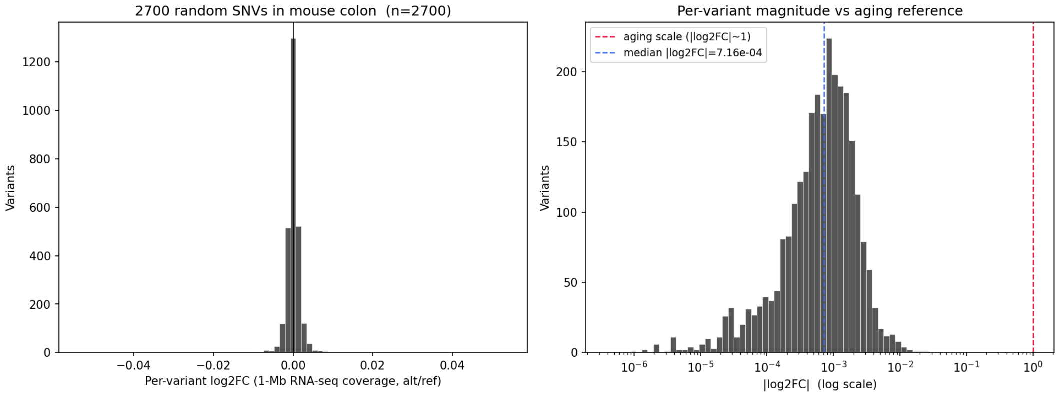


**Figure S4: Per-variant $\log_2$FC distribution from the mouse mm10 random-SNV simulation.** Companion histogram to Supplementary Figure S2. The distribution is tightly centred near zero, mirroring the human hg38 result. *Species: mouse* (mm10).

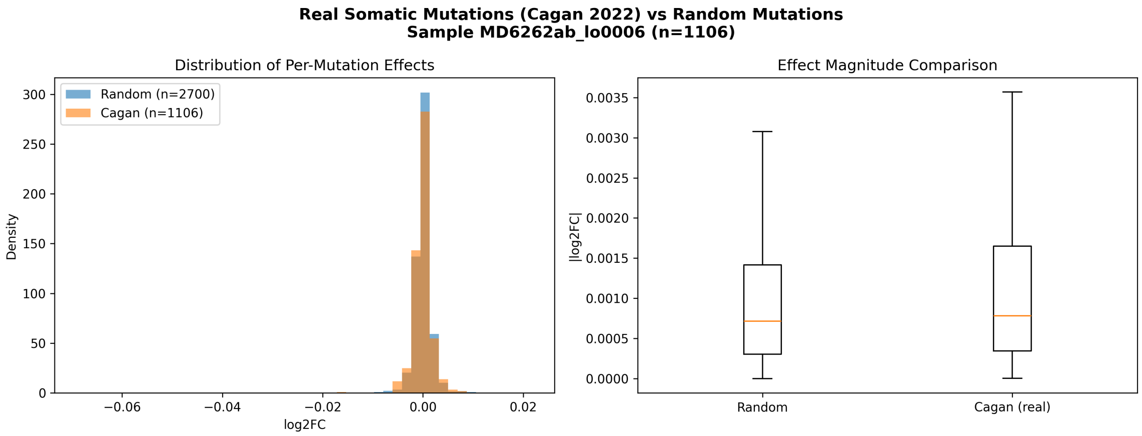


**(a)** Comparison of per-variant $\log_2$FC distributions in the mouse cohort (primary sample MD6262ab_lo0006, $n = 1{,}106$ SNVs against the matched mm10-native 2,700-random-SNV baseline; colon UBERON:0001157, mm10-native predictions). The mouse Cagan and random distributions overlap closely (mean $|\log_2 \mathrm{FC}|$ $1.34 \times 10^{-3}$ vs $1.12 \times 10^{-3}$, 1.20×).

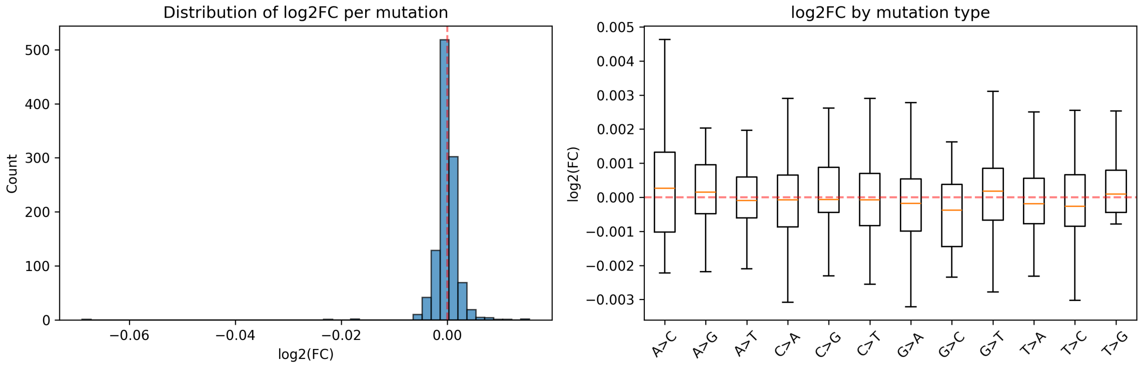


**(b)** Per-variant $\log_2$FC distribution for the same mouse sample, with 2-fold-change threshold $|\log_2 \mathrm{FC}| = 1$ marked.

**Figure S5: Mouse-cohort equivalent of Figure 7.** *Species: mouse.* Retained because the mouse cohort is larger ($n = 54$ crypts; 9,177 SNVs across five detailed samples, scored mm10-natively) than the three-donor human cohort shown in the main-text figure; conclusions are identical in both species.

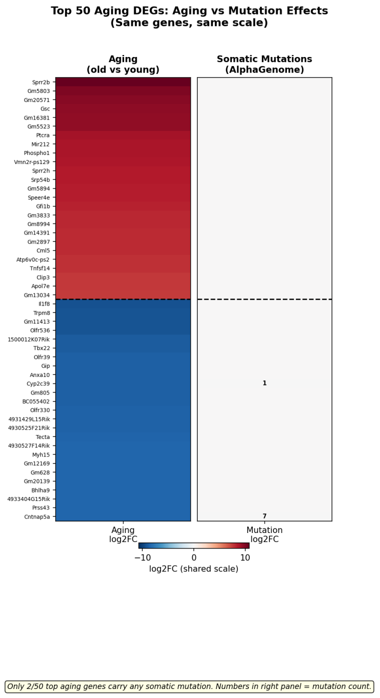

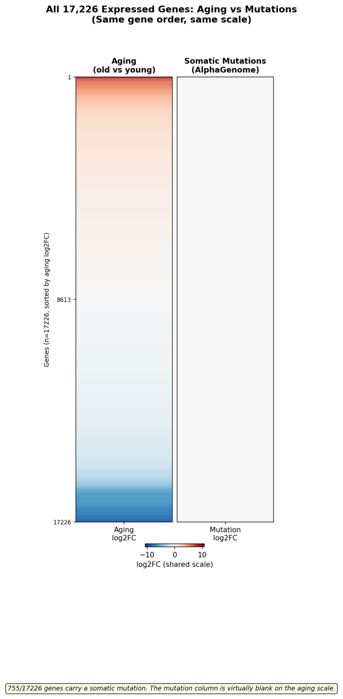


**(a)** Top 25 up- + top 25 down-regulated aging DEGs.

**(b)** All ∼17,000 expressed genes, sorted by aging $\log_2$FC.

**Figure S6: Mouse-on-mouse Cagan-vs-aging heatmap (R4 supplement).** 1,781 real somatic mutations from Cagan sample MD6267ab_lo0015 plotted side-by-side with Tabula Muris Senis aging $\log_2$FC on a shared diverging colour scale, in mouse large-intestine epithelium. (*Left*) Top 25 up + 25 down aging DEGs; the aging column paints the full $[-10, +12]$ gradient while the mutation column is entirely neutral. (*Right*) All $\sim$ 17k expressed genes ordered by aging $\log_2$FC; mutation column blank along the entire axis despite 755 of those genes carrying $\geq 1$ real somatic mutation. Max aging-to-mutation ratio $\approx 840\times$, median $\approx 990\times$ at 1-Mb-window scoring. *Species: mouse* (Tabula Muris Senis; Cagan *et al.* 2022 mouse sample MD6267ab_lo0015).

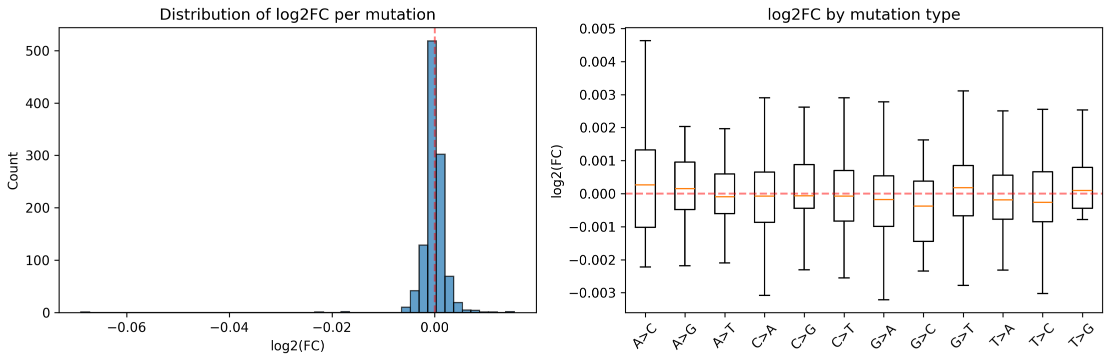


**(a)** Distribution of per-indel predicted expression $\log_2$FC for primary mouse sample MD6264o_lo0009 ($n = 457$ indels). Histograms split insertions and deletions; rug marks at bottom indicate individual indels. The distribution median sits at $|\log_2\text{FC}| \approx 0.03$, with a tail extending toward $|\log_2\text{FC}| \sim 1$ for a small number of splice- or CDS-disrupting events.

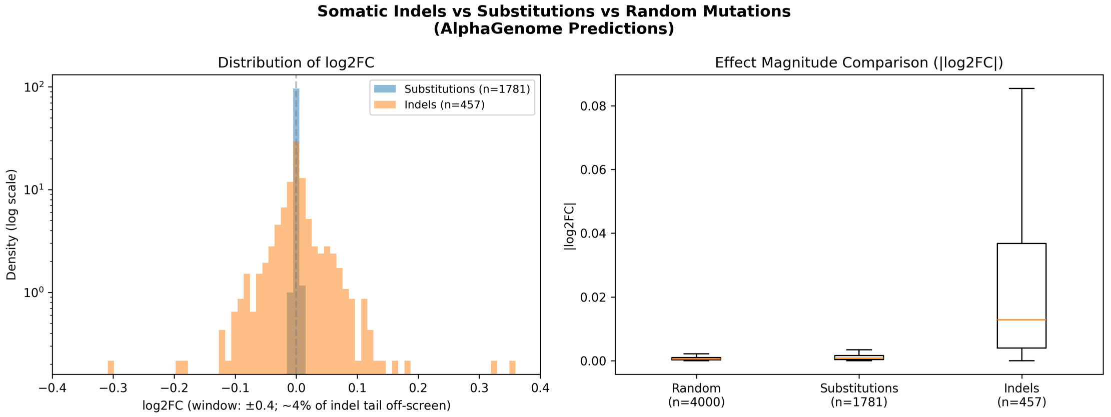


**(b)** Direct mouse indel-vs-substitution comparison within the same sample: overlaid distributions of per-event $|\log_2 \text{FC}|$ and box plots of absolute effect, with medians annotated. The 24.4× per-event ratio is evident, as is the heavier indel tail reflecting occasional frame- or splice-disrupting events. Both distributions remain below the aging-scale $\log_2$FC reference.

**Figure S7: Mouse-cohort counterpart of main-text Figure 9 ($n = 8$ mouse samples, 3,233 indels in total; primary detail sample MD6264o_lo0009, $n = 457$).** The mouse cohort confirms the per-event indel/SNV amplification observed in the human cohort: mean $|\log_2 \text{FC}|$ across 409 indels of sample MD6267ab_lo0003 was 0.030 vs 0.0012 for substitutions in the same sample (24.4× larger per mutation). The per-gene worst case observed across all 3,233 mouse indels in all 8 samples was $|\sum \log_2 \text{FC}| = 0.82$ at *Spertl* (4 indels in the same gene), still below the magnitude of a single top aging DEG. *Species: mouse* (Cagan *et al.* 2022 mouse colonic crypts).

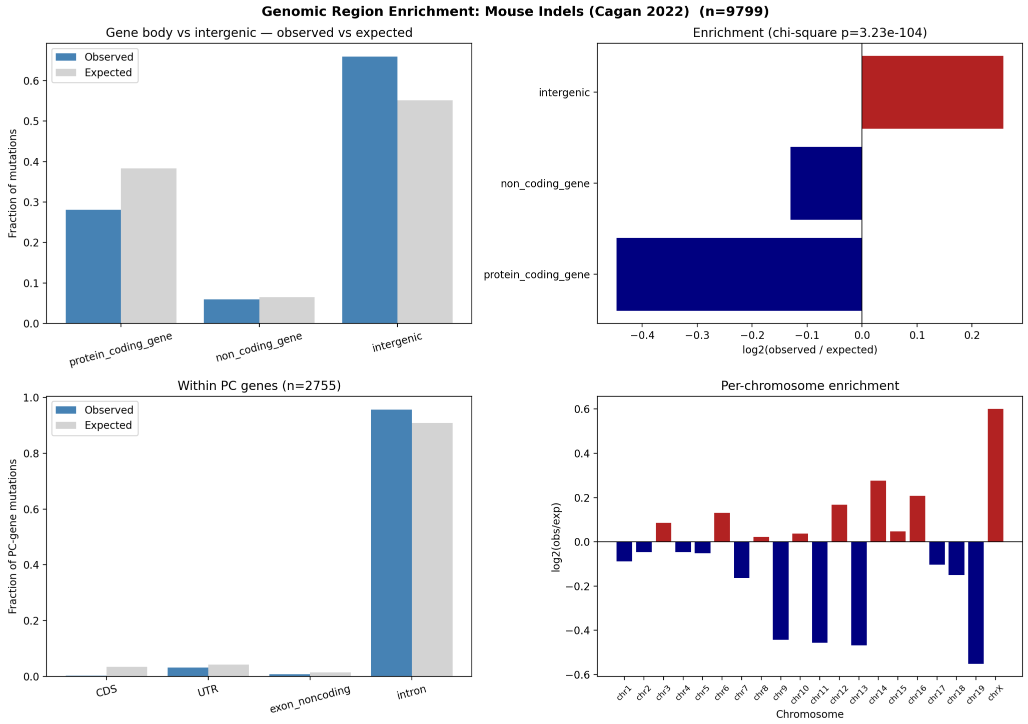


**Figure S8: Mouse-cohort counterpart of main-text Figure 10 ($n = 54$ samples, 9,799 indels).** The same purifying-selection signature seen in the human cohort, with much sharper magnitudes owing to the larger sample size: 12× CDS depletion ($\log_2 = -3.58$, only 8 of 2,755 protein-coding-gene indels in CDS), modest intergenic enrichment ($\log_2 = +0.26$), and a top-level $\chi^2$ goodness-of-fit p-value of $3.2 \times 10^{-104}$ that is power-driven by the 9,799-indel sample size (the same fractional deviations against $\sim 3\times$ fewer indels in human gave $p = 4.4 \times 10^{-2}$ at the top level). *Species: mouse* (Cagan *et al.* 2022, 54 mouse colonic crypts; GENCODE vM25 mm10 annotations).

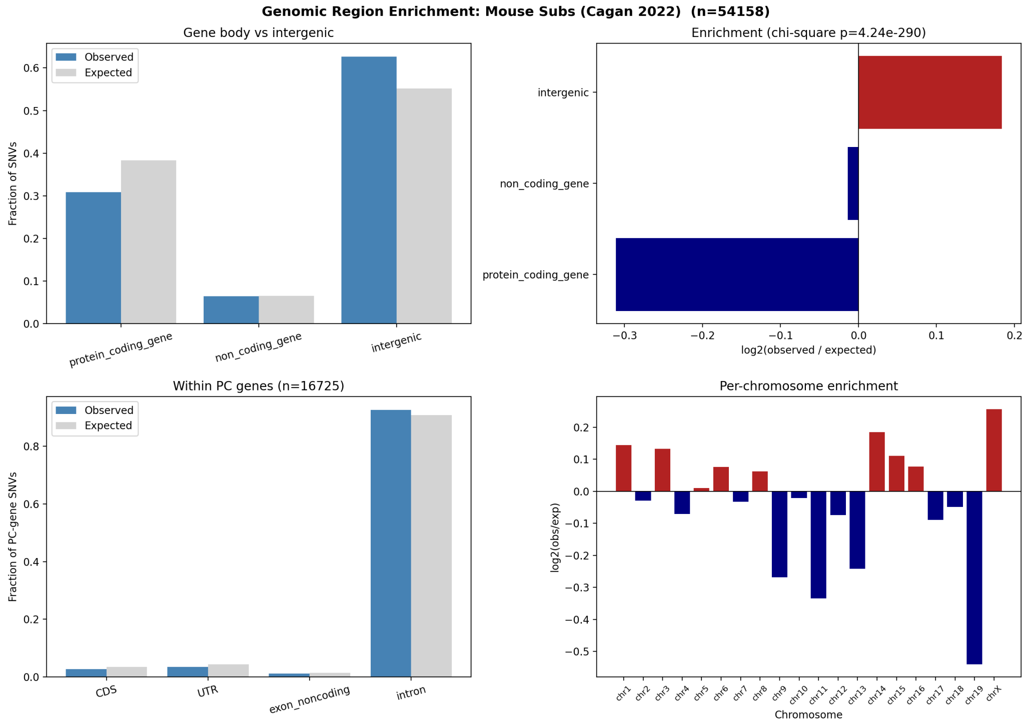


**Figure S9: Mouse-cohort counterpart of main-text Figure 11 ($n = 54$ samples, 54,158 SNVs).** Region-class enrichment of mouse SNVs reproduces the human direction at slightly weaker per-event magnitude: CDS $\sim 1.27\times$ depleted ($\log_2 = -0.34$, 460 of expected 583 hits), UTRs and exon-noncoding regions modestly depleted ($\log_2 \approx -0.32$ to $-0.36$), introns near-neutral, intergenic modestly enriched ($\log_2 = +0.18$). Within-PC $\chi^2\, p = 1.0 \times 10^{-15}$; top-level $\chi^2\, p = 4.2 \times 10^{-290}$ (the latter dominated by the 54,158-SNV sample size; the biologically meaningful quantity is the modest $\log_2$ effect size, which is roughly an order of magnitude smaller than the indel result of Supplementary Figure S8). *Species: mouse* (Cagan *et al.* 2022, 54 mouse colonic crypts; GENCODE vM25 mm10 annotations).
