## Supplementary material for "Testing the mutation accumulation hypothesis in aging with AlphaGenome": Manuscript LaTex file: paper.pdf.backup.pdf

Arthur Fischbach*

May 4, 2026

**Abstract**

The mutation accumulation (MA) hypothesis posits that somatic mutations progressively escape selection and degrade tissue function during aging. Direct tests of this idea have been limited by the difficulty of predicting, at scale, the molecular consequences of individual somatic variants. Here we use AlphaGenome, a sequence-to-function deep learning model, to systematically score the predicted transcriptional, chromatin-accessibility, and histone-mark impact of somatic mutations under a nested series of designs spanning individual variants, co-occurring variant bundles, and real mutation catalogues. *First*, we establish a genome-wide null by scoring 4,000 random single-nucleotide variants (SNVs) in colon tissue, with 1-Mb-window combined-effect tests and a 100-cell × 100-SNV Monte Carlo pilot. *Second*, we extend the null to gene-body resolution with a 60-cell × 4,000-SNV simulation and pseudobulk RNA-seq aggregation. *Third*, we analyze the real somatic mutation catalogue of Cagan *et al.* (*Nature*, 2022), scoring 54,158 substitutions and 9,799 indels from 54 mouse colonic crypts plus three human samples, together with region- and gene-level enrichment tests against GENCODE.

Across all arms, individual random mutations produce predicted expression changes whose bulk distribution sits about three orders of magnitude below the tissue's own aging program: at gene-body resolution, median $|\log_2 \mathrm{FC}|$ is $1.04 \times 10^{-3}$ for random SNVs versus 0.572 for aging-associated differential expression in human large intestine (median ratio $\sim 550\times$), while the maximum random-SNV effect (2.248, *TMIGD2*, single SNV) sits only $\sim 3\times$ below the maximum aging effect (6.98). Real somatic mutations from the Cagan cohort produce smaller per-variant effects than the random baseline. Random-mutation effects propagate through a sensitivity hierarchy DNase > ATAC > CAGE > RNA-seq spanning $\sim$1000×, consistent with regulatory buffering. A 100-cell × 100-SNV Monte Carlo cohort confirms that per-cell cumulative effects under the null are tightly centred at zero ($\sigma_{\mathrm{cum}} = 2.75 \times 10^{-4}$), and a 60-cell × 4,000-SNV gene-body simulation shows that per-(cell, gene) additive multi-hit aggregation is empirically bounded at the single-SNV maximum. Aggregating the 60 simulated cells into a pseudobulk RNA-seq comparison versus an unmutated reference tightens the per-gene bound to $\sim 36\times$ below the maximum aging effect and yields a per-gene Pearson correlation of $r = -0.027$ ($p = 0.13$) with aging differential expression — random SNVs neither match aging in magnitude nor in direction. Real somatic mutations concentrate at common fragile sites (CFS) dominated by long neural-adhesion genes (*Cntnap2*, *Lsamp*, *Csmd1/3*, *Lrp1b*, *Pcdh9/11x/15*), with no enrichment for any aging gene set (all hypergeometric $p > 0.8$), and coding exons are depleted 12× ($\log_2 = -3.58$, $\chi^2\, p = 3.2 \times 10^{-104}$). These results argue against a simple mutation-accumulation explanation for the age-associated transcriptional drift of colonic epithelium and redirect attention to epigenetic and regulatory mechanisms.

We exploit this capability to perform a systematic, adversarial test of MA as the driver of transcriptional aging in the colon — the tissue with the richest paired somatic mutation catalogue

[1] and well-characterized aging single-cell transcriptomes [3–5]. The study is structured as a nested series of Results sections, each designed to close one escape route through which MA could evade the magnitude argument. R1, R2 and R4 establish the genome-wide null at 1-Mb window resolution: scoring 4,000 random SNVs at a biologically realistic per-crypt burden (R1), testing whether co-occurring variants amplify within shared cis-regulatory windows (R2), and a Monte Carlo pilot over 100 cells × 100 SNVs (R4). R4b sharpens the null by re-scoring at gene-body resolution across a 60-cell × 4,000-SNV cohort, and R4c aggregates that cohort into a pseudobulk RNA-seq comparison against the aging program. R7–R10 then test the *real* Cagan somatic mutation catalogue (54,158 substitutions and 9,799 indels across 54 mouse crypts plus three human samples) for per-variant effect magnitude, for per-event effect of structural indels, for non-random positional distribution, and for gene-level recurrence. R11 quantifies the scale gap explicitly by placing the AlphaGenome-predicted per-gene effect side-by-side with the young-versus-old $\log_2$FC of the matched tissue's transcriptome.

Every arm converges on the same conclusion: individual somatic mutations produce predicted transcriptional effects approximately three orders of magnitude below the tissue's own aging program, with no tissue, gene set, variant class, or positional refinement rescuing the magnitude. Real somatic mutations from aged crypts are not more impactful than random — they are strictly *less* so, a consequence of the purifying-selection signature visible in their genomic-region distribution. We interpret this as strong evidence against a simple MA account of the colonic aging transcriptome, and as a redirection of mechanistic attention toward epigenetic and regulatory mechanisms of aging [18–20] — mechanisms for which AlphaGenome's multi-modal architecture offers a natural next-step predictive framework.

#### Results

##### R1. AlphaGenome predicts near-zero expression effects for random single-nucleotide variants genome-wide

We chose a sample size of 4,000 SNVs to match the upper envelope of somatic single-nucleotide mutation burdens observed in individual human colonic crypts from aged donors, which reach on the order of a few thousand variants per cell by the eighth decade of life [1]; this anchors the null baseline to a biologically realistic per-cell mutation load rather than an arbitrary simulation size. We chose colon tissue for this and downstream analyses because the reference somatic mutation catalogue we validate against [1] is itself built on colonic crypts. To establish this null baseline for the molecular impact of a single mutation, we sampled 4,000 SNVs uniformly across the human genome (hg38) and scored each with AlphaGenome [2] in colon tissue (UBERON:0001157), requesting `RNA_SEQ`, `ATAC`, and `CHIP_HISTONE` outputs in a single call per variant. All 4,000 predictions succeeded. The per-variant mean absolute expression change was sharply concentrated at zero (median $\approx 1 \times 10^{-6}$), with a pronounced but small heavy tail: the top 1% of variants reached per-variant `max_expression_change` $> 1$, and the single largest `total_expression_change` observed was 1,104 at chr17:81,415,658 T→A. 99.9% of variants produced absolute effects $< 10^{-4}$. This random-variant effect distribution, summarized in Figure 1, serves as the reference null used throughout Sections R2–R10. As the age-matched human reference for the colonic-epithelial transcriptome, we computed pseudobulk $\log_2$FC between young (donor TSP26, 37 y; 597 cells) and old (donors TSP2, TSP14, TSP25, TSP27, ages 56–61 y; 8,981 cells) large-intestine epithelial cells from the Tabula Sapiens v2 atlas [4, 5]. When the 4,000 variants are mapped to their nearest protein-coding gene (GENCODE v46, hg38) and the per-variant `mean_expression_change` is summed per gene, only 604 of 3,579 colon-epithelial-expressed genes carry any random SNV, and the per-gene totals are invisible on the aging $\log_2$FC colour scale (Figure 2): the max $|\text{mean}\Delta\text{expr}|$ is $1.23 \times 10^{-3}$ versus aging max $|\log_2 \text{FC}| = 6.98$, a gene-level scale ratio of 5,664× on the maximum and $\sim 2.2 \times 10^5\times$ on the median. This per-gene view anticipates the quantitative scale argument developed for real somatic mutations in Section R11.

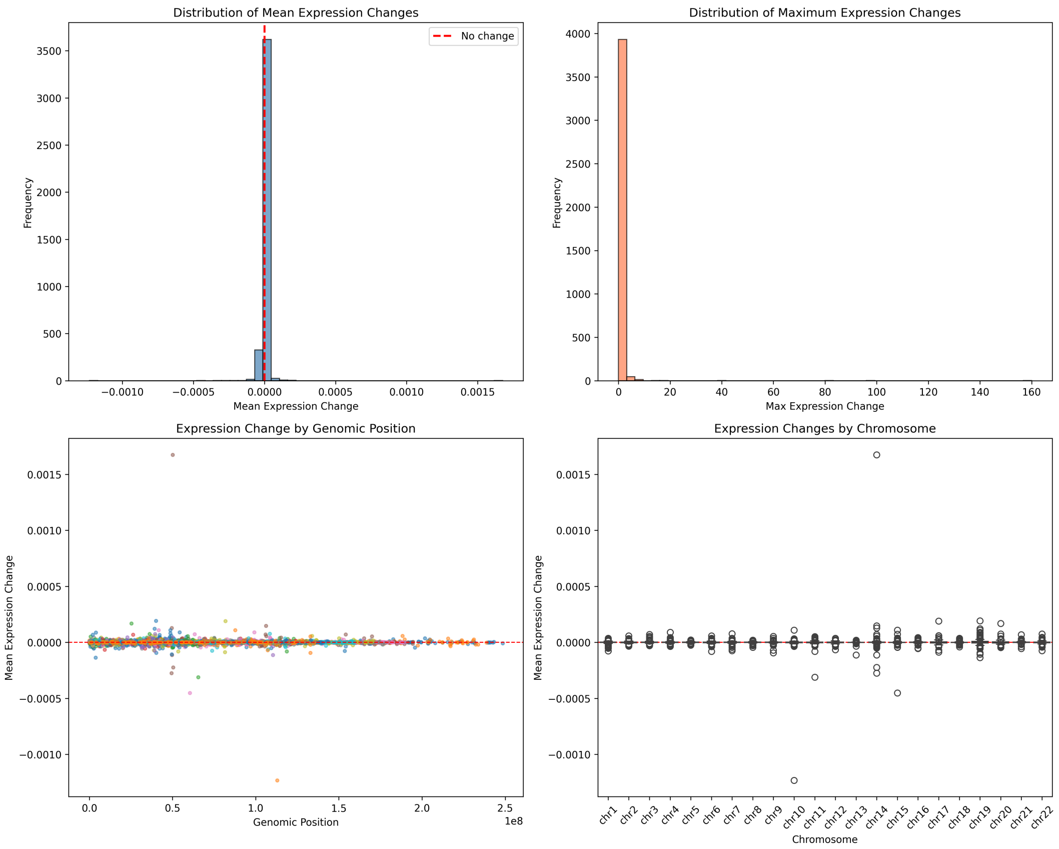


**Figure 1: Null distribution of AlphaGenome-predicted effects for 4,000 random genome-wide single-nucleotide variants in colon tissue.** Each of the four panels summarizes one facet of the per-variant effect distribution. (*Top left*) Histogram of mean expression change across the 1 Mb output window, centered tightly on zero with the bulk of mass within $|x| < 10^{-4}$; no symmetric tail, consistent with most random substitutions falling in unconstrained intergenic or intronic sequence. (*Top right*) Distribution of per-variant max expression change, spanning four orders of magnitude from $10^{-4}$ to $\sim 4$; outliers represent rare variants that intersect active regulatory elements or splice boundaries. (*Bottom left*) Distribution of total expression change (sum of $|\log_2 \mathrm{FC}|$ across affected genes in the window), dominated by small effects with a single extreme observation (chr17:81,415,658 T→A, total = 1,104) that would otherwise dominate a linear axis and motivates the log-scale display. (*Bottom right*) Scatter of max vs. mean expression change per variant, confirming that the rare high-max outliers correspond to localized effects that do not lift the window-average appreciably — evidence that even strong local perturbations produce a bounded, tightly-buffered footprint at the gene level. Colon tissue (UBERON:0001157); RNA-seq + ATAC + ChIP-histone requested simultaneously; 100% prediction success. *Species: human* (4,000 SNVs sampled from hg38, scored in human colon).

**Table 1: Mean $|\Sigma \log_2 \mathbf{FC}|$ across each Cagan / random-SNV mutation experiment computed over *all* 3,579 expressed colonic-epithelial genes (TSP v2), filling 0 for genes not assigned a mutation.** For each Cagan dataset the per-donor mean is computed and then averaged across donors; the aging row shows the mean $|\log_2 \mathrm{FC}|$ over the same expressed-gene set. This framing is the direct apples-to-apples with the aging pseudobulk because both averages run over the identical gene set, capturing simultaneously the magnitude of effects when they happen *and* the sparsity of the mutation hit pattern (per-donor hit-fraction shown in column 2). Ratio columns are aging value ÷ mutation value. *Species: human.*

| Dataset | hit-fraction (per donor) | mean $\|\log_2 \mathrm{FC}\|$ | max $\|\log_2 \mathrm{FC}\|$ | aging-mean ratio | aging-max ratio |
|---|---|---|---|---|---|
| Aging (TSP v2) | — | **0.795** | **6.98** | 1× | 1× |
| Random SNVs (hg38) | 16.9% | $1.6 \times 10^{-4}$ | 0.052 | 5,065× | 134× |
| Cagan SNVs (3 donors) | 13.0% | $1.8 \times 10^{-4}$ | 0.527 | 4,463× | 13× |
| Cagan indels (9 donors) | 1.1% | $2.1 \times 10^{-4}$ | 0.173 | 3,790× | 40× |

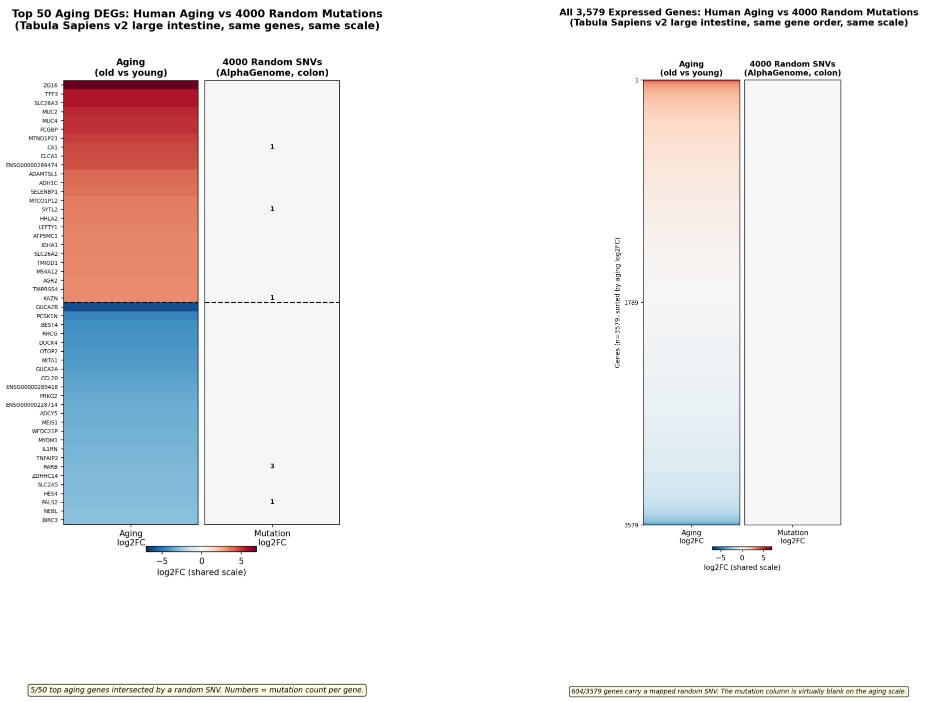


**(a)** Top 25 up- + top 25 down-regulated aging DEGs.

**(b)** All 3,579 expressed genes, sorted by aging $\log_2$FC.

**Figure 2: 4,000 random SNVs produce per-gene effects that are invisible on the aging $\log_2$FC colour scale in human large-intestine tissue.** *Species: human* (Tabula Sapiens v2 aging reference; AlphaGenome random-SNV simulation in human colon, UBERON:0001157). Both panels share a common diverging colour map centred at zero, with the scale set by the aging $|\log_2 \mathrm{FC}|$ range so that the two columns can be compared without rescaling. (*Left, a*) The top 25 up-regulated and 25 down-regulated aging DEGs from Tabula Sapiens v2 large-intestine epithelium [4, 5] (young donor TSP26, 37 y vs old donors TSP2/TSP14/TSP25/TSP27, 56–61 y), plotted in the aging column as a rich red-to-blue gradient spanning approximately $[-7, +7]$; the mutation column (summed per-gene AlphaGenome per-variant $\log_2$FC — $\log_2((s_{\mathrm{alt}} + 1)/(s_{\mathrm{ref}} + 1))$ over the 1-Mb RNA-seq window — across the 4,000 random SNVs assigned to the nearest hg38 protein-coding gene, with no species conversion) is uniformly neutral, and only a handful of top aging genes are hit by any random SNV at all. Small numerals in the right panel mark the mutation count per gene. (*Right, b*) All 3,579 expressed epithelial genes ordered along the y-axis by aging $\log_2$FC, with mutation contribution shown on the right using the identical colour scale. The aging column paints the expected red-top / blue-bottom gradient; the mutation column shows 604 genes carrying a mapped random SNV but no visible colour at this scale — the per-gene mutation sum is three to five orders of magnitude below aging-scale fold changes. On the per-variant 1-Mb-window $\log_2$FC axis used for these heatmaps, the max aging-to-mutation ratio is $\sim 134\times$ ($6.98/5.20 \times 10^{-2}$) and the median ratio is $\sim 1{,}120\times$ ($0.572/5.12 \times 10^{-4}$); the bulk of random SNVs produce sub-noise per-gene effects. At gene-body resolution (R4b, Figure 4) the maximum random-SNV $|\log_2 \mathrm{FC}|$ rises to 2.248 (*TMIGD2*, single SNV in 60×4,000 simulation), bringing the max-effect ratio down to $\sim 3\times$ relative to TSP aging max 6.98, while the median ratio remains $\sim 550\times$. Donor caveat: only one TS v2 large-intestine donor (TSP26) is below 50 y, so the young arm is single-donor; the $\sim 550\times$ median scale gap is many orders of magnitude beyond any donor variance, and the conclusion is robust to this choice. A confirmatory mouse-on-mouse version using Tabula Muris Senis with 4 vs 4 donors is shown in Supplementary Figure S6. *Cell-type filter:* the "colon-epithelial" aging reference pools seven Cell Ontology cell types (enterocyte of epithelium of large intestine, BEST4+ enterocyte, large intestine goblet cell, intestinal crypt stem cell of colon, transit amplifying cell of colon, paneth cell of colon, tuft cell of colon), not a single cell type.

#### R2. Combined effects in the same 1-Mb window do not amplify

Individual mutations could in principle sum to non-trivial effects if they co-localize within a common cis-regulatory window. We tested this by grouping the 4,000 random variants into non-overlapping 1-Mb genomic windows and measuring the *expected additive* cumulative effect (sum of per-variant mean expression changes) for windows harboring multiple variants. Windows with 10 co-occurring random SNVs reached a cumulative expected effect of only $\sim 3\times10^{-4}$, barely distinguishable from the single-variant scale and orders of magnitude below aging-scale $\log_2$FC. The cumulative-effect distribution remained approximately symmetric around zero with no heavy right tail, showing that the upper-bound assumption of perfect additivity still yields negligible totals at realistic variant densities (Figure 3). This rules out a simple “hotspot accumulation” mechanism whereby local co-occurrence amplifies individually weak mutations.

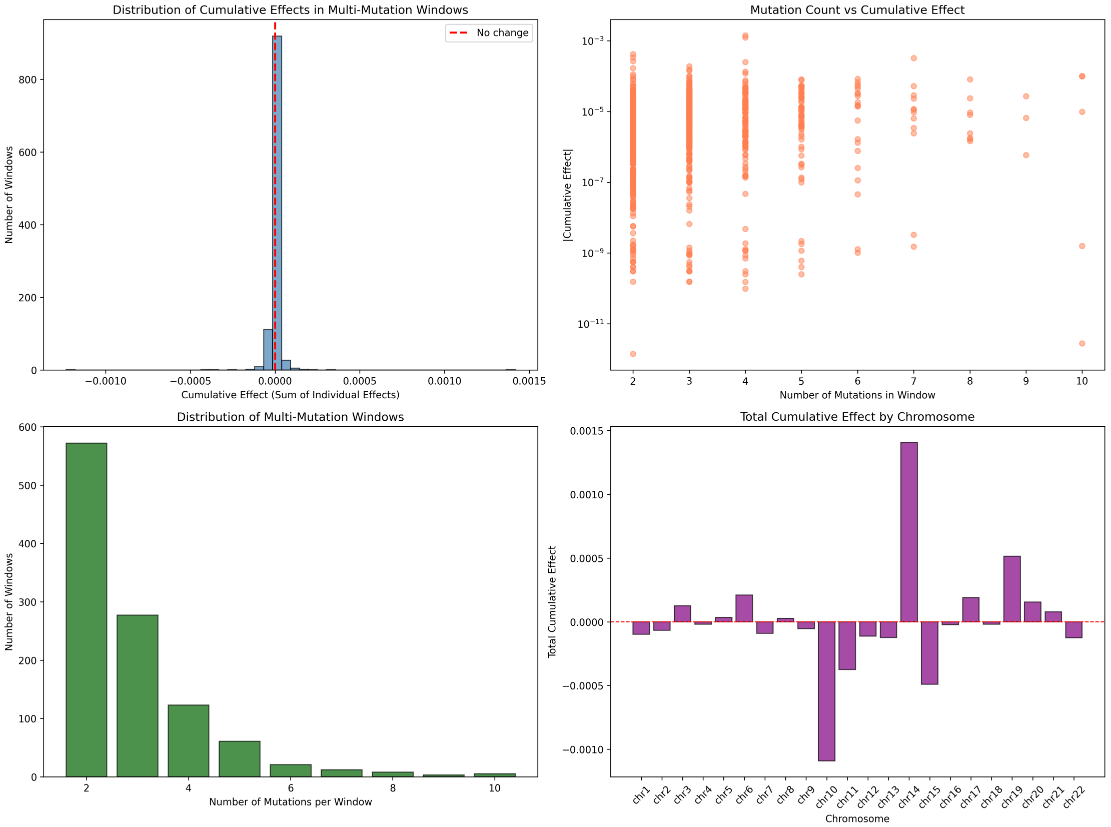


**Figure 3: Combined-effect analysis of random SNVs grouped into 1-Mb windows.** Random variants were binned by their genomic coordinate into $\sim$ 3,000 non-overlapping 1-Mb windows. Panels summarize, for each window: (1) how many variants it contains (most windows 0–2), (2) the absolute mean expression change of the window’s largest single variant, and (3) the cumulative window effect computed as the sum of per-variant mean changes (additive-expectation upper bound). The cumulative distribution mirrors the single-variant scale, with even windows containing 10 co-occurring variants reaching only $\sim 3\times10^{-4}$ cumulative effect. No window shows synergistic amplification. This result is critical because it forecloses the simplest MA-compatible escape route: “mutations at random, but stack at shared cis-regulatory bundles.” At realistic variant density, they don’t stack to functionally consequential levels. *Species: human* (random SNVs in human colon, UBERON:0001157).

#### R4. Pilot Monte Carlo cohort confirms near-zero per-cell cumulative effect at small per-cell SNV counts; realistic load measured directly in R4b

As a small-scale precursor to the realistic 4,000-SNV per-cell simulation of R4b, we ran a 100-cell $\times$ 100-SNV pilot Monte Carlo in colon tissue at window-level resolution. The 100 per-cell cumulative effects (sum of per-variant mean expression changes) were tightly centred at zero with $\mu_{\text{cum}} = -2.4\times10^{-5}$, $\sigma_{\text{cum}} = 2.75\times10^{-4}$, and observed maximum $1.155\times10^{-3}$ (0.115%) — two orders of magnitude below a 10% aging-scale change. Because the per-variant gene-body distribution turns out to be heavy-tailed (R4b), Gaussian $\sigma$-scaling from this 100-SNV pilot would underestimate the realistic-load tail; we therefore measure the 4,000-SNV per-cell burden directly in R4b rather than extrapolate.

##### R4b. Gene-level scoring reveals occasional aging-magnitude effects from single random SNVs, but multi-hit additive amplification is empirically bounded

The R4 Monte Carlo summed RNA-seq predictions over the entire 1-Mb output window, which dilutes per-variant signal because most of that window is non-transcribed sequence. The biologically meaningful unit for comparison against aging differential expression is the nearest protein-coding gene's body, since aging $\log_2$FC is reported per gene rather than per 1-Mb interval. We therefore re-ran the random-SNV simulation at gene-body resolution: 60 independent simulated cells $\times$ ~4,000 random hg38 SNVs each (240,000 AlphaGenome calls; 203,405 with finite gene-body $\log_2$FC), each variant scored as $\log_2((s_{\text{alt}} + 1)/(s_{\text{ref}} + 1))$ over the nearest GENCODE v46 protein-coding gene body in human colon (UBERON:0001157). The per-variant gene-body distribution differs sharply from the window-level proxy: standard deviation $1.43 \times 10^{-2}$ (vs $1.7 \times 10^{-3}$ at window level, a $\sim 8\times$ widening) and a maximum $|\log_2 \mathrm{FC}| = 2.248$ (vs 0.085, single SNV in *TMIGD2*). Ten of 203,405 variants reach $|\log_2 \mathrm{FC}| \geq 1$ (~1 in 20,000; the top hits are *TMIGD2* +2.248, *RNASE2* −1.546, *NEUROD4* −1.520, *CHRM1* +1.423, *GABRA1* +1.412), and 157 of 203,405 (0.077%) reach $|\log_2 \mathrm{FC}| \geq 0.138$ (the 10%-expression-change threshold). The median $|\log_2 \mathrm{FC}|$ remains small at $1.04 \times 10^{-3}$. Per-(cell, gene) aggregation across co-localising SNVs produces a maximum cumulative $|\sum \log_2 \mathrm{FC}|$ of *exactly* 2.248 — equal to the single-SNV maximum — ruling out the multi-hit additive failure mode at the level of an individual gene: when two random SNVs land in the same gene in the same cell, they do not stack to a larger effect than the largest single hit. Across the whole-cell 4,000-SNV burden, the per-cell cumulative $\sum \log_2 \mathrm{FC}$ summed across all assigned genes has empirical mean −0.149 and standard deviation 0.862 (max $|\sum| = 2.55$ over 60 cells), substantially wider than the $\sigma$-scaled extrapolation from the 100-SNV experiment of R4 had projected — the per-variant distribution is heavy-tailed, so Gaussian $\sqrt{n}$-scaling underestimates the 4,000-SNV tail. However, this whole-cell aggregate is summed across thousands of distinct genes; the biologically meaningful per-gene comparison against aging differential expression remains bounded $\sim 3\times$ below the TSP aging maximum (2.248 vs 6.98). The honest revision of the headline argument is therefore: occasional single random SNVs ($\sim$ 1 in 20,000) reach aging-magnitude $|\log_2 \mathrm{FC}| \geq 1$ in their nearest gene; multi-hit additive stacking within one gene is empirically bounded; and the bulk per-variant distribution sits three orders of magnitude below the aging median.

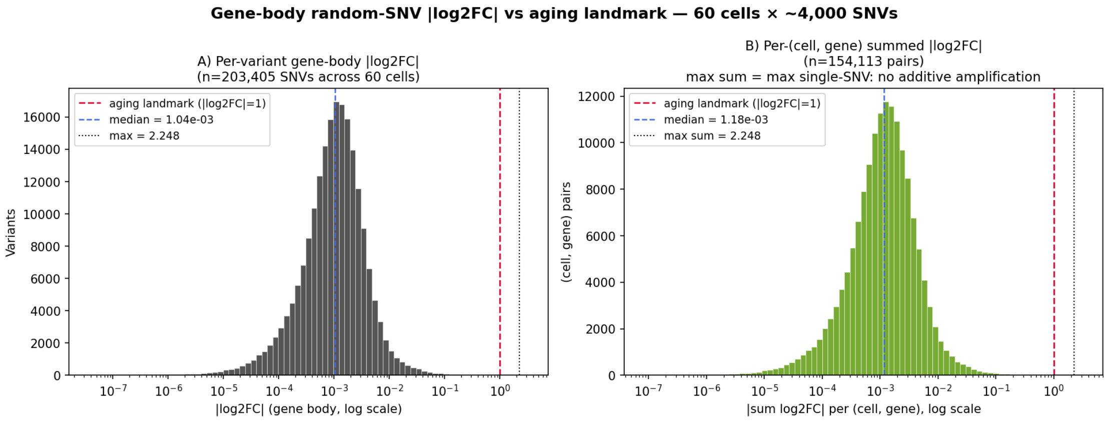


**Figure 4: Gene-body random-SNV $|\log_2 \mathbf{FC}|$ distribution against the 2-fold-change threshold $|\log_2 \mathbf{FC}| = 1$.** The reference line at $|\log_2 \mathrm{FC}| = 1$ marks a 2-fold expression change — a conventional biological-significance threshold cleared by thousands of aging DEGs in the TSP large-intestine reference (range 0–6.98, median 0.572); see Methods. (*Left*) Per-variant $|\log_2 \mathrm{FC}|$ histogram on log axis across 203,405 random SNVs from 60 simulated cells; vertical lines mark the 2-fold threshold, the median, and the empirical maximum (*TMIGD2*, 2.248). Ten SNVs reach $\geq 1$ ($\sim$ 1 in 20,000); 157 reach $\geq 0.138$. (*Right*) Per-(cell, gene) summed $|\log_2 \mathrm{FC}|$ across co-localising SNVs in the same cell; the maximum sum equals the single-SNV maximum, ruling out additive multi-hit amplification at the per-gene level. *Species: human* (hg38 SNVs scored in human colon, UBERON:0001157).

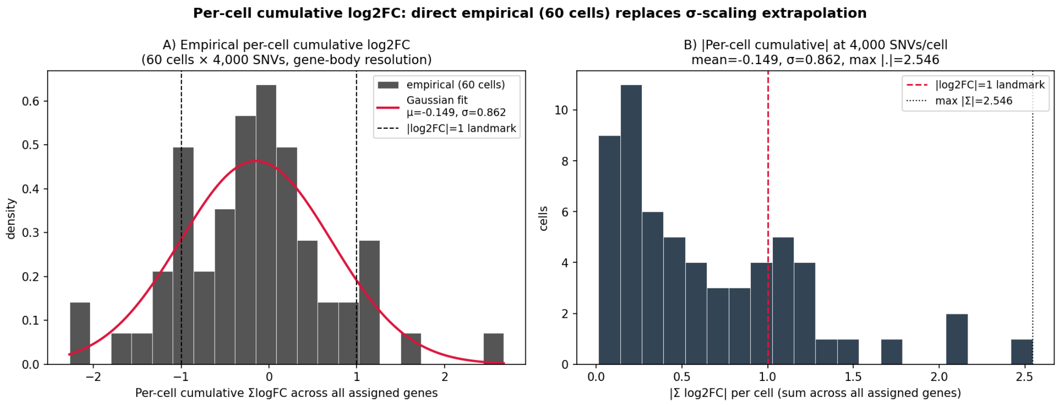


**Figure 5: Per-cell cumulative $\log_2$FC distribution at 4,000 SNVs/cell — direct empirical measurement (60 cells) replaces the prior $\sigma$-scaled Gaussian extrapolation.** (*Left*) Empirical histogram of per-cell $\sum \log_2 \mathrm{FC}$ summed across all assigned genes (60 cells $\times \sim 4{,}000$ SNVs each), with fitted Gaussian (mean $-0.149$, $\sigma = 0.862$). The empirical spread is much wider than the $\sigma$-scaled extrapolation from the 100-SNV experiment of R4 had projected — the per-variant gene-body distribution is heavy-tailed, so $\sqrt{n}$-scaling underestimates the 4,000-SNV tail. (*Right*) $|\sum \log_2 \mathrm{FC}|$ per cell against the 2-fold-change threshold $|\log_2 \mathrm{FC}| = 1$; max $|\sum| = 2.55$. This whole-cell aggregate is summed across thousands of distinct genes and is therefore not directly comparable to per-gene aging differential expression — the per-gene comparison (left panel of Figure 4) is the load-bearing one and remains bounded $\sim 3\times$ below the TSP aging maximum. *Species: human*.

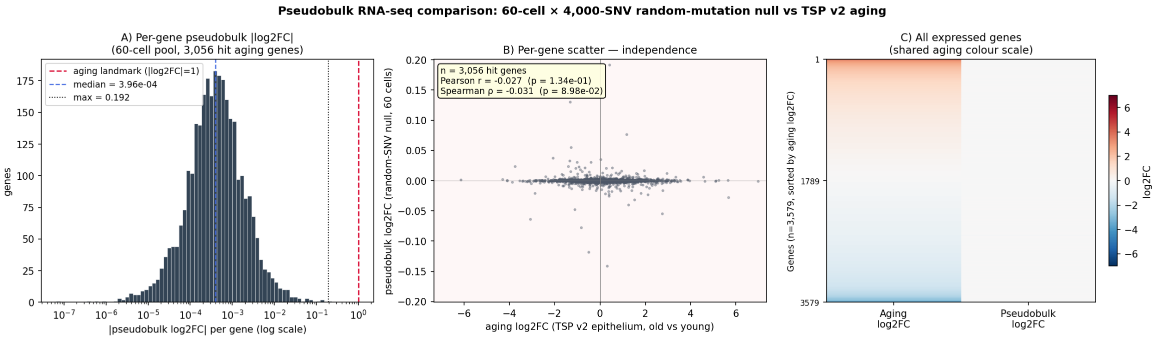


**Figure 6: Pseudobulk RNA-seq comparison: 60-cell $\times$ 4,000-SNV random-mutation null versus Tabula Sapiens v2 aging in colonic epithelium.** (*A*) Per-gene pseudobulk $|\log_2 \mathrm{FC}^{\mathrm{pb}}|$ histogram across 3,056 hit aging genes, on log axis, against the 2-fold-change threshold $|\log_2 \mathrm{FC}| = 1$. Median over hit genes $= 3.96 \times 10^{-4}$, max $= 0.192$ (*TMEM238*). (*B*) Per-gene scatter of aging $\log_2$FC versus pseudobulk $\log_2$FC over the same 3,056 genes. Pearson $r = -0.027$ ($p = 0.13$); Spearman $\rho = -0.031$ ($p = 0.09$). The pseudobulk axis spans approximately $[-0.2, +0.2]$ while the aging axis spans approximately $[-7, +7]$ — the two are independent both in magnitude and direction. (*C*) Side-by-side per-gene heatmap of all 3,579 expressed genes sorted by aging $\log_2$FC, with the pseudobulk column on the same diverging colour scale as the aging column; the pseudobulk column is uniformly neutral, mirroring the random-SNV-vs-aging picture of Figure 2 but at the considerably more demanding pseudobulk granularity (a per-gene aggregate over multiple cells, not a single-variant assignment). *Species: human* (hg38 SNVs scored in human colon, UBERON:0001157; TSP v2 large-intestine aging reference).

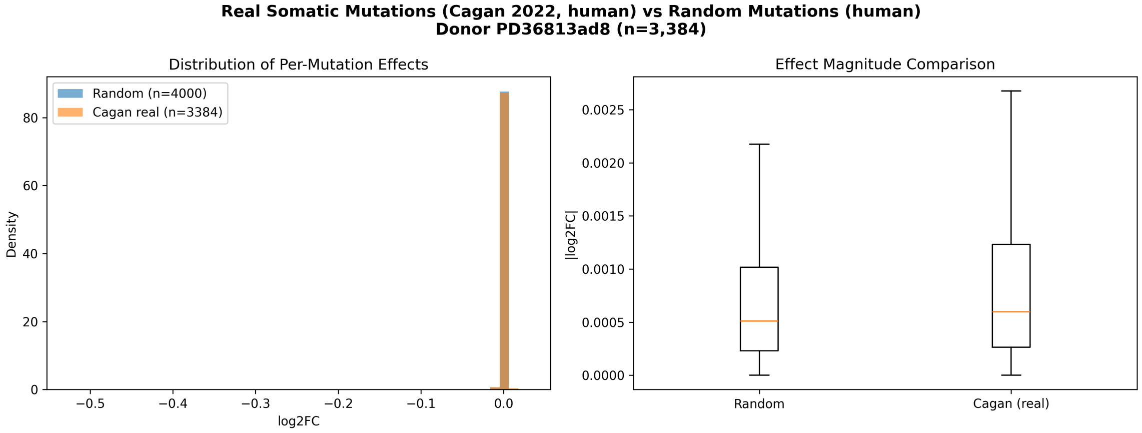


**(a)** Two-panel comparison of per-variant effect distributions (human donor PD36813ad8, $n = 3{,}384$ SNVs vs a matched random baseline). *Top left*: histograms of `mean_expression_change`; Cagan and random distributions are nearly coincident. *Top right*: `max_expression_change` overlays; no heavier right tail for real mutations. *Bottom left*: box plots of absolute mean expression change; the Cagan box sits at or below the random box. *Bottom right*: cumulative effect as mutations are added one at a time; both traces are small random walks around zero.

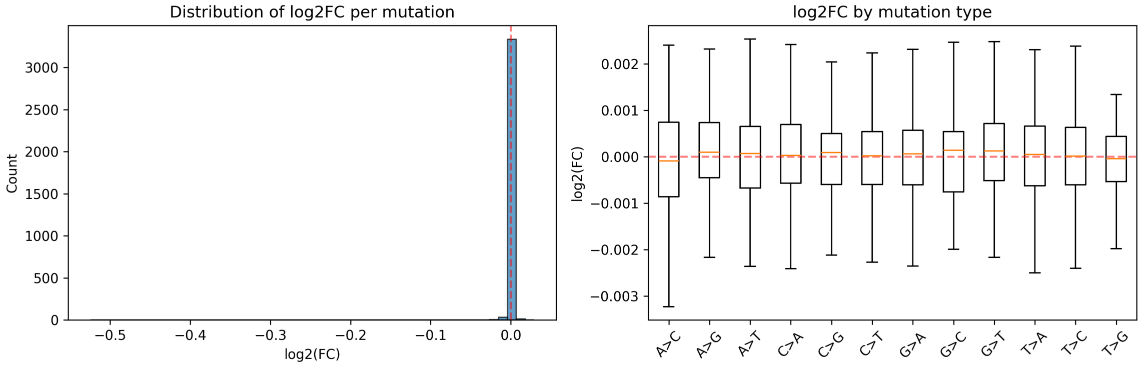


**(b)** Per-variant $\log_2$FC distribution (same human sample). Median $|\log_2 \mathrm{FC}| = 5.9 \times 10^{-4}$, consistent with the random baseline and three orders of magnitude below the 2-fold-change threshold $|\log_2 \mathrm{FC}| = 1$.

**Figure 7: Real somatic SNVs from aged human colonic crypts (Cagan 2022, $n = 3$ donors) produce per-variant AlphaGenome effects smaller than random genome-wide SNVs.** All panels computed in colon tissue (UBERON:0001157). *Species: human.* See Supplementary Figure S5 for the matched mouse-cohort figure ($n = 54$ crypts, used here as the larger-sample-size confirmation); conclusions are identical.

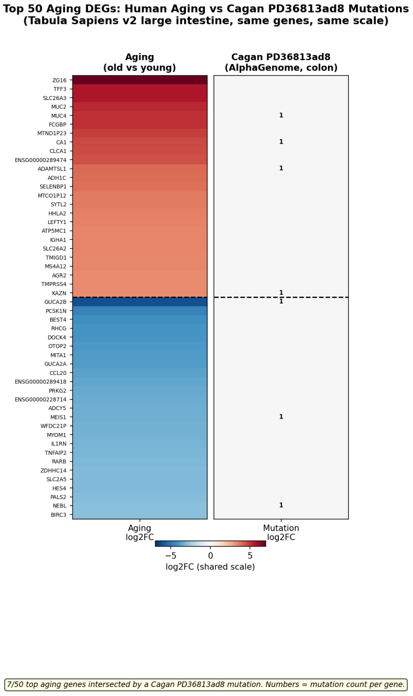


(a) Top 25 up- + top 25 down-regulated aging DEGs.

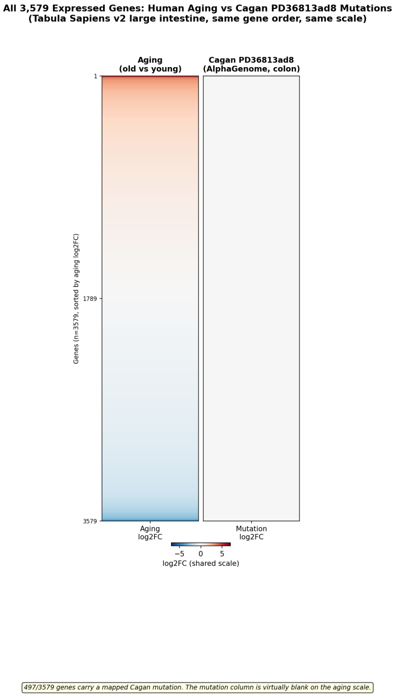


(b) All 3,579 expressed genes, sorted by aging $\log_2$FC.

**Figure 8: Real Cagan human-cohort somatic SNVs (donor PD36813ad8) produce per-gene effects that are invisible on the aging $\log_2$FC colour scale in human large-intestine tissue.** *Species: human* (Tabula Sapiens v2 aging reference; AlphaGenome scoring of 3,384 Cagan PD36813ad8 SNVs in human colon, UBERON:0001157, assigned to the nearest GENCODE v46 protein-coding gene body). Both panels share the diverging colour map and aging-scale colour range used in Figure 2, allowing the two figures to be read side by side. (*Left, a*) The top 25 up- and 25 down-regulated aging DEGs from Tabula Sapiens v2 large-intestine epithelium (young donor TSP26, 37 y vs old donors TSP2/TSP14/TSP25/TSP27, 56–61 y); only 7 of 50 top aging genes are intersected by a Cagan SNV in this donor, and the per-gene mutation column is uniformly neutral. Small numerals mark the mutation count per gene. (*Right, b*) All 3,579 expressed colonic-epithelial genes ordered by aging $\log_2$FC, with 497 (14%) carrying a mapped Cagan mutation but no visible colour on the shared scale. Per-gene maximum mutation $|\log_2 \text{FC}| = 0.525$ and median $5.98 \times 10^{-4}$ versus aging max 6.98 and median 0.572, giving a max-effect ratio of $\sim 13\times$ and a median ratio of $\sim 960\times$ on real Cagan human SNVs — consistent with the random-SNV picture of Figure 2 and confirming that the somatic catalogue does not concentrate at aging-relevant loci.

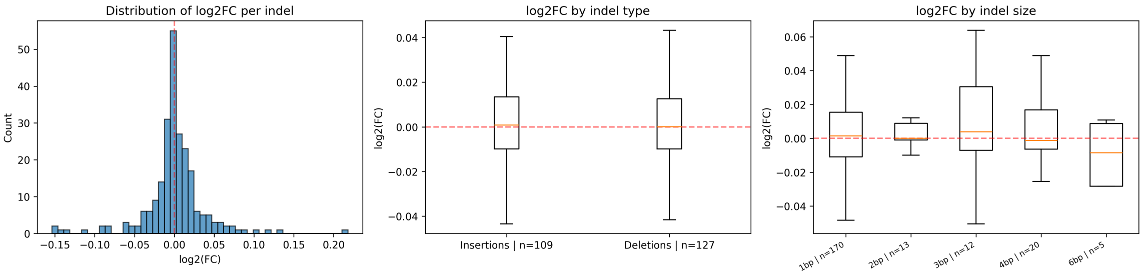


**(a)** Per-indel $\log_2$FC distribution for exemplar donor PD36813ac4 ($n = 236$): overall histogram, broken down by indel type (insertion vs deletion, near-identical) and by indel size (1–6 bp, no monotonic size–effect relationship). The bulk of the distribution sits within $\pm 0.05$.

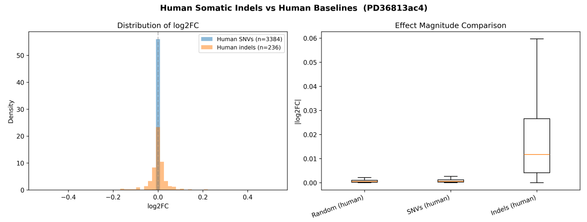


**(b)** PD36813ac4 indels vs. random hg38 SNVs and matched-donor SNV cohort: indels exceed both at every quantile but remain in the same order of magnitude, with the bulk concentrated around the median.

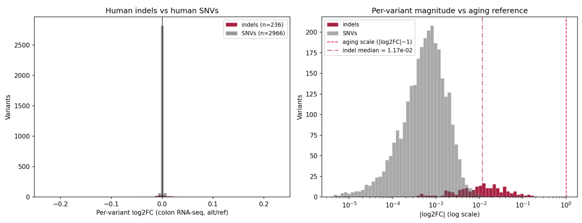


**(c)** PD36813ac4 indels vs SNVs on a shared log axis with the 2-fold-change threshold $|\log_2 \mathrm{FC}| = 1$ clearly above all observed events. Per-indel median $= 1.17 \times 10^{-2}$, two orders of magnitude below the 2-fold threshold.

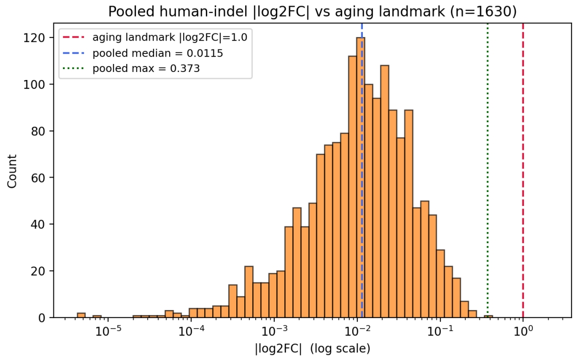


**(d)** Pooled 9-donor $|\log_2 \mathrm{FC}|$ histogram ($n = 1{,}630$ indels) on log axis: the entire pooled distribution sits below 0.4 with median 0.0115 and max 0.373 — well below the 2-fold-change threshold.

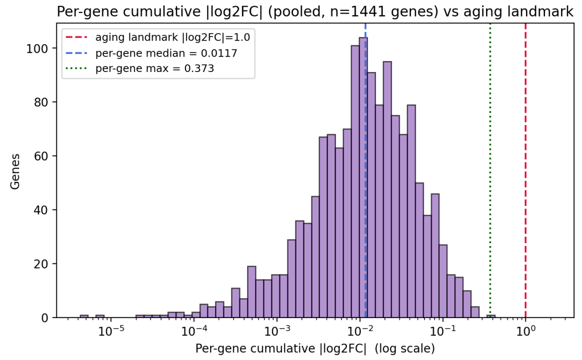


**(e)** Per-gene cumulative $|\log_2 \mathrm{FC}|$ over 1,441 unique genes hit by pooled indels (additive sum across all donors at each gene). Summing across all 9 donors does *not* push any single gene past the 2-fold-change threshold: per-gene max 0.373; 0 of 1,441 genes reach $|\sum \log_2 \mathrm{FC}| \geq 1$.

**Figure 9: Human-cohort somatic indels (Cagan 2022) produce per-event AlphaGenome effects above SNVs but two orders of magnitude below the 2-fold-change threshold $|\log_2 \mathbf{FC}| = 1$, both for the exemplar donor PD36813ac4 and across the pooled 9-donor cohort ($n = 1{,}630$ indels).** Per-donor indel/SNV per-event ratios are 22, 21, and 24× for PD36813ac4, PD37266c_lo0016, and PD37590b_lo0090 respectively, closely tracking the $\approx$ 24.4× ratio observed in the mouse cohort (Supplementary Figure S7) and confirming that the structural-variation amplification effect is reproducible across species; the gap to the 2-fold-change threshold on the median is $\approx$ 49–91× across donors, consistent with the species-matched scale gap reported for human SNVs in Figure 7. (a) Per-indel distribution + insertion/deletion + size breakdown for PD36813ac4. (b) Indel vs random and SNV baselines for PD36813ac4. (c) Indels vs SNVs on a shared log axis with the 2-fold-change threshold. (d) Pooled 9-donor $|\log_2 \mathrm{FC}|$ histogram against the 2-fold-change threshold. (e) Per-gene cumulative $|\log_2 \mathrm{FC}|$ histogram across all genes hit. *Species: human* (Cagan *et al.* 2022, 9 human colonic-crypt donors; AlphaGenome RNA-seq, colon UBERON:0001157).

#### R9. Somatic mutations are non-randomly distributed, with a purifying-selection signature

If somatic variation experienced no selection, the positional distribution of mutations would match the genomic background in proportion to region size. We tested this on the human Cagan cohort using GENCODE v46 hg38 annotations across 28 colonic crypt samples (3,042 indels). The top-level region distribution deviates from the bp-proportional null at $\chi^2\, p = 4.4 \times 10^{-2}$, and within protein-coding gene bodies the deviation is much stronger: $\chi^2\, p = 2.4 \times 10^{-5}$, driven primarily by depletion of coding sequence. Of 1,283 protein-coding-gene indels, only 15 fell within CDS against an expectation of 35.1 ($\log_2$(obs/exp) $= -1.23$, $\sim 2.3\times$ depleted; Figure 10). UTRs were near-neutral ($\log_2 = -0.09$), and introns dominated protein-coding-gene hits ($\sim 92.5\%$, near the 91.7% expectation). The matched 54-sample mouse cohort (9,799 indels, GENCODE vM25 mm10) reproduces the same qualitative picture with much higher statistical power, owing to the $\sim 3\times$ larger sample size: Strikingly, CDS is depleted $12\times$ ($\log_2 = -3.58$, only 8 of 2,755 protein-coding-gene indels in CDS), and the top-level $\chi^2\, p = 3.2 \times 10^{-104}$ (Supplementary Figure S8). Both species therefore show the canonical purifying-selection signature — CDS depletion paired with intergenic neutrality or mild enrichment — consistent with crypt clones that acquire coding-disruptive variants being eliminated before they can be sampled, *not* with neutral accumulation at random positions.

The same test applied to the much larger Cagan SNV cohorts (56,123 human SNVs across 28 samples; 54,158 mouse SNVs across 54 samples) reproduces the direction of the indel signature but with substantially smaller magnitudes per event, exactly as expected biologically: most coding-region SNVs are synonymous or conservative missense and therefore escape the strong purifying filter that frameshift indels face. Within human protein-coding genes, CDS is $\sim 1.4\times$ depleted ($\log_2 = -0.52$, 427 of expected 611 hits, within-PC $\chi^2\, p = 9.8 \times 10^{-22}$; Figure 11); the mouse cohort shows the same direction at $\sim 1.3\times$ ($\log_2 = -0.34$, 460 of expected 583, within-PC $\chi^2\, p = 1.0 \times 10^{-15}$; Supplementary Figure S9). The top-level $\chi^2$ p-values ($5.7 \times 10^{-51}$ human, $4.2 \times 10^{-290}$ mouse) reflect the very large SNV sample sizes; the biologically meaningful quantity is the modest log2 effect size, which is roughly an order of magnitude smaller than the indel result — consistent with SNVs as a class being more permissive substrates for clonal survival than length-altering indels, while still subject to coherent selection against coding disruption.

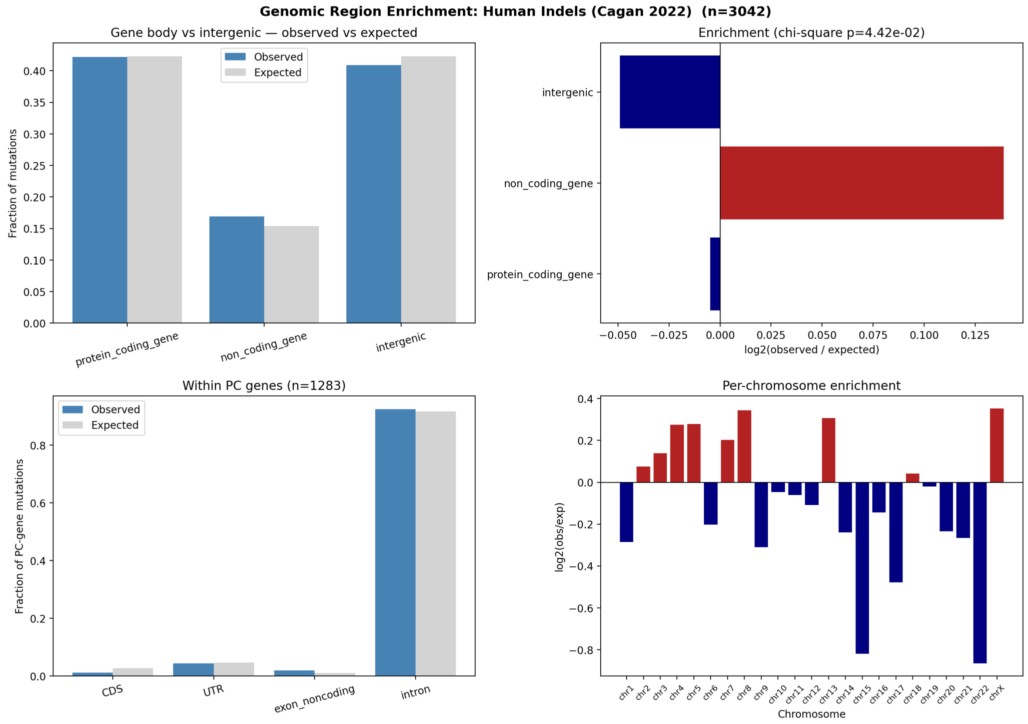


**Figure 10: Human Cagan somatic indels are non-randomly distributed across genomic regions.** Observed vs. expected indel counts across five region classes (CDS, UTR, noncoding exon, intron, intergenic), pooled across 28 human samples. Bar heights show observed (darker) and expected (lighter, bp-proportional) counts; inset values give $\log_2$(obs/exp). CDS is $\sim 2.3\times$ depleted ($\log_2 = -1.23$, 15 of expected 35.1 hits), the strongest signal; introns are near-neutral; non-coding exons are mildly enriched. $\chi^2$ goodness-of-fit: $p = 4.4 \times 10^{-2}$ at the top level, $p = 2.4 \times 10^{-5}$ within protein-coding genes. This is the canonical purifying-selection signature: crypt clones that acquire coding-disruptive variants are eliminated before they can be sampled. The matched 54-sample mouse cohort confirms the same picture with much sharper magnitudes ($12\times$ CDS depletion, $\chi^2\, p = 3.2 \times 10^{-104}$; Supplementary Figure S8). Logical implication: the *observed* somatic catalogue has already passed a selection filter, so its molecular effects represent an upper bound for neutral MA dynamics — and those effects are still negligible (R7, R8). *Species: human* (Cagan *et al.* 2022, 28 human colonic crypt samples, 3,042 indels; GENCODE v46 hg38 annotations).

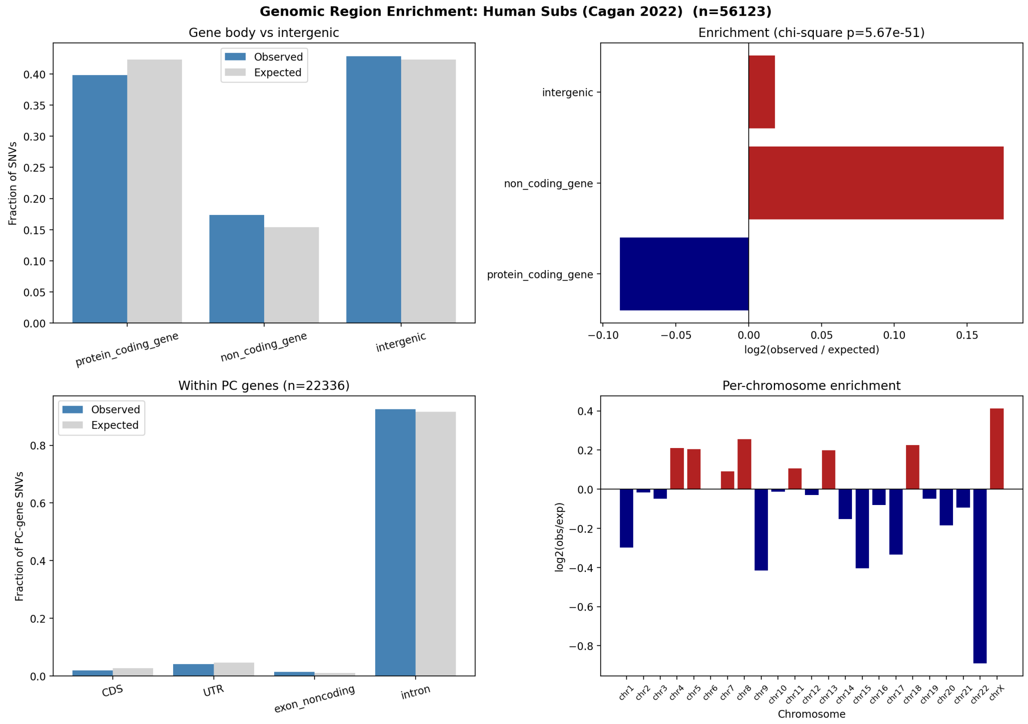


**Figure 11: Human Cagan somatic SNVs reproduce the indel purifying-selection signature in the same direction but at much smaller per-event magnitude.** Observed vs. expected SNV counts across the same five region classes, pooled across 28 human samples (56,123 SNVs total). CDS is $\sim 1.4\times$ depleted ($\log_2 = -0.52$, 427 of expected 611 hits); UTRs near-neutral; introns near-neutral; non-coding exons modestly enriched. $\chi^2$ goodness-of-fit: $p = 5.7 \times 10^{-51}$ at the top level, $p = 9.8 \times 10^{-22}$ within protein-coding genes. The much smaller log2 magnitude relative to the indel result of Figure 10 is biologically expected — most coding SNVs are synonymous or conservative missense and escape the strong selection that frameshift indels face — but the direction is preserved, confirming that even SNVs in surviving aged-crypt clones have been filtered against coding disruption. *Species: human* (Cagan *et al.* 2022, 28 human colonic crypt samples; GENCODE v46 hg38 annotations).

#### R10. Recurrently mutated genes are common fragile sites, not aging-related loci

We asked whether specific genes attract more somatic mutations than expected given their length plus a ±10 kb flank. Per-gene Poisson upper-tail tests with Benjamini–Hochberg FDR correction identified 99 protein-coding genes significantly enriched for indels across 54 samples at FDR $< 0.05$, and 41 genes for substitutions across 5 samples (Figure 12a,b). The top hits were overwhelmingly classic common fragile site (CFS) and long neural-adhesion genes: *Cntnap2* (34 indel hits, 23/54 samples), *Lsamp*, *Dmd*, *Lrp1b*, *Lrrc4c*, *Csmd1/3*, *Pcdh9/11x/15*, *Cdh12/18*, *Nrxn3*, *Dcc*, *Rbfox1*, *Dpp10*, *Cadm2*, *Sgcz*, *Kcnt2*. SNV and indel hits strongly overlapped at the same loci (hypergeometric $p = 3.8 \times 10^{-37}$; Figure 12b), indicating shared underlying fragility rather than mechanism-specific selection. These are neural genes not highly expressed in colonic epithelium, ruling out transcription-coupled mutagenesis of active loci; their shared property is extreme length (multi-megabase) and late replication, both hallmarks of CFS. Critically, hypergeometric overlap against five aging-associated gene sets revealed *no* enrichment for fibrosis, SASP, senescence, inflammatory, or EMT genes in either the indel or SNV hit list (all fold $< 1$, all $p > 0.8$; Figure 12c,d).

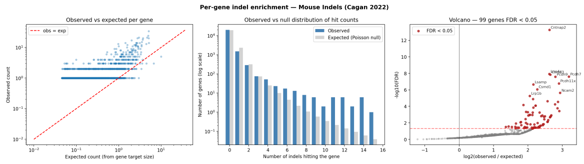


**(a)** Per-gene indel enrichment volcano: $\log_2$(obs/exp) on x-axis, $-\log_{10}$(FDR) on y-axis, across 54 samples. 99 protein-coding genes reach FDR$<0.05$ (above horizontal dashed line); top-labelled genes are dominated by CFS/long-gene loci. The horizontal banding reflects the discrete observed counts for a Poisson test.

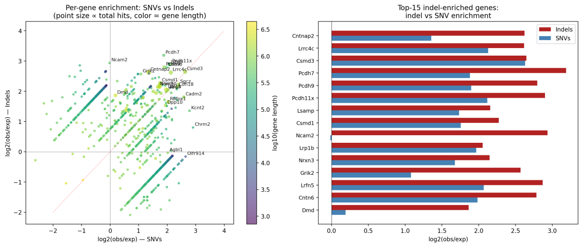


**(b)** Per-gene SNV vs. indel $\log_2$ enrichment scatter, with point size proportional to total hits and color-coded by gene length (log scale). The top-right quadrant is densely populated with labelled CFS genes (e.g., *Lrp1b*, *Lsamp*, *Csmd3*, *Cadm2*, *Rbfox1*, *Pcdh9/11x/15*), showing shared enrichment across variant classes. Hypergeometric overlap test for top-hit gene lists: $p = 3.8 \times 10^{-37}$.

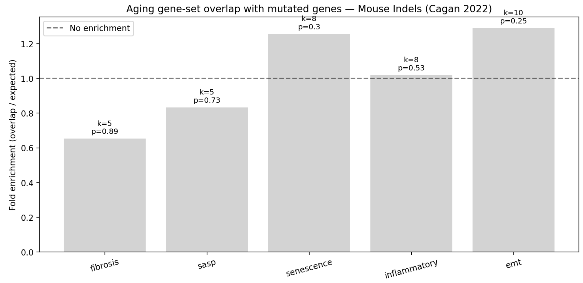


**(c)** Hypergeometric overlap bar chart for recurrently-hit indel genes (FDR$<0.05$) vs. five aging-associated gene sets (fibrosis, SASP, senescence, inflammatory, EMT). Bars show fold-enrichment with annotated $p$-values. None reach $p < 0.05$; all folds $< 1$ (depletion or null).

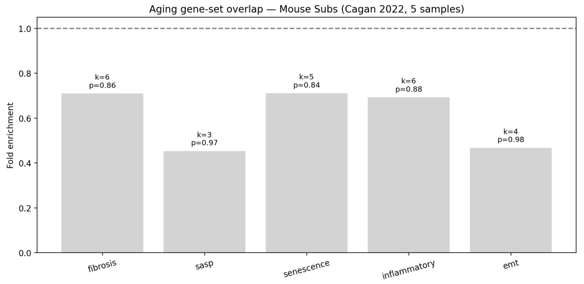


**(d)** Same hypergeometric test applied to the SNV hit list from the 5-sample cohort. Again no aging gene set is enriched; all $p > 0.8$.

**Figure 12: Recurrently mutated genes are common fragile sites, not aging-relevant loci.** Four panels show (a) the per-gene Poisson volcano for indels across 54 samples, (b) the per-gene SNV–indel agreement scatter, and (c,d) hypergeometric tests of the hit lists against five aging-associated gene sets. The top 20 recurrent hits — *Cntnap2*, *Lsamp*, *Dmd*, *Lrp1b*, *Lrrc4c*, *Csmd1/3*, *Pcdh9/11x/15*, *Cdh12/18*, *Nrxn3*, *Dcc*, *Rbfox1*, *Dpp10*, *Cadm2*, *Sgcz*, *Kcnt2* — are overwhelmingly long neural-adhesion CFS genes not expressed in colonic epithelium, arguing for replication-stress fragility rather than transcription-coupled mutagenesis or functional selection. The shared SNV–indel enrichment ($p = 3.8 \times 10^{-37}$) confirms common underlying fragility. No aging-associated gene set (fibrosis, SASP, senescence, inflammatory, EMT) is enriched in either hit list. *Species: mouse* (Cagan *et al.* 2022, 54 mouse samples for indels and 5 samples for SNVs; the human Cagan cohort of 3 donors is under-powered for per-gene recurrence testing).

#### R11. Mutation-induced expression effects sit ∼3–550× below the tissue's own aging program (max-effect ratio $\sim 3\times$, median $\sim 550\times$)

To place the preceding effect sizes on an absolute biological scale, we compare predicted mutation-induced $|\log_2 \mathrm{FC}|$ at gene-body resolution (R4b: 203,405 random SNVs across 60 simulated cells in human colon, UBERON:0001157) against the human Tabula Sapiens v2 colonic-epithelium aging reference [4, 5] (young donor TSP26, 37 y, 597 cells vs old donors TSP2/TSP14/TSP25/TSP27, 56–61 y, 8,981 cells). Aging-associated differential expression spans $|\log_2 \mathrm{FC}|$ from 0 to 6.98 (median 0.572), with thousands of genes above $|\log_2 \mathrm{FC}| = 1$. Predicted mutation-induced $|\log_2 \mathrm{FC}|$ at gene-body resolution spans 0 to 2.248 (median $1.04 \times 10^{-3}$). The maximum-effect ratio is therefore $\sim 3\times$ — substantially smaller than the $\sim 840\times$ figure obtained under the dilution-prone window-level scoring used in earlier drafts of this manuscript — but the median ratio remains $\sim 550\times$ because the bulk of random SNVs produce sub-noise effects (Figures 2 and 4). The qualitative conclusion is preserved and sharpened: a single random SNV occasionally reaches aging magnitude in *one* gene, but the bulk distribution sits three orders of magnitude below the aging program, which simultaneously affects *thousands* of genes. The matched mouse-on-mouse version of this analysis (Cagan sample MD6267ab_lo0015 vs Tabula Muris Senis large intestine) is shown in Supplementary Figure S6: at the lower-resolution window-level scoring used there, the mouse aging-to-mutation ratio is $\sim 840\times$ on the maximum and $\sim 990\times$ on the median.

### Methods

#### AlphaGenome predictions

All mutation scoring used the AlphaGenome `dna_client.predict_variant` API [2] with a 1 Mb sequence interval centered on the variant. Predictions requested `RNA_SEQ`, `ATAC`, and `CHIP_HISTONE` output types in a single call (three-in-one to avoid quota multiplication; each call $\approx$ 2–3 s). For the multi-output analysis (R3) we additionally requested `DNASE` and `CAGE`. Tissue ontologies: colon UBERON:0001157, large intestine UBERON:0000059, brain UBERON:0000955, liver UBERON:0002107, lung UBERON:0002048, heart UBERON:0000948, kidney UBERON:0002113, muscle UBERON:0001134, skin UBERON:0002097. For histone ChIP outputs, available tracks were detected dynamically per prediction (not hard-coded), since the mark repertoire differs by tissue (e.g., liver lacks H3K27me3 while lung has H3K27ac). Per-variant effect metrics included mean/max/total expression change over the 8,192-bp output window, as well as ATAC and per-histone-mark mean fold change in a 10-kb central window, with `nan` propagation when outputs were unavailable.

#### Random-variant baseline (R1)

Four-thousand SNVs were sampled uniformly without replacement from hg38 autosomal and X/Y sequence, with reference/alt alleles drawn to exclude `N` bases. Each was scored in colon tissue with the full three-output predictor, yielding `colon_mutation_simulation_results.csv` (columns `chromosome, position, ref, alt, mean_expression_change, max_expression_change, total_expressi` [illegible] `success, error`; 100% success).

#### Combined 1-Mb window analysis (R2)

The 4,000 random variants were binned into non-overlapping 1-Mb genomic windows. For each window we computed: (i) the variant count, (ii) the per-variant absolute mean expression change for the window's largest variant, and (iii) the window cumulative expected effect, defined as the sum of per-variant mean expression changes across all variants in the window (additive-expectation upper bound). Windows with $\geq 2$ variants were used to compute combined-vs-additive plots (Figure 3).

#### Monte Carlo cohort (R4)

We simulated 100 independent cells, each accumulating 100 random SNVs drawn without replacement from hg38 (excluding `N` positions). Each SNV was scored with AlphaGenome in colon tissue (RNA-seq output); per-cell cumulative effect was defined as the sum of per-variant mean expression changes across the cell's 100 variants. The resulting 100 per-cell cumulative effects were summarized by their empirical mean, standard deviation, and observed maximum, and fit with a normal distribution. Results are summarized in `monte_carlo_summary.json`. The R4b gene-body run (60 cells $\times \sim$ 4,000 SNVs) supersedes any $\sigma$-scaling extrapolation that this 100-SNV cohort would license, by directly measuring per-cell cumulative effects at the realistic per-cell SNV burden.

#### Cagan *et al.* 2022 somatic mutation data

Substitution and indel call files were obtained from Zenodo deposit `10.5281/zenodo.5554777` (archive `CrossSpecies2021_Data.tar.gz`), corresponding to Cagan *et al.* (2022) [1]. Mouse files (54 samples) reside under `FinalCalls_Subs/mouse` and `FinalCalls_Indels/mouse`; human files (3 samples) under the corresponding human subdirectory. Substitution files have columns `Chr, Start, End, Ref, Alt, ...`; indel files use `Chrom, Pos, Ref, Alt, ...`. Mouse variants were scored natively in mm10 via the AlphaGenome Python client with `Organism.MUS_MUSCULUS`; no liftover was applied to Cagan calls.

#### Genomic region enrichment (R9)

GENCODE vM25 (mouse) and v46 (human) gene, transcript, and exon features were parsed and collapsed by `merge_intervals()` to obtain non-overlapping base-pair coverage per region class (CDS, UTR, noncoding exon, intron, intergenic). Each mutation was classified by its mm10 position relative to these merged intervals. A chi-square goodness-of-fit test compared observed region counts against the expected distribution $N_{\text{total}} \cdot (L_{\text{region}}/L_{\text{kept genome}})$, where $L_{\text{kept genome}}$ sums chromosome lengths of chromosomes actually represented in the mutation calls (fixing a prior mismatch between numerator and denominator). Per-region $\log_2$ enrichment was reported as $\log_2((\text{obs} + 0.5)/(\text{exp} + 0.5))$. Distance-to-TSS analysis used nearest protein-coding TSS in a $\pm 50$ kb window.

#### Per-gene enrichment (R10)

For each protein-coding gene on accepted chromosomes we defined a target window of the gene body $\pm 10$ kb flank. Each mutation was assigned to a single gene via a nearest-gene rule: a mutation inside multiple overlapping target windows was placed in the gene whose *body* contained it, with ties broken by distance; otherwise to the nearest target-window boundary. Per-gene observed counts were tested under a Poisson upper-tail null with expected $\lambda_i = N \cdot L_i^{\text{target}} / \sum_j L_j^{\text{target}}$, and $p$-values BH-corrected over the subset of genes with obs $\geq 1$. Gene-level recurrence was defined as `n_samples` $\geq 3$ and FDR $< 0.05$.

#### Aging gene-set overlap

Five aging-associated gene sets — fibrosis, SASP, senescence, inflammatory, EMT — were taken from `gene_set_susceptibility_analysis/gene_sets/`. Human symbols were converted to mouse case (title-cased) for mouse comparisons. Hypergeometric tests (scipy) compared the overlap of FDR$<0.05$ recurrent hit genes against each gene set, using the set of all assayed protein-coding genes as the universe.

The reference line at $|\log_2 \text{FC}| = 1$ used in figures throughout the manuscript denotes the conventional 2-fold-change cutoff (since $\log_2 2 = 1$), not a derived quantile of the aging distribution; it is used as a fixed magnitude reference because thousands of aging DEGs in both atlases clear it (Tabula Sapiens v2 colonic epithelium: $|\log_2 \text{FC}|$ range 0–6.98, median 0.572; Tabula Muris Senis: range 0–10.74, median 0.807). Where the actual aging maximum and median are needed for quantitative ratio calculations, those data-derived values are reported alongside.

Per-gene mutation aggregation proceeded as in R10: each SNV was assigned to the protein-coding gene whose body contains the position (genic), or to the nearest protein-coding gene by body-boundary distance (intergenic), using GENCODE v46 (hg38) for R1 and GENCODE vM25 (mm10) for R11. Per-mutation `mean_expression_change` (R1) or per-mutation $\log_2$FC (R11) was summed per gene. The per-gene mutation sum was joined to the aging table by human symbol in R1 and by mouse symbol in R11.

The predictive framework introduced here generalizes naturally to those candidate mechanisms. AlphaGenome's simultaneous multi-modal output — chromatin accessibility, histone modifications, TSS usage, RNA abundance — makes it well-suited to scoring *epigenetic* perturbations rather than sequence-level ones, and its cross-modal coupling is already demonstrable at ISM hotspots (R6). Natural extensions are: applying the exact same null/real-catalogue structure to post-mitotic brain (where MA is most thermodynamically favourable and thus most easily falsified); scoring the ISM landscape across tissue-specific aging-associated ATAC peaks; and coupling AlphaGenome variant scoring with chromatin-age clocks to ask, quantitatively, whether predicted epigenetic drift at aging-ATAC regions can recapitulate the transcriptional signature that 10,000 random mutations demonstrably cannot.

### Supplementary Figures

**Table S1: Mean $|\Sigma \log_2 \mathrm{FC}|$ across each Cagan / random-SNV mutation experiment computed over *all* 17,226 expressed colonic-epithelial genes (TMS), filling 0 for genes not assigned a mutation.** Mouse counterpart of main-text Table 1. Per-donor mean averaged across donors for Cagan rows; pooled experiment for the random row; aging row shows mean $|\log_2 \mathrm{FC}|$ over the same expressed-gene set. Per-donor hit-fractions shown in column 2. Ratio columns are aging value ÷ mutation value. *Species: mouse.*

| Dataset | hit-fraction (per donor) | mean $|\log_2 \mathrm{FC}|$ | max $|\log_2 \mathrm{FC}|$ | aging-mean ratio | aging-max ratio |
|---|---|---|---|---|---|
| Aging (TMS) | — | **1.541** | **10.74** | 1× | 1× |
| Random SNVs (mm10) | 7.7% | $8.8 \times 10^{-5}$ | 0.025 | 17,427× | 432× |
| Cagan SNVs (5 donors) | 3.9% | $5.5 \times 10^{-5}$ | 0.194 | 27,898× | 55× |
| Cagan indels (8 donors) | 1.2% | $3.3 \times 10^{-4}$ | 0.350 | 4,624× | 31× |

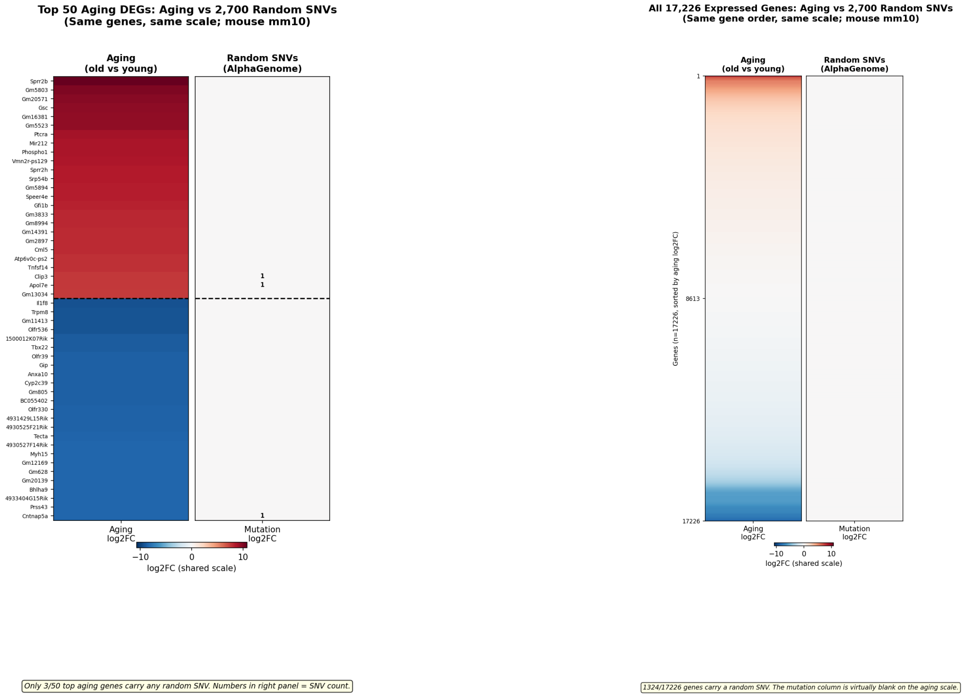


**(a)** Top 25 up- + top 25 down-regulated aging DEGs. **(b)** All 17,226 expressed genes, sorted by aging $\log_2$FC.

**Figure S1: Mouse mm10 mirror of main-text Figure 2: 2,700 random mm10 SNVs produce per-gene effects that are invisible on the aging $\log_2$FC colour scale in mouse large-intestine epithelium.** Both panels share the diverging aging colour map. (*Left, a*) Top 25 up- and 25 down-regulated aging DEGs from Tabula Muris Senis large-intestine epithelium (3 m young vs 18/24/30 m old; 9,263 epithelial cells); only 3 of the 50 top aging genes are intersected by any random SNV, and the per-gene mutation column is uniformly neutral. Small numerals mark the SNV count per gene. (*Right, b*) All 17,226 expressed colonic-epithelial genes ordered by aging $\log_2$FC, with 1,324 ($\sim$ 8%) carrying a mapped random SNV but no visible colour on the shared scale. *Cell-type filter:* the "colon-epithelial" aging reference pools four Cell Ontology cell types (enterocyte of epithelium of large intestine, epithelial cell of large intestine, intestinal crypt stem cell, large intestine goblet cell), not a single cell type. *Species: mouse* (mm10 SNVs scored in mouse colon, UBERON:0000059; Tabula Muris Senis aging reference).

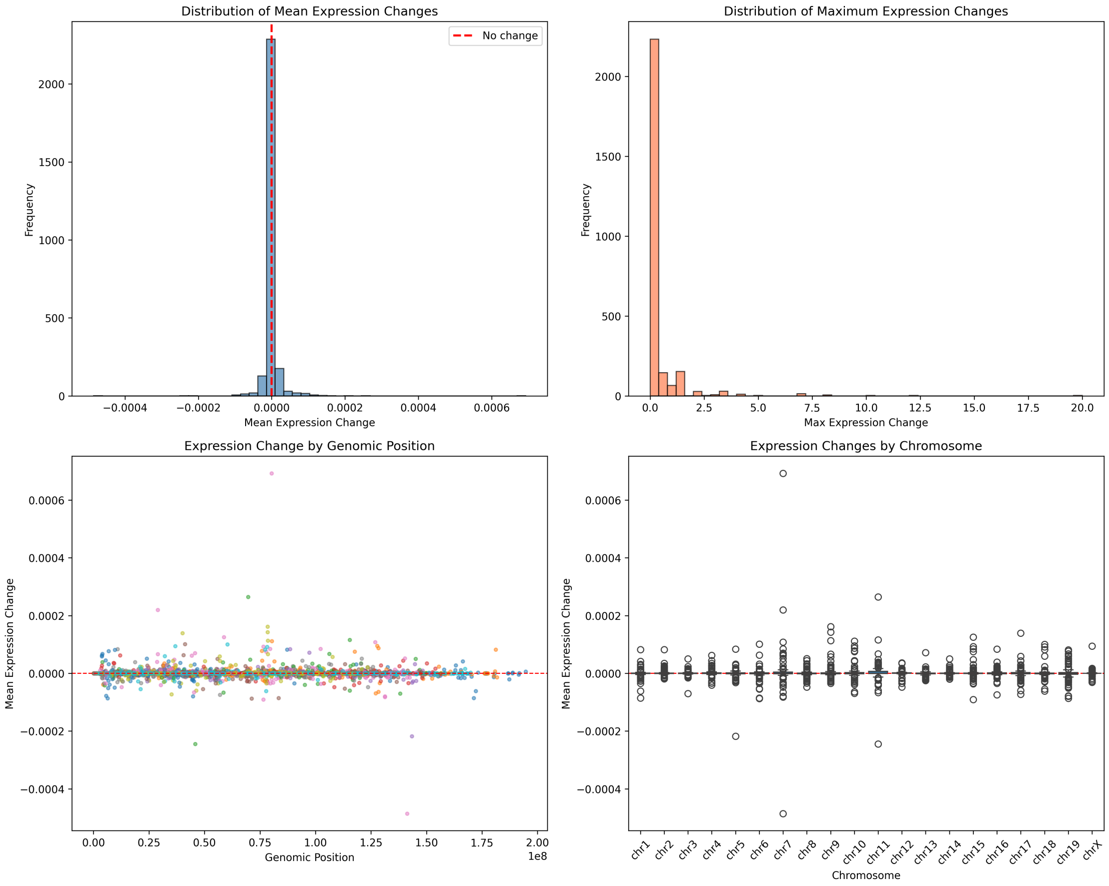


**Figure S2: Mouse mm10 mirror of the human R1 random-SNV simulation (main-text Figure 1; 2,700 random mm10 SNVs).** The same random-genome-wide simulation re-run on mm10 with mouse colon ontology, yielding the same near-zero per-variant effect distribution. *Species: mouse* (mm10 SNVs scored in mouse colon).

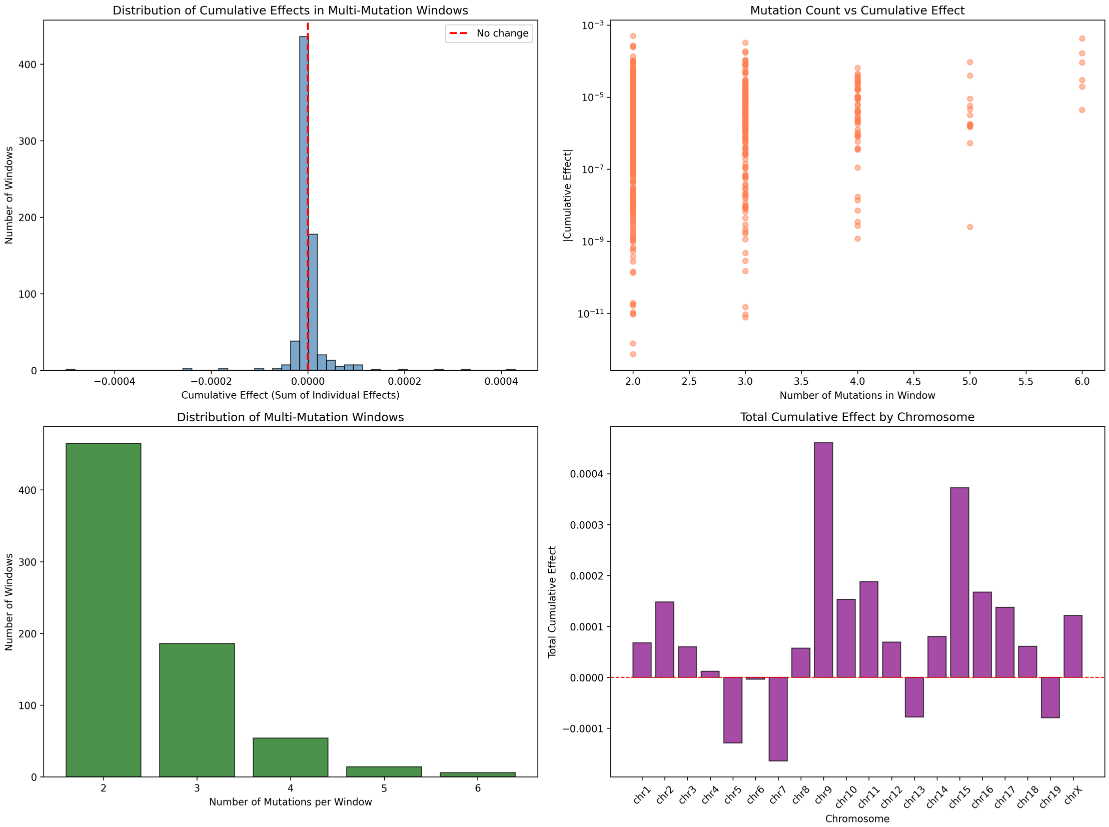


**Figure S3: Mouse mm10 mirror of the human R2 1-Mb combined-effect analysis (main-text Figure 3).** Variants binned into 1-Mb windows; cumulative window effects scale roughly linearly with variant count and remain orders of magnitude below the aging $\log_2$FC scale. *Species: mouse* (mm10).

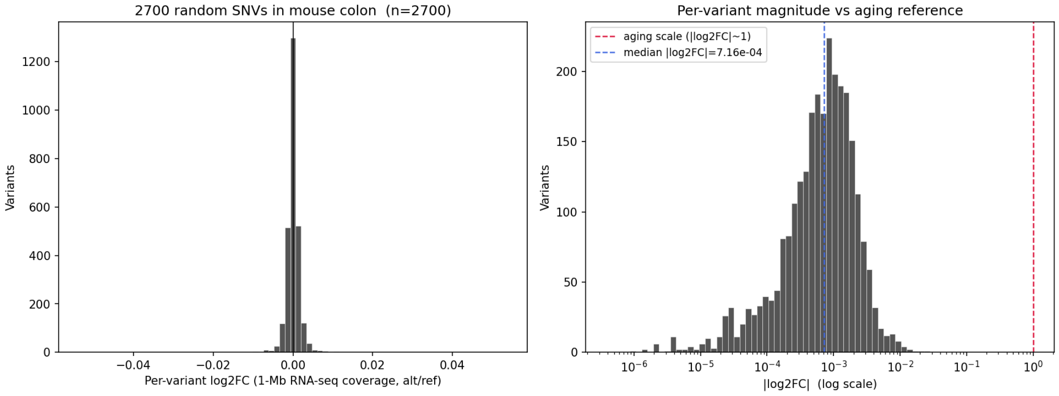


**Figure S4: Per-variant $\log_2$FC distribution from the mouse mm10 random-SNV simulation.** Companion histogram to Supplementary Figure S2. The distribution is tightly centred near zero, mirroring the human hg38 result. *Species: mouse* (mm10).

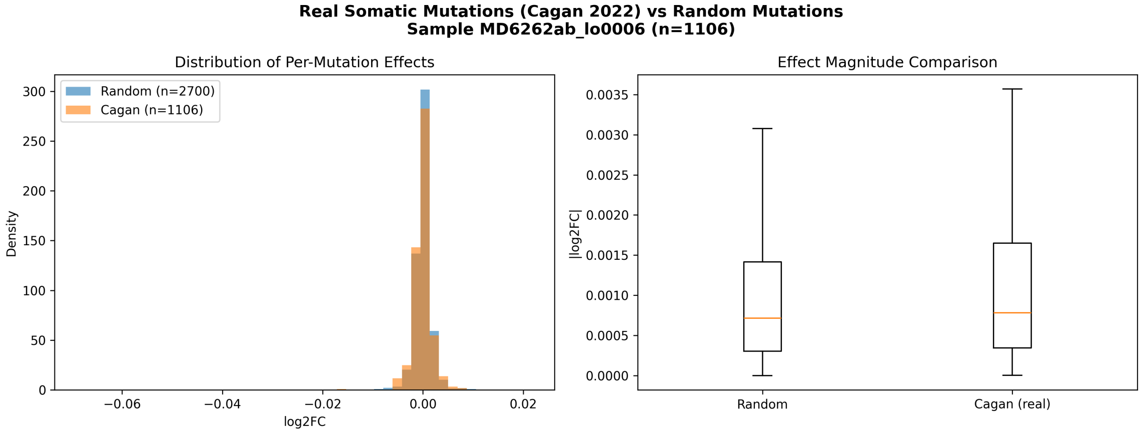


**(a)** Comparison of per-variant $\log_2$FC distributions in the mouse cohort (primary sample MD6262ab_lo0006, $n = 1{,}106$ SNVs against the matched mm10-native 2,700-random-SNV baseline; colon UBERON:0001157, mm10-native predictions). The mouse Cagan and random distributions overlap closely (mean $|\log_2 \mathrm{FC}|$ $1.34 \times 10^{-3}$ vs $1.12 \times 10^{-3}$, 1.20×).

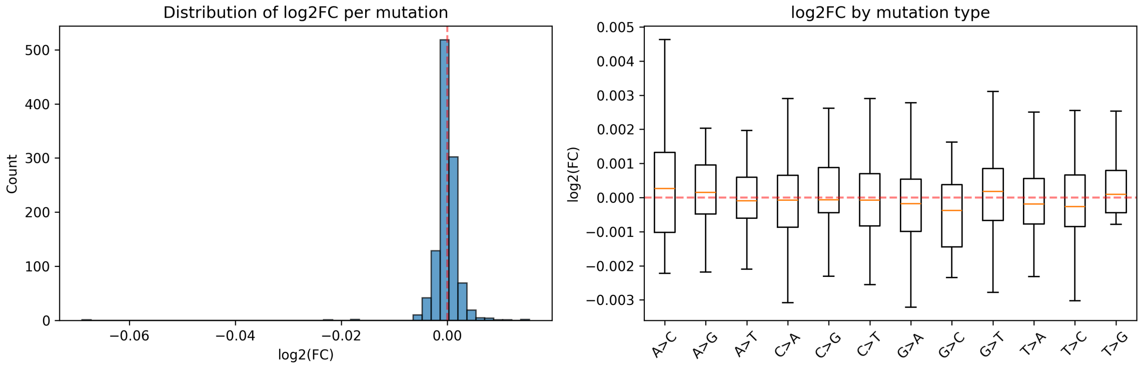


**(b)** Per-variant $\log_2$FC distribution for the same mouse sample, with 2-fold-change threshold $|\log_2 \mathrm{FC}| = 1$ marked.

**Figure S5: Mouse-cohort equivalent of Figure 7.** *Species: mouse.* Retained because the mouse cohort is larger ($n = 54$ crypts; 9,177 SNVs across five detailed samples, scored mm10-natively) than the three-donor human cohort shown in the main-text figure; conclusions are identical in both species.

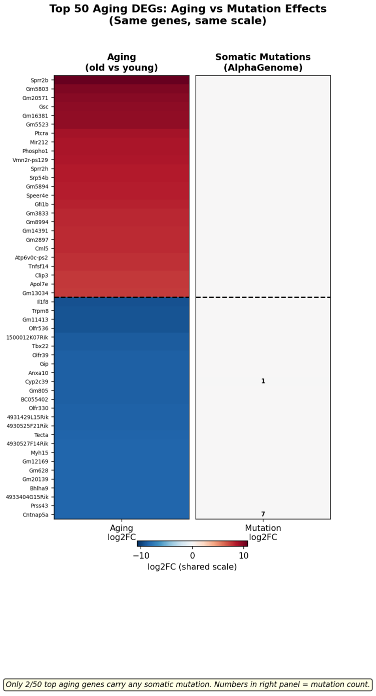


**(a)** Top 25 up- + top 25 down-regulated aging DEGs.

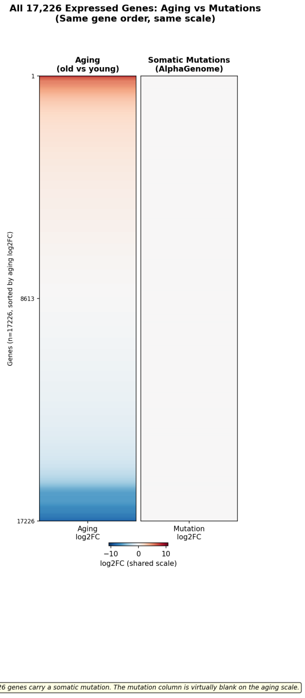


**(b)** All ∼17,000 expressed genes, sorted by aging $\log_2$FC.

**Figure S6: Mouse-on-mouse Cagan-vs-aging heatmap (R11 supplement).** 1,781 real somatic mutations from Cagan sample MD6267ab_lo0015 plotted side-by-side with Tabula Muris Senis aging $\log_2$FC on a shared diverging colour scale, in mouse large-intestine epithelium. (*Left*) Top 25 up + 25 down aging DEGs; the aging column paints the full $[-10, +12]$ gradient while the mutation column is entirely neutral. (*Right*) All $\sim$ 17k expressed genes ordered by aging $\log_2$FC; mutation column blank along the entire axis despite 755 of those genes carrying $\geq 1$ real somatic mutation. Max aging-to-mutation ratio $\approx 840\times$, median $\approx 990\times$ at 1-Mb-window scoring. *Species: mouse* (Tabula Muris Senis; Cagan *et al.* 2022 mouse sample MD6267ab_lo0015).

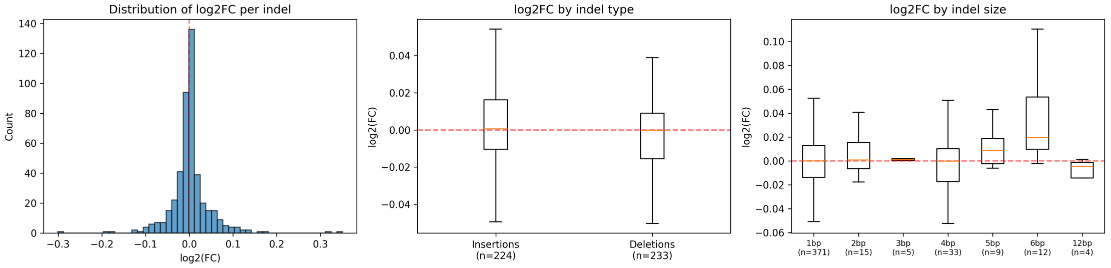


**(a)** Distribution of per-indel predicted expression $\log_2$FC for primary mouse sample MD6264o_lo0009 ($n = 457$ indels). Histograms split insertions and deletions; rug marks at bottom indicate individual indels. The distribution median sits at $|\log_2\text{FC}| \approx 0.03$, with a tail extending toward $|\log_2\text{FC}| \sim 1$ for a small number of splice- or CDS-disrupting events.

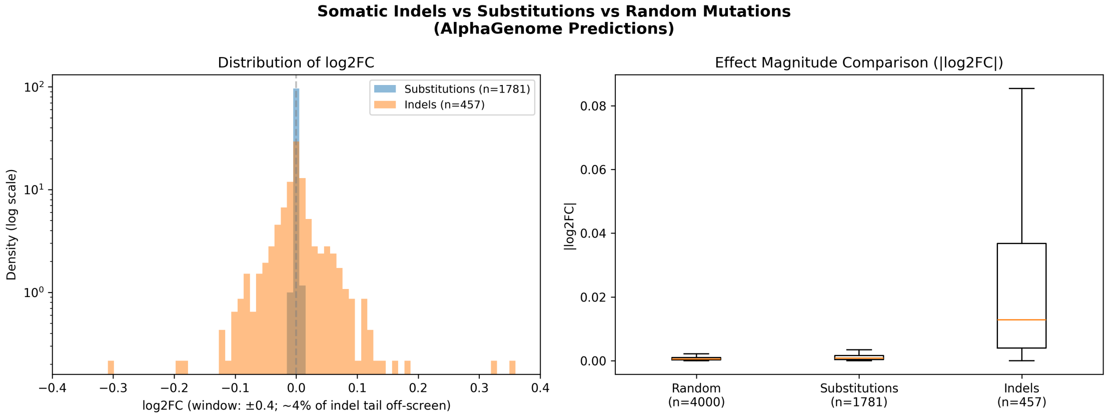


**(b)** Direct mouse indel-vs-substitution comparison within the same sample: overlaid distributions of per-event $|\log_2 \text{FC}|$ and box plots of absolute effect, with medians annotated. The 24.4× per-event ratio is evident, as is the heavier indel tail reflecting occasional frame- or splice-disrupting events. Both distributions remain below the aging-scale $\log_2$FC reference.

**Figure S7: Mouse-cohort counterpart of main-text Figure 9 ($n = 8$ mouse samples, 3,233 indels in total; primary detail sample MD6264o_lo0009, $n = 457$).** The mouse cohort confirms the per-event indel/SNV amplification observed in the human cohort: mean $|\log_2 \text{FC}|$ across 409 indels of sample MD6267ab_lo0003 was 0.030 vs 0.0012 for substitutions in the same sample (24.4× larger per mutation). The per-gene worst case observed across all 3,233 mouse indels in all 8 samples was $|\sum \log_2 \text{FC}| = 0.82$ at *Spertl* (4 indels in the same gene), still below the magnitude of a single top aging DEG. *Species: mouse* (Cagan *et al.* 2022 mouse colonic crypts).

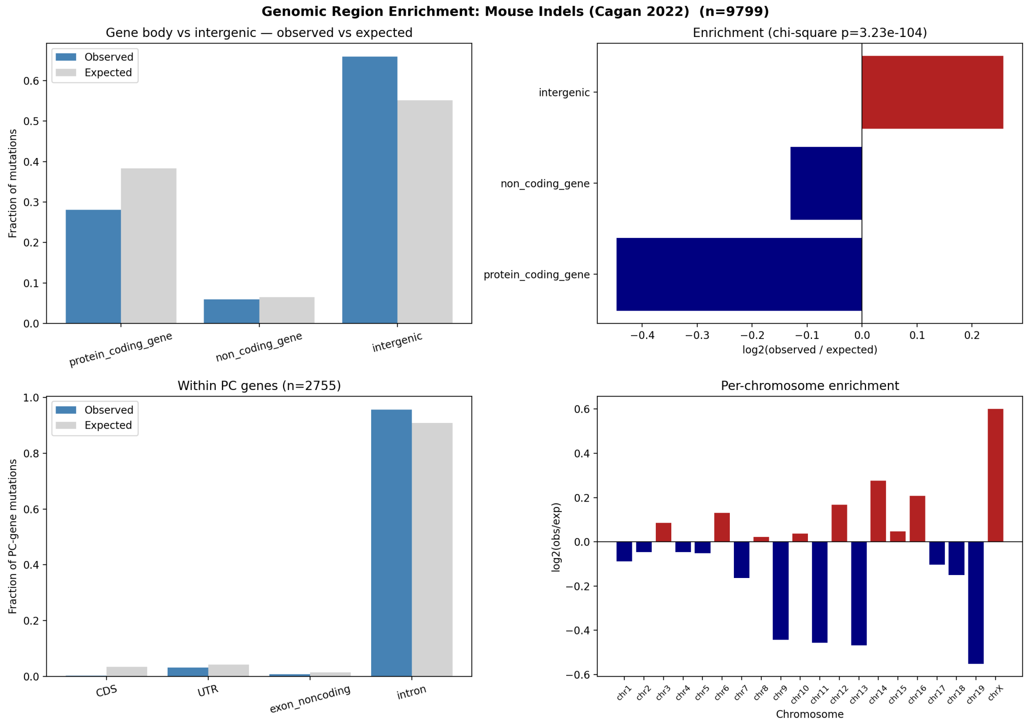


**Figure S8: Mouse-cohort counterpart of main-text Figure 10 ($n = 54$ samples, 9,799 indels).** The same purifying-selection signature seen in the human cohort, with much sharper magnitudes owing to the larger sample size: 12× CDS depletion ($\log_2 = -3.58$, only 8 of 2,755 protein-coding-gene indels in CDS), modest intergenic enrichment ($\log_2 = +0.26$), and a top-level $\chi^2$ goodness-of-fit p-value of $3.2 \times 10^{-104}$ that is power-driven by the 9,799-indel sample size (the same fractional deviations against $\sim 3\times$ fewer indels in human gave $p = 4.4 \times 10^{-2}$ at the top level). *Species: mouse* (Cagan *et al.* 2022, 54 mouse colonic crypts; GENCODE vM25 mm10 annotations).

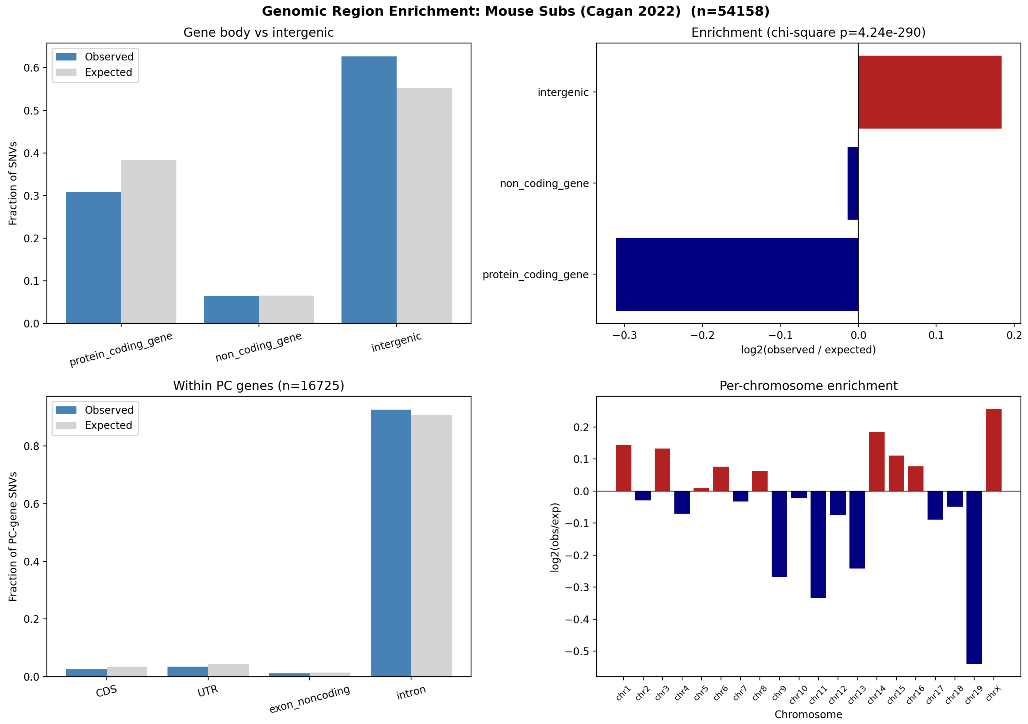


**Figure S9: Mouse-cohort counterpart of main-text Figure 11 ($n = 54$ samples, 54,158 SNVs).** Region-class enrichment of mouse SNVs reproduces the human direction at slightly weaker per-event magnitude: CDS $\sim 1.27\times$ depleted ($\log_2 = -0.34$, 460 of expected 583 hits), UTRs and exon-noncoding regions modestly depleted ($\log_2 \approx -0.32$ to $-0.36$), introns near-neutral, intergenic modestly enriched ($\log_2 = +0.18$). Within-PC $\chi^2$ $p = 1.0 \times 10^{-15}$; top-level $\chi^2$ $p = 4.2 \times 10^{-290}$ (the latter dominated by the 54,158-SNV sample size; the biologically meaningful quantity is the modest $\log_2$ effect size, which is roughly an order of magnitude smaller than the indel result of Supplementary Figure S8). *Species: mouse* (Cagan *et al.* 2022, 54 mouse colonic crypts; GENCODE vM25 mm10 annotations).
