## Supplementary figures and images for "Testing the mutation accumulation hypothesis in aging with AlphaGenome"

### 02_pooled_abs_log2fc_logscale.png

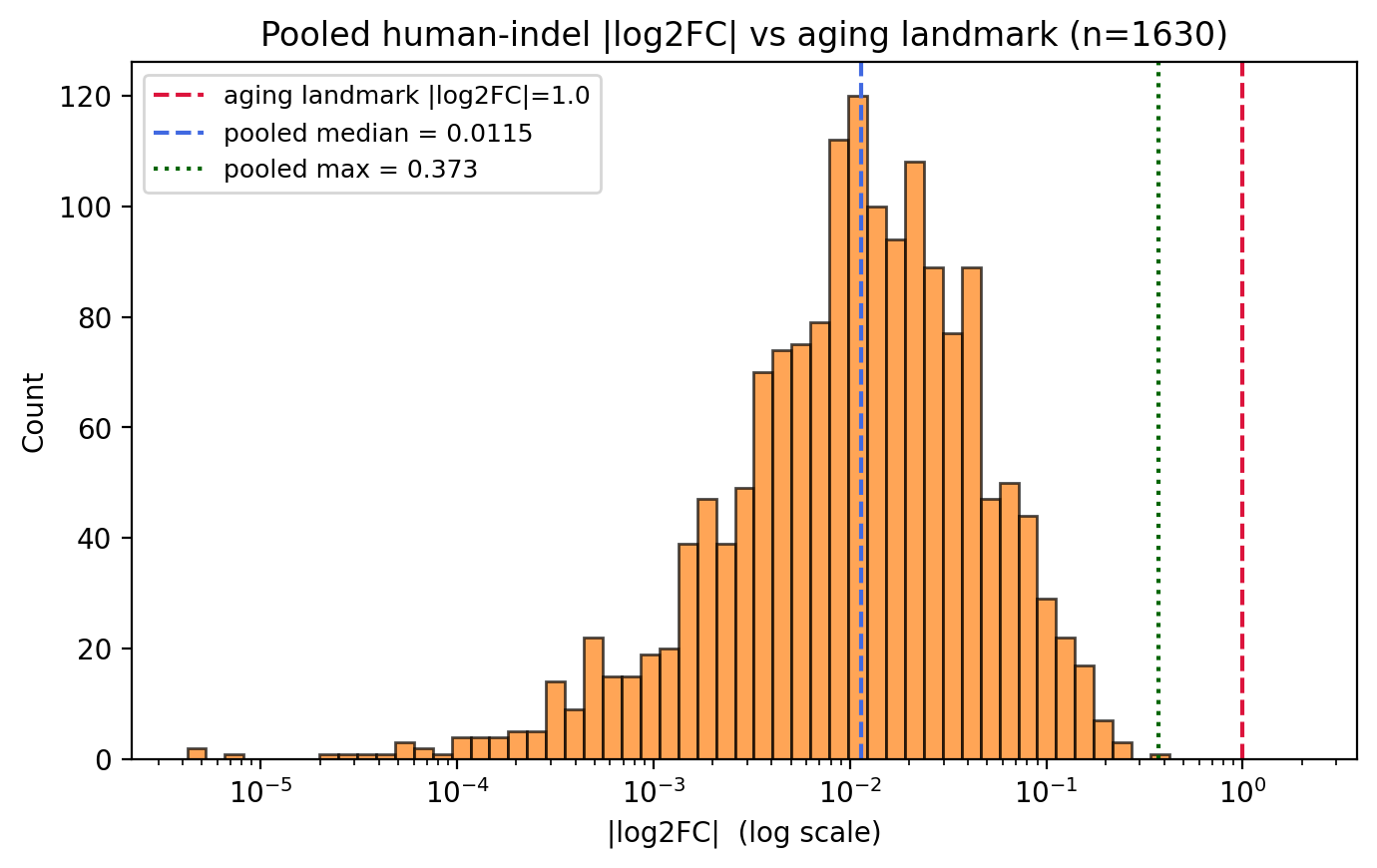

### 09_per_gene_cumulative_vs_aging.png

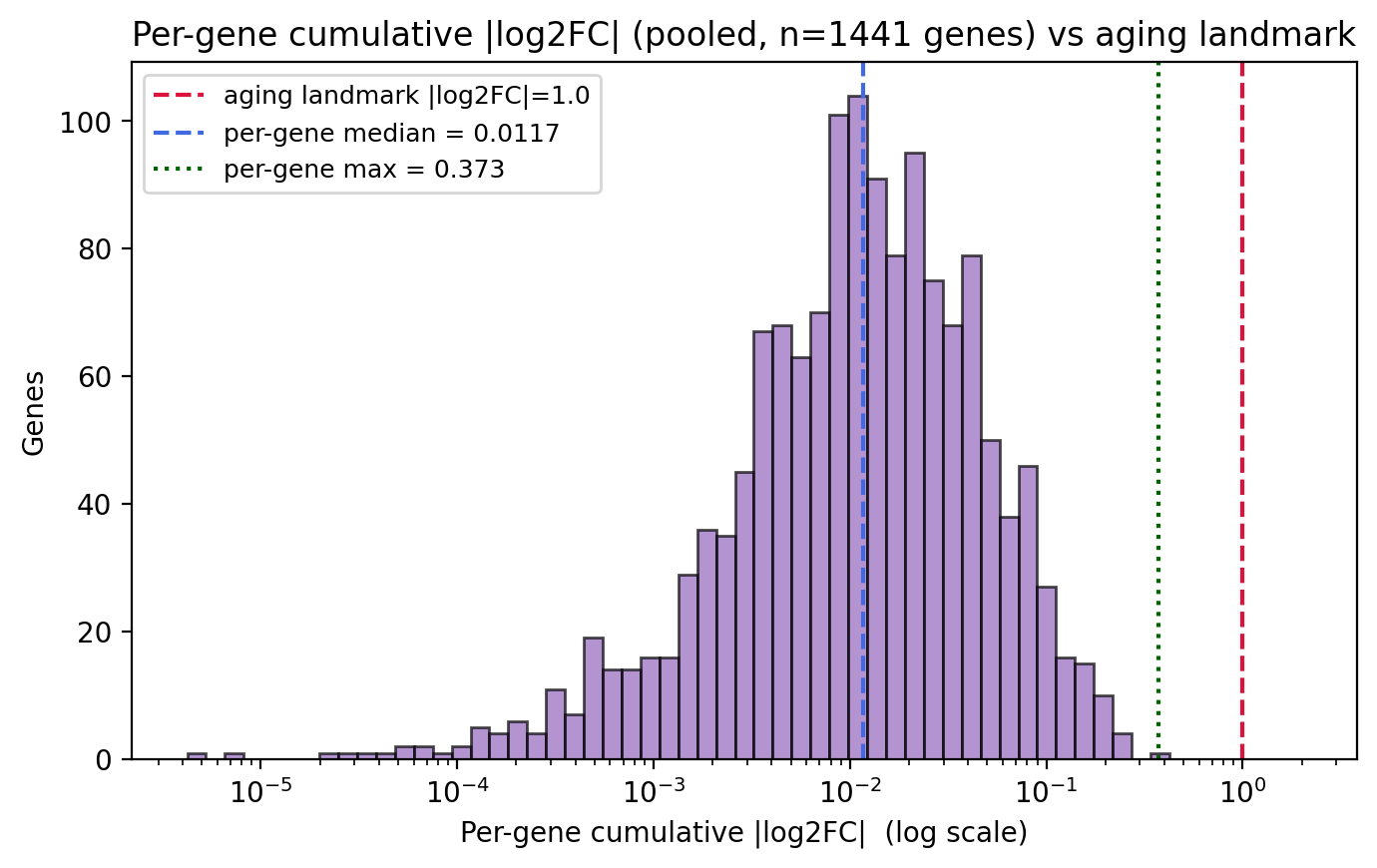

### aging_vs_mutations_all_genes.png

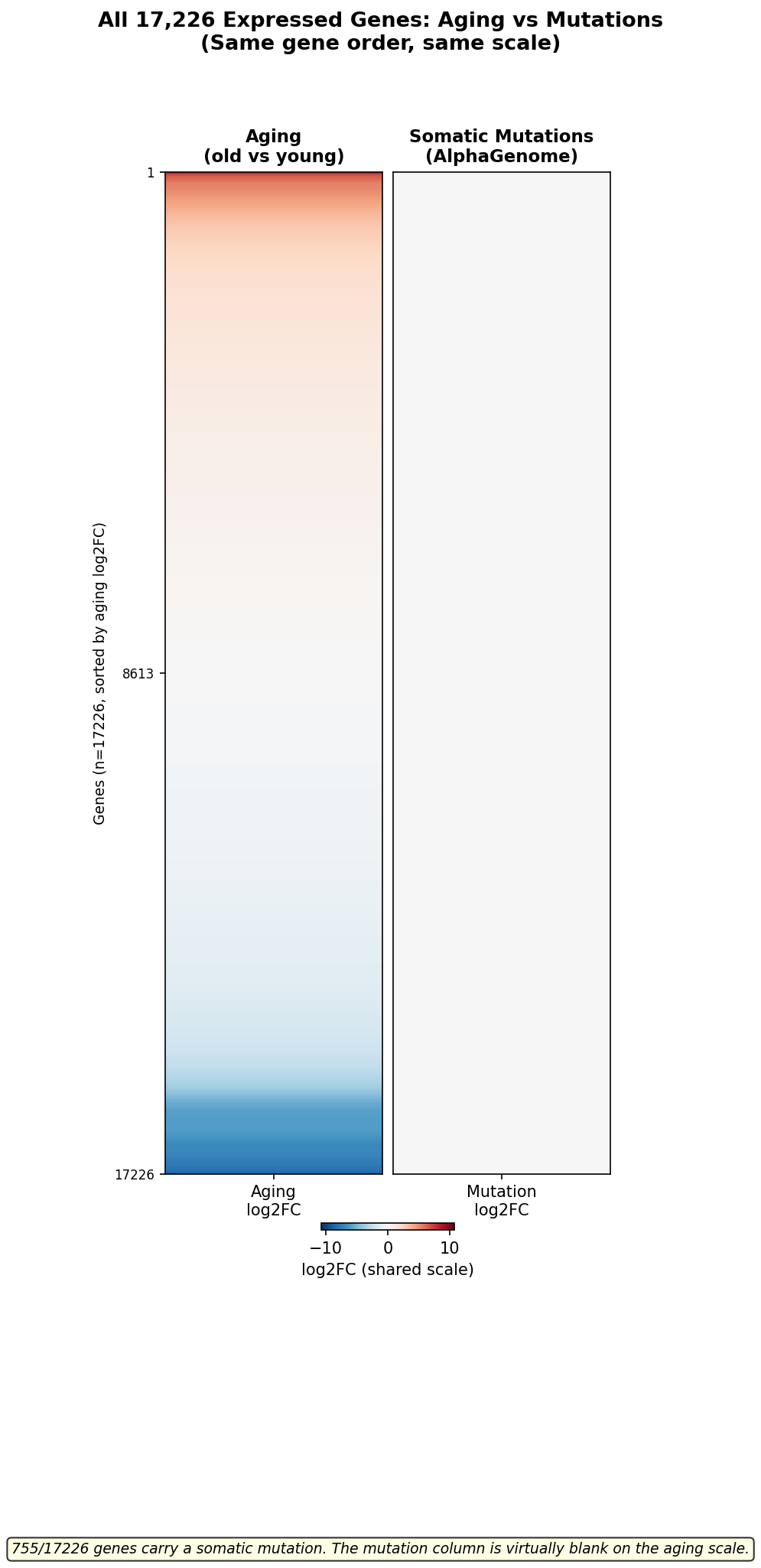

### aging_vs_mutations_gene_level_top50.png

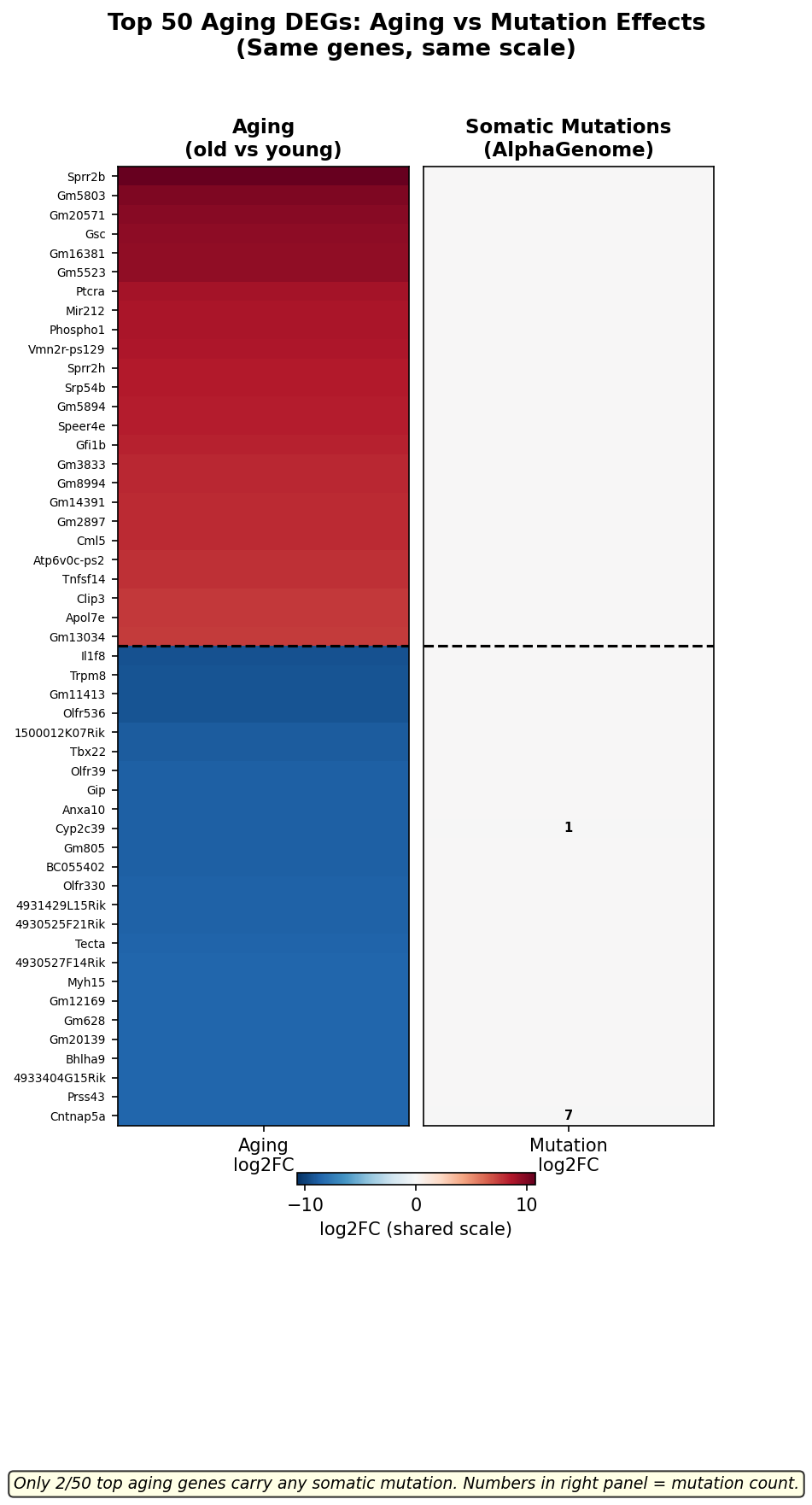

### aging_vs_mutations_human_all_genes.png

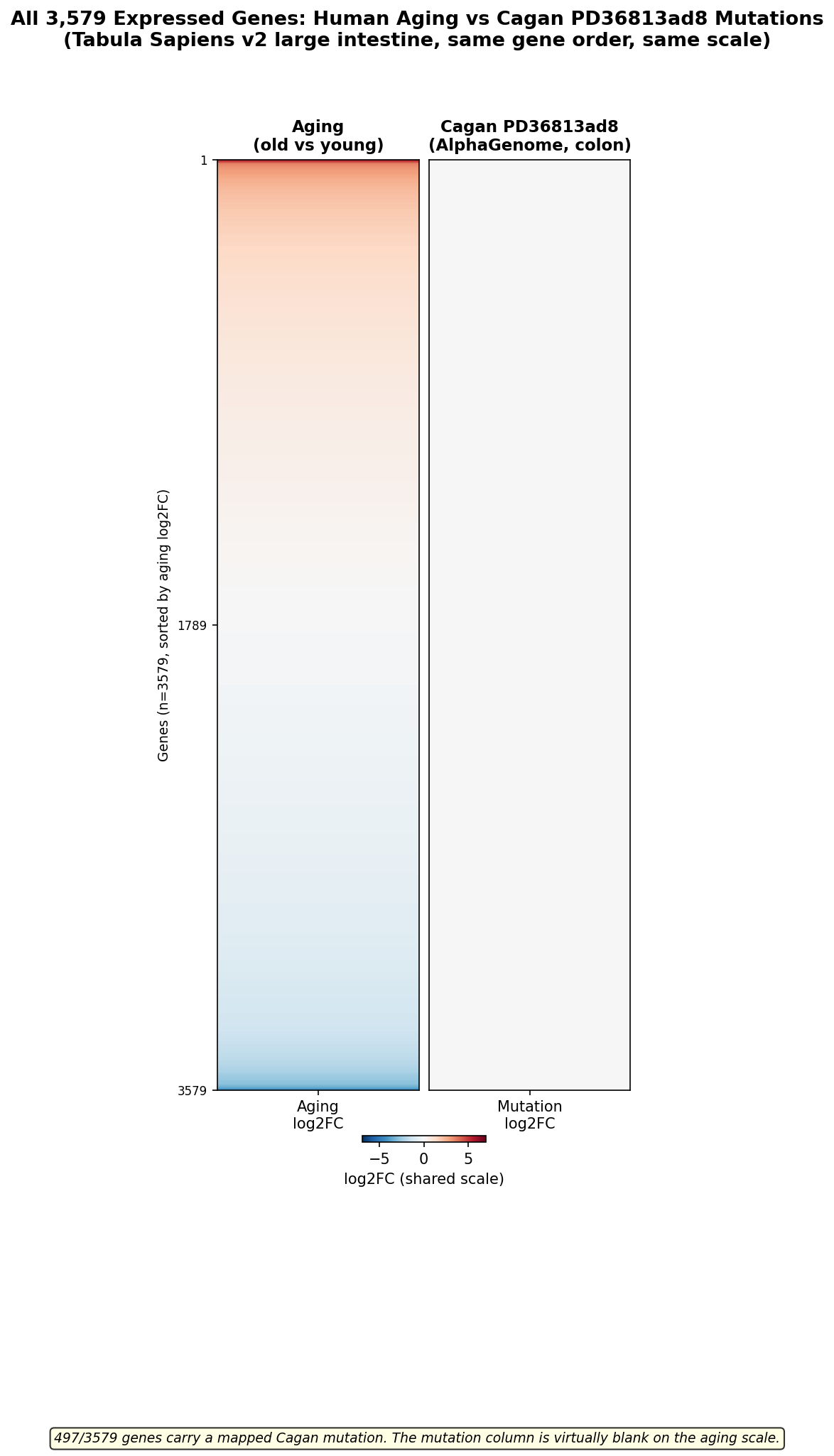

### aging_vs_mutations_human_gene_level_top50.png

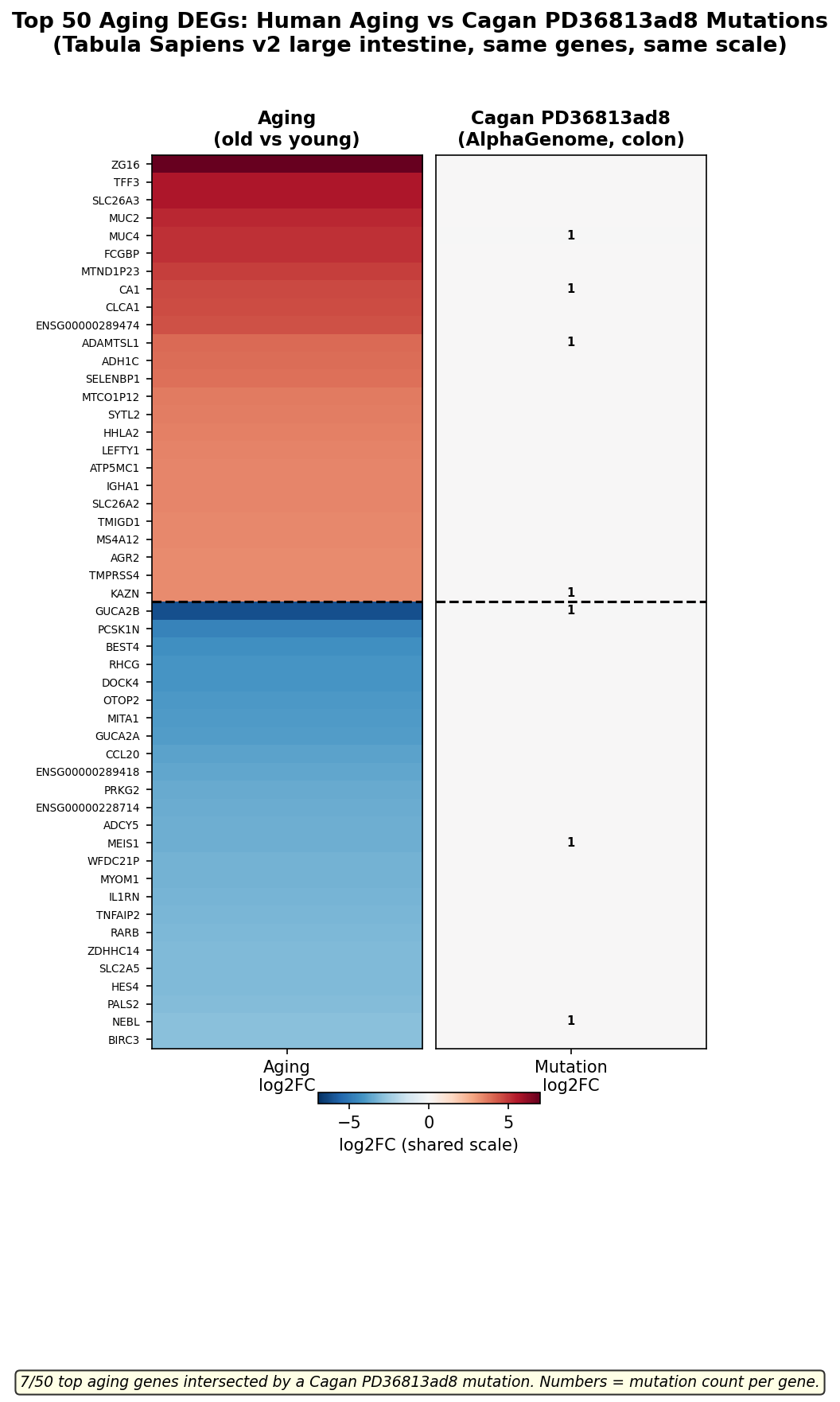

### aging_vs_random2700_mouse_all_genes.png

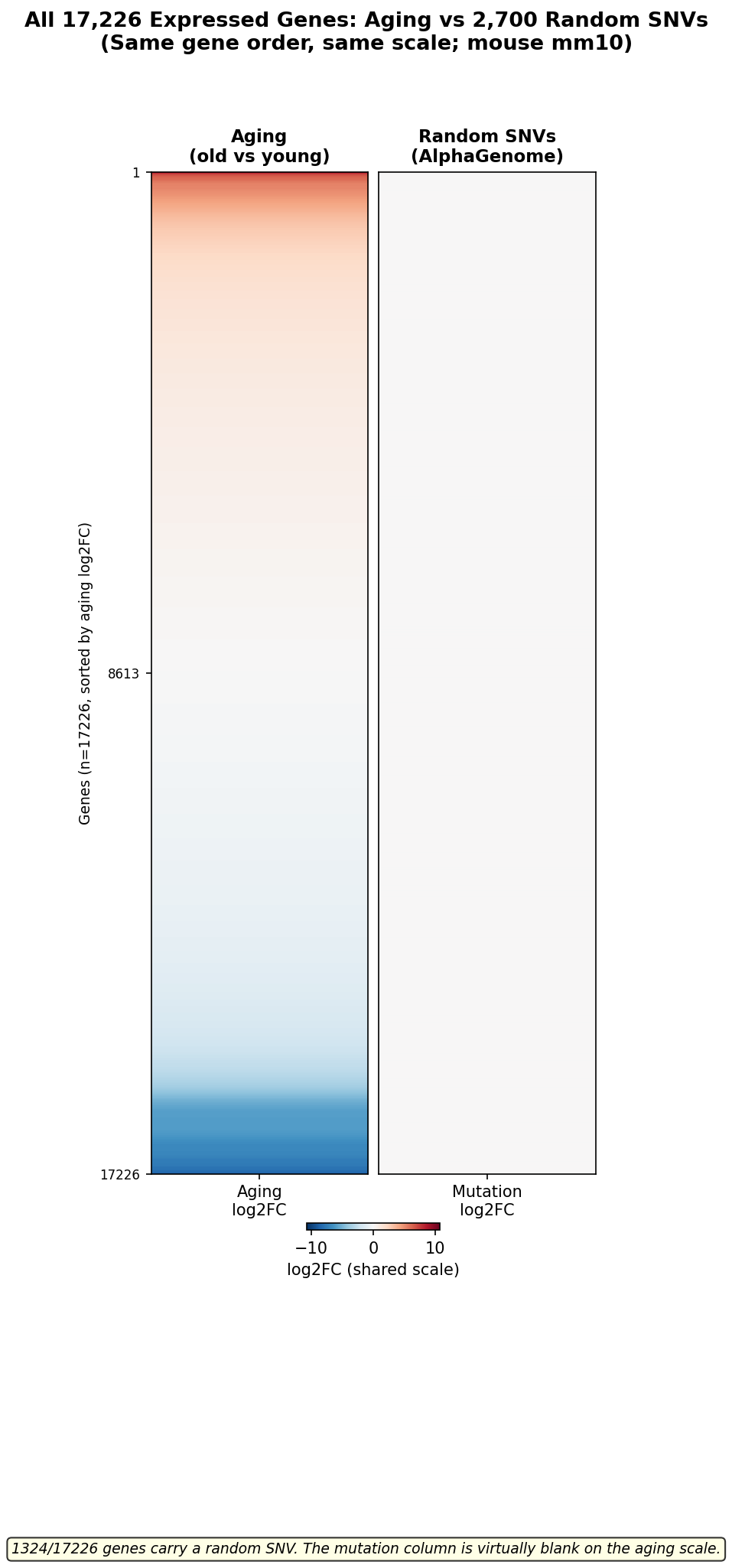

### aging_vs_random2700_mouse_gene_level_top50.png

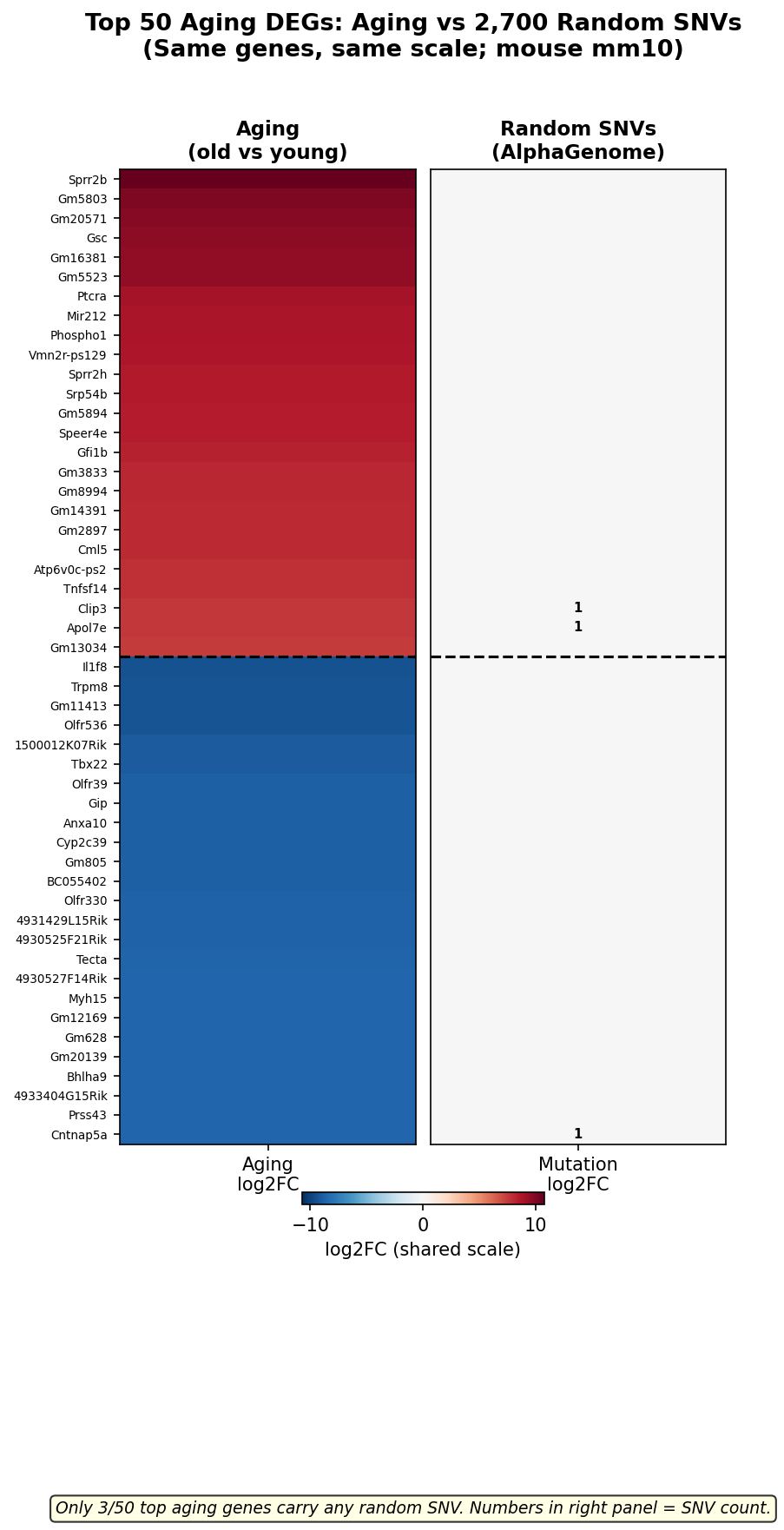

### aging_vs_random4000_human_all_genes_log2fc.png

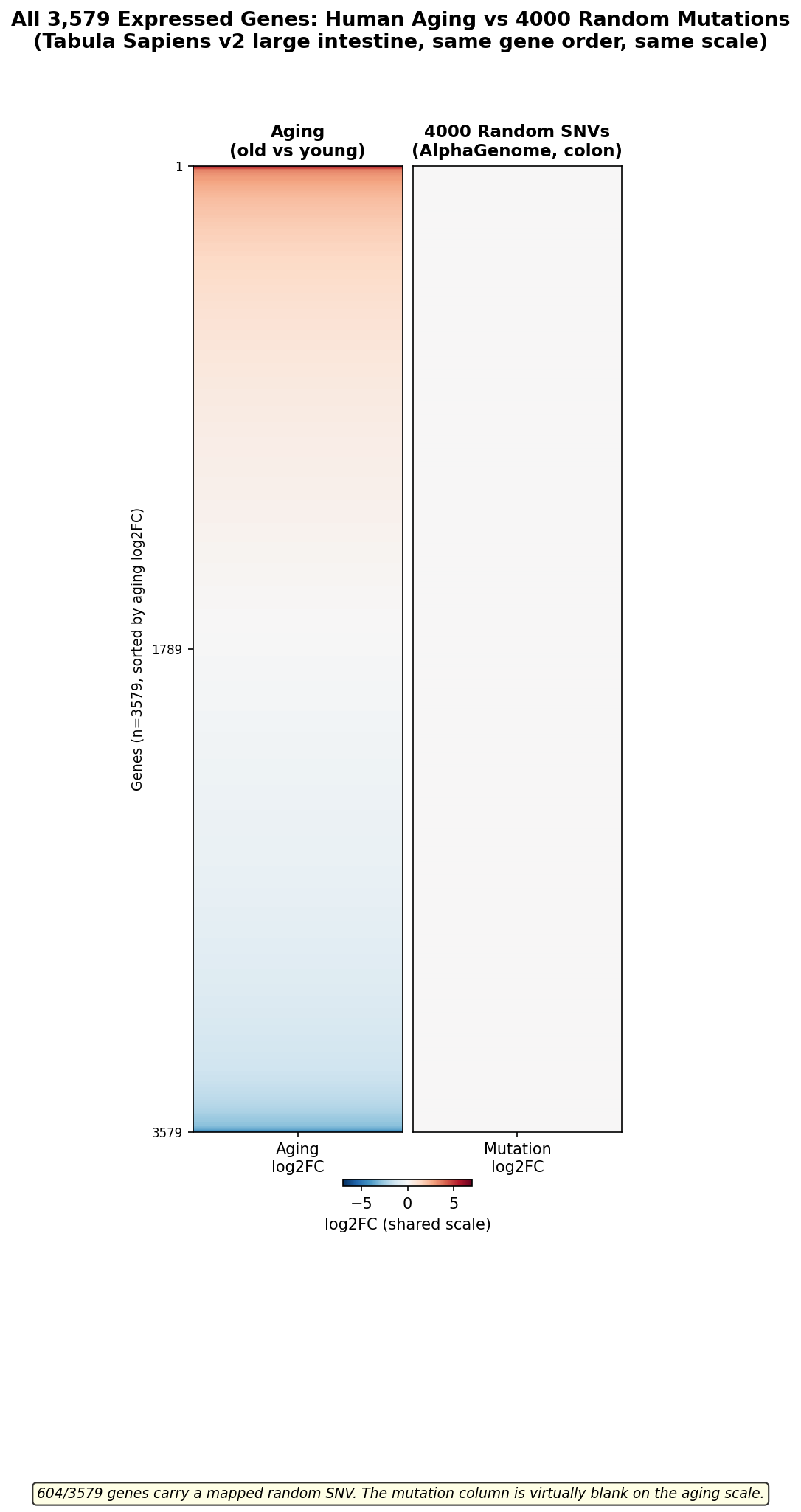

### aging_vs_random4000_human_gene_level_top50_log2fc.png

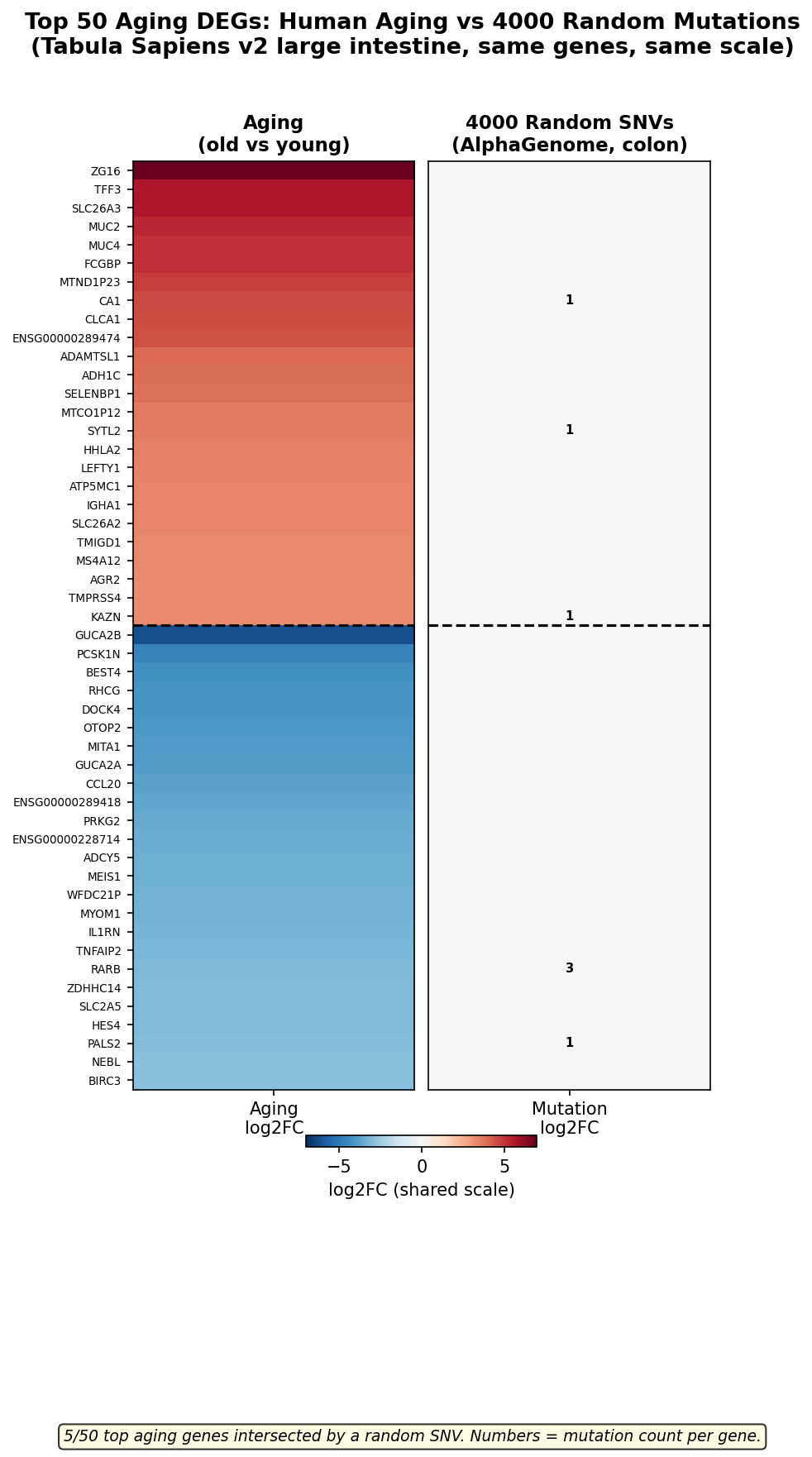

### cagan_vs_random_comparison.png

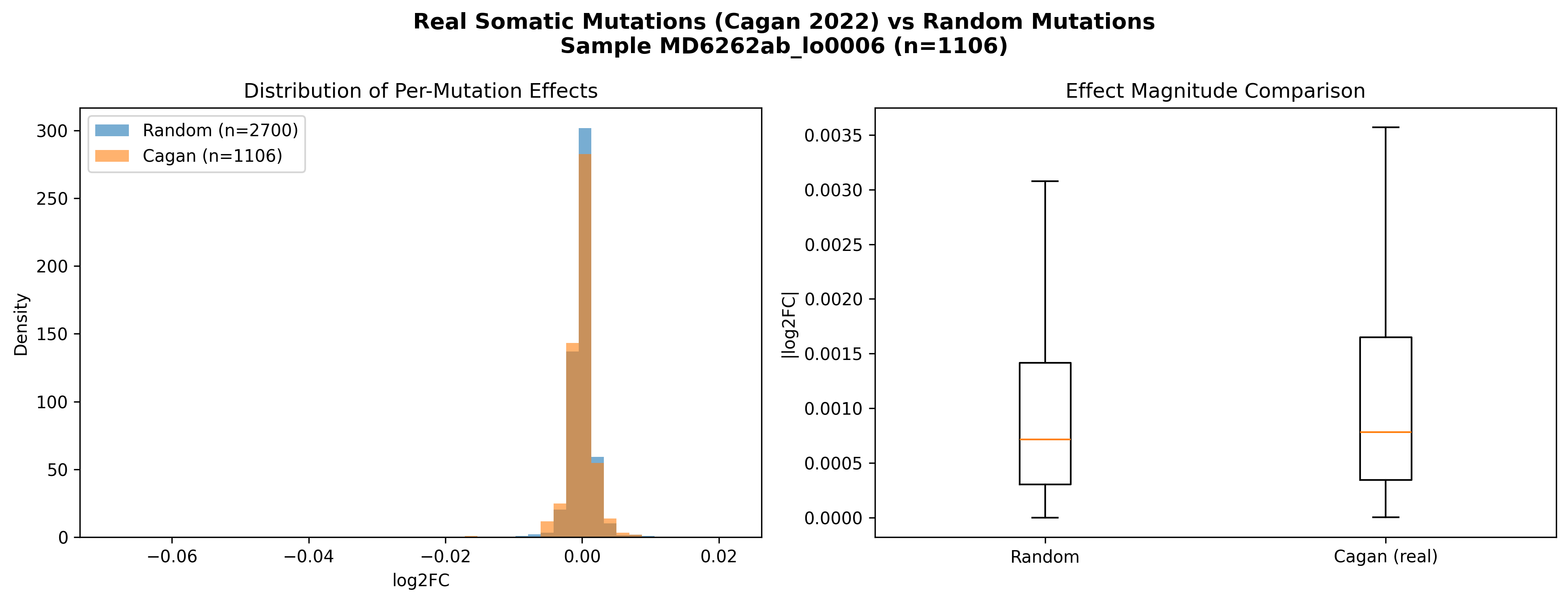

### colon_mutation_simulation_plots.png

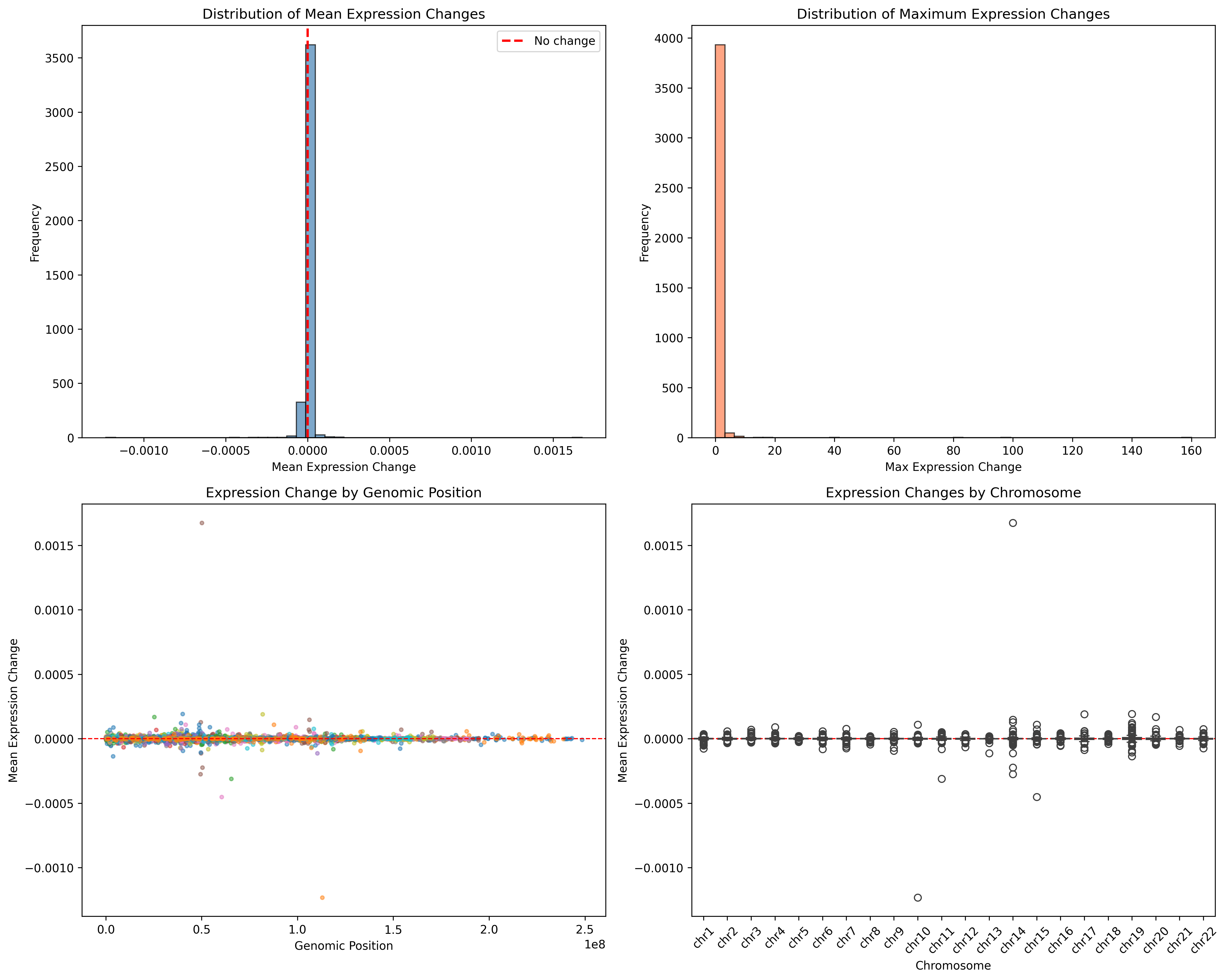

### combined_effects_analysis.png

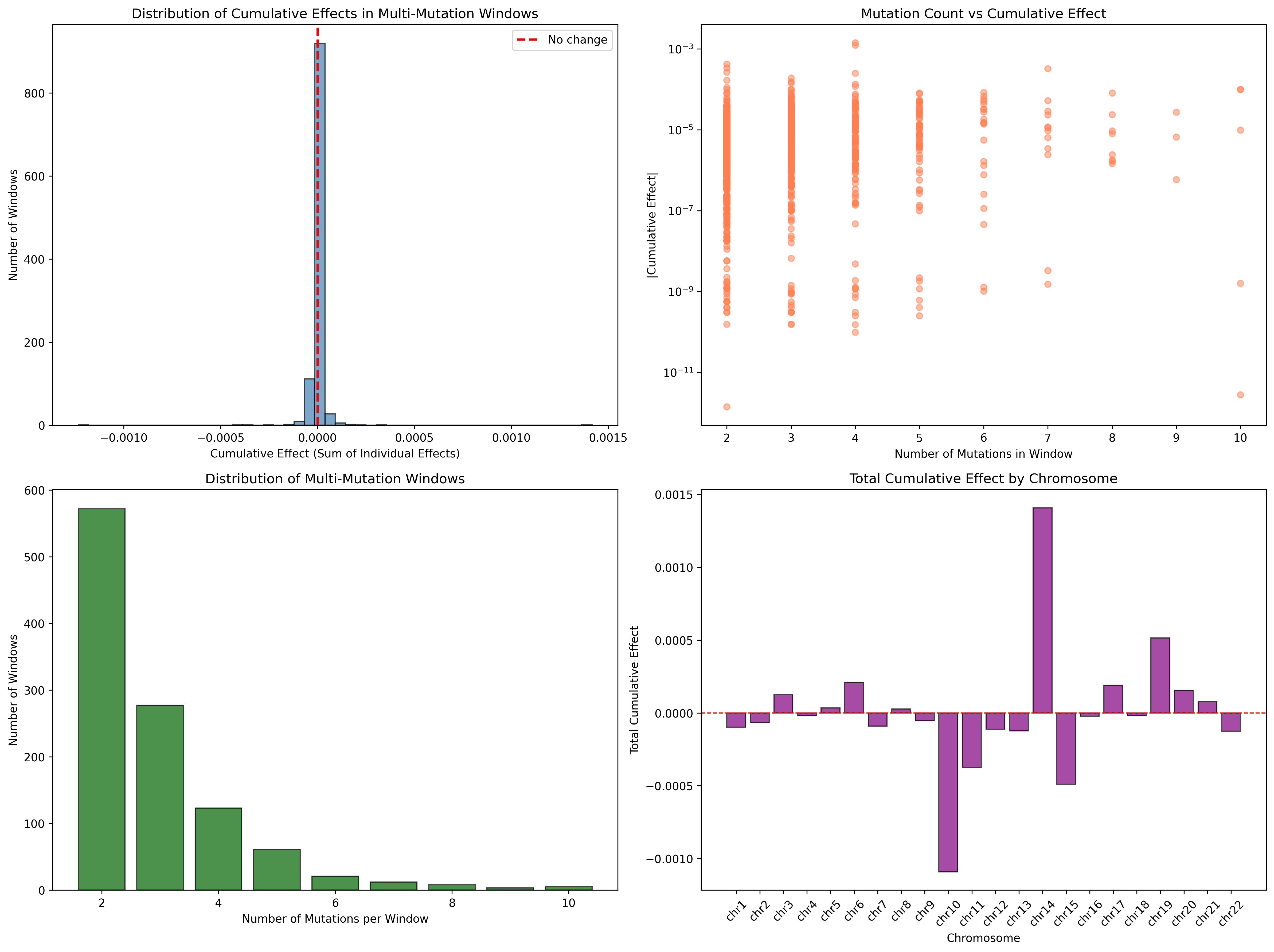

### enrichment_Human_Indels_Cagan_2022.png

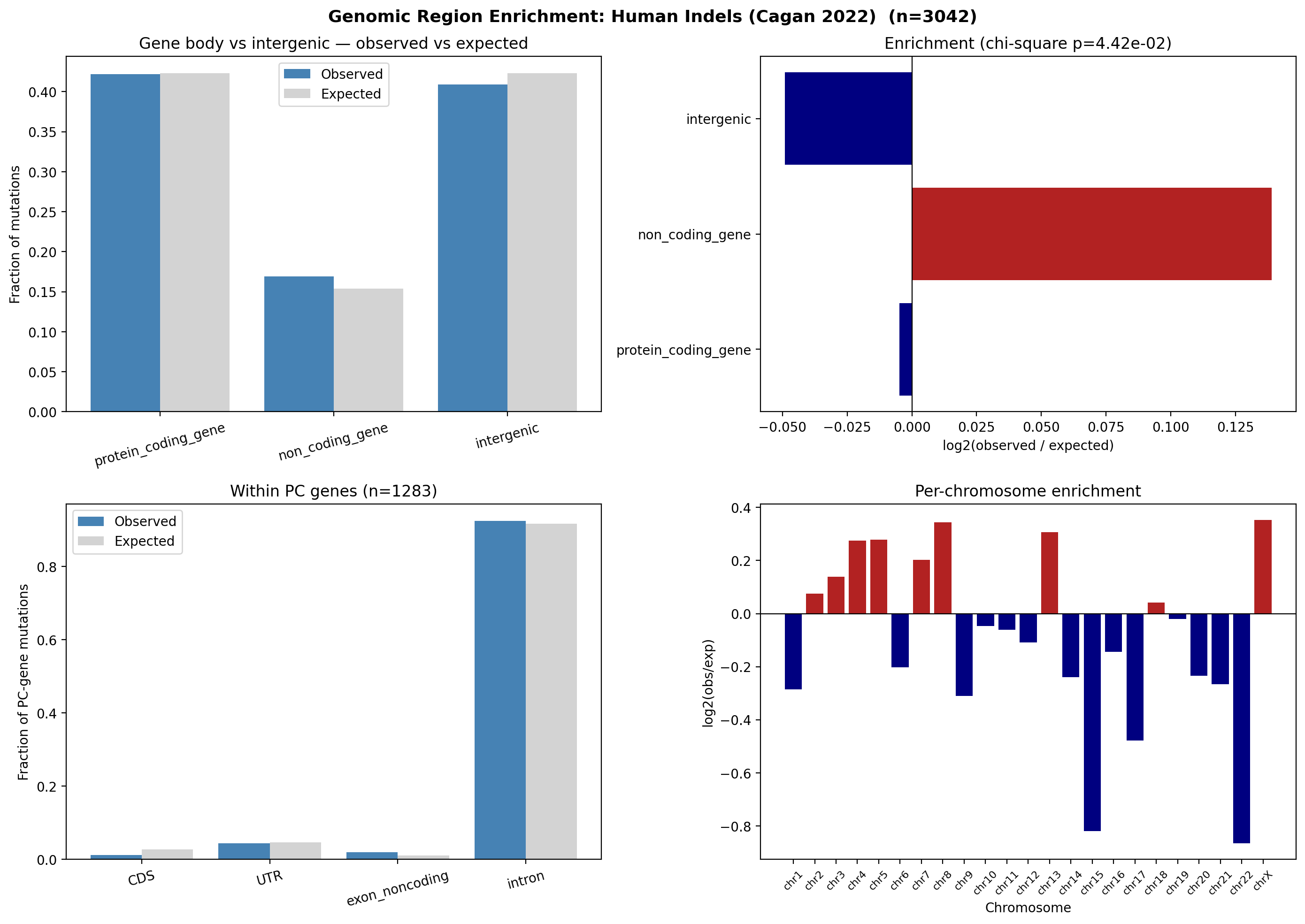

### enrichment_Human_Subs_Cagan_2022.png

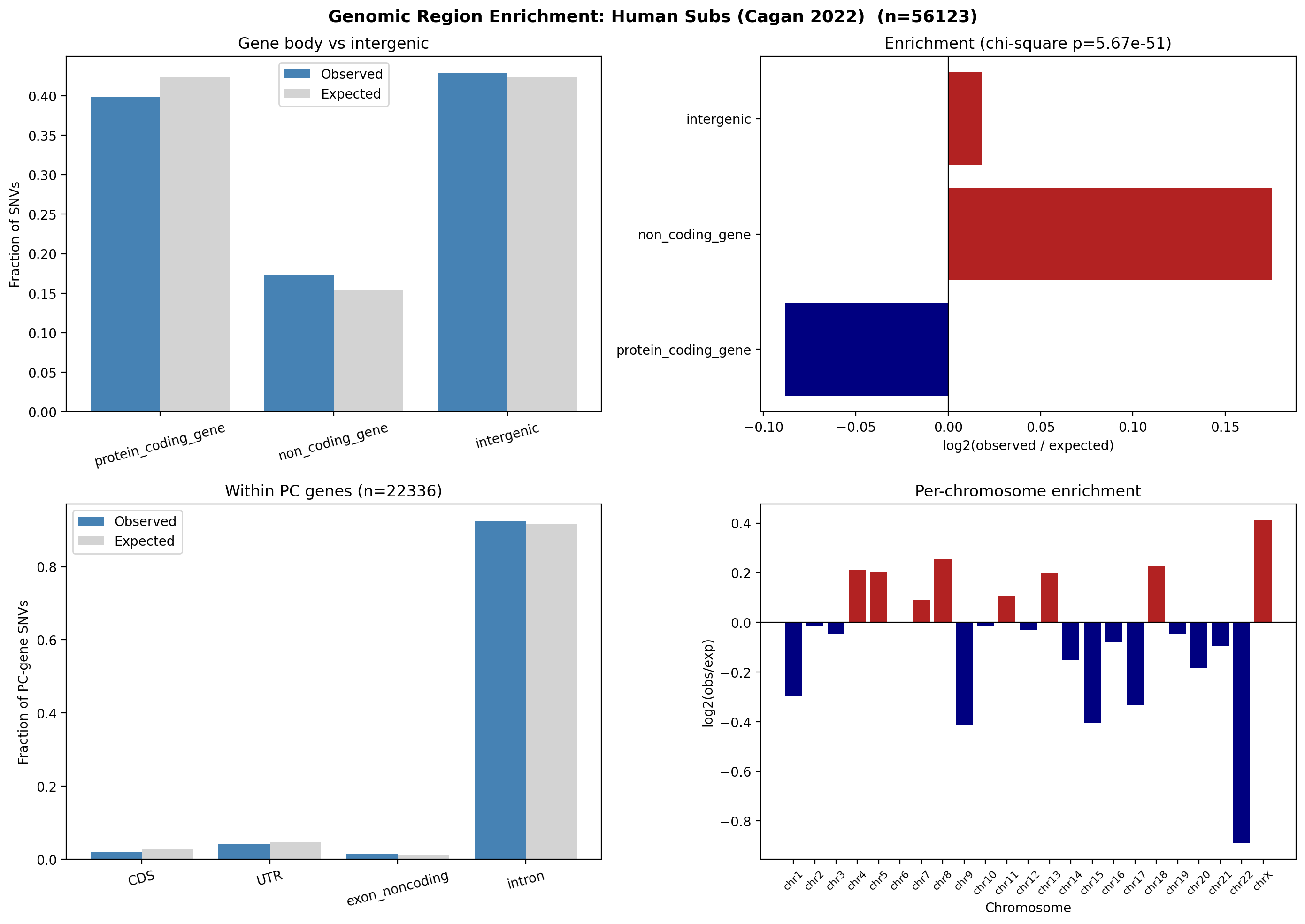

### enrichment_Mouse_Indels_Cagan_2022.png

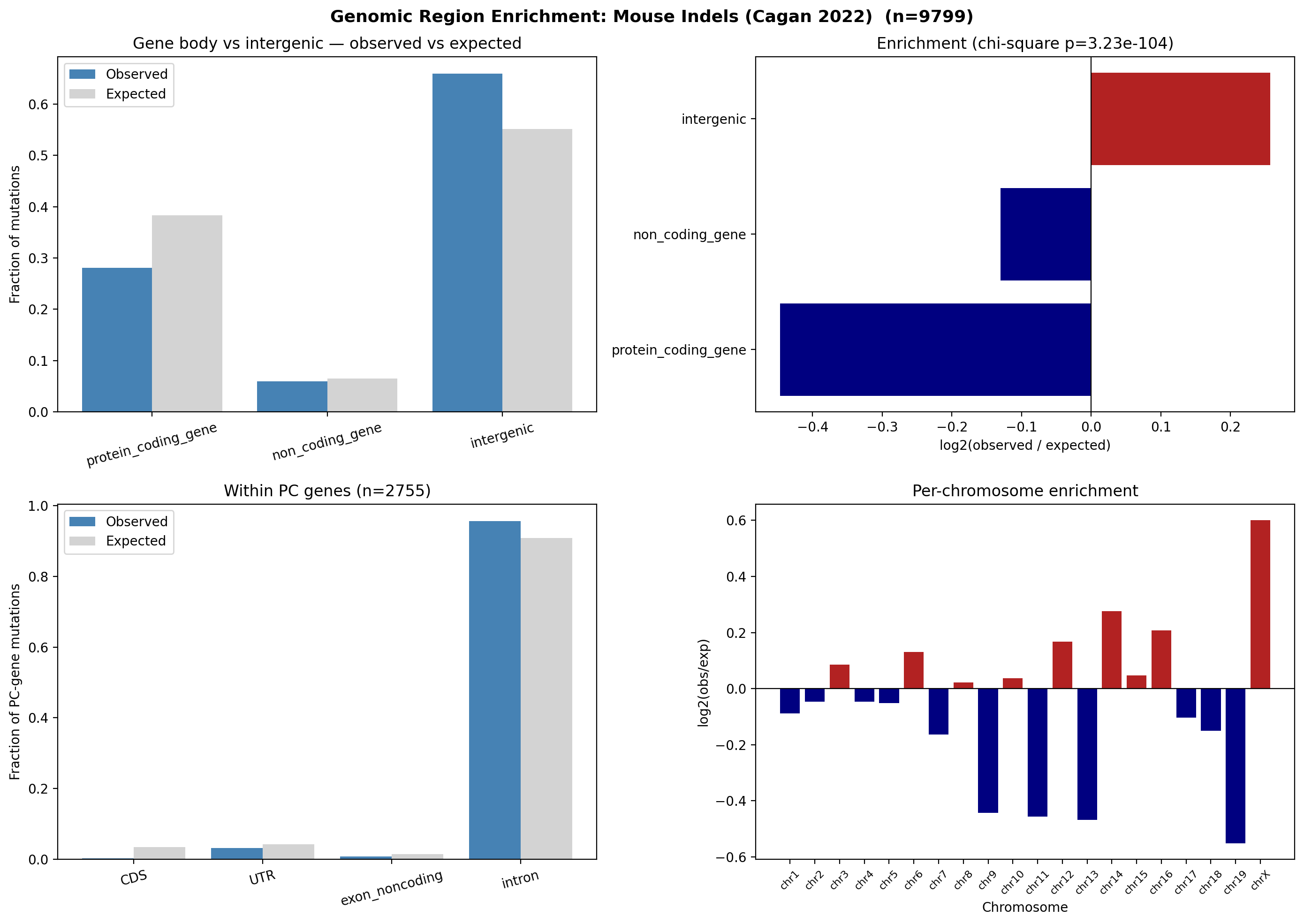

### enrichment_Mouse_Subs_Cagan_2022.png

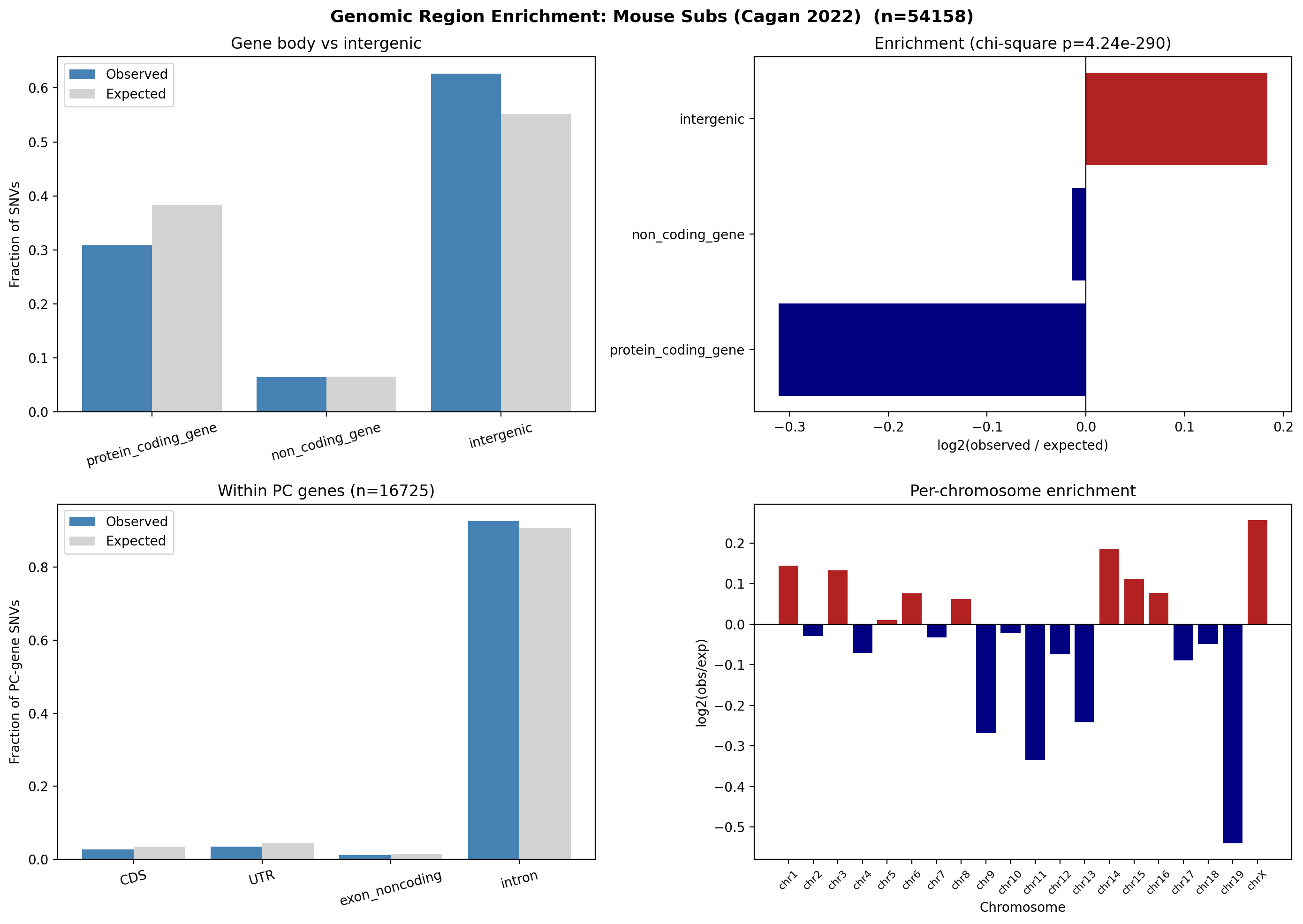

### gene_level_log2fc_vs_aging_landmark.png

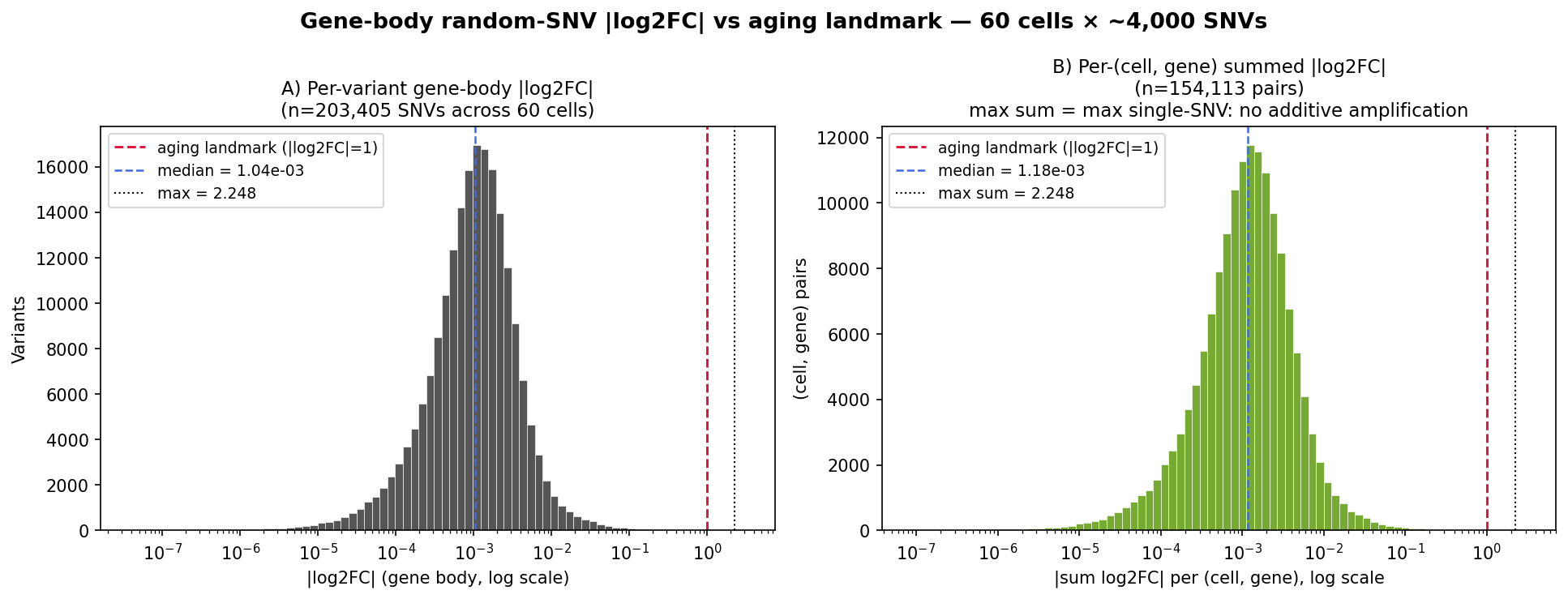

### gene_set_overlap.png

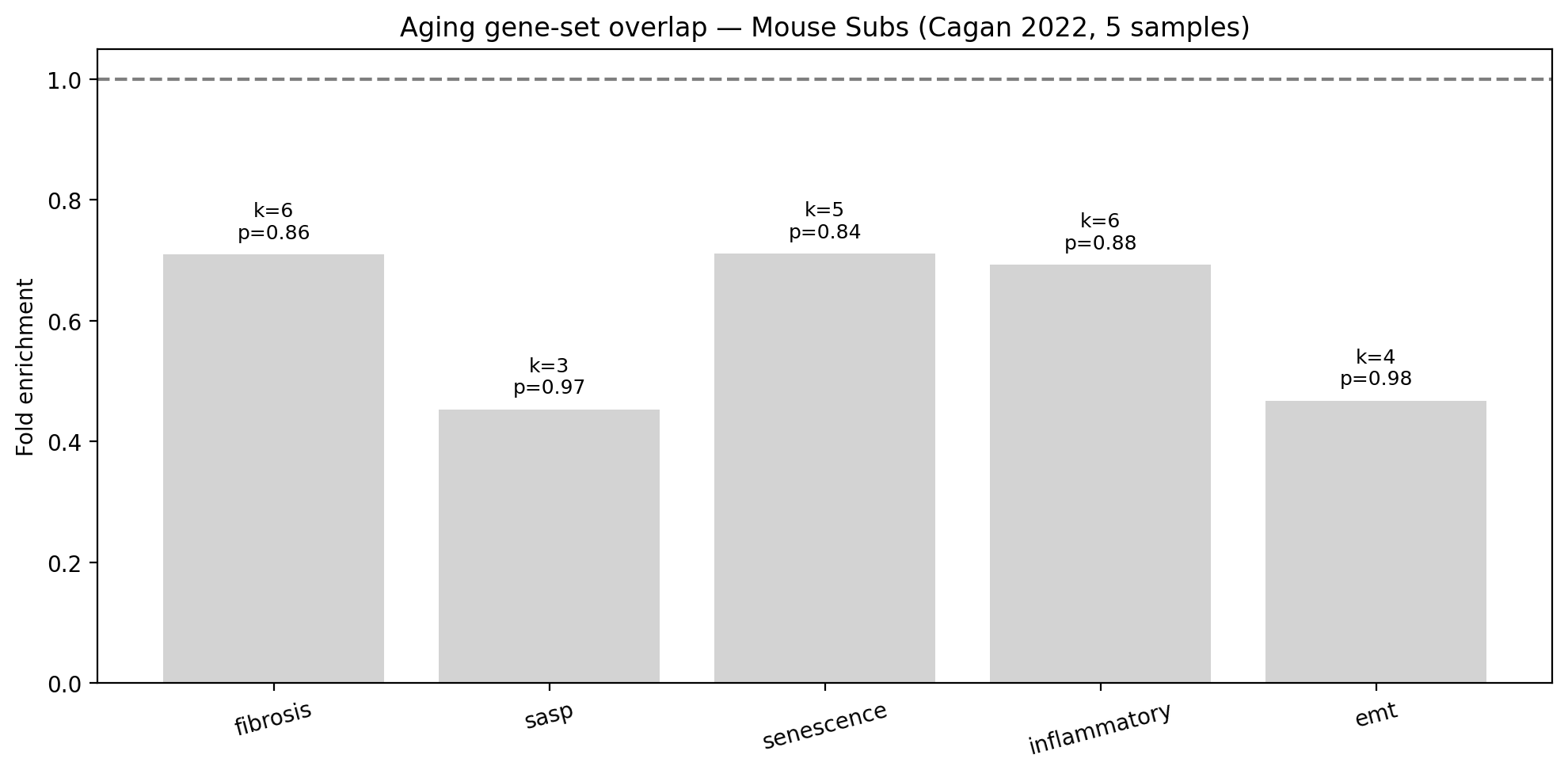

### gene_set_overlap_Mouse_Indels_Cagan_2022.png

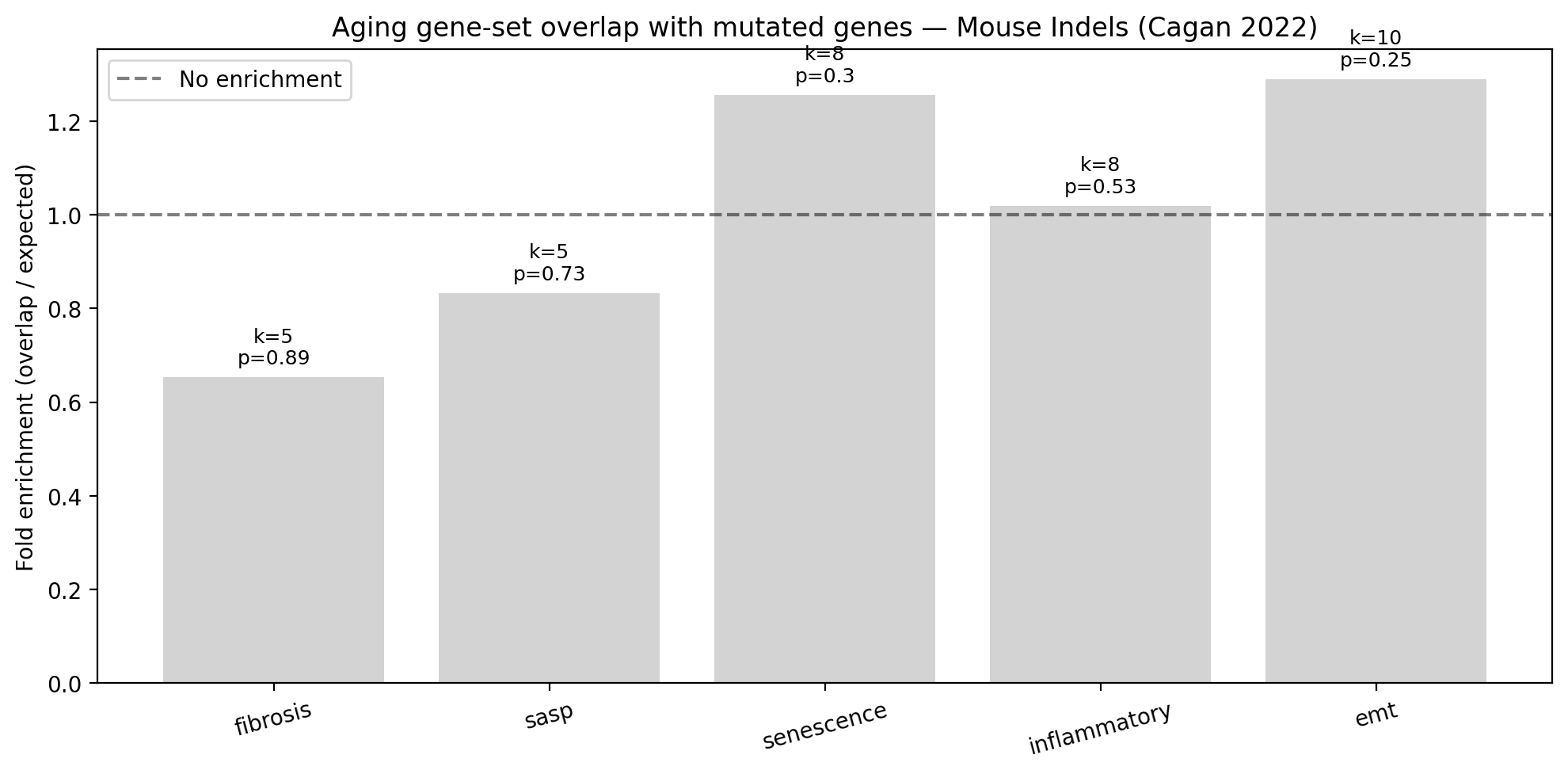

### human_cagan_vs_random_comparison.png

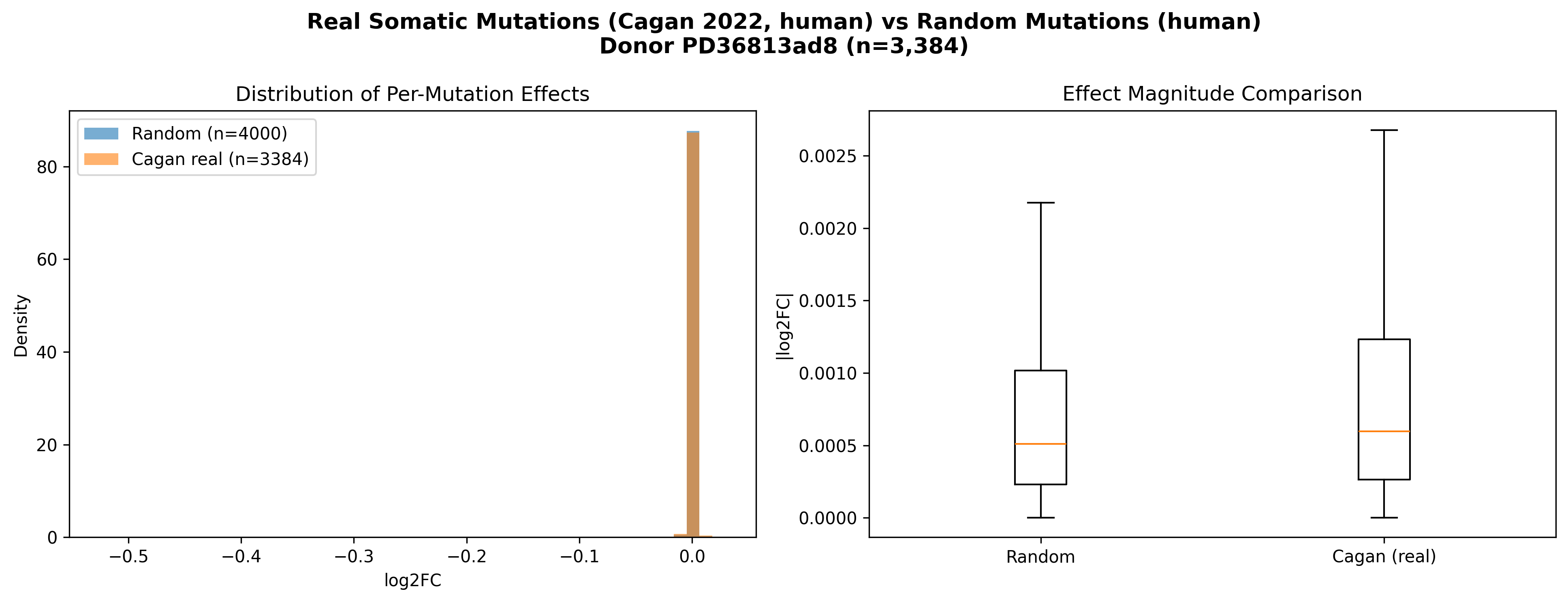

### human_log2fc_analysis.png

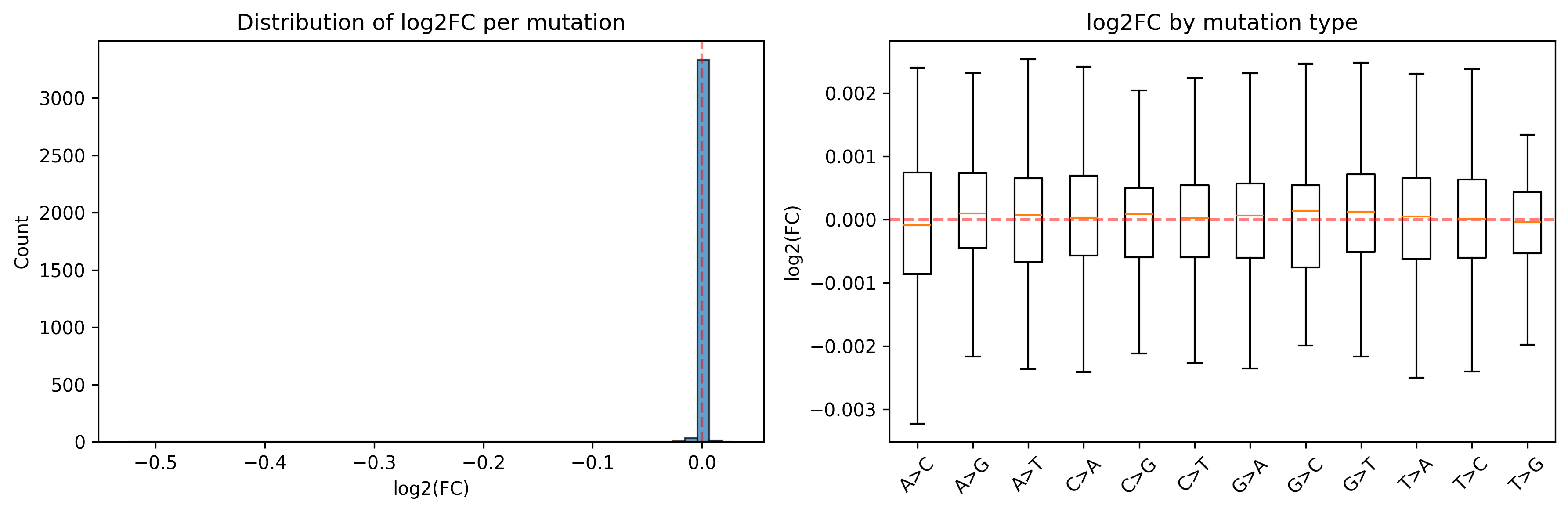

### indels_vs_subs_comparison.png

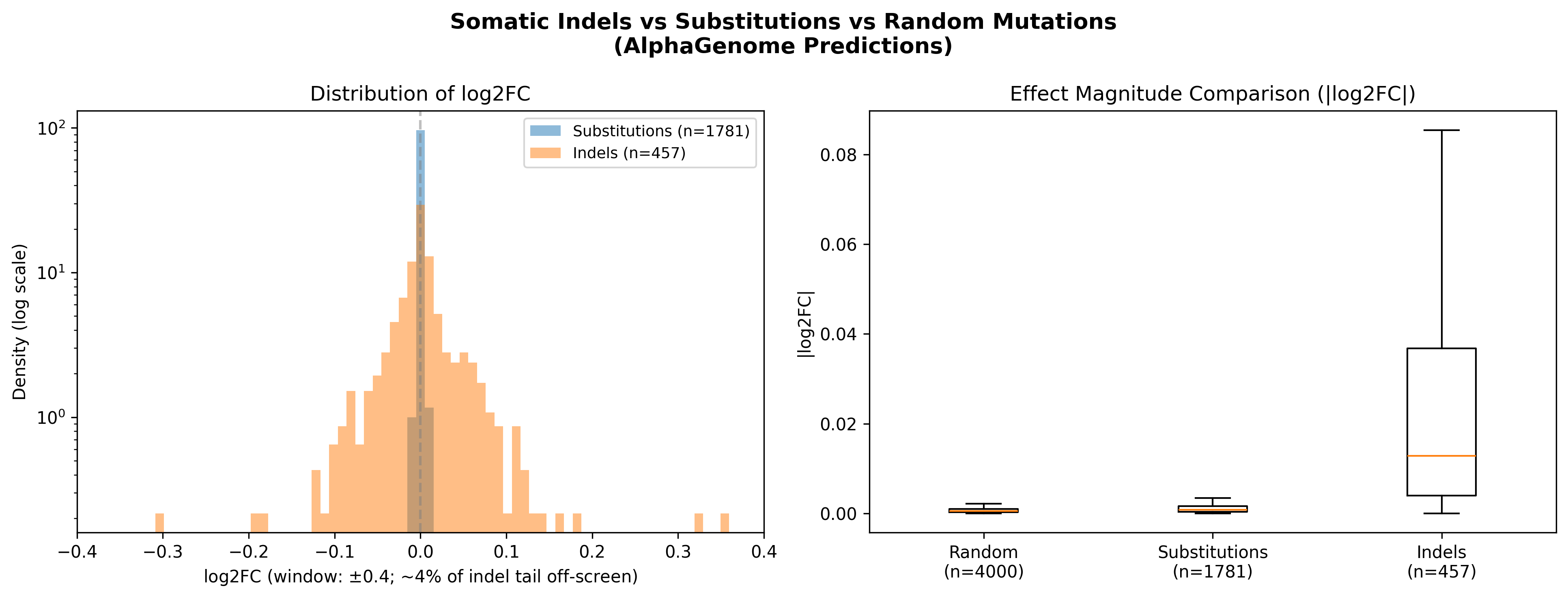

### log2fc_analysis.png

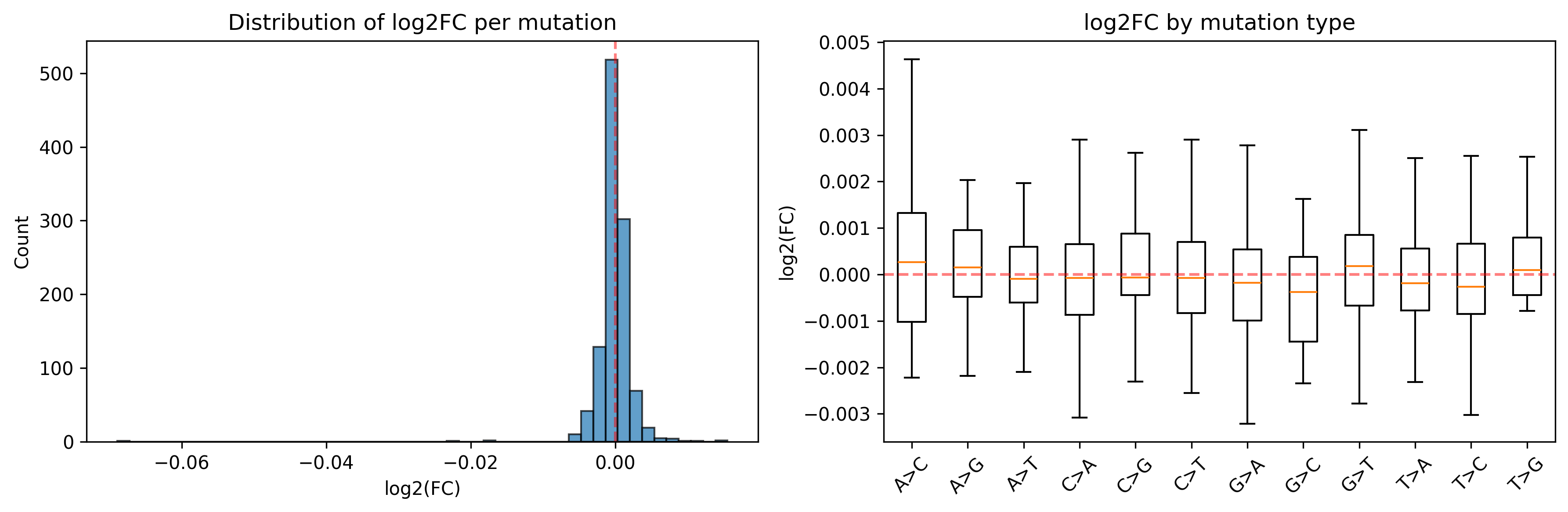

### mm10_colon_mutation_log2fc_distribution.png

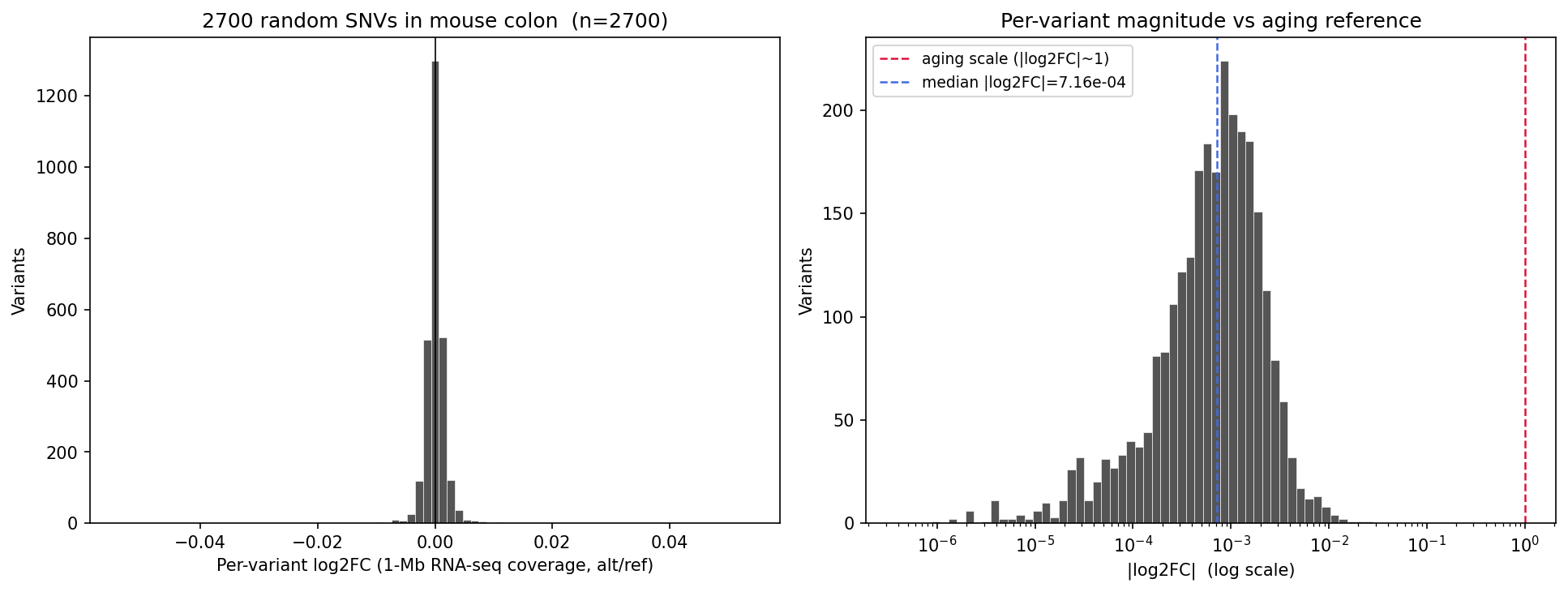

### mm10_colon_mutation_simulation_plots.png

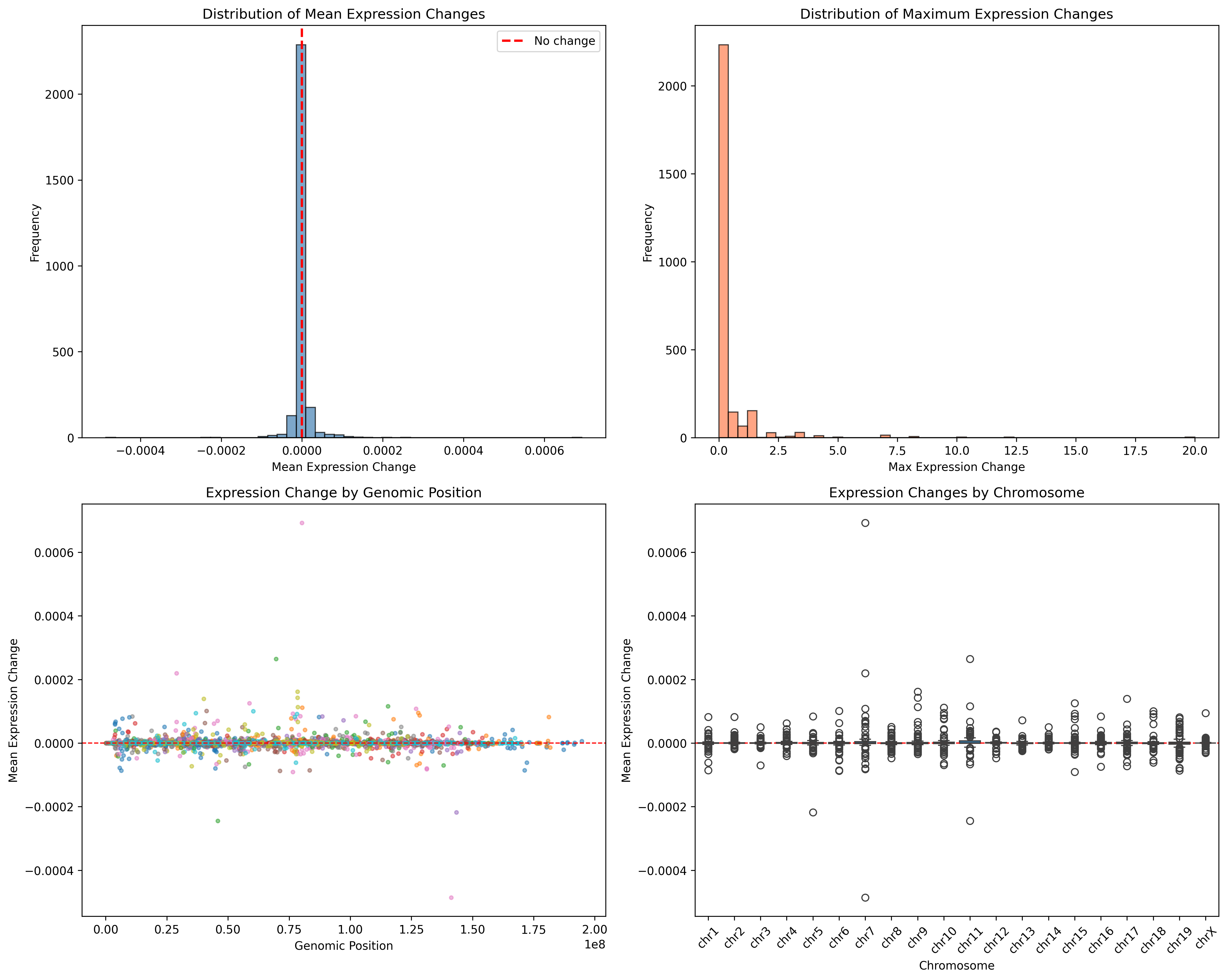

### mm10_combined_effects_analysis.png

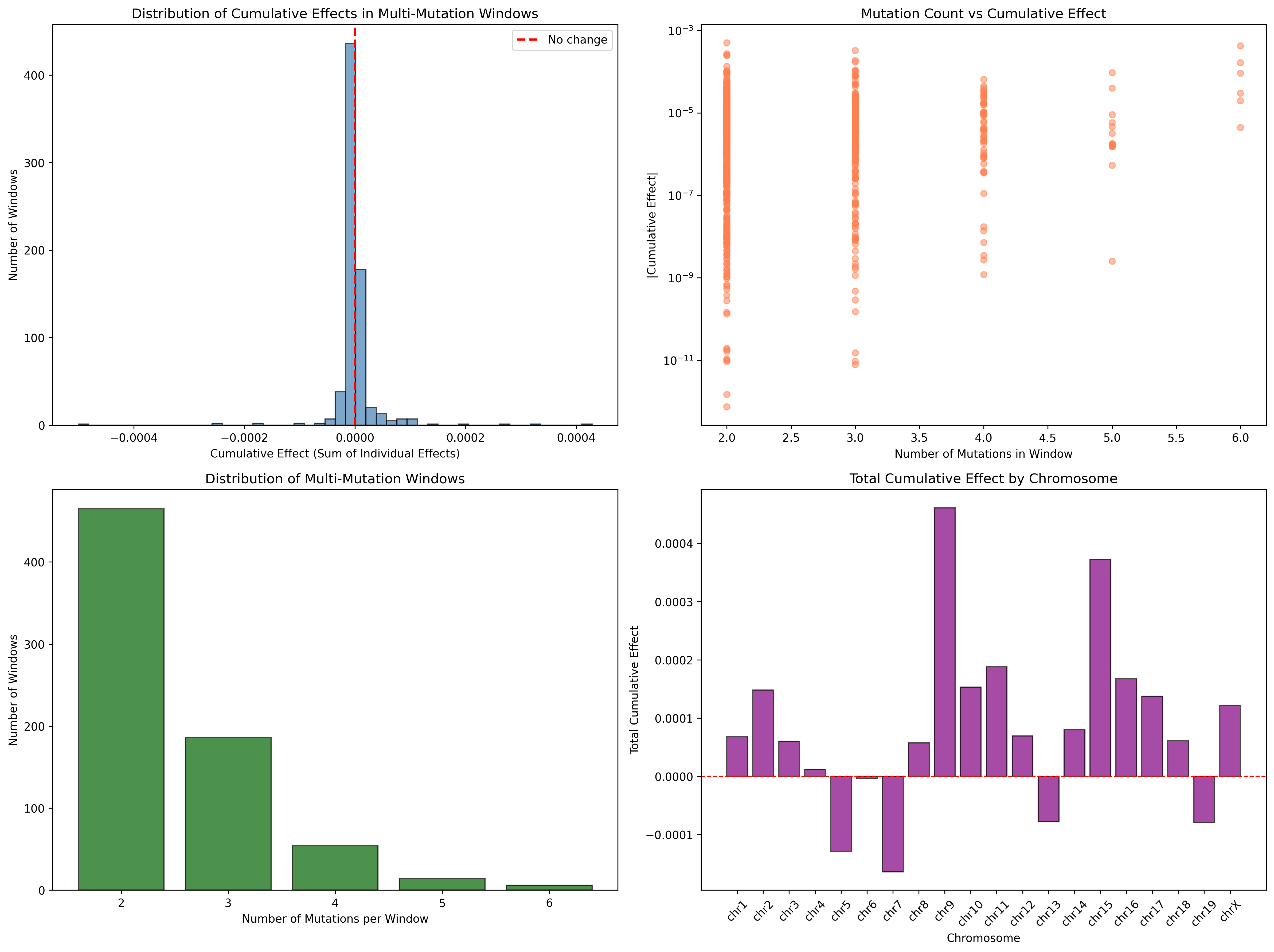

### monte_carlo_log2fc_extrapolation.png

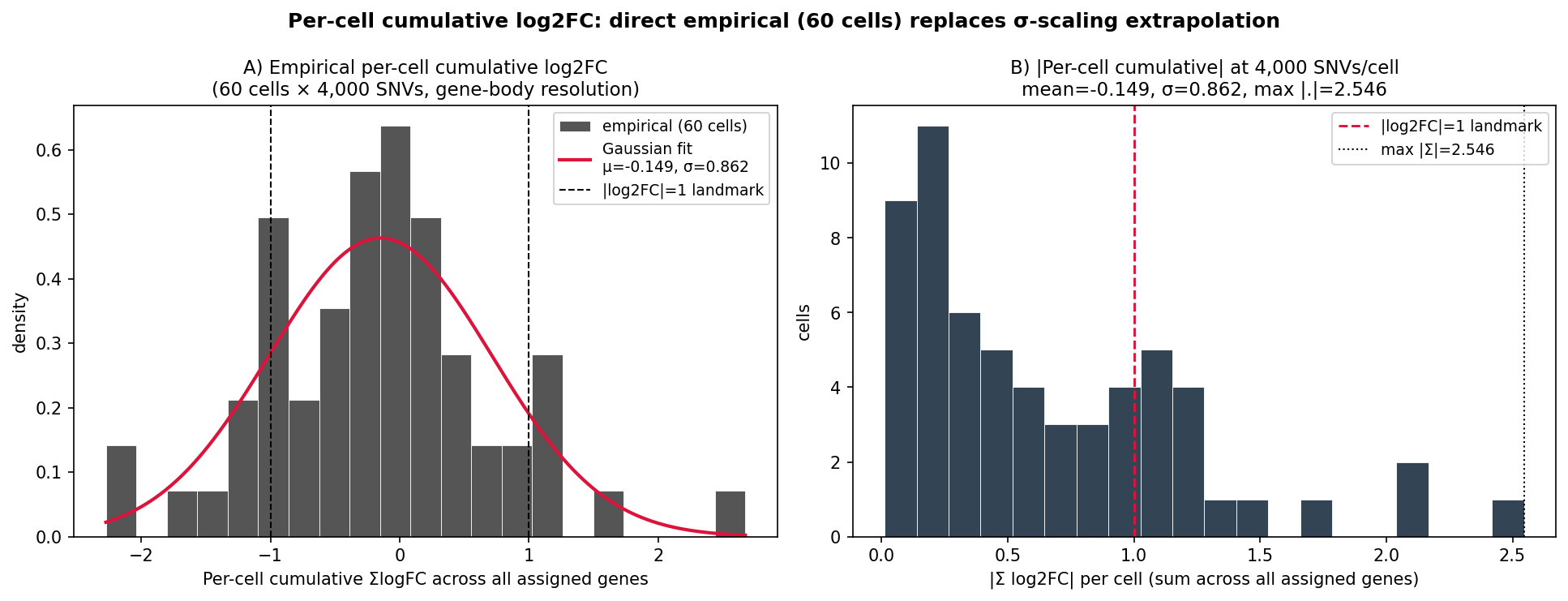

### PD36813ac4_human_indel_log2fc_analysis.png

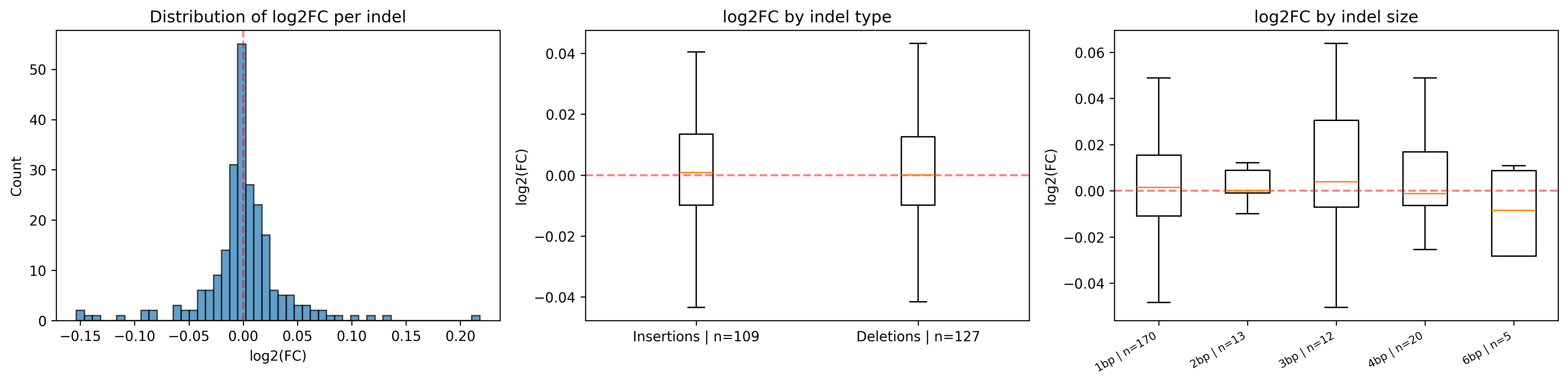

### PD36813ac4_human_indels_log2fc_paper_panel.png

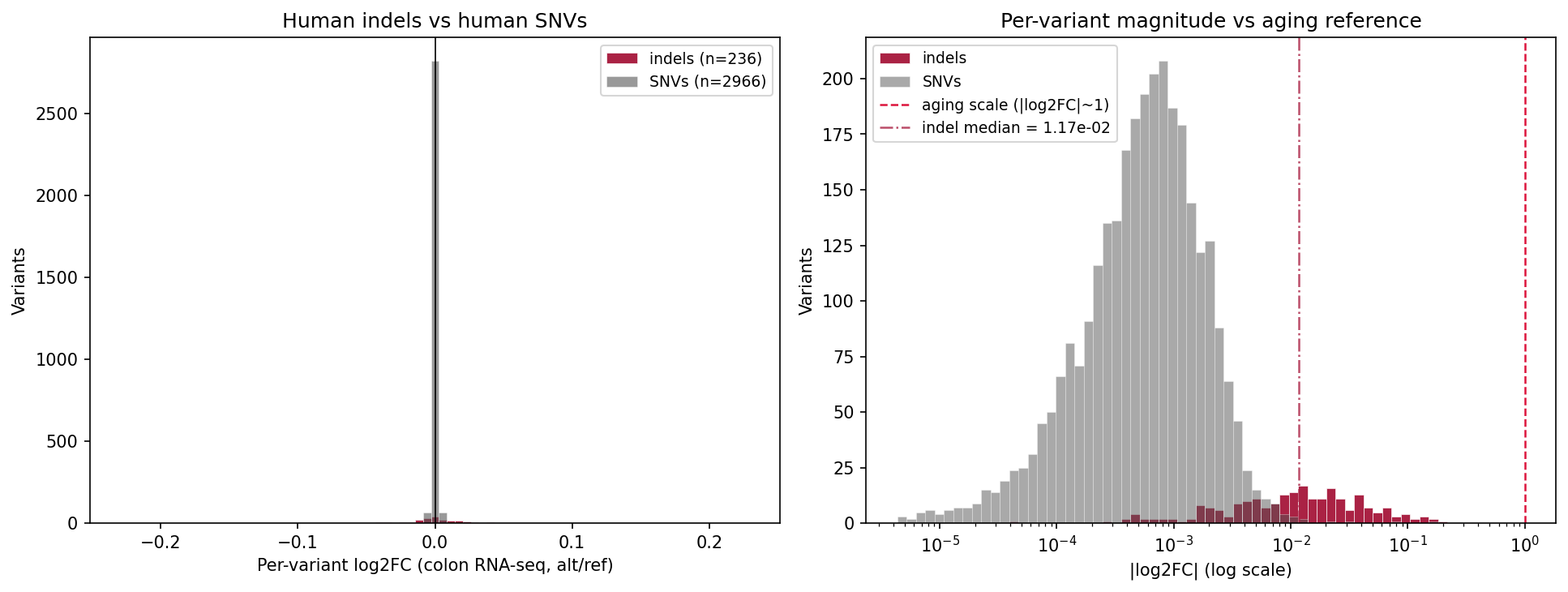

### PD36813ac4_human_indels_vs_baselines.png

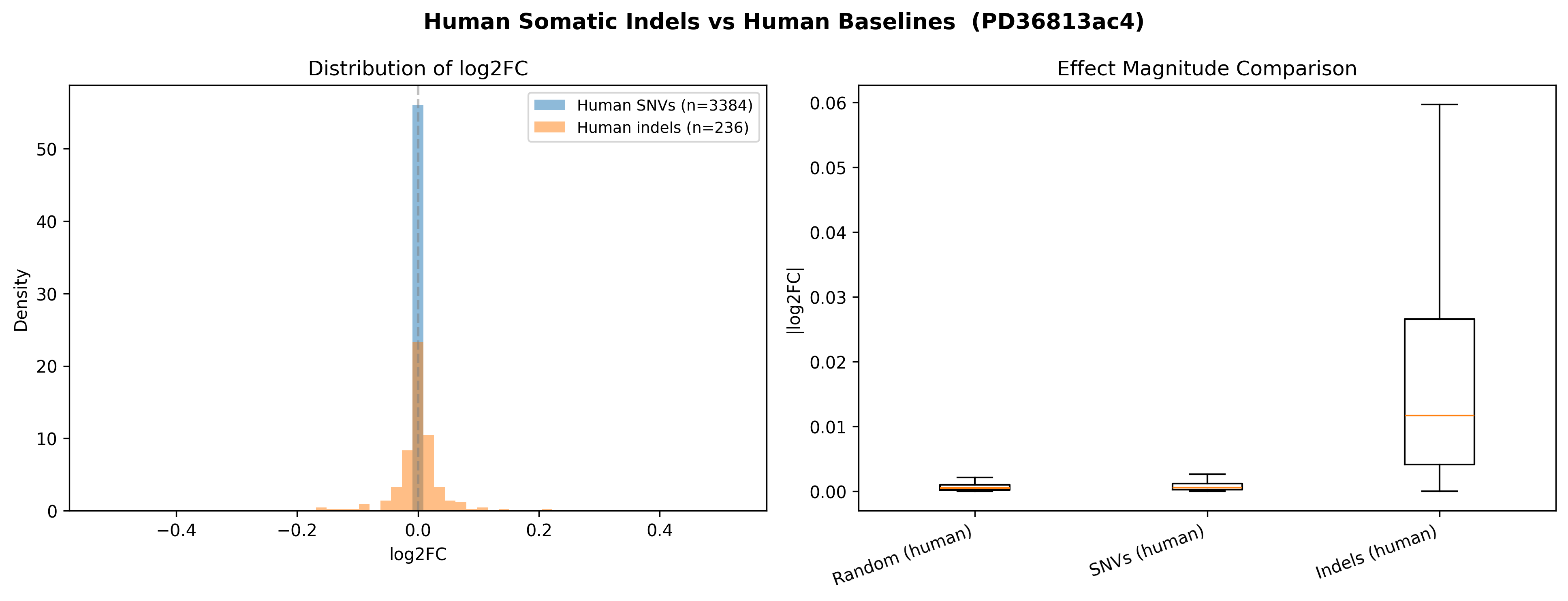

### per_gene_enrichment_Mouse_Indels_Cagan_2022.png

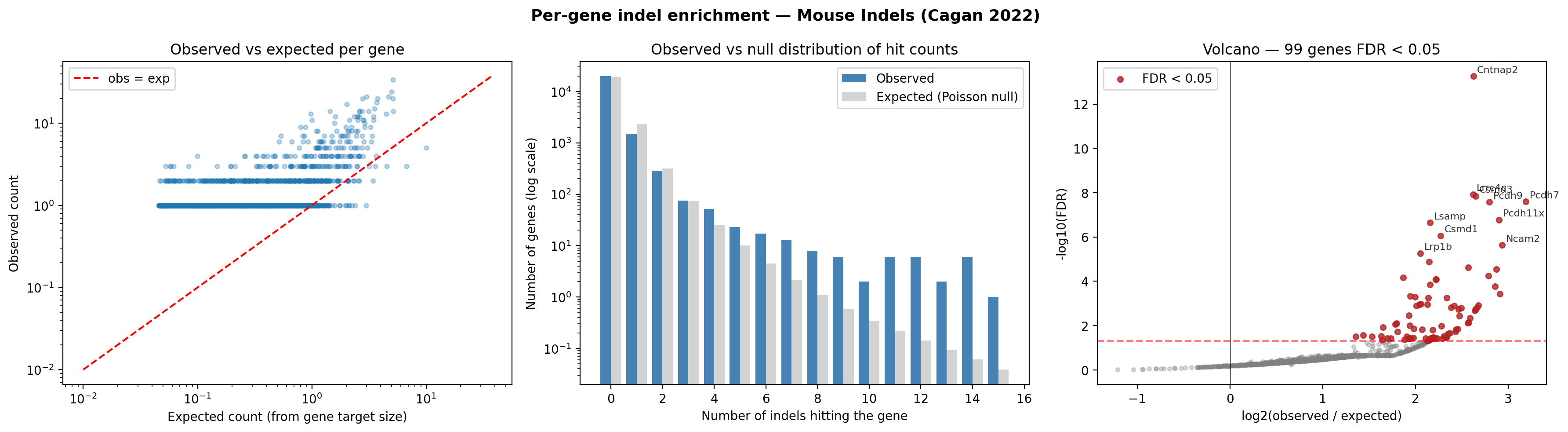

### pseudobulk_60cells_vs_aging.png

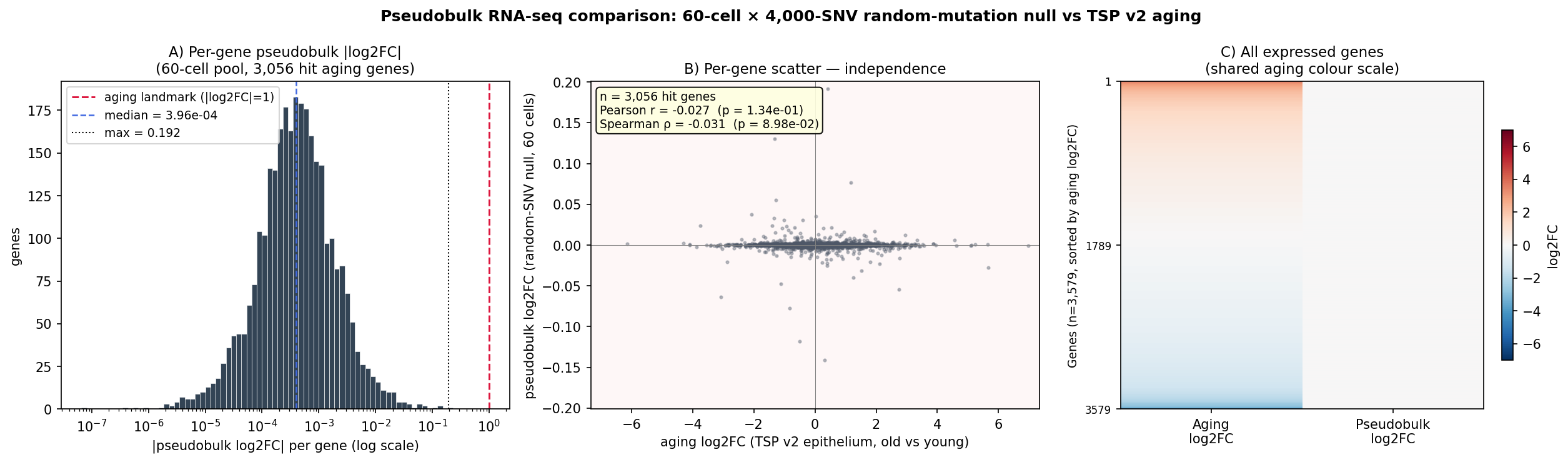

### snv_vs_indel_comparison.png

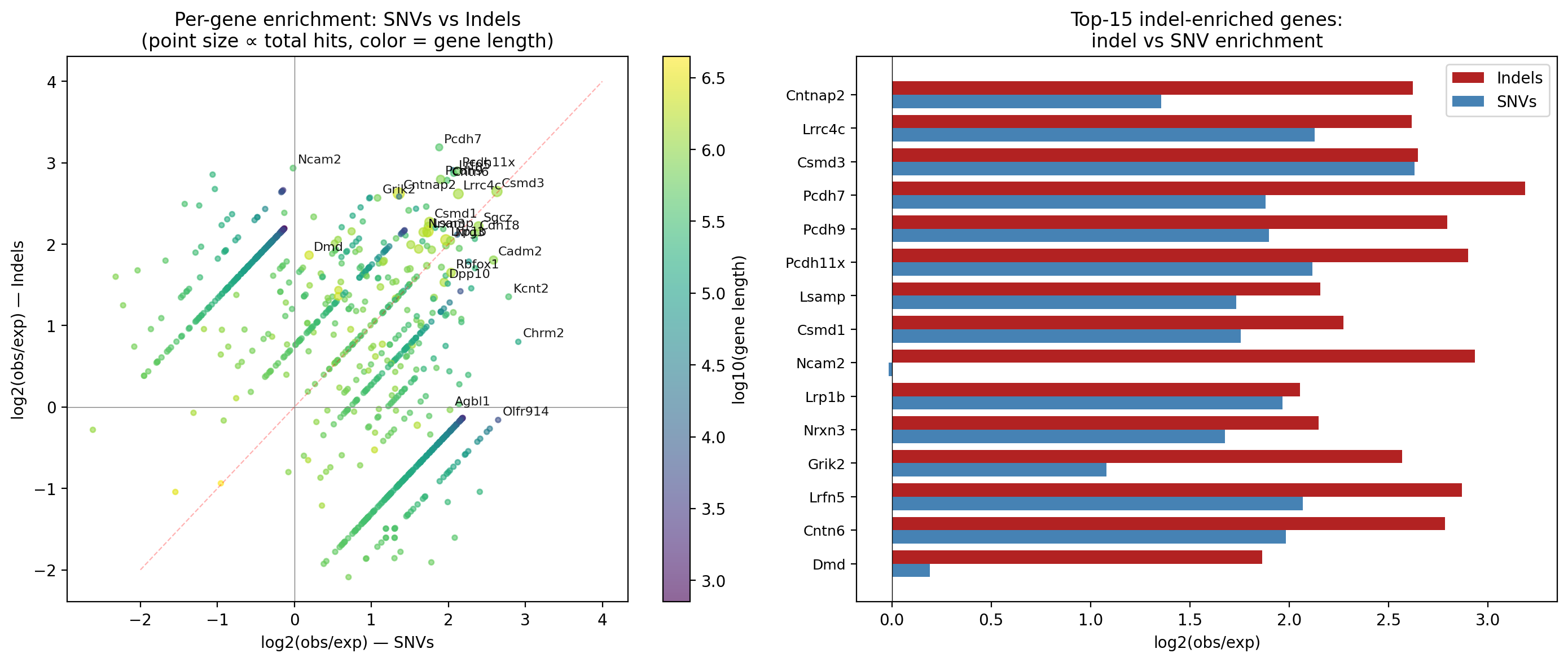
